# Early-life obesogenic environment integrates immunometabolic and epigenetic signatures governing neuroinflammation

**DOI:** 10.1101/2023.04.21.537874

**Authors:** Perla Ontiveros-Ángel, Julio David Vega-Torres, Timothy B. Simon, Vivianna Williams, Yaritza Inostroza-Nives, Nashareth Alvarado-Crespo, Yarimar Vega Gonzalez, Marjory Pompolius, William Katzka, John Lou, Fransua Sharafeddin, Ike De la Peña, Tien Dong, Arpana Gupta, Chi T. Viet, Marcelo Febo, Andre Obenaus, Johnny D. Figueroa

## Abstract

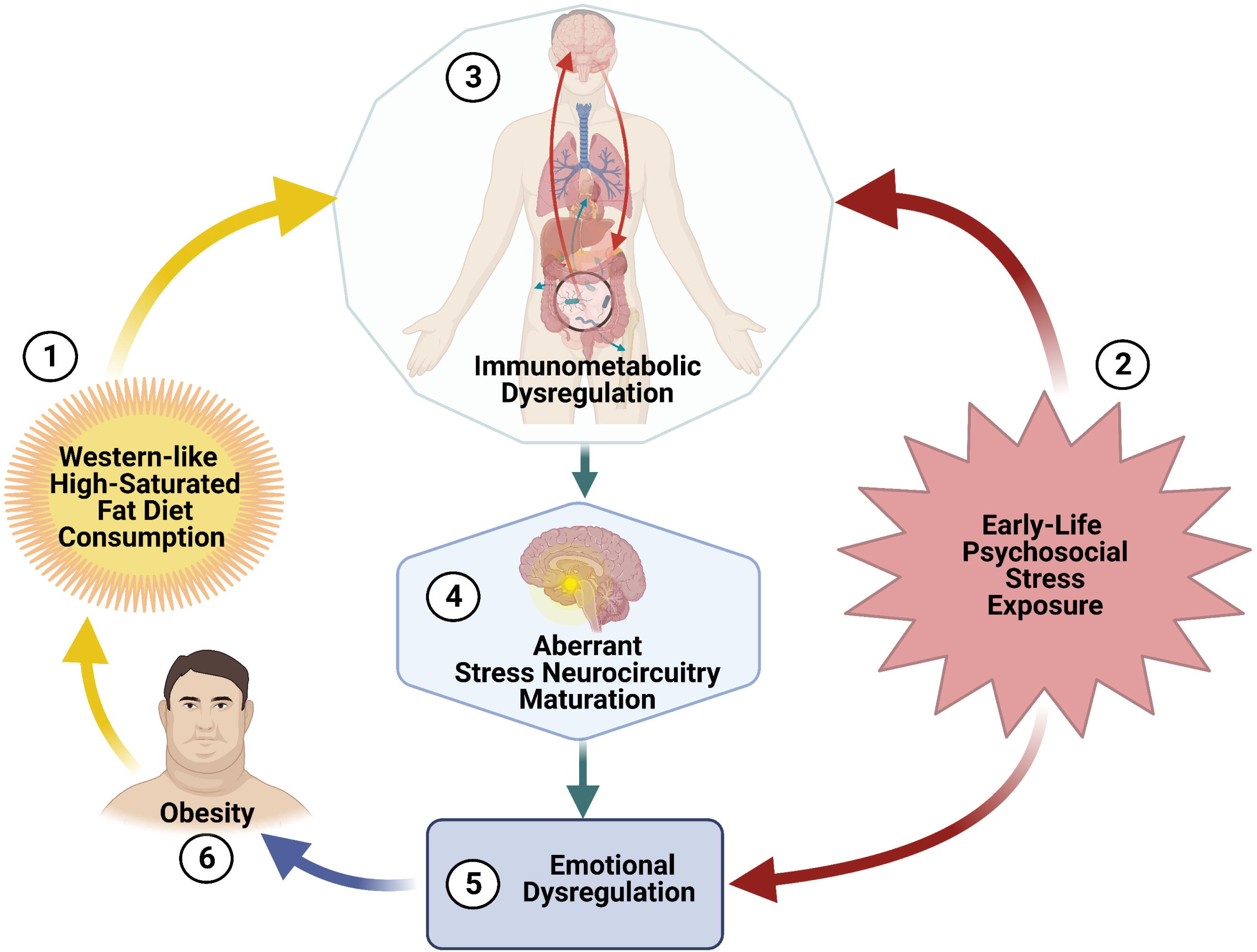
Early life trauma and obesogenic diet effects of feeding control. Consumption of a Western-like high-saturated fat diet (WD, 42% kcal from fat) during adolescence in combination with (2) Exposure to early-life psychosocial stress leads to (3) changes in brain neurocircuitry and metabolic dysregulation. These alterations lead to (4) stress susceptibility, (5) emotional and feeding dysregulation, and (6) obesity. Dysregulation of feeding control and obesity leads to increased hedonic feeding and engages individuals in a cycle of aberrant feeding behaviors.

**Background:** Childhood overweight/obesity is associated with the development of stress-related psychopathology. However, the pathways connecting childhood obesity to stress susceptibility remain poorly understood. Here, we used a systems biology approach to determine linkages underlying obesity-induced stress susceptibility.

**Methods:** Sixty-two (62) adolescent Lewis rats (PND21) were fed for four weeks with a Western-like high-saturated fat diet (WD, 41% kcal from fat) or a matched control diet (CD, 13% kcal from fat). Subsequently, a group of rats (*n* = 32) was exposed to a well-established 31-day model of predator exposures and social instability (PSS). The effects of the WD and PSS were assessed with a comprehensive battery of behavioral tests, DTI (diffusion tensor imaging), NODDI (neurite orientation dispersion and density imaging), high throughput 16S ribosomal RNA gene sequencing for gut microbiome profiling, hippocampal microglia morphological and gene analysis, and gene methylation status of the stress marker, FKBP5. Parallel experiments were performed on human microglial cells (HMC3) to examine molecular mechanisms by which palmitic acid primes these cells to aberrant responses to cortisol.

**Results:** Rats exposed to the WD and PSS exhibited deficits in sociability indices and increased fear and anxiety-like behaviors, food consumption, and body weight. WD and PSS interacted to alter indices of microstructural integrity within the hippocampal formation (subiculum) and subfields (CA1). Microbiome diversity and taxa distribution revealed that WD/PSS exposure caused significant shifts in the diversity of gut dominant bacteria and decreased the abundance of various members of the *Firmicutes* phylum, including *Lachnospiracae NK4A136.* Interestingly, the WD and PSS synergized to promote hippocampal microglia morphological and gene signatures implicated in neuroinflammation. These alterations were associated with changes in the microbiome, and in the expression and methylation status of the corticosterone receptor chaperone rat gene *Fkbp5*. HMC3 responses to cortisol were markedly disrupted after incubating cells in palmitate, shown by morphological changes and pro-inflammatory cytokine expression and release. Notably, these effects were partly mediated by the human FKBP5 gene.

**Conclusions:** The combination of psychosocial stress and poor diet during adolescence has a deleterious synergistic impact on brain health. This study enhances our understanding of mechanisms and adaptations by which obesogenic environments shape the maturational trajectories of common neurobiological correlates of resilience.

**Highlights:**

- Obesogenic diet consumption during adolescence leads to stress-induced anxiety-like behaviors in rats.
- Exposure to an obesogenic environment during adolescence alters indices of hippocampal microstructural integrity.
- Obesogenic diet and chronic stress promote selective gut microbiota dysbiosis.
- Obesogenic diet and chronic stress synergize to expand putative pro-inflammatory microglia populations in the CA1 subfield of the hippocampus.
- Obesogenic diet and chronic stress influence hippocampal *Fkbp5* gene methylation status at specific sites.
- FKBP5 integrates microglial pro-inflammatory signals under obesogenic conditions.

## Introduction

Childhood obesity is a severe medical problem affecting over 340 million children and adolescents worldwide.^1^ While obesity is a complex disease, consuming Western-style diets (WD) significantly contributes to the global pediatric obesity epidemic.^2^ Obesity and consuming imbalanced obesogenic WD, typically rich in saturated fats and simple sugars, have emerged as risk factors for developing psychiatric disorders.^3–7^

Recently, the prolonged COVID-19-related lockdowns, social instability, and uncertainty stress have negatively impacted dietary practices and increased childhood stress and obesity prevalence rates.^8–10^ Exposure to these impinging obesogenic environmental factors during childhood may have profound long-term effects on the brain and shape adult behavior and health. Thus, there is a clear need to delineate biological pathways and mechanisms linking early obesogenic environments to adult mental health outcomes.

While many complex linkages connect childhood obesity to psychopathology, stress susceptibility is a significant pathway.^11^ There is extensive literature describing shared pathological pathways in obesity and stress-related psychiatric disorders.^4, 12^ Obesity influences inflammatory mediators, the sympathetic nervous system, and the hypothalamic-pituitary-adrenal (HPA) axis. Recent investigations highlight the potential involvement of extra-hypothalamic brain areas in obesity-mediated stress susceptibility.^13–16^

Reduced hippocampal volume has emerged as a prominent anatomical endophenotype in human obesity.^13–16^ The hippocampus is critical in terminating the physiological stress responses via feedback inhibition of the hypothalamic-pituitary-adrenal (HPA) axis.^17–19^ Thus, alterations in this brain region may contribute to the stress-related psychiatric co-morbidities associated with obesity. In line with these studies, we reported that access to an obesogenic diet during adolescence: 1) reduces the volume of the hippocampus,^20^ 2) impairs the maturation of corticolimbic circuits,^21^ 3) enhances behavioral vulnerabilities to psychosocial stressors,^20–22^ and 4) increases oxidative stress and neuroinflammation,^23^ even in the absence of an obesogenic phenotype.^24^ However, less is known about the mechanisms contributing to hippocampal impairments that may result in stress susceptibility in individuals exposed to obesogenic environments during adolescence.

Obesity is associated with reduced synaptic density in humans^25^ and rodents.^26, 27^ Several genes implicated with human obesity are critical in sustaining synaptic integrity, supporting the evidence for links between synaptic integrity and the disease.^28^ Microglia are the innate immune cells of the brain. These highly dynamic cells play significant roles in axonal remodeling, synaptic pruning,^29^ and hippocampal tissue volume.^30^ Microglia are highly responsive to stress,^31, 32^ obesogenic diets,^24, 26, 27^ and are involved in the behavioral outcomes associated with obesity.^26, 33, 34^ However, the underlying molecular mechanisms regulating microglia activities in obesogenic environments remain poorly understood.

Microglia respond to local signals within the brain and receive input from the periphery, including dietary lipids, cytokines, hormones, and other immune modulators. Obesogenic dietary factors like palmitate and stress hormones such as glucocorticoids control microglia morphology and functions.^35–39^ In addition, preclinical findings suggest that the gut microbiome plays a pivotal role in regulating microglial maturation and function,^40–42^ which can impact brain maturation.^43, 44^ Because both obesity and stress are also associated with lasting epigenetic changes, a possible hypothesis is that access to an obesogenic diet during the critical maturational period of adolescence could confer stress susceptibility later in life through epigenetic effects on microglial molecules involved in immunometabolic function.

An essential modulator of these interactions is FKBP5,^45–48^ which codes for the FK506-binding protein 5-51 (FKBP51) and has emerged as a promising drug target for stress-related disorders.^49–51^ New studies demonstrate a robust association between the FKBP5/FKBP51 and microglia activities^52^ and overall hippocampal structure and function.^53–55^ Thus, FKBP51 may play a role in predisposing obese individuals to stress by regulating microglia and the structural integrity of the hippocampus.

This study builds on prior research from our group and investigated the impact of obesogenic environments on adolescent brain maturation. Adolescent access to an obesogenic diet and chronic psychosocial stress proved an excellent method to identify the lasting consequences of these environmental factors on sociability, anxiety-like behavior, startle responses, and fear learning. In addition to alterations in hippocampal structure, microbiome composition, and neuroinflammation, the obesogenic diet and stress interacted to influence *FKBP5* expression and methylation states in the hippocampus. *In vitro* analyses demonstrated that palmitate primes microglia to altered pro-inflammatory responses to cortisol. This study has translational and clinical relevance that may lead to alleviating symptoms in individuals who have been subsumed into the insidious cycle of dietary obesity and chronic psychosocial stress. Our results demonstrate that exposure to obesogenic conditions during adolescence adversely affect behavior, brain structure, and stress-related biomarkers.

## Methods

### Animals

Experimental procedures involving animals were performed in compliance with protocol No. 20-171, approved by the Institutional Animal Care and Use Committee (IACUC) at the Loma Linda University Health School of Medicine. This protocol follows institutional regulations consistent with the National Institutes of Health Guide for the Care and Use of Laboratory Animals and the ARRIVE guidelines for reporting animal research.^56^ Great efforts were made to reduce the animal number and minimize animal suffering and discomfort. Female Lewis dams with 8 male pups (postnatal day 15, PND15) were obtained from Charles River Laboratories (Portage, MI, USA). Upon arrival, female rats were housed with their pups with *ad libitum* access to food and water. The rats were kept in standard housing conditions (12-h light/dark cycle with lights on at 7:00 AM, 21 ± 2°C, and relative humidity of 45%). Adolescent male pups (PND21) were weaned, matched across diet groups based on their body weight and startle responses, and paired housed with *ad libitum* access to their assigned diet and water for the duration of the study. The body weights were recorded weekly, and food consumption was quantified at least twice weekly.

The food consumed was measured at the same time of the day (0900-1100 h). Uneaten food (cage top and spillage) was weighted and recorded, and fresh food was added to the top of the cage. Food intake was expressed as kilocalories (kcal) per day.

### Study Design

Sixty-two (62) adolescent Lewis rats (PND21) were weaned, carefully matched by body weight and acoustic startle reactivity, and randomized to receive either a Western-like high-saturated fat diet (*WD*, *n* = 30) or an ingredient-matched purified control diet (*CD*, *n* = 27). Five female rats were identified and removed from the study. Any male rat that interacted with a female was also removed from the study. The final total sample number was 52: CD *n* = 24 and WD *n* = 28, all males. The WD (*Product No. F7462*; Lot No. 248348) is 4.6 kcal/gram and 41.4% kcal from fat) and the CD (*Product No. F7463*; Lot No. 248347) 3.8 kcal/gram and 16.5% kcal from fat) were obtained from Bio-Serv (Frenchtown, NJ, USA). The macronutrient composition and fatty acid profiles are detailed in previous studies and summarized in Supplemental Table 1.^57, 58^ The rats consumed the diets for 4 weeks were further subdivided into four groups using random assignments (at PND51): **(1)** control diet, unexposed (CDU); **(2)** control diet, exposed (CDE); **(3)** Western diet, unexposed (WDU); **(4)** Western diet, exposed (WDE); *n = 14* each. Exposed groups underwent a well-established 31-day psychosocial stress (PSS) model, including predator encounters and social instability.^59^ The effects of WD and PSS were assessed using a comprehensive battery of standard behavioral tests and an array of stress and inflammation markers measures. A day after completing the behavioral assessments, a subset of rats was fasted for 12 hours to investigate the metabolic consequences of the experimental perturbations. All the rats were then euthanized, and plasma and brain tissue were collected. Experiments were performed in 4 independent rat sub-cohorts (each with a similar number of animals and containing all study groups). A complete timeline of the study is included in **Figure 1**.

**Figure 1.**
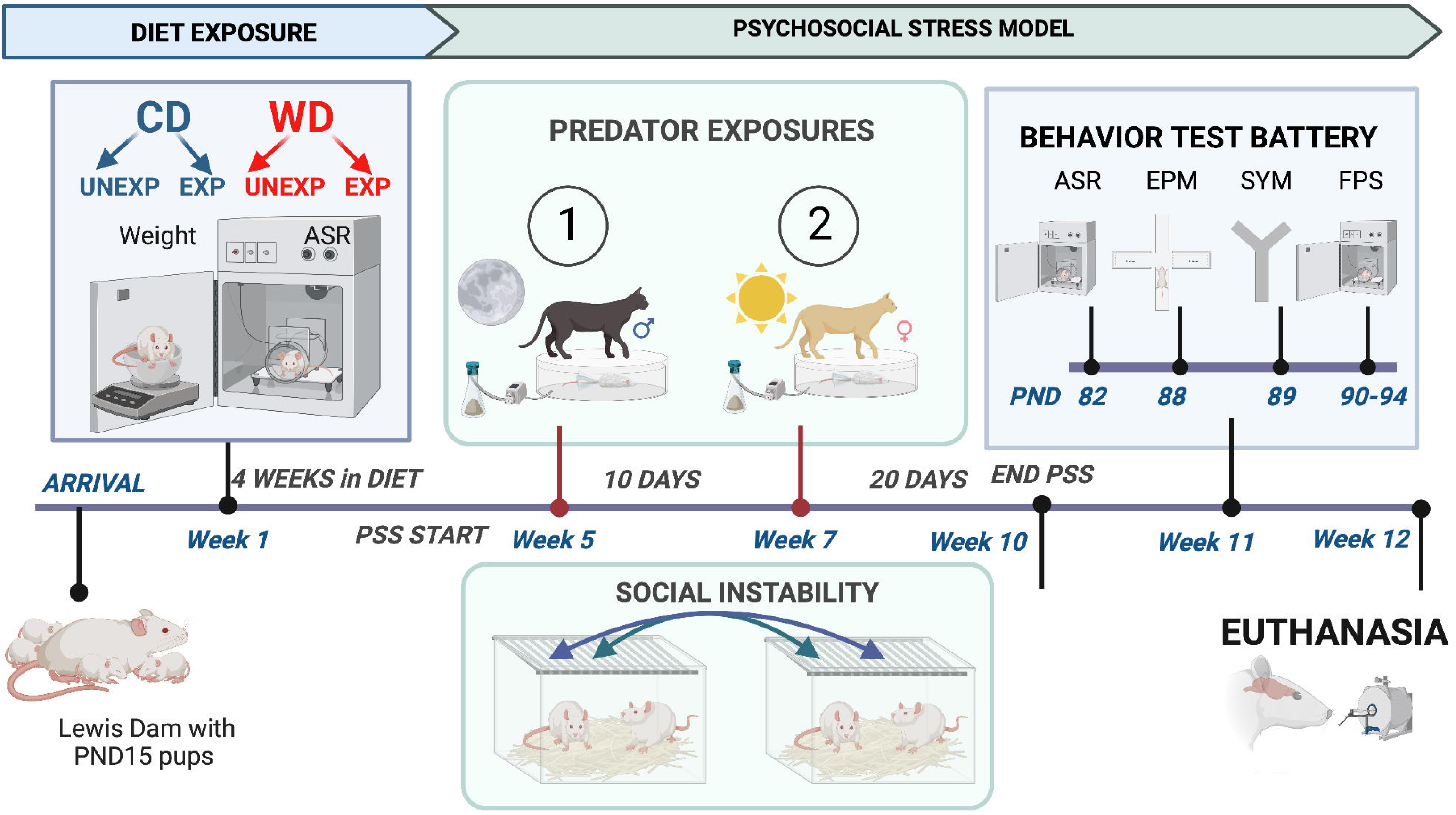
Experimental timeline and procedures. Study timeline including weaning and group assignment, diet exposure for four weeks, psychosocial stress model including two predator exposures and social instability, and behavioral evaluations. Abbreviations: CD control diet, WD western diet, EPM elevated plus maze, SYM Social Y Maze, FPS fear-potentiated startle, ASR Acoustic Startle Reflex, CD Control Diet, EPM Elevated Plus Maze, EXP Exposed, FPS Fear Potentiated Startle, PND Postnatal Day, PSS Psychosocial Stress, SYM Social Y Maze, UNEXP Unexposed, WD Western Diet.

### Psychosocial stress model (PSS)

To evaluate whether access to an obesogenic diet disrupts adolescent trauma’s behavioral and physiological consequences, we adapted a model of post-traumatic stress disorder (PTSD). The psychosocial stress (PSS) model includes trauma-inducing exposures to a natural predator and social instability.^59^

### Predator exposure

A set of domesticated juvenile cats were used (one male and one female). Cats were transported to the Animal Care Facilities 30 minutes before the start of each exposure. Retrieval and transportation of the cats were kept consistent and included the collection of soiled cat litter (3-day old), sifted for stools. The cat’s litter was weighted before and after, allowing the cats to use it. This measure allowed us to quantify and normalize the approximate urine amount, facilitating reproducibility between subcohorts.

The rats in the exposed groups were immobilized in plastic DecapiCones (Cat. No. NC9679094, Braintree Scientific; MA, USA) and placed in a perforated wedge-shaped plexiglass pie cage designed to deliver aerosols (Cat. No. RPC-1 AERO, Braintree Scientific; MA, USA; Diameter 41cm × Height 6.75 cm). The container with the soiled cat litter was connected to a nebulizer to aid in aerosol delivery and placed in the metal enclosure (91.4 cm × 58.4 cm × 63.5 cm, Amazon Basics, Amazon, USA) with the freely moving cat for a duration of 1 hour. A total of two (2) exposures were performed, separated by 10 days. The first exposure was performed during the light cycle (between 0800 h and 0900 h), and the second exposure took place during the dark cycle (between 2000 h and 2100 h). Unexposed controls were transported and kept in a separate behavior room for 1 h, once during the light cycle and once during the dark cycle.

### Social instability

The rats in the PSS group were exposed to unstable housing conditions for 31 days, starting on the day of the first cat exposure, and accomplished by changing cage partners daily, at random times of the day. The exposed groups were never housed with the same partner on consecutive days during the social instability protocol. The rats were returned to their original partner at the end of the protocol.

Rats in the unexposed group were kept undisturbed and housed with their same partner for the duration of the study. This model provides ecological and environmental validity, mimicking intense psychosocial stress with lasting consequences in the brain, behavior, and physiology.^59–61^

### Behavioral Test Battery

We use a test battery to investigate behavioral stress responses in rats that consume obesogenic diets (Figure 1). Our standard battery includes acoustic startle measures (and its plasticity: habituation and potentiation), elevated plus-maze, and the Y maze.^57, 58, 62–64^ In this study, we incorporated a variation of the Y maze that captures social behaviors. Behavioral tests were generally performed from least invasive to most invasive to minimize carryover effects. Behavioral measures were recorded from PND82-94. In our hands, this sensitive and comprehensive test battery is a powerful tool to unmask how obesogenic conditions impact behavior.^57, 58, 62–64^

### Acoustic Startle Reflex (ASR)

ASR measurements were performed in the SR-LAB™ Startle Response Dual Cabinet System (Cat. No. 2325-0401; San Diego Instruments, San Diego, CA, USA). The SR-Lab system provides sound-proofed and well-ventilated enclosures that facilitate the reproducible assessment of the acoustic startle. The auditory stimuli intensities and response sensitivities were calibrated before commencing the experiments with a dynamic calibration system (Calibrator Serial No. 006340; San Diego Instruments, San Diego, CA, USA) to ensure comparable stabilimeter sensitivity across test chambers. Sound levels were calibrated using a dB(A) scale calibrator (Cat. No. 407730, Extech Instruments, Nashua, New Hampshire, USA).

The rats were tested in pairs in neighboring test chambers. The rats are placed inside a clear nonrestrictive tubular plexiglass stabilimeter. A high-frequency loudspeaker mounted 34 cm above the plexiglass cylinder produces the acoustic stimuli, with peak and average amplitudes of the startle response detected by a piezoelectric accelerometer. At the onset of the startling stimulus, 100 (1-ms) readings are recorded, and the average amplitude is used to determine the magnitude of the startle response (measured in arbitrary units). The testing sessions were 22 minutes long and started with a 5-min habituation period (background noise = 60 decibels, dB. The rats were then presented with 30 tones (10 tones at one intensity: 105 dB) using a 30-s inter-trial interval (ITI). The acoustic stimuli had a 20 ms duration. After each testing session, the chambers and enclosures were thoroughly cleaned, and the rats returned to their home cages. ASR responses were normalized by body weight and reported as Weight-corrected ASR (Maximum startle magnitude in mV, V_Max_, divided by body weight, in grams, at testing day).

### Fear-Potentiated Startle (FPS)

Hippocampal-dependent fear learning was assessed with a trace fear-potentiated startle (FPS) experimental paradigm. FPS responses were assessed inside the SR-LAB™ Startle Response Dual Cabinet System (Product No. 2325-0401; San Diego Instruments, San Diego, CA, USA). Each chamber contained a plexiglass enclosure with a calibrated piezoelectric motion sensor (Shock lever tester Product No. 2325-0358; San Diego Instruments, San Diego, CA, USA).

The startle reflex has a nonzero baseline and is graded in magnitude. Accelerometer readings were obtained at 1 ms intervals for 200 ms after the startle-inducing acoustic stimulus and recorded using SR-LAB startle software. Each enclosure contained grid floors capable of delivering foot shocks (Potentiated Startle Kit, Part. No. 2325-0404, San Diego Instruments, CA). Each FPS session started with a 5-min acclimation period (background noise = 60 dB). Fear conditioning was achieved by training the rats to associate a conditioned light stimulus (CS) with an aversive unconditional stimulus (US) consisting of a 0.6 mA foot shock.

Fear conditioning involved 10 CS + US pairing presentations. Each trial included a 4 s exposure to the light (CS) followed by a 500-ms foot shock, with 3 s between stimuli. Light-shock pairings were presented in a quasi-random manner (Inter-trial interval; ITI = 3-5 min). Fear learning was evaluated 24 h after the fear conditioning session. This cued fear acquisition testing session started with a brief 5 min acclimation period (background noise = 60 dB). The rats were first presented with 15 startle-inducing tones (leaders; 5 each at 90 dB, 95 dB, and 105 dB) delivered alone at 30 s ITI. These leader trials were used to familiarize the rats with the acoustic stimuli, facilitating more accurate startle reactivity measurement during the session. Subsequently, the rats were presented with 60 test trials. For half of these test trials, a 20 ms tone was presented alone (tone alone trials; 10 trials for each tone: 90 dB, 95 dB, and 105 dB). For the other half, the tone onset (20 ms duration) occurred 3 s after the offset of the 4 s light; 10 trials for each pairing: 90 dB, 95 dB, and 105 dB. The 60 trials were divided into 10 blocks of 6 test trials each, including 3 tone-alone trials and 3 light + tone trials. To conclude the testing session, the rats were presented with 15 startle-inducing tones (trailers; 5 each at 90 dB, 95 dB, and 105 dB) delivered at 30 s ITI. These trailer trials are commonly compared to the leader trials and used to investigate startle habituation during the testing session. All the trials in this session were presented in a quasi-random order (ITI = 30 s). The startle-eliciting tones had a 20 ms duration.

For each rat, we calculated the mean startle magnitude in the absence (tone alone startle) and in the presence of the learned light stimulus (light + tone startle) and the percent difference between these means. The rats were then exposed to two extinction-training sessions starting 24 h after the fear-learning testing session. This daily extinction training session consisted of 15 CS alone presentations (light without shock or noise bursts) with a duration of 3700 ms (ITI = 30 sec). One day after the second fear extinction training, we determined fear extinction acquisition using the same FPS session that was used to measure fear acquisition. We assessed fear learning and fear extinction learning by comparing the startle amplitude from light + tone trials (CS + US) relative to tone alone trials (US). FPS data were reported as the proportional change between US and CS + US, using the below formula.^65^

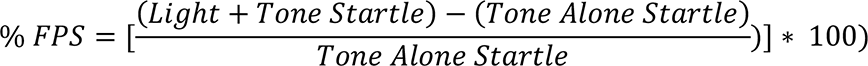

Conditioned fear responses from control animals were associated with potential hippocampal-related alterations. Increased potentiated startle responses (relative to controls) usually indicate heightened fear, while decreased FPS suggests deficits in fear acquisition, retention, and/or expression.

### Elevated plus maze (EPM)

The EPM was used as an additional proxy for anxiety-like behaviors, mobility, and risk-assessment behaviors. The near infrared-backlit (NIR) EPM consisted of two opposite open arms (50.8 × 10.2 × 0.5 cm) and two opposite enclosed arms (50.8 × 10.2 × 40.6 cm) (Cat No. ENV-564A; Med Associates Inc., Fairfax, VT). The arms were connected by a central 10 × 10 cm square-shaped area. The maze was elevated 72.4 cm from the floor. Behaviors were recorded in a completely dark room (< 10 lux). The rats were placed on the central platform facing an open arm and allowed to explore the EPM for 5 min. The apparatus was cleaned after each trial (70% ethanol, rinsed with water, and then dried with paper towels). The behaviors were recorded via a monochrome video camera equipped with a NIR filter and tracked using Ethovision® XT (Noldus Information Technology, Leesburg, VA). In rats, changes in the percentage of time spent on the open arms (OA) indicate changes in anxiety,^66^ and the number of closed arms (CA) entries is the best measure of locomotor activity.

### Social Y-Maze

Sociability was assessed with a modified Y maze implementing a protocol adapted from Vuillermot et al.,^67^ The test was previously used to measure rodent social interactions.^67, 68^ The Social Y-Maze (MazeEngineers, Stokie, IL) is a plexiglass Y-maze with a triangular center (8 cm sides) and three identical arms (50 cm × 9 cm × 10 cm; length × width × height). One arm functioning as the start arm, while the other two are equipped with rectangular wired mesh cages to hold the live conspecific or inanimate object (Multicolored LEGO® pieces, Billund, Denmark.

The protocol was conducted using one social interaction test trial without habituation training. The animal was placed into the start arm and allowed to explore the maze for 9 min freely. The unfamiliar sex and age-matched conspecific rat was placed into one rectangular wire mesh cage while the inanimate object was put into the other wire mesh cage.

The conspecifics and objects were counterbalanced across arms and treatment groups. A camera was mounted directly above the maze to record each test session. Video recordings were analyzed using Ethovision XT tracking software (Noldus Information Technology, Leesburg, VA). The maze was cleaned and allowed to dry between trials. The results were reported as the number of social interactions and ratio of interactions between conspecific and object.

### *Tissue* Collection

Animals from the four (4) experimental groups were randomly divided into two (2) main euthanasia groups (*n* = 57). The first group was designated for neuroimaging and neurohistological studies (*n* = 28). The second group was selected for molecular analyses (*n* = 29). All rats were deeply anesthetized with an intraperitoneal overdose injection of Euthasol (150 mg/kg Virbac, Fort Worth, TX). In the first group, rats were transcardially perfused with an isotonic solution at 9.25% in distilled water as a prewash and 4% paraformaldehyde (PFA; fixative, Cat No. 15714-S, Electron Microscopy Sciences, Hatfield, PA, USA) using the Perfusion Two ^TM^ Automated Pressure Perfusion System (Leica Biosystems, Buffalo Grove, IL).

Brains were preserved in the cranial vault after fixation and postfixed in 4% PFA, washed and stored at 4 °C in 0.1 M PBS with 0.05% Sodium azide until processing for imaging and histology. In the second group, rats were perfused transcardially with ice-cold PBS. These rats were rapidly decapitated, their brains isolated, and both the left and right hippocampi were dissected. Hippocampal tissue was immediately submerged in RNAlater™ Stabilization Solution (Cat. No. AM7021; Thermofisher, Waltham, MA, USA) to protect cellular RNA. Samples were stored at −80 °C until testing.

### Metabolic and Inflammatory Profiling Fecal Corticosterone

Fecal samples were collected at different time points across the experimental timeline to determine corticosterone (CORT) levels. Feces were collected 24 h after weaning (baseline), 24 h before and after each exposure to the predator, at the end of the PSS model, and after behavioral assessments. The last collection took place the day the rats were euthanized. Cages were changed 24 h before each collection. Fecal boli were collected and stored at −80 °C until extraction. The extraction protocol has been previously described.^69^ Samples were defrosted, weighted, and pulverized before the ethanol-based extraction. Subsequently, the levels of CORT were measured by ELISA (Cat. No. ADI-900-097; Enzo Life Sciences, Ann Arbor, MI, USA) according to the manufacturer’s instructions. Samples were diluted with kit assay buffer (1:5 dilutions) and ran in duplicates. The absorbance was measured at 405 nm with 570 nm correction using the SpectraMax i3X detection platform (Molecular Devices, Sunnyvale, CA). Corticosterone levels were determined as the percentage bound using a standard curve with a detection range between 20,000 - 31.25 pg/mL.

### Heart Rate and Blood Pressure Measurements

Before euthanasia, measurements of cardiovascular function were taken to determine the effect of WD and PSS on heart rate reactivity. Measurements were acquired using a rodent tail-cuff plethysmography blood pressure system (MRBP IITC Life Science, Woodland Hills, CA). Briefly, the rats were placed in cylindrical restraints with careful tail placement through a cuff/sensor endplate. After a period of acclimation (10 min), the rats were then placed in the testing chamber, where the tail blood pressure cuff/sensor was connected to the air and sensor ports. The tail was taped in place to minimize movement. Data were monitored for artifacts (movement) or variability in the detection range. Measurements that were tested outside the quality control parameters were repeated, with careful attention to tail position. System operation and data acquisition are performed with the Blood Pressure Data Acquisition Software (Part No. 31, IITC Life Science, Woodland Hills, CA). Data was exported using Excel (Microsoft, Redmond, WA, USA) and analyzed using GraphPad Prism version 9.4.1.681 for Windows (GraphPad Software, San Diego, California USA, www.graphpad.com. Measurements of heart rate were reported as beats per minute (bpm). Systolic and diastolic arterial pressure was reported in millimeters of mercury (mm of Hg).

### Fasting Blood Glucose

A group of rats was fasted for 16 h, and blood glucose levels were measured in anesthetized rats by cutting the tail tip (CDU, *n* = 13; CDE, *n* = 14; WDU, *n* = 15; WDE, *n* = 15). Glucose levels were measured from tail blood using a glucometer (OneTouch UltraMini® manufactured by LifeScan Inc, Milpitas, CA, USA) and reported as mg per dL. The rats were subsequently euthanized via a Euthasol injection (150 mg/kg, Virbac, Fort Worth, TX), transcardial perfusion, and decapitation.

### Bead-based Cytokine Detection

We used the multianalyte bead-based immunoassay LEGENDplex™ (Biolegend, San Diego, CA, USA) to determine inflammatory profiles. Three (3) mL of cardiac blood were collected from the right atrium into EDTA-coated tubes (Greiner, Cat. No. 454021, Fisher Scientific, Waltham, MA, USA). Whole blood samples were centrifuged at 1000 rpm for 15 min (with no brakes), and the plasma portion was collected. Samples were kept at −80°C until testing. Evaluation of circulating cytokine and chemokines (IL-10, IL- 1α, IL-6, IL-1β, TNF-α, IL-33, GM-CSF, CXCL-1, CCL2, IL-18, IL-12p70, IL-17a, and INF-γ) was performed according to the manufacturer’s recommendations. Samples were diluted at 1:4 with Assay Buffer and tested in triplicates. Cytokine levels were determined using antibodies for each analyte covalently conjugated to a set of microspheres in a carbodiimide crosslinking reaction and detected by a cocktail of biotinylated antibodies.

Following the binding of streptavidin–phycoerythrin conjugate, the fluorescent reporter signal was measured using a MACSQuant® Analyzer 10 (Miltenyi Biotech, Bergisch Gladbach, Germany). FCS files were evaluated with the LEGENDplex™ Qognit 53790 Data Analysis Suite (Biolegend, San Diego, CA, USA). A five-parameter logistic curve-fitting method was used to calculate concentrations from Median Fluorescence Intensity values. The results are reported as pg/mL.

### FKBP5 Methylation Profiling

Left hippocampal tissue was shipped to EpigenDx™ (Hopkinton, MA, USA) for qRT-PCR and targeted next-generation bisulfite sequencing (tNGBS).

### *In silico* Assay Design Process

Rat FKBP5 gene target panel NGS152 was screened for methylation percentage in various regulatory regions. Each regulatory element was carefully evaluated, and gene sequences containing the target of interested were acquired from the Ensembl genome browser annotated (**S Fig 16**). The target sequences were re-evaluated against UCSC genome browser for DNA repeat sequences, including LINE, SINE, LTR elements. Sequences containing repetitive elements, low sequence complexity, high thymidine content, and overall CpG density were excluded from the *in-silico* design process resulting in 23 out of 28 preselected assays per gene. Five (5) assays failed PCR optimization. A table in **Supplementary Figure 1 7** shows the list of the 23 optimized targets.

### DNA bisulfite modification and Methylation Analysis

Hippocampal tissue samples were lysed and bisulfite modified with EZ-96 DNA Methylation-Direct Kit (Zymo Research, CA, USA) following manufacturer recommendations. Bisulfite-modified DNA samples were eluted using M-elution buffer (20 µl in 46 µl respectively). Modified samples were amplified using simplex or multiplex PCRs. Reactions (20 µl) included 0.5 units of Qiagen HotStartTaq (Cat. No. 203207, QIAgen, Hilden, Germany), 0.2 µM primers, and 3 µl of bisulfite-treated DNA. All PCR products were verified and quantified using QIAxcel Advanced System (Cat. No. 9002123, QIAgen, Hilden, Germany. PCR products from the same sample were pooled and purified using QIAquick PCR Purification Kit Columns (Cat. No. 28104, QIAgen, Hilden, Germany). Assays were grouped in 5 batches at varying annealing temperatures (56°C, 59°C, and 61°C). Samples were run under cycling conditions: 95°C for 15 min; 45 x (95°C 30 s, Ta°C 30 s, 68°C 30 s); 68°C 5 min; 4°C ∞.

### Data Sequencing and Analysis

Enriched template positive EpigenDx™ library molecules were sequenced on the Ion S5™ sequencer (specs) using an Ion 530™ sequencing chip (Cat. No. A27212, ThermoFisher Scientific, Waltham, MA, USA). FASTQ files were aligned to a local reference database using open source Bismark Bisulfite Read Mapper with Bowtie2 alignment algorithm.^70^ Methylation levels were calculated in Bismark as the ratio of methylated reads over the total reads.

### Microbiome Sequencing and Analysis

Stool samples were collected and immediately stored at −80°C. DNA was extracted using the ZymoBIOMICS DNA Microprep Kit (Zymo Research, USA) per manufacturers’ instructions. The V4 region of the 16S ribosomal RNA gene was amplified by PCR using the 515F-806R primer set. Samples underwent 250 × 2 paired-end sequencing on an Illumina MiSeq (Illumina, San Diego, CA, USA).^71^ Raw fastq files were processed using the DADA2 pipeline in R, which assigns taxonomy using the SILVA 132 database and default parameters.^72^ After pre-processing in R utilizing DADA2, the data were incorporated into QIIME 2 version 2019.10.^73^

To remove sparse amplicon sequence variants (ASV), ASV were filtered if their relative abundance was below 0.0001%, like our previous publications.^74, 75^ Sequence depths ranged from 18,434 to 36,958 per sample, with a median value of 26,216. Alpha diversity was calculated using the Shannon index (a metric that combines both species richness and species evenness) and Chao1 index (a metric of species richness) through QIIME2. For alpha diversity, data was rarefied to 18,433 reads. The statistical significance of the Shannon index and Chao1 index was calculated using analysis of variance (ANOVA). Beta diversity was determined using the robust Aitchison distance metric in QIIME2 using the DEICODE package.

This newer distance metric can better discriminate differences than other distance metrics such as UniFrac or Bray-Curtis.^76^ Differences in beta diversity were determined using permutational multivariate analysis of variance through the ‘adonis’ package in R (version 4.1.2).^74^ Differential abundance testing at various taxonomic levels was performed using DESeq2, which utilizes negative binomial modeling to account for the sparse data in microbiome sequencing.^77^ P-values were converted to q-values to adjust for multiple hypothesis testing.

### Magnetic resonance imaging MRI acquisition

A group of rats was randomly selected for neuroimaging studies (CDU, *n* = 6; CDE, *n* = 7; WDU, *n* =8; WDE, *n* = 7). The rats were deeply anesthetized with an intraperitoneal overdose injection of Euthasol (150 mg/kg; Virbac, Fort Worth, TX). Transcardiac perfusion was done with a prewash with 9.25% sucrose in distilled water and 4% paraformaldehyde (PFA; fixative, Cat No. 15714-S, Electron Microscopy Sciences, Hatfield, PA, USA) using the Perfusion Two^TM^ Automated Pressure Perfusion System (Leica Biosystems, Buffalo Grove, IL). The brains were preserved in the cranial vault after fixation and postfixed in 4% PFA, washed, and stored at 4°C in 0.1 M PBS with 0.05% Sodium azide until imaging. The experimenters were blinded to the group designation during MRI acquisition and analyses.

Encapsulated rat brains were prepared for scanning inside the skull to preserve the intact shape, as previously reported.^78^ Tissue was immersed in FC-40 (Fluorinert™) inside a 50 ml polypropylene conical tube and allowed at least 12 h to equilibrate to room temperature before scanning. A custom plastic fixture was inserted into the tube to physically stabilize the sample and minimize movement during gradient pulsations. Brain scans were collected in ultrahigh-resolution 11.1 Tesla/40 cm bore MRI scanner (Magnex Scientific Ltd., Oxford, UK) with a Resonance Research Inc. gradient set (RRI BFG- 240/120-S6, maximum gradient strength of 1000 mT/m at 325 Amps and a 200 µs risetime; RRI, Billerica, MA) and controlled by a Bruker Paravision 6.01 console (Bruker BioSpin, Billerica, MA) as previously reported.^79^ An in-house 3.2 x 7.6 cm quadrature birdcage tuned to 470.7MHz (1H resonance) was used for B1 excitation and signal detection (Advanced Magnetic Resonance Imaging and Spectroscopy Facility, Gainesville, FL).

Diffusion-weighted scans were acquired using a 4-shot, 2-shell spin echo planar diffusion imaging (DTI EPI) sequence in Bruker Paravision, with TR = 4 seconds, TE = 19 ms, number of averages = 4, gradient duration δ = 3 ms, diffusion time Δ = 8 ms, 77 images with 3 different diffusion weightings, eight b=0, 23 directions with b=600 s/mm2, and 46 directions with b=1200 s/mm^2^. A navigator signal was used by the Bruker reconstruction software to improve signal stability in the 4-shot EPI. Image saturation bands were placed on either side and below the brain to suppress non-brain signals during image acquisition. Images were collected with 1602 voxels x 50 slices at 0.15 mm2 x 0.5 mm slice thickness.

### Diffusion MRI processing

Diffusion MRI scans were processed using tools available in the FMRIB software library – FSL.^80^ Eddy correction was used for adjusting slight movements during image acquisition, and gradient files were rotated according to the motion correction vectors. After eddy correction, tensor element reconstruction and estimates of 1st, 2^nd^, and 3rd eigenvectors and eigenvalues (λ1, λ2, λ3, where λ1 is axial diffusivity and the average of λ2 and λ3 provides radial diffusivity values) was performed using weighted least squares regression on DTIFIT in FSL.^81^ Thus, generating independent images of mean, axial, and radial diffusivities (MD, AD, and RD, respectively) and Fractional Anisotropy (FA).

### NODDI processing

NODDI processing incorporates a multi-compartment model of the diffusion signal. The model includes an isotropic fraction of free unrestricted and hindered water (e.g., cerebral spinal fluid) and an anisotropic model that contains an intracellular fraction with zero radius infinite cylinders (high diffusion coefficients along the length of the principal axis only), and an extracellular fraction with nonzero diffusion perpendicular to the principal axis. The latter adds morphological information to the tissue model of the extracellular environment surrounding axons and dendrites (occupied by neurons and non-neuronal cell bodies).^82^ Analysis of dMRI scans following processing with the NODDI model was implemented within the Accelerated Microstructure Imaging via Convex Optimization (AMICO) framework,^83^ which accelerates the model fitting procedure in standard Linux PC running Python Software. The fitting of the intracellular volume fraction (i.e., zero-radius cylinders or “sticks”) involves the use of a Bingham-Watson series used in statistical shape analysis^84, 85^ and the results of the fitting process provide an index of the degree of orientation dispersion of so-called neurites. For the parameter fitting procedure, we assumed a value of ex vivo intrinsic diffusivity of 0.6 µm2/ms and isotropic diffusivity of 2.0 µm^2^/ms.^86^ The whole brain model-fitting generates maps of an intracellular volume fraction (ICVF: relative concentration of zero-radius cylinders modeling “neurites”), orientation dispersion (ODI: 0 for no dispersion as in highly organized parallel fiber bundles to a maximum of 1 for the highest degree of dispersion, as in cerebral cortical grey matter), and the isotropic free water fraction (ISO: 0 low isotropy to maximum isotropy of 1).^87^

### Image normalization and rat brain parcellation

DTI and NODDI scalars and normalized Jacobian matrix values reflecting structural brain differences between the groups were analyzed in primary regions of interest (ROI). B0 scan masks outlining rat brain boundaries were generated in MATLAB using Three-Dimensional Pulsed Coupled Neural Networks (PCNN3D)^88^ and adjusted manually using ITKSNAP.^89^ B0 aligned to a parcellated rat brain template.^90^ Images were cropped and linearly registered to the rat template using FSL linear registration tool (FLIRT),^91^ using a correlation ratio search cost, full 180-degree search terms, 12 degrees of freedom, and trilinear interpolation. The linear registration output was then warped nonlinearly to the template space using Advanced Normalization Tools (ANTs).^92^ The resulting deformation field images were used to generate Jacobian determinant maps to evaluate subject-to-atlas registration quality and to quantify the effects of regional structural differences. B0-to-atlas linear and nonlinear transformation matrices were applied to scalar maps later. All data were extracted and summarized in Excel (Microsoft, Redmond, WA, USA) and analyzed using GraphPad Prism version 9.4.1.681 for Windows (GraphPad Software, San Diego, California USA, www.graphpad.com). Given our established interest in investigating hippocampal vulnerabilities to obesogenic environments, our analyses focus on this region.

### Neurohistology Tissue Preparation

Following the MRI, a subgroup of brains was randomly selected from each group for histological analyses (*n* = 16; 4 rat brains per group). The brains were removed from the cranial vault after MRI and washed with phosphate-buffered saline (PBS). Brains were cryoprotected with sucrose (30%) and exposed to decreasing sucrose concentration for three consecutive days. Subsequently, the brains were rinsed and stored at 4°C in PBS before embedding in the Tissue-Tek^®^ O.C.T.^TM^ compound (Sakura, Torrance, CA, United States).

### Immunohistochemistry

Four (4) brains per group were sectioned with a cryostat (CM3050-S, Leica Biosystems, Vista, CA, USA) at 25 µm thickness in the coronal plane. Regions of interest (ROI) included dorsal and ventral areas of the CA1 region of the hippocampus of both hemispheres (−3.25 and −5.0 from Bregma). The sectioning pattern was 1:4, with three (3) consecutive brain sections used for immunohistochemistry and fourth sections used for immunofluorescence. Sections were mounted in glass slides, and immunohistochemistry staining was performed with the microglia/macrophage marker, ionized calcium-binding adaptor protein molecule 1 (Iba-1). First, sections were incubated in a quenching buffer (10% methanol, 1% hydrogen peroxide in PBS) to block endogenous peroxide activity. Blocking buffer containing secondary antibody’s host (5% Goat Serum) with 1% Triton-X in PBS was used to prevent nonspecific binding of antibodies to the tissue. Sections were then incubated overnight with a polyclonal rabbit anti-Iba1 primary antibody (1:750, rabbit anti-Iba1, Cat. No. 019-19741, Wako, USA) followed by secondary biotinylated antibody incubation (1:200, goat anti-rabbit IgG, Cat. No. BA-1000; Vector Labs Inc., Burlingame, CA, USA). Controls for nonspecific binding consisted in omitting the primary antibody (S Fig 5). Vectastain Elite ABC HRP kit and DAB peroxidase substrate kit with nickel (Cat. No. PK-6100 and SK-4100; Vector Labs Inc., Burlingame, CA, USA) were used to visualize staining according to manufacturer instructions. Following staining, slides were dehydrated in increasing concentrations of ethanol (70, 95, and 100%) and rinsed in Xylene to prepare for mounting with Permount Mounting Medium (Cat. No. 50-277-98, Fisher Chemical, Waltham, MA, USA). After applying glass coverslips, slides were allowed to dry overnight and stored at 4°C for subsequent visualization.

### Microglial Morphological Characterization Image Acquisition

Blinded investigators performed image acquisition and analyses as previously reported.^58^ Iba-1/DAB-stained tissue sections were imaged using a Keyence BZ-9000 series microscope (Keyence Corporation, Osaka, Japan) in brightfield at a 40× magnification. To obtain a single high-quality full-focused composite image that showed detailed magnification of ramified processes of the cells and facilitated image segmentation, a multi-plane virtual-Z mode with 10 images (1 mm thickness) in 20 mm depth of tissue section was captured using the BZ9000 BIOREVO sectioning algorithm. Hippocampal dorsal and ventral CA1 areas were scanned to obtain at least 9 sets of images per region of interest per hemisphere covering a field of view area of 3.6 mm. Each acquired image included at least six (6) Iba-1+ cells and was saved as a Tag Image File Format (TIFF) file.

### Image Processing and segmentation

Images were processed unbiasedly and systematically to obtain a filled image using the public domain software *ImageJ.*^93^ For this purpose, a single high-quality image was obtained by merging the 10 (8-bit) images from each z-stack. The composite images were processed under a minimum threshold to soften the background and enhance the contrast. Each object (cell) was manually edited to obtain a cell mask consisting of continuous pixels. A set threshold of pixel number addition (no more than 4 pixels) to join processes belonging to a selected cell was established and performed systematically. When applicable, a 2-pixel set threshold was applied to separate ramifications to neighboring cells. This step was carefully performed under the view of the original z-stack images to avoid bias. Subsequently, the image is binarized to obtain a black-and-white image by applying a standardized software feature. The final filled shape image containing the masks was saved as a TIFF and later analyzed to quantify the morphological changes of microglial cells.

### Fractal Analysis

FracLac^94^ for *ImageJ* was used to quantify the morphological changes of microglia. Foreground pixel quantification of the filled binary images per object (cell) was done with the Box Counting (BC) feature, which counts the number of boxes containing foreground pixels and the successively smaller caliber grids. BC box size scale was obtained as a Power Series where the base is raised to the exponent added to it to make successive sizes.

Finally, the slope for each object was the average of 12 measurements with different and random grid placements. Box Counting, Convex, and Hull Area measurements were exported using Excel (Microsoft, Redmond, WA, USA) for record and tabulation. GraphPad Prism version 9.4.1.681 (GraphPad Software, San Diego, California USA) was used for statistical analyses.

### In Vitro assessments Cell Culture

The Human Microglial Clone 3 (HMC3) cell line used in this study was purchased from the American Type Culture Collection (ATCC^®^, CRL-3304 ^TM^, Manassas, VA, USA). The cells were cultured in minimum essential medium (MEM) (Cat. No.11095080; Gibco, ThermoFisher Scientific, Waltham, MA, USA), supplemented with 10% FBS (F4135, Sigma Aldrich, St. Louis, MO, USA) and 100 U/ml penicillin/streptomycin (Cat. No. 15140122; Gibco, ThermoFisher Scientific, Waltham, MA, USA) at 37°C in a 5% CO_2_ humidified atmosphere. We used HMC3 at passages 2 to 10. The passage number of HMC3 used in this study was not allowed to exceed passage number 10. Briefly, twelve hours before treatment, cells were serum starved in MEM with 0.2% FBS. At the time of treatment, cells were washed with PBS and incubated with vehicle, cortisol, and/or palmitic acid in MEM with 0.2% FBS. The cells were harvested for analysis after a 24 h incubation period.

### Cell Viability Assay

Cell viability was assayed by 3-(4,5-dimethylthiazol-2-yl)-2,5-diphenyltetrazolium bromide (MTT, Cat. No. M6494, ThermoFisher Scientific, Waltham, MA, USA) to differentiate the range of toxic and nontoxic palmitic acid concentrations. HMC3 cells were seeded in 96-well plates at a density of 1 × 10^4^ cells per well for 24 h. Cells were incubated with palmitic acid at concentrations ranging from 50 to 800 μM or with vehicle (ethanol) for 24 h. MTT reagent (0.4 mg/mL in PBS) was added to each well, and cells were incubated at 37 °C, 5% CO_2_ for 4 h. The culture supernatants were discarded, and 200 μL of DMSO was added to each well. The absorbance at 570 nm was measured using a microplate reader. MTT assay was performed 4 times, and each assay was done in duplicate. The half maximal inhibitory concentration (IC_50_) value of palmitic acid to the cells was obtained.

### RNA Extraction and Quantitative Real-Time PCR

Total RNA was prepared with 1 mL of TRIzol™ Reagent (Cat. No.15596026, ThermoFisher Scientific, Waltham, MA, USA) according to the manufacturer’s instructions. The High-Capacity cDNA Reverse Transcription Kit (Cat. No. 4368814, Applied Biosystems, Waltham, MA, USA) was used to make 20 μL of cDNA from 2 μg of RNA. Gene expression was analyzed using real-time PCR using TaqMan gene expression assay for FKBP5, IL6, TNFα, NFKB, and GAPDH in the StepOnePlus™ Real-Time PCR System (Applied Biosystems, Waltham, MA, USA). The ΔΔ cycle threshold method was used to determine mRNA levels. Gene expression was normalized to GAPDH levels.

### Gene Silencing

HMC3 cells were plated at 2.5 × 105 cells/well cell density in a six-well plate using media without FBS or antibiotics. Transfection of FKBP5 siRNA (Cat. No. 4390824, Applied Biosystems, Waltham, MA, USA) was performed using Lipofectamine™ 3000 Transfection Reagent (Cat. No. L3000015, ThermoFisher Scientific, Waltham, MA, USA) according to the manufacturer’s instructions. Twenty-four hours after siRNA transfection, cells were treated with hydrocortisone or palmitic acid for 24 h. Cells were harvested, centrifuged at 1500 revolutions per minute (rpm) for 5 min at RT, and subjected to RNA extraction.

### Cytokine Quantification

To determine the concentration of IL-6 or TNFα in cell culture media of control or treated HMC3 cells, ELISA kits (Cat. No. KHC0061, KAC1751, respectively, ThermoFisher Scientific, Waltham, MA, USA) were used following the manufacturer’s instructions.

### Confocal Microscopy

HMC3 cells were cultured in two-well Nunc Lab-Tek II chamber slides. After 24 h of treatment, the cells were fixed in a 4% paraformaldehyde solution for 15 min and permeabilized with 0.5% Triton™ X-100 in DPBS for 10 min at room temperature. The cells were blocked with 1% bovine serum albumin for 30 min at room temperature, then probed with CD68 Monoclonal Antibody, eFluor 570, eBioscience (Cat. No. 41-0687-82, ThermoFisher Scientific, Waltham, MA, USA) for 2 h at room temperature. Cells were fixed again in 4% PFA before applying ProLong® Diamond Antifade Mountant with DAPI (Cat. No. P36935, Life Technologies, Carlsbad, CA, USA).

### Determination of ROS Generation

ROS generation of HMC3 cells was measured by the Muse Oxidative stress kit using the Muse® Cell Analyzer (MilliporeSigma, Burlington, MA) fluorescent-based analysis. The protocol was performed according to the manufacturer’s recommendations. Briefly, HMC3 cells were treated with hydrocortisone and or palmitic acid and incubated for 24 h. 1 × 10^7^ cells/mL samples were prepared in 1X assay buffer and treated with Oxidative stress reagent, based on dihydroethidium (DHE) used to detect ROS that is oxidized with superoxide anion to procedure the DNA-binding fluorophore ethidium bromide which intercalates with DNA resulting in red fluorescence. This method was validated using N-Acetyl-L-Cysteine (NAC) as ROS inhibitor and Antimycin A as ROS inducer. To evaluate whether cellular fluorescence reflects ROS generation, three different experiments with duplicate samples were incubated with 5 mM NAC or vehicle for 30 min, and then cells were stimulated with 200 µM Antimycin A or 100 nM Hydrocortisone or 50 µM Palmitic Acid for 30 min to ensure that the cellular fluorescence reflected ROS generation. We ran stimulated, unstimulated, with and without dihydroethidium (DHE) samples.

### Statistical Analysis

We analyzed the data using GraphPad Prism version 8.0. Shapiro-Wilk statistical analyses were used to determine sample distribution. The Brown-Forsythe test was used to test for the equality of group variances. When appropriate, two-way analysis of variance (ANOVA) was used to examine the effect of the diet type, stress, and interaction between factors on outcome measures. Multiple comparisons were made using Dunnett’s (following Welch’s ANOVA) or Sidak’s (repeated measures two-way ANOVA) tests. The ROUT method was used to investigate outliers. We considered differences significant if *p* < .05. The data is shown as the mean ± standard error of the mean (S.E.M.). We also conducted a posthoc power analysis with the *G*Power* program.^95^ For two-way ANOVA analyses, the statistical power (1-β) for data including the four study groups was 0.79 for detecting a medium-size effect (*d* = 0.41). The power exceeded 0.99 for the detection of large effect size (*d* = 0.8).

## RESULTS

This study investigated brain, behavior, and immunometabolic responses to stress in rats exposed to an obesogenic diet during adolescence. We postulated that intake of an obesogenic Western-like high saturated fat diet (WD) during adolescence would alter responses to chronic psychosocial stress (PSS). *F* statistics and post hoc test results are detailed in **Tables 1-9** and **Supplemental Tables 2-7**.

**Table 1.** Detailed summary of behavioral assessment findings and statistics. Bold denotes significant effects following three-way ANOVA and post hoc comparisons between CD UNEXP, CD EXP, WD UNEXP, and WD EXP. Sample size = 14 rats/group (before outlier testing). Abbreviations: CD control diet, WD western diet, EPM elevated plus maze, SYM Social Y Maze, FPS fear-potentiated startle, ASR Acoustic Startle Reflex, SYM Social Y Maze, UNEXP Unexposed, WD Western Diet.

|  | **ASR**<br>(% change from baseline) | **FPS**<br>(% FPS) | **EPM**<br>(% OA duration) | **SYM**<br>(# social interactions) | **SYM**<br>Frequency (Conspecific: Object) |
| --- | --- | --- | --- | --- | --- |
| Source of Variation | F Statistic, posthoc | F Statistic, posthoc | F Statistic, posthoc | F Statistic, posthoc | F Statistic, posthoc |
| **WD x PSS** | F (1, 51) = 0.204, $p=0.6533$ | F (1, 34) = 0.034, $p=0.853$ | F (1, 53) = 2.649, $p=0.109$ | F (1, 50) = 0.442, $p=0.508$ | F (1, 47) = 0.065, $p=0.7993$ |
| **PSS** | **F (1, 51) = 4.708, $p=0.0347$** | **F (1, 34) = 7.511, $p=0.009$** | F (1, 53) = 2.832, $p=0.098$ | **F (1, 50) = 5.293, $p=0.025$** | F (1, 47) = 1.664, $p=0.2034$ |
| **WD** | **F (1, 51) = 5.305, $p=0.0254$** | F (1, 34) = 0.008, $p=0.927$ | **F (1, 53) = 11.37, $p=0.001$** | **F (1, 50) = 5.168, $p=0.027$** | **F (1, 47) = 4.180, $p=0.0465$** |
| **CDU vs WDU** | $p=0.2306$ | | $p=0.6022$ | $p=0.1882$ | |
| **CDU vs CDE** | $p=0.6411$ | | $p>0.9999$ | $p=0.1816$ | |
| **CDU vs WDE** | $p=0.9997$ | | **$p=0.0045$** | **$p=0.0134$** | |
| **WDU vs CDE** | **$p=0.0105$** | | $p=0.6270$ | $p>0.9999$ | |
| **WDU vs WDE** | $p=0.2372$ | | $p=0.0974$ | $p=0.6421$ | |
| **CDE vs WDE** | $p=0.5513$ | | **$p=0.0050$** | $p=0.6543$ | |

**Table 2.** Statistical Information on microglial morphological measurements in Figure 3,4. Bold denotes significant effects following three-way ANOVA and post hoc comparisons between CD UNEXP, CD EXP, WD UNEXP, and WD EXP. Sample size: Neurohistology: 4 rats/group, Gene Expression: 3 rats/group.

| | *Iba-1+* cells | Lacunarity | Span Ratio | TNF-α<br>HPC GE |
| --- | --- | --- | --- | --- |
| Source of Variation | F Statistic,<br>posthoc | F Statistic,<br>posthoc | F Statistic,<br>posthoc | F Statistic,<br>posthoc |
| **WD x PSS** | -- | F (1, 12) = 0.3175,<br>*p* = 0.5835 | F (1, 12) = 0.001,<br>*p* = 0.9666 | F (1, 8) = 1.691,<br>*p* = 0.2297 |
| **PSS** | F (1, 13) = 1.311,<br>*p* = 0.2729 | F (1, 12) = 0.6753,<br>*p* = 0.4272 | **F (1, 12) = 10.87,**<br>***p* = 0.0064** | **F (1, 8) = 23.00,**<br>***p* = 0.0014** |
| **WD** | **F (1, 13) = 11.85,**<br>***p* = 0.0044** | **F (1, 12) = 11.270,**<br>***p* = 0.0057** | F (1, 12) = 1.565,<br>*p* = 0.2348 | **F (1, 8) = 8.237,**<br>***p* = 0.0208** |
| **CDU vs WDU** | ***p* = 0.0200** |  | *p* = 0.7975 | *p* = 0.0715 |
| **CDU vs CDE** | *p* = 0.6697 |  | *p* = 0.1383 | ***p* = 0.0110** |
| **CDU vs WDE** | ***p* = 0.0287** |  | ***p* = 0.0325** | ***p* = 0.0028** |
| **WDU vs CDE** | *p* = 0.3996 |  | *p* = 0.4964 | *p* = 0.5539 |
| **WDU vs WDE** | *p* = 0.6697 |  | *p* = 0.1523 | *p* = 0.1399 |
| **CDE vs WDE** | ***p* = 0.0200** |  | *p* = 0.8277 | *p* = 0.6940 |

**Table 3.** Statistical Information on microglial distribution by population phenotype in Figure 4. Chi-squared analysis (1982) was performed, and distribution is denoted in number and percentage over three distinct phenotypical distributions. Sample size: Neurohistology: 16,487 *Iba-1+* cells, 4 rats/group.

|  | **Hyper-ramified** | **Intermediate** | **Hypo-ramified** |
| --- | --- | --- | --- |
| Experimental groups | # *Iba-1*<sup>+</sup> \| (% population) | # *Iba-1*<sup>+</sup>, (% population) | # *Iba-1*<sup>+</sup>, (% population) |
| **WD EXP** | 1,417 \| (22) | 4392 \| (70) | 505 \| (8) |
| **WD UNEXP** | 844 \| (17) | 3587 \| (75) | 383 \| (8) |
| **CD EXP** | 356 \| (12) | 1,501 \| (51) | 1,064 \| (37) |
| **CD UNEXP** | 349 \| (14) | 1,263 \| (52) | 826 \| (34) |

**Table 4.** Statistical information from microbiome analysis in Figure 4. Bold denotes significant effects following three-way ANOVA and post hoc comparisons between CD UNEXP, CD EXP, WD UNEXP, and WD EXP.Sample size: 10 rats/group.

|  | **Ls NK4A136**<br>Relative abundance | **PCA** | **Richness**<br>Chao1 | **Diversity**<br>Shannon | ***Prevotellacea***<br>Relative abundance |
| --- | --- | --- | --- | --- | --- |
| Source of Variation | F Statistic, posthoc | t test | F Statistic, posthoc | F Statistic, posthoc | F Statistic, posthoc |
| **WD x PSS** | **F (1, 34) = 8.91, $p = 0.005$** | -- | F(1, 34) = 0.003, $p = 0.959$ | F (1, 34) = 0.15, $p = 0.069$ | F(1, 34) = 0.2, $p = 0.591$ |
| **PSS** | **F (1, 34) = 5.86, $p = 0.021$** | **t (34), $p < 0.001$** | F (1, 34) = 0.03, $p = 0.856$ | F (1, 34) = 1.7, $p = 0.190$ | F(1, 34) = 0.02, $p = 0.879$ |
| **WD** | **F (1, 34) = 9.98, $p = 0.003$** | t (34), $p = 0.780$ | F (1, 34) = 2.51, $p = 0.122$ | F (1, 34) = 0.007, $p = 0.934$ | **F(1, 34) = 9.6, $p = 0.003$** |
| **CDU vs WDU** | **$p = 0.0005$** |  |  |  |  |
| **CDU vs CDE** | **$p = 0.0048$** |  |  |  |  |
| **CDU vs WDE** | **$p = 0.0019$** |  |  |  |  |
| **WDU vs CDE** | $p = 0.954$ | | | | |
| **WDU vs WDE** | $p = 0.9743$ | | | | |
| **CDE vs WDE** | $p = 0.9994$ | | | | |

**Table 5.** Statistical information from endocrine reactivity analysis. Fecal corticosterone metabolites were measured, Bold denotes significant effects following three-way ANOVA and post hoc comparisons. Sample size: t test: CD, *n* = 12, WD *n* = 13; FCM 4 weeks (CDU, *n* = 4; CDE, *n* = 4; WDU, *n* = 5; WDE, *n* = 5). FCM 4 weeks +1 day (CDU, *n* = 4; CDE, *n* = 4; WDU, *n* = 5; WDE, *n* = 5). FCM 5 weeks +1 day (CDU, *n.*= 6; CDE, *n* =6; WDU, *n* = 7; WDE, *n* = 6).

|  | FCM | FCM | FCM |
| --- | --- | --- | --- |
|  | 4 weeks in WD | 4 weeks in WD +<br>1 day after first PSS | 5 weeks in WD +<br>1 day after second PSS |
| Source of Variation | t test, p | F Statistic, post-hoc | F Statistic, post-hoc |
| **WD vs CD** | **t (22) = 3.559, $p = 0.0018$** | -- | -- |
| **WD x PSS** | | **F (1, 14) = 5.185, $p = 0.0390$** | F (1, 21) = 1.398, $p = 0.2502$ |
| **PSS** |  | **F (1, 14) = 17.96, $p = 0.0008$** | **F (1, 21) = 5.829, $p = 0.0250$** |
| **WD** | | **F (1, 14) = 10.90, $p = 0.0052$** | F (1, 21) = 0.9038, $p = 0.3526$ |
| **CDU vs WDU** |  | **p = 0.0071** | p = 0.4343 |
| **CDU vs CDE** |  | p = 0.5688 | p = 0.8274 |
| **CDU vs WDE** |  | p = 0.9096 | p = 0.7418 |

**Table 6.** Statistical information from hippocampal FKBP5 gene expression and methylation main effects. Hippocampal gene expression and the percentage of Fkbp5 methylation in all experimental groups. Bold denotes significant effects following three-way ANOVA and post hoc comparisons. Sample size: 3 rats/group.

|  | **Fkbp5** | **Fkbp5** |
| --- | --- | --- |
|  | **HPC GE** | **HPC % Methylation** |
| Source of Variation | F Statistic, post-hoc | F Statistic, post-hoc |
| WD | **F (1, 8) = 21.6, p = 0.0016** | **F (1, 32) = 10.82, p = 0.0024** |
| PSS | **F (1, 8) = 8.46, p = 0.0196** | F (1, 32) = 3.79, p = 0.0603 |
| WD x PSS | F (1, 8) = 0.02, p = 0.8879 | F (1, 32) = 0.58, p = 0.4498 |
| CpG |  | **F (3, 32) = 53.7, p < 0.0001** |
| CpG x WD |  | F (3, 32) = 0.10, p = 0.9540 |
| CpG x PSS |  | F (3, 32) = 0.67, p = 0.5759 |
| CpG x WD x PSS |  | F (3, 32) = 0.47, p = 0.7027 |

**Table 7.** Statistical Information of TNF-α gene expression, protein release and ROS in HCM3 cells following PA+CORT treatment. TNF-α gene expression was significantly affected by the interaction of PA + CORT (*p* = 0.0002) and by CORT treatment alone (*p* = 0.0146). TNF-α protein release was significantly affected by the interaction of PA + CORT (*p* = 0.0021) and by CORT (*p* = 0.0002) and PA (*p* = 0.0096). ROS was significantly influenced by the interaction of PA + CORT (*p* = 0.0459) and by CORT (*p* < 0.0001) and PA (*p* < 0.0001). Bold denotes significant effects following three-way ANOVA and post hoc comparisons. Sample size: gene and protein measurements: 3 rats/group; ROS: 4 rats/group.

|  | **TNF-α** | **TNF-α** | **ROS** |
| --- | --- | --- | --- |
|  | Gene expression | Protein release | RFU |
| Source of Variation | F Statistic, post-hoc | F Statistic, post-hoc | F Statistic, post-hoc |
| CORT:PA | **F (1, 11) = 30.84, p = 0.0002** | **F (1, 12) = 15.19, p = 0.0021** | **F (1, 12) = 4.957, p = 0.0459** |
| CORT | **F (1, 11) = 8.384, p = 0.0146** | **F (1, 12) = 28.50, p = 0.0002** | **F (1, 12) = 37.02, p < 0.0001** |
| PA | F (1, 11) = 0.02779, p = 0.8706 | **F (1, 12) = 9.459, p = 0.0096** | **F (1, 12) = 96.35, p < 0.0001** |
| VEH:VEH vs. VEH:PA | **p = 0.0102** | p = 0.936 | **p = < 0.0001** |
| VEH:VEH vs. CORT:VEH | p = 0.3187 | p = 0.7415 | **p = 0.0004** |
| VEH:VEH vs. CORT:PA | p = 0.1687 | **p = 0.0003** | **p = < 0.0001** |
| VEH:PA vs. CORT:VEH | p = 0.2987 | p = 0.4143 | p = 0.0878 |
| VEH:PA vs. CORT:PA | **p = 0.0003** | **p = 0.0001** | p = 0.0755 |
| CORT:VEH vs. CORT:PA | **p = 0.0114** | **p = 0.0017** | **p = 0.0008** |

**Table 8.** Measurements of HCM3 ROS production and M2 polarization following PA+CORT treatment in Figure 7. Sample size: *n* = 4 rats/group

|  | **ROS**<br>RFU | **M1** | **M2** |
| --- | --- | --- | --- |
| Source of Variation | F Statistic, post-hoc | % ROS | % ROS |
| VEH | -- | 73.07 | 24.40 |
| CORT:PA | **F (1, 12) = 4.957, p = 0.0459** | 35.83 | 62.30 |
| CORT | **F (1, 12) = 37.02, p < 0.0001** | 54.93 | 43.00 |
| PA | **F (1, 12) = 96.35, p < 0.0001** | 40.63 | 56.97 |
| VEH:VEH vs. VEH:PA | **p = < 0.0001** |  |  |
| VEH:VEH vs. CORT:VEH | **p = 0.0004** |  |  |
| VEH:VEH vs. CORT:PA | **p = < 0.0001** |  |  |
| VEH:PA vs. CORT:VEH | p = 0.0878 |  |  |
| VEH:PA vs. CORT:PA | p = 0.0755 |  |  |
| CORT:VEH vs. CORT:PA | **p = 0.0008** |  |  |

**Table 9.**
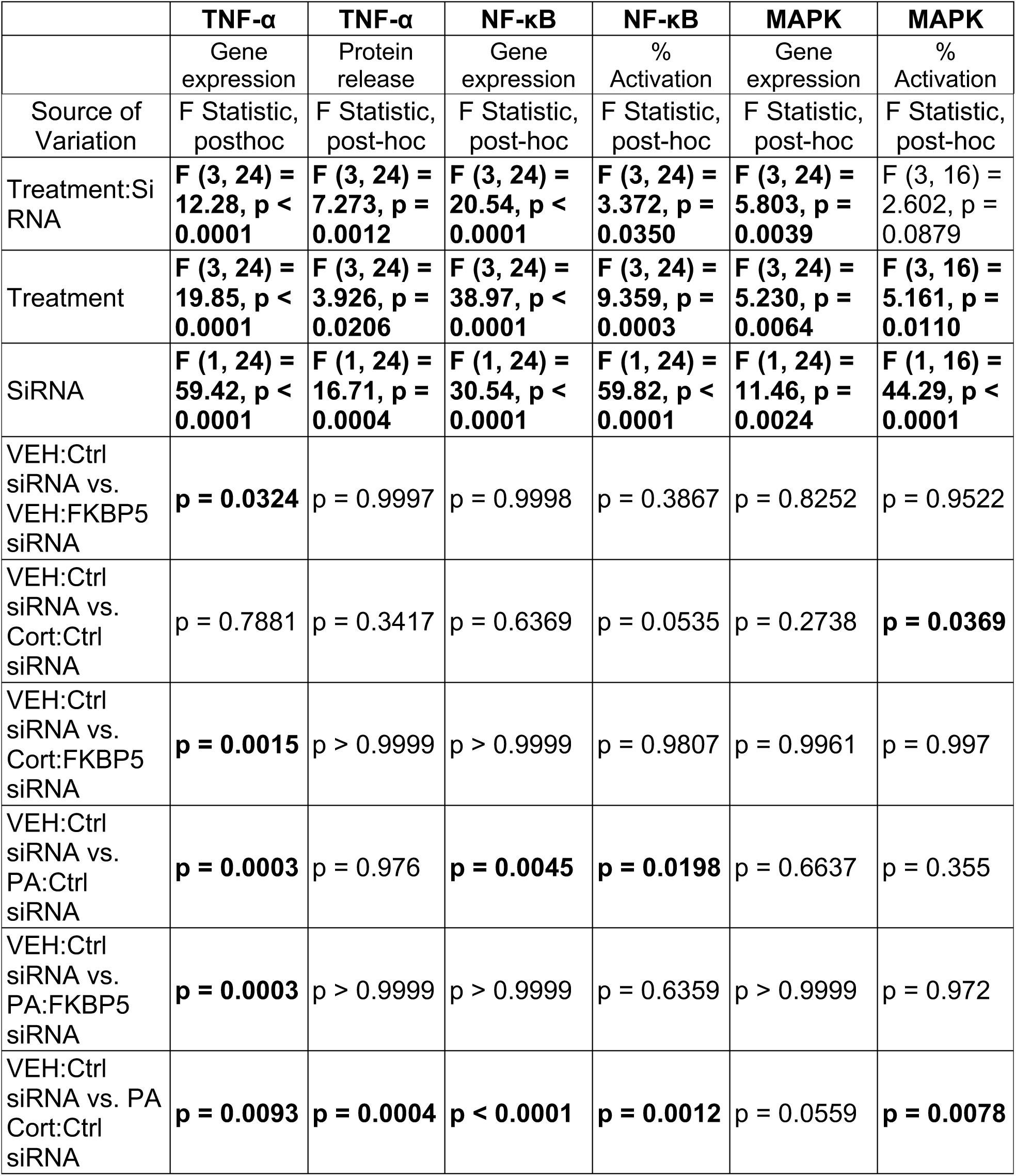
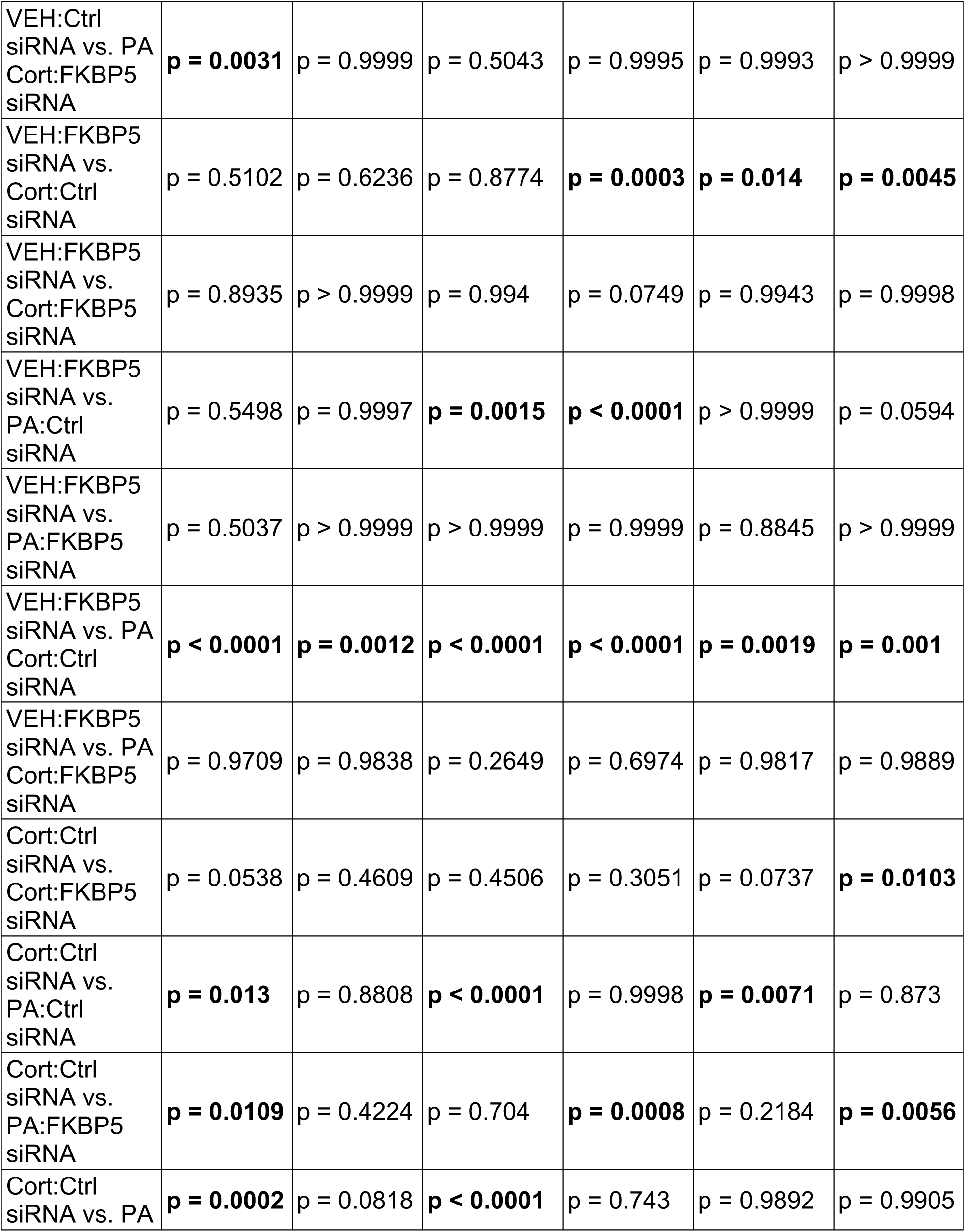
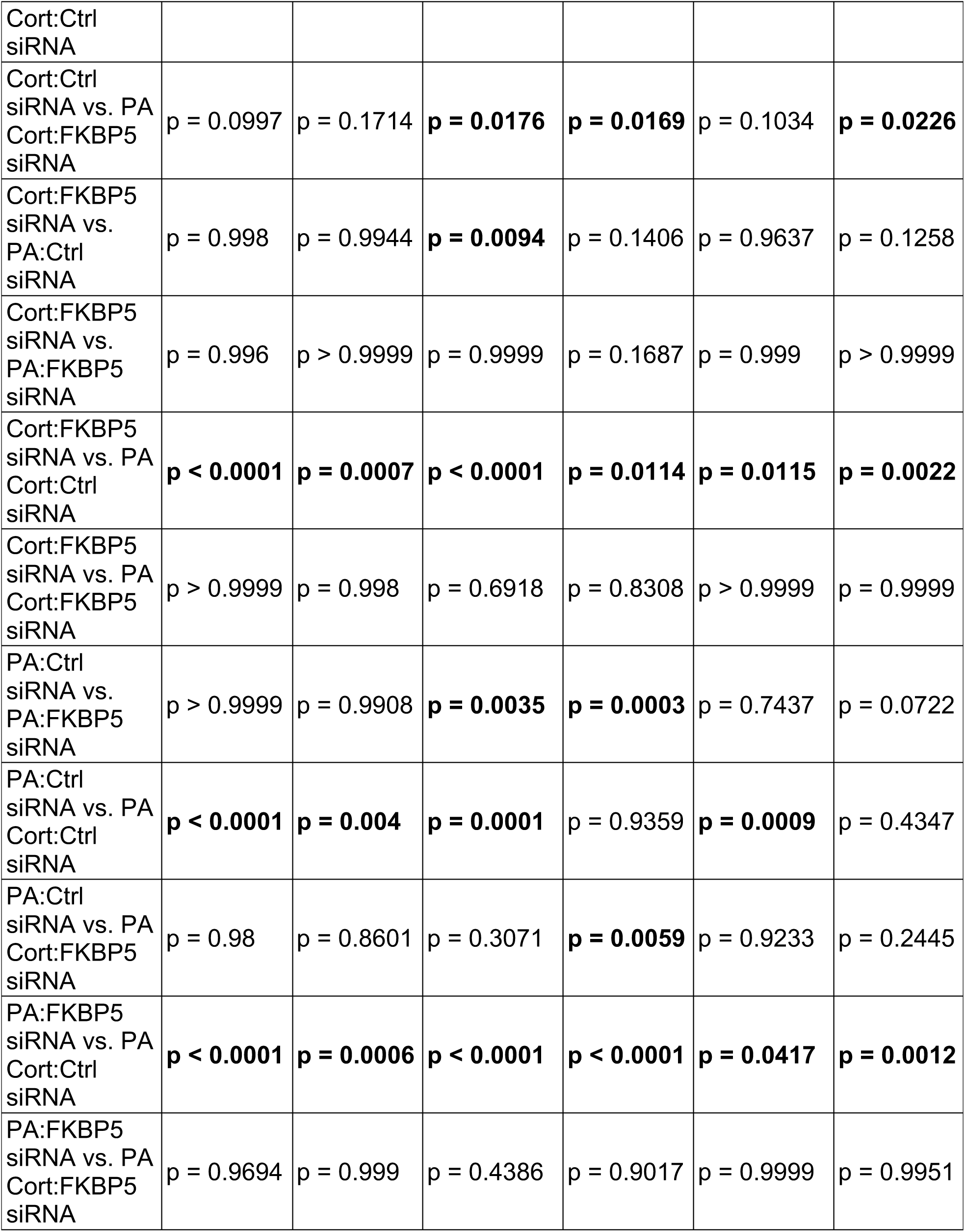
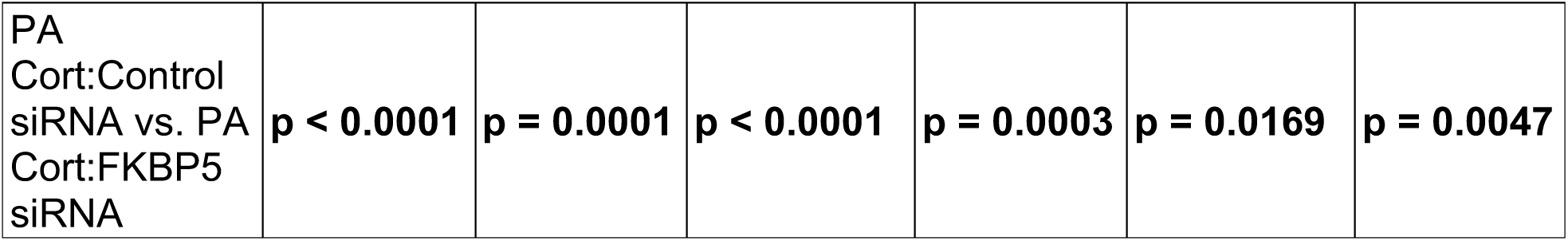
Statistical Information of Selected gene and protein expression in HCM3 following PA+CORT treatment after FKBP5 SiRNA in Figure 8. Sample size: *n* = 6 rats/group

### Exposure to obesogenic conditions during adolescence increases body weight and caloric intake

Body weight and food intake were measured bi-weekly to determine the effects of the obesogenic conditions on growth and caloric intake. We found that WD and PPS promoted significant weight gain. Statistical information for growth charts and caloric intake is detailed in **Supplemental Table 2**.

The percent change in body weight after the completion of the stress model (PND91) showed that both WD (*p* < 0.0001) and PSS (*p* = 0.0004) are significant factors contributing to weight increase (**S Fig 1A**). Analyses revealed a significant effect in body weight increase by diet alone (*p* = 0.0125), by stress alone in the WD-fed group (*p* = 0.0192), and among the exposed groups regardless of diet type (*p* = 0.0041). No significant increase in body weight was observed in the stressed rats that consumed the CD (*p* = 0.1028). Notably, WD and PSS synergized to increase weight gain significantly (*p <* 0.0001) (**S Fig 1A**). Two-way ANOVA demonstrated that PSS significantly contributed to increased caloric intake (*p* = 0.0033), suggesting a potential pathway for increased body weight gain (**S Fig 1B**).

### WD and PSS induce distinct effects on cardiometabolic function

Given the well-established effect of obesogenic diets and stress on cardiometabolic health, we measured fasting blood glucose levels (FBG), heart rate (HR), and blood pressure (BP) at endpoint. We found that PSS decreased FBG (*p* = 0.0010), particularly in the rats that consumed the CD (*p* = 0.0196) (**S Fig 2**). This hypoglycemic response suggests adrenal fatigue and metabolic dysregulation, which are commonly associated with severe stress.

**Figure 2.**
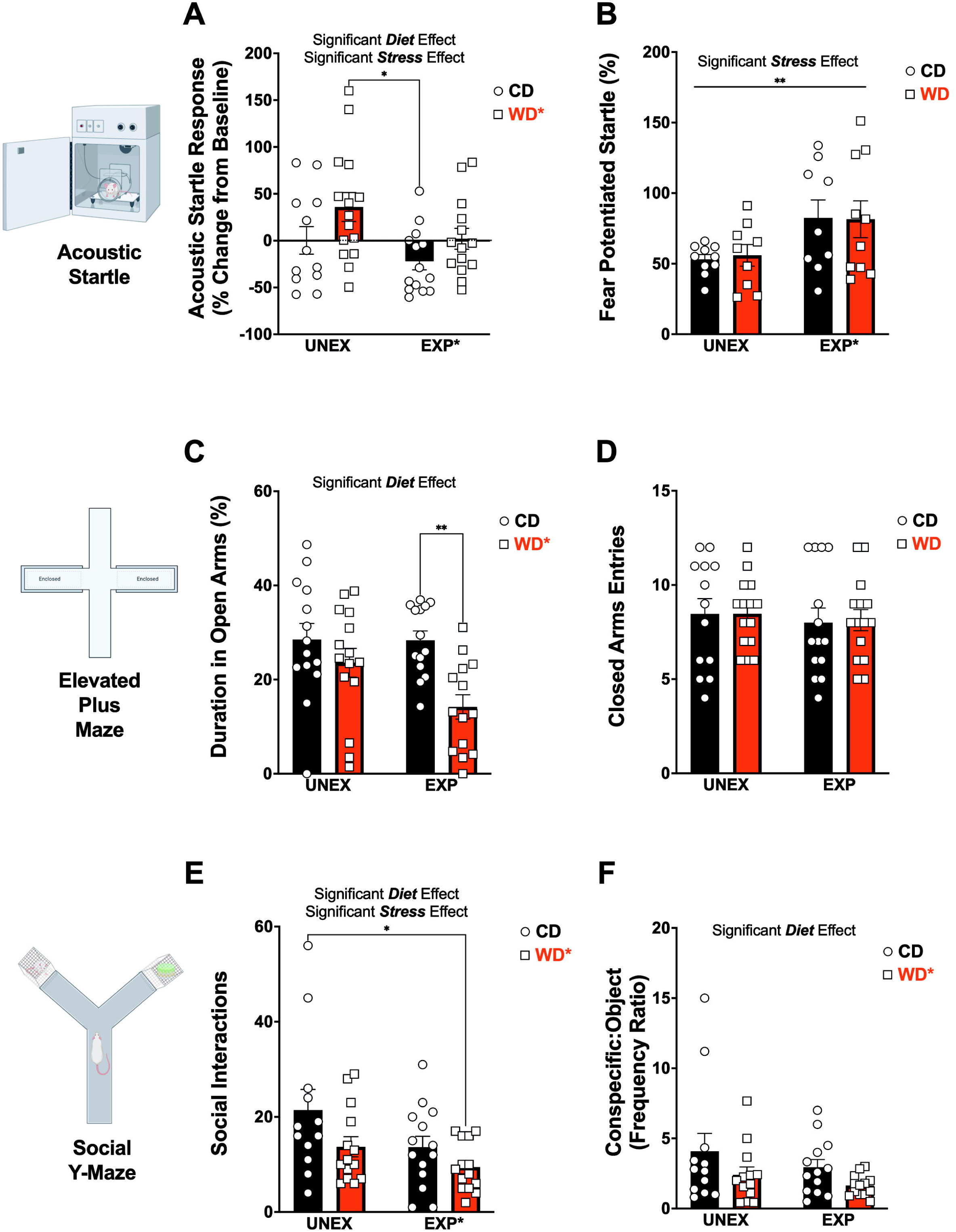
Behavioral phenotypes associated with early access to an obesogenic diet and psychosocial stress. **(A)** Acoustic Startle Response (ASR) chamber schematic and response in percent change from baseline. Stress (*p* = 0.0347) and diet (*p* = 0.0254) significantly contribute to the ASR response. CD-fed animals show a significant decrease in ASR (*p* = 0.0105) when exposed to the stress paradigm compared to the WD diet alone. **(B)** Fear potentiated startle (FPS) percentage was significantly increased with exposure to stress (*p* = 0.0097) **(C)** Elevated plus maze (EPM) schematic and duration in open arms. WD consumption significantly reduces time spent in Open Arms (*p* = 0.0014). WD and PSS significantly decrease time spent in the OA when compared with the CD EXP rats (*p* = 0.0050). **(D)** Number of closed arms entries in EPM was not affected by diet or stress (*p* > 0.0050). **(E)** Social Y Maze (SYM) number of social interactions was significantly decreased by and diet (*p* = 0.0273) and stress (*p* = 0.0256) with WD EXP animals having the most dramatic decrease (*p* = 0.0134). **(F)** Frequency of interactions between the conspecific or the object in SYM was significantly altered by diet. For all behavioral samples: sample size = 14 rats/group (before outlier testing).

Measures of cardiovascular status showed no significant interactions or main effect of WD or PSS in mean arterial blood pressure and systolic pressure. However, the rats that consumed the WD exhibited decreased diastolic blood pressure (*p* = 0.0195) and heart rate (*p* = 0.0096) (**Sup Fig 3A-D**).

**Figure 3.**
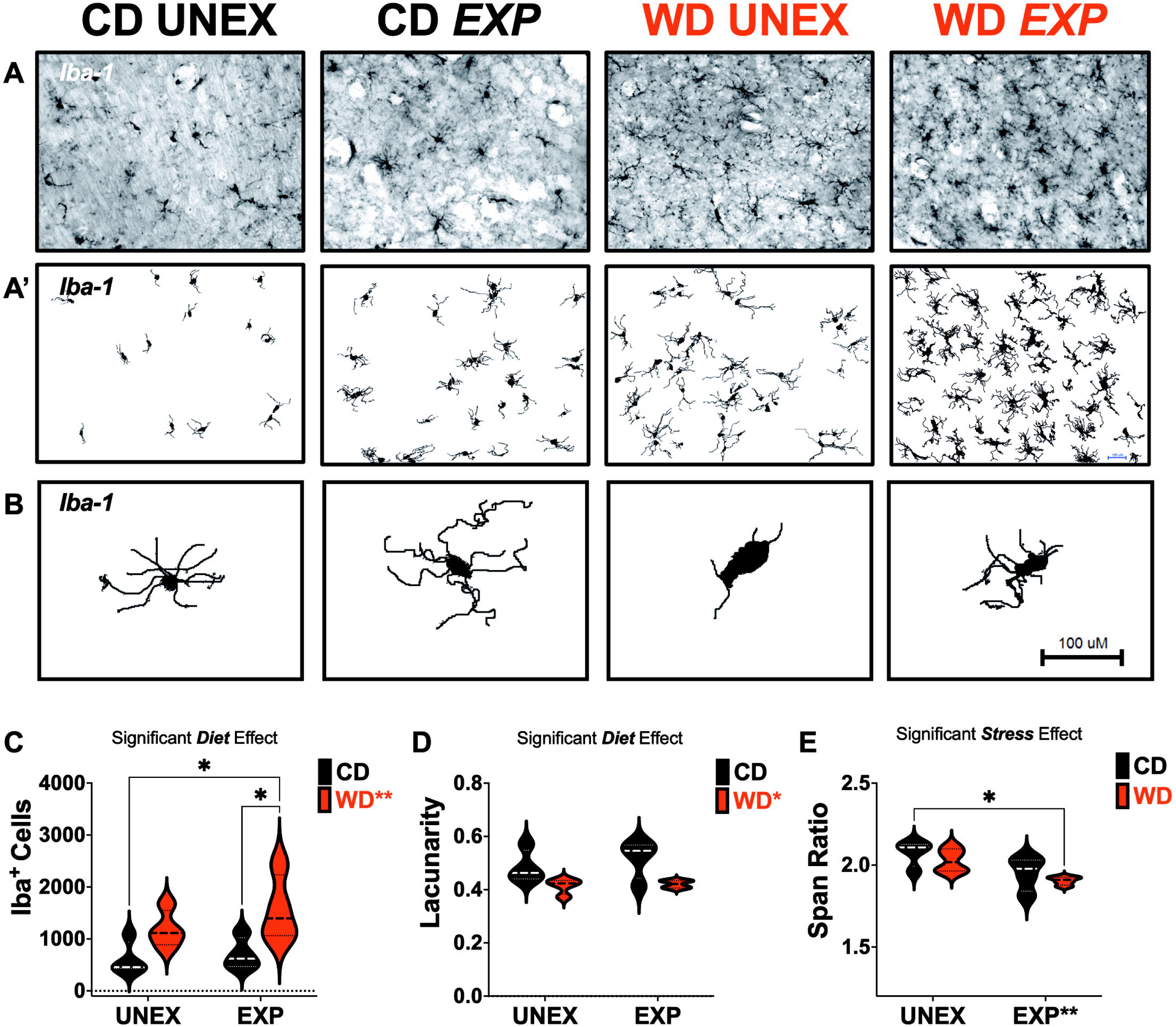
Obesogenic environment alters microglial number and induces a proinflammatory phenotype in hippocampal CA1. **(A)** Representative images of full-focused composite 10× brightfield images from hippocampal CA1 were acquired and stained for Iba-1. **(A’)** Corresponding binarized images were processed for quantitative morphometric analysis. **(B)** Representative microglia morphologies in each study group. **(C)** Diet significantly contributed to the total number of Iba1+ cells (*p* = 0.0044) in the microglial population numbers. Multiple comparison analyses showed that WD significantly increased the number of Iba1+ cells (*p* = 0.0200), and chronic stress synergized to increase the WD effect (*p* = 0.0287). **(D)** Lacunarity was significantly affected by diet (*p* = 0.0057) and decreased by WD. **(E)** Span ratio measurements were significantly affected by Stress (*p* = 0.0064) and decreased in WD EXP rats (*p* = 0.0325). Sample size = 4 rats/group. Scale bars: 100 microns.

### WD intake during adolescence increases the susceptibility to stress-induced anxiety-like behaviors

Previous work from our laboratory and others has demonstrated that exposure to an obesogenic diet during adolescence alters the neural and behavioral substrates implicated with emotional regulation. In this study, we asked whether those effects would be exacerbated when animals are exposed to an obesogenic diet in combination with a robust model of chronic psychosocial stress.

### The obesogenic conditions have differential effects on the acoustic startle reflex maturation

Acoustic startle response (ASR) is increased during states of anxiety and fear in rats and humans. Multiple studies lead to the concept that the magnitude of ASR can be regarded as an indicator of the emotional state of an organism.^96^ In this study, we used the ASR as a tool to longitudinally assess changes in emotional reactivity. We found that the PSS attenuated (*p* = 0.0347) and the WD increased (*p* = 0.0254) the ASR magnitude during adulthood relative to baseline values from early adolescence. These results validate prior findings and demonstrate competing influences of the WD and PSS on emotional regulation during adolescence.

### Exposure to PSS during adolescence increases the magnitude of the fear-potentiated startle

We used the trace fear-potentiated startle (FPS) paradigm to assess hippocampal-dependent conditioned fear learning and fear extinction. Our results showed that exposure to PSS significantly increased the FPS during the learning testing session (*p* = 0.0097) (**Fig 2B**). Interestingly, we found a significant interaction effect on FPS responses during the fear extinction testing session (*p* = 0.0216) (**S Fig 4C**). Unexposed rats that consumed the WD exhibited deficits in fear extinction relative to unexposed rats that consumed the CD. Conversely, PSS-exposed rats that consumed the WD exhibited higher fear extinction learning relative to the rats that consumed the CD.

**Figure 4.**
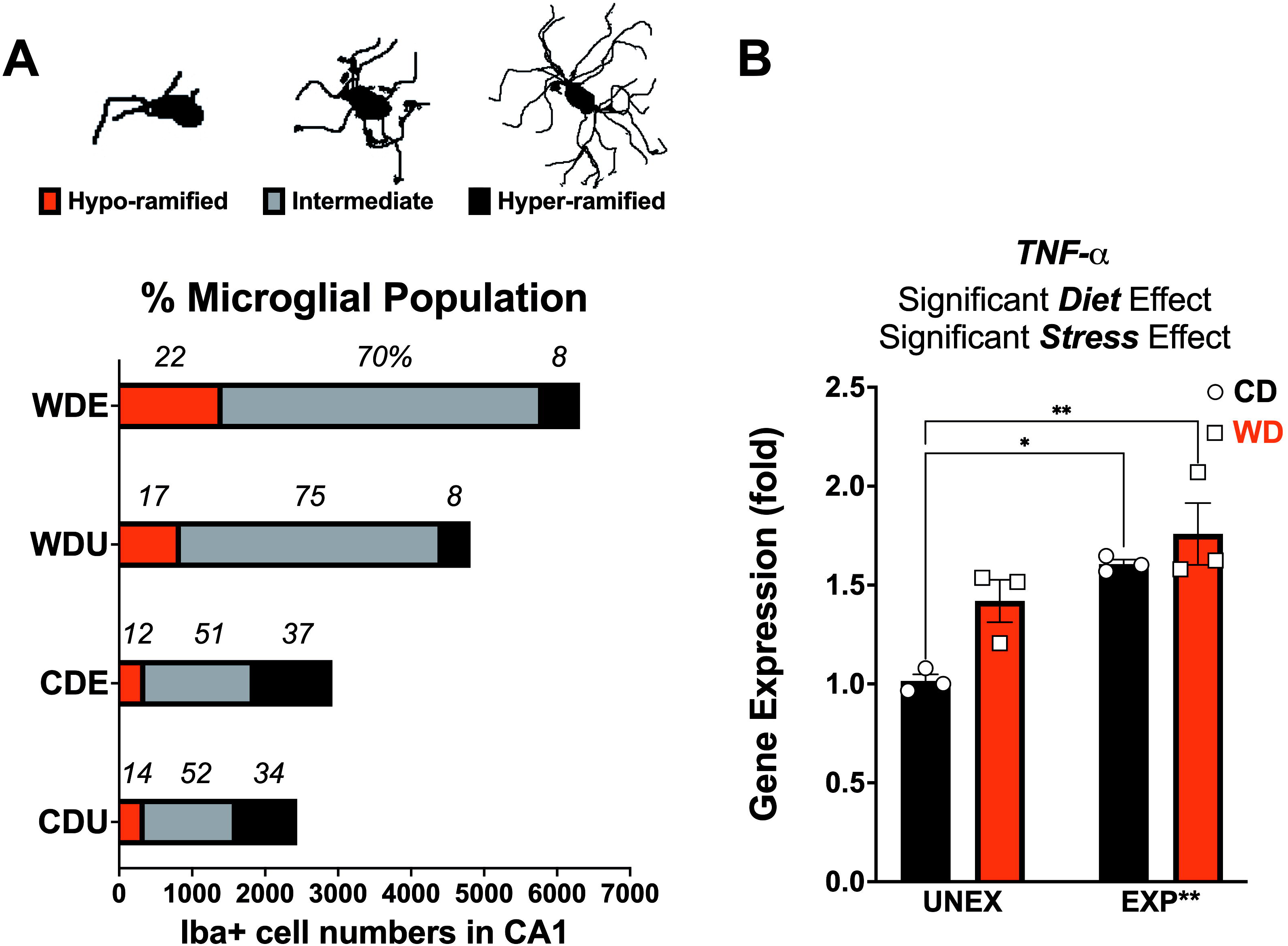
Obesogenic environments uncover Hypervigilant microglia population in the CA1 subfield of the hippocampus. **(A)** Representative images of the 3 distinct morphological phenotypes: Hyper-ramified, Intermediate, and Hypo-ramified identified. Microglial cell distribution in the percentage of cell populations showing diet and stress increased hyper-ramified and intermediate phenotypes (n, χ², df, p; 6,487, 1982, 6, *p* < 0.0001). **(B)** Hippocampal TNF-α gene expression showed to be significantly influenced by both diet (*p* = 0.0208) and stress (*p* = 0.0014). Stress significantly increased CD (*p* = 0.0110), and WD (*p* = 0.0028) fed animals. Sample size: neurohistology: 4 rats/group, gene expression: 3 rats/group.

### WD intake increases vulnerability to PSS-induced anxiety-like behaviors in the EPM

The elevated plus-maze (EPM) test is widely used to study anxiety-like behavior in rats. We examined EPM behaviors in adult rats after the exposure to the obesogenic conditions. We found that WD consumption reduced the percent of time in the open arms (*p* = 0.0014) (**Fig 2C**). Evaluation of the total number of closed arms entries as a proxy of motor activity showed no significant difference among all experimental groups (**Fig 2D**). This finding indicates that the indices of anxiety observed are not related to changes in ambulation. Other ethologically relevant behaviors were measured and corroborated the anxiogenic effect of the WD on EPM measures (**S Fig 4**). PSS rats that consumed the WD exhibited reduced frequency in the nose dipping zone relative to CD rats exposed to PSS (*p* = 0.0003) (**S Fig 4E**). Similarly, WD consumption significantly reduced the number of stretch attend postures (*p* < 0.0001) (**S Fig 4F**). Altogether, these showed that WD rats exhibited increased anxiety-like behaviors in the EPM, mainly when rats were also exposed to PSS.

### PSS exposure during adolescence reduces indices of sociability

We used the social Y maze to enable observation and evaluation of WD and PSS effects on social choices. We found a significant effect of PSS (*p* = 0.0256) and WD (*p* = 0.0273) in the number of social interactions with a caged counterpart (**Fig 2E**). We also discovered that the ratio of interactions between the conspecific and the object was significantly reduced in WD rats (*p* = 0.0465) (**Fig 3F**). Interestingly, the total distance traveled in the 9-min test was reduced by PSS relative to unexposed controls (*p* = 0.0328) (**S Fig 4G**). The sociability index was calculated to correct the total distance traveled and analyses revealed that PSS reduced this metric (*p* = 0.0060) (**S Fig 4H**). In summary, these results demonstrate that this early trauma model that incorporates social instability significantly impacts social behaviors in adult rats. A complete report of the statistical details from behavioral assessments can be found in **Table 1 and Supplementary Table 3**.

### Adolescent WD and PSS exposure influence hippocampal microglia in adult rats

Microglia are a dynamic population of resident immune cells in the brain responsible for regulating development and maintenance. Several studies, including ours, reported on microglia’s phenotypic and functional heterogeneity under obesogenic conditions. Specifically, our previous multimodal imaging and quantitative microglial morphometric analysis showed a significant response to a short exposure to an obesogenic diet in the CA1 region of the hippocampus. Interestingly, these changes were associated with increased anxiety-like behaviors in rats.^24^

In this study, we investigate the effects of chronic stress and an obesogenic diet in the CA1 hippocampal with a multiparameter quantitative morphometric analysis of Iba1+ microglia and a comprehensive neuroanatomical integrity approach (**Figure 3, 4**).

### WD consumption increases hippocampal microglia cell numbers and promotes a hyper-ramified phenotype

Morphological analyses of 16,250 Iba1+ cells, exploring counts, dimensions, and fractal heterogeneity, revealed precise effects of the WD and PSS on hippocampal microglia. Representative composite Iba1+ microglial fields (**Fig 3A**) and their binarized masks (**Fig 3A’**) show distinct morphological features analyzed using a FracLac Box Counting Algorithm. In short, grids are systematically scaled over each object (cell) to determine their fractal dimensions (change in pixel value over a spatial pattern). The increase in grid count and complexity allow differentiating cells by morphological features, as shown in representative cells for each experimental group (**Fig 3B**). Quantification of the total number of Iba1+ cells demonstrated that WD increased microglial cell numbers (*p* = 0.0044). Notably, PSS and WD synergized to increase hippocampal microglia cell numbers relative to unexposed controls (*p* = 0.0287), suggesting that the inflammatory response to the stress is potentiated by the obesogenic diet (**Fig 3C**).

Principal component analysis (PCA) was performed to identify morphological features driving group differences. In agreement with prior findings, we identified lacunarity, span ratio, density, and circularity as important differentiating features. Evaluating lacunarity, a measure of morphological heterogeneity sensitive to changes in particular features such as soma size relative to process length, we found that WD significantly decreased this metric (*p* = 0.0325), suggesting a proinflammatory microglial phenotype (**Fig 3D**). Previous fractal evaluations of microglia morphology have found that lacunarity decreases as the distribution becomes more scattered by the cell cycling to a more activated state.^97^ Span ratio measures, a parameter that accounts for microglial dimensions and shape and accounts for the ratio of the major and minor axes of the outermost points of the cell, revealed that PSS exposure significantly decreased the span ratio (*p* = 0.0064) (**Fig 3 E)**. Evaluation of microglial density as a measure of cell solidity showed that the WD significantly decreased microglial density (*p* = 0.0285) and mean radius (*p* = 0.0352), a measure of cell roundness (**S Fig 8 B, C**). Circularity describes a biologically relevant shape feature, as it accounts for measures of the ratio of convex hull area to the perimeter. In our study, circularity was significantly affected by PSS (*p* = 0.0250) and increased (*p* = 0.045) in rats that consumed an obesogenic WD and were exposed to PSS (**S Fig 8D**). These findings prompted us to further investigate relationships between these parameters to understand better the biology of the responses to PSS and WD. We identified lacunarity (form factor) and solidity as the main features impacted by the experimental manipulations (**S Fig 8A**).

### An obesogenic environment increases a hyperactive-like microglia population in the CA1 subfield of the hippocampus

We used *K*-means clustering to classify cells into three phenotypes based on lacunarity: hyper-ramified, intermediate, and hypo-ramified. The cell distribution in **Fig 4A** shows that the WD increased hypo-ramified phenotypes relative to controls (approximately 7% increase). PSS amplified the effect of the WD in expanding this cell population relative to controls (22% vs. 14%). Our data suggest that microglia appear prepared to transition into fully hypo-ramified (hyperactive-like) states when conditioned with the obesogenic diet. We sought to confirm this notion by measuring the mRNA expression of the tumor necrosis factor-alpha (TNF-α) in the hippocampus. In agreement with the morphological findings, the WD (*p* = 0.0208) and PSS (*p* = 0.0014) increased TNF-α gene expression in the hippocampus. The combination of the obesogenic conditions (PSS and WD) had a robust effect on TNF-α upregulation relative to controls (*p* = 0.0028) (**Fig 4C). Table 2 and Supplementary Table 4** detail the microglial morphology statistical measures.

### The obesogenic conditions affect peripheral cytokine levels selectively

Obesity and stress are associated with a marked dysregulation in the circulating levels of several inflammatory mediators that can influence microglia activities. We evaluated a panel of thirteen peripheral cytokines using a bead-based flow cytometry immunoassay. Considering main effects, WD significantly affected the circulating levels of IL-6 (decreased; *p* = 0.0115) and the granulocyte-macrophage colony-stimulating factor GM-CSF (increased; *p* = 0.0009). PSS increased IL1-α levels in plasma (*p* = 0.0047). Notably, the WD and PSS interacted to alter the levels of IL1-α (*p* = 0.0033), IL- 10 (*p* = 0.0175), GM-CSF (*p* = 0.0091), IL-12p70 (*p* = 0.0211), and IL-33 (*p* = 0.0169). **Supplemental Figures 13, 14, and Tables 12 and 13 include** the results and statistics of the cytokines tested in this study.

### Neuroimaging helps to establish links between structural abnormalities in hippocampal subfields and microglia morphology

Understanding the factors contributing to alterations to normal neural development that manifest as behavioral disorders in childhood and the mechanisms mediating these changes are essential for proper diagnosis and intervention. We investigated the relationship between the morphological features predictive of microglial-mediated hippocampal dysfunction with a multimodal imaging approach using magnetic resonance imaging (MRI)-derived diffusion tensor imaging (DTI) and neurite orientation dispersion and density imaging (NODDI) analysis. Representative images of the parameters investigated can be found in **Supplemental Figure 9**. A list of descriptive concepts for each parameter measured can be found in **Supplemental Table 5**.

The diffusion tensor model of DTI is based upon three orthogonal axes of diffusion, yielding a diffusivity index from which fractional anisotropy (FA) can be estimated. NODDI distinguishes restricted diffusion in the intracellular compartment, hindered diffusion in the extracellular compartment, and free diffusion in cerebrospinal fluid (CSF), from which parameter maps can be estimated. **Supplemental Figure 10** shows heatmaps illustrating the impact of the WD and PSS on each neuroimaging metric. We selected the eleven (11) hippocampal subfield and formation regions from the atlas. **Supplemental Tables 6 and 7** include the statistical details of these analyses.

Analyses of hippocampal subfields and formation regions revealed that FA was significantly altered by the WD (*p* < 0.0001), PSS (*p* = 0.0022), and the interactions between these factors (*p* = 0.0219), indicating changes in structural integrity. Mean, axial, and radial diffusivity indices were also significantly disturbed by the WD (mean: *p* = 0.0304) (axial: *p* = 0.0357) (radial: *p* = 0.0284) and PSS (mean: *p* = 0.0296) (axial: *p* = 0.0333) (radial: *p* = 0.0295). These findings demonstrate restricted water motion due to microstructural changes. Indices of orientation dispersion, intracellular volume fraction, and Log Jacobian showed to be significantly modified by the WD (orientation dispersion: *p* < 0.0001) (intracellular volume fraction: *p* < 0.0001) (Jacobian: *p* = 0.0023), and interaction effects (orientation dispersion: *p* = 0.0131) (intracellular volume fraction: *p* = 0.0057) (Jacobian: *p* = 0.0270). Details of these analyses are found in **Supplemental Figure 10 and Supplemental Table 6**.

### WD and PSS promote gut microbiota dysbiosis associated with pro-inflammatory microglia and structural neuroadaptations in the hippocampus

Preclinical and clinical evidence supports the concept of bidirectional brain-gut microbiome interactions. We aimed to determine the effects of adolescent WD and PSS in gut microbial composition (**S Fig 11**). Detailed statistics can be found in **Supplemental Table 8**. PCA of the unweighted Unifrac distance matrix from the rarefied data was used to evaluate the presence of clusters or groupings based on the operational taxonomic unit (OTU)-level microbial features. These analyses indicated that diet type was a significant contributor to their microbial signatures (*p* < 0.001) and discriminated between the two groups regardless of exposure (*p* = 0.78) (**S Fig 11A**). Unexpectedly, global parameters of relative bacterial richness (*p* = 0.12) (**S Fig 11B**) and diversity (*p* = 0.19) (**S Fig 11C**) were similar among groups. Taxonomic classification at the family (**S Fig 11D**) and genus (**S Fig 11E**) levels showed a selective distribution of Firmicutes and Bacteroidetes in each condition. Interestingly, the relative abundance of Bacteroidaceae–Prevotellaceae, a bacterial family associated with the production and transport of secondary metabolites, including short-chain fatty acids (SCFA), was significantly decreased by the exposure to PSS (*p* = 0.0040) (**S Fig 11F**). It has been shown that a reduction in SCFAs leads to increased gut permeability,^98^ thus increasing exposure to bacterial endotoxins and neuroinflammation. Microbiome evaluation revealed that the WD (*p* = 0.0033), PSS (*p* = 0.0210), and the interactions between these factors (*p* = 0.0052) significantly contributed to the relative abundance of *Lachnospiraceae NK4A136.* Relative to unexposed controls, the WD (*p* = 0.0005), PSS (*p* = 0.0048), and the combination of WD and PSS (*p* = 0.0019) significantly decreased the relative abundance of *Lachnospiraceae NK4A136* (**Fig 5A**). Statistical details of microbiome analysis can be found in **Table 4**.

**Figure 5.**
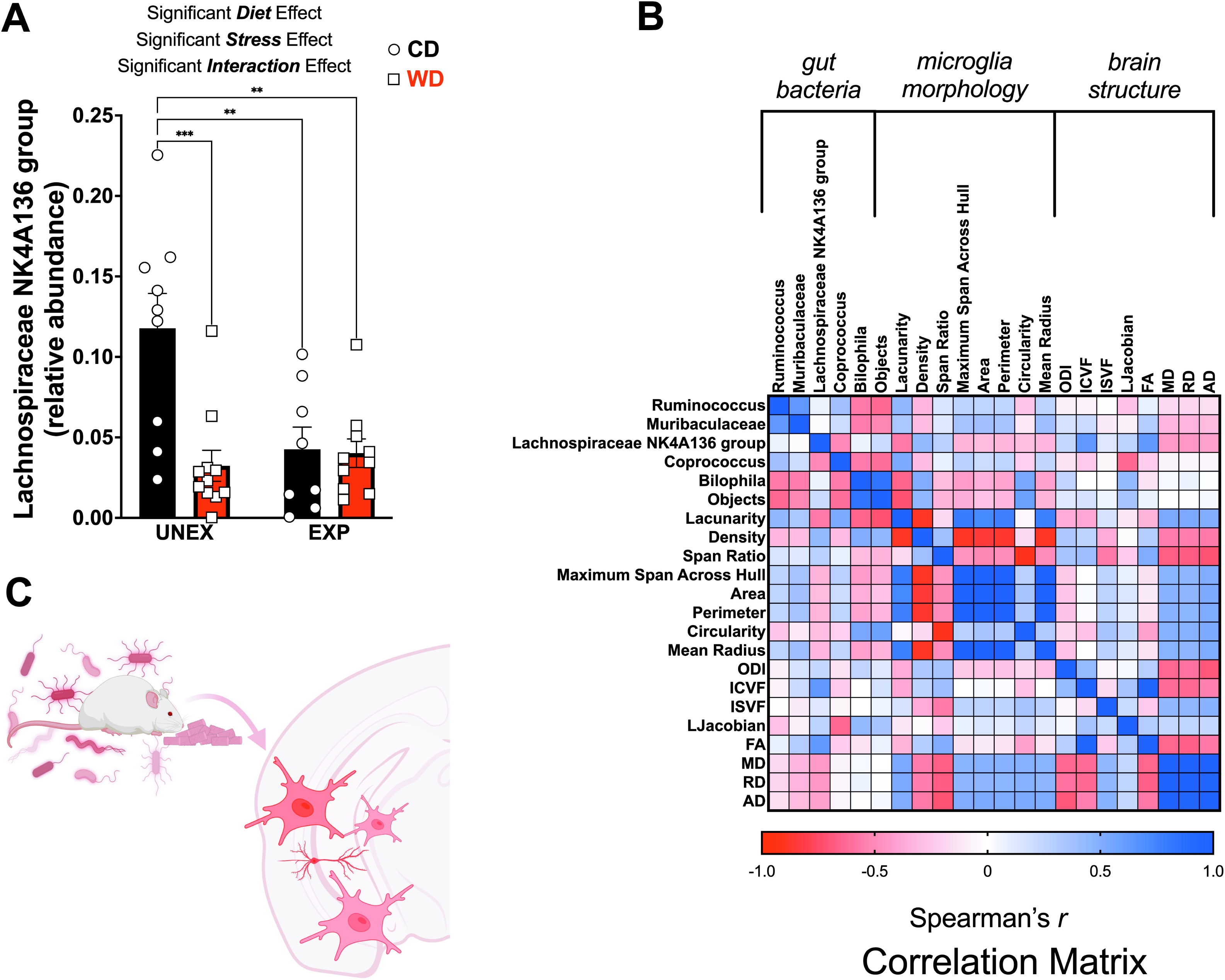
WD diet and chronic stress promote effects in the gut-brain axis. **(A)** Microbiome evaluation revealed diet (*p* = 0.0033), stress (*p* = 0.0210), and interactions of diet and stress (*p* = 0.0052) significantly contributed to the relative abundance of Lachnospiraceae NK4A136. Multiple comparisons analysis showed a significant decrease of Lachnospiraceae NK4A136 by WD (*p* = 0.0005), Stress (*p* = 0.0048), and the combination of WD and PSS (*p* = 0.0019). **(B)** Spearman’s rank correlation coefficient analysis showed significant associations between brain structure, microglial morphology, and the relative abundance of several members of the Firmicutes phyla. **(C)** Schematic of the hypothesis from the previous findings: microbiome influences microglia; in turn, microglia may impact hippocampal structure. Sample size: 4 rats/group.

Spearman’s rank correlation coefficient analysis showed differential associations within brain microstructure parameters and key microglial morphological features with microbiome dysbiosis (**Fig 5B**). While no direct links are established, these data validate the notion that the microbiome influences microglia activities, which in turn can change brain structure and function. These results suggest that diet and stress exert selective adaptive effects on gut microbiome diversity and potential microbiota remodeling, increasing susceptibility to metabolic and cognitive deficits (**Fig 5C**).

### Obesogenic diet and chronic stress influence neuroendocrine response

We sought to evaluate the neuroendocrine response by measuring the primary glucocorticoid in rodents, corticosterone, as metabolites in excreta (fecal corticosterone metabolites, FCM). In male rodents, FCM accounts for 75% to 90% of the corticosterone eliminated from the body, with the remainder excreted in the urine. A range of stressors can activate the HPA axis and promote corticosterone release, facilitating prompt adaptation to environmental demands. Chronic adaptive states promote a state of increased allostatic load, which can alter hippocampal plasticity and contribute to the development of neuropsychiatric disorders.^99^

We found that WD intake reduced FCM *(p* = 0.0018) at four weeks of diet consumption (**Fig 6A**). The WD *(p* = 0.0052), PSS *(p* = 0.0008), and the interaction between those factors *(p* = 0.0039) were found to significantly influence FCM levels 24 h after the first predator exposure (**Fig 6B**). Of importance, only WD rats exhibited increased FCM 24 h after the predator stress exposure relative to unexposed controls *(p* = 0.0012). Predatory stress significantly increased FCM levels at 24 h following the second exposure *(p* = 0.0250) (**Fig 6C**). Statistical detailed information can be found in **Table 5**.

**Figure 6.**
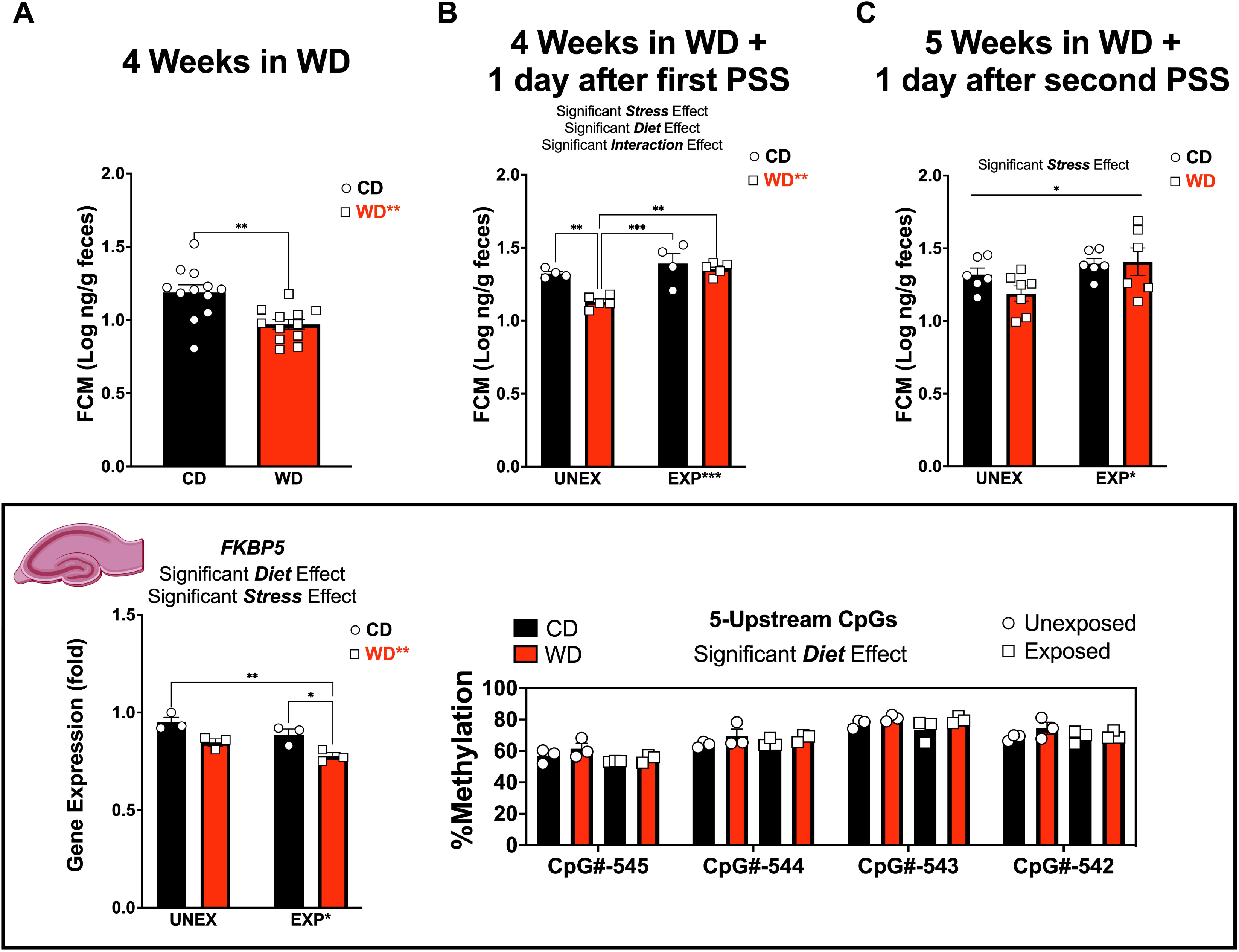
Obesogenic diet and chronic stress influence FCM and hippocampal *Fkbp5* gene methylation status at specific sites. **(A)** Fecal corticosterone metabolite (FCM) release was significantly decreased by diet after 4 weeks of WD consumption (*p* = 0.0018). **(B)** FCM was significantly influenced by the interaction of diet and stress (*p* = 0.0390) and by diet (*p* =0.0052) and stress (*p* = 0.0008) alone at 1 day after the first psychosocial stress exposure. **(C)** 5 weeks and 1 day after the second psychosocial stress exposure, FCM remained significantly influenced by stress (*p* = 0.0250). **(D)** Gene expression of fkbp5 in the hippocampus was significantly decreased by diet (*p* = 0.0016) and stress (*p* = 0.0196). **(E)** Diet had a significant effect (*p* = 0.0024) in the percent of methylation in four (4) significant (*p* < 0.0001) 5 Upstream CpGs (CpG#-545, CpG#-544, CpG#-543, CpG#-542). Sample size: 4 rats/group.

### Obesogenic diet and chronic stress influence hippocampal Fkbp5 gene methylation status at specific sites

The Fkbp5 gene encodes the Fk506 binding protein 5, an essential regulator of glucocorticoid activity and acute stress response. Interaction of FKBP5 with intronic regulatory glucocorticoid response elements (GREs) leads to activation and increased transcription. As part of a negative feedback loop, FKBP5 attenuates glucocorticoid signaling via a dynamic process of methylation/de-methylation. Early life traumatic experiences are thought to imprint long-standing patterns of demethylation,^100^ however, the specific CpG sites and how diet interacts with chronic stress to alter epigenetic marks remained understudied. We found that the WD *(p* = 0.0016) and PSS (*p* = 0.0196) significantly decreased hippocampal Fkbp5 mRNA levels (**Fig 6C**). The WD had a significant effect (*p* = 0.0024) in the percent of methylation in several upstream CpGs (CpG#-545, CpG#-544, CpG#-543, CpG#-542) (**Fig 6D). Table 6** shows the detailed statistical analysis.

### FKBP5 integrates microglial pro-inflammatory signals under obesogenic conditions

To study inflammatory mechanisms linking obesity to stress susceptibility, we used human immortalized microglial cell lines clone 3 (HMC3).^101^ HCM3 cells have been extensively characterized with respect to cell morphology, antigenic profile, and cell function and reliably show most of the original antigenic properties and express specific microglial markers. Palmitic acid (PA) is the most abundant free saturated fatty acid present in the WD. This fatty acid produces a metabolic, inflammatory response associated with damaging mechanisms such as oxidative stress. We sought to determine whether PA would replicate the dysregulated responses of endocrine and immune biomarkers observed in our model. HCM3 cells culture and treatment conditions were optimized to determine cell viability and the half-maximal inhibitory concentration (IC_50_). HCM3 cells treated with increasing concentrations of PA (0-800 µM) showed no significant difference in the percent of viability between cells treated with PA or PA + hydrocortisone. IC_50_ for PA was determined to be 33.52 µM, and PA + hydrocortisone was 50.84 µM. HMC3 cells treated with increasing concentrations of hydrocortisone (0- 1000 nM) and viability assays showed a 50% of growth inhibition (IC_50_) at 119.3 nM (**S Fig 21**). HCM3 cells were primed with PA (50 µM) or vehicle for 24 hours, and hydrocortisone (100 nM) was added for 24h. (**Fig 7A**). TNF-α acts as a pro-inflammatory cytokine during the acute phase stress response and exerts potent neuromodulatory effects. Two focal effects of TNF-α are increased inflammation and immunomodulation. To this end, we examined TNF-α gene expression in cell lysates and evaluated its protein release in the supernatant. TNF-α gene expression was significantly increased by CORT treatment alone (*p* = 0.0146). Interestingly, while PA alone decreased TNF-α gene expression (*p* = 0.0102), PA pretreatment and CORT increased its TNF-α mRNA levels (*p* = 0.0003) (**Fig 8B**). TNF-α protein release levels were significantly influenced by CORT (*p* = 0.0002), PA (*p* = 0.0096), and the interaction of PA + CORT (*p* = 0.0021) (**Fig 7B, 7C**). We found that PA + CORT significantly increased secreted TNF-α protein levels, supporting the increased mRNA levels. Microglia produce reactive oxygen species (ROS), essential in cell death by activating c-Jun N-terminal kinase. ROS formation was expressed as RFU and normalized to protein (**Fig 8C**). ROS was significantly increased by CORT (*p* < 0.0001), PA (*p* < 0.0001), and by the combination of PA+CORT (*p* = 0.0459). In support of these findings, we observed increased M2 polarization under PA+CORT conditions (**Fig 7D, E**). These results were confirmed via immunofluorescence staining of HCM3 with microglia activation marker cluster of differentiation 68 (CD68) and imaged with confocal microscopy. The obesogenic environment caused by priming with PA and exposure to CORT showed the characteristic morphological changes of microglial activation with enhanced phagocytic activity (**S Fig 22, 23**). Statistical analyses are detailed in **Tables 7 and 8**.

**Figure 7.**
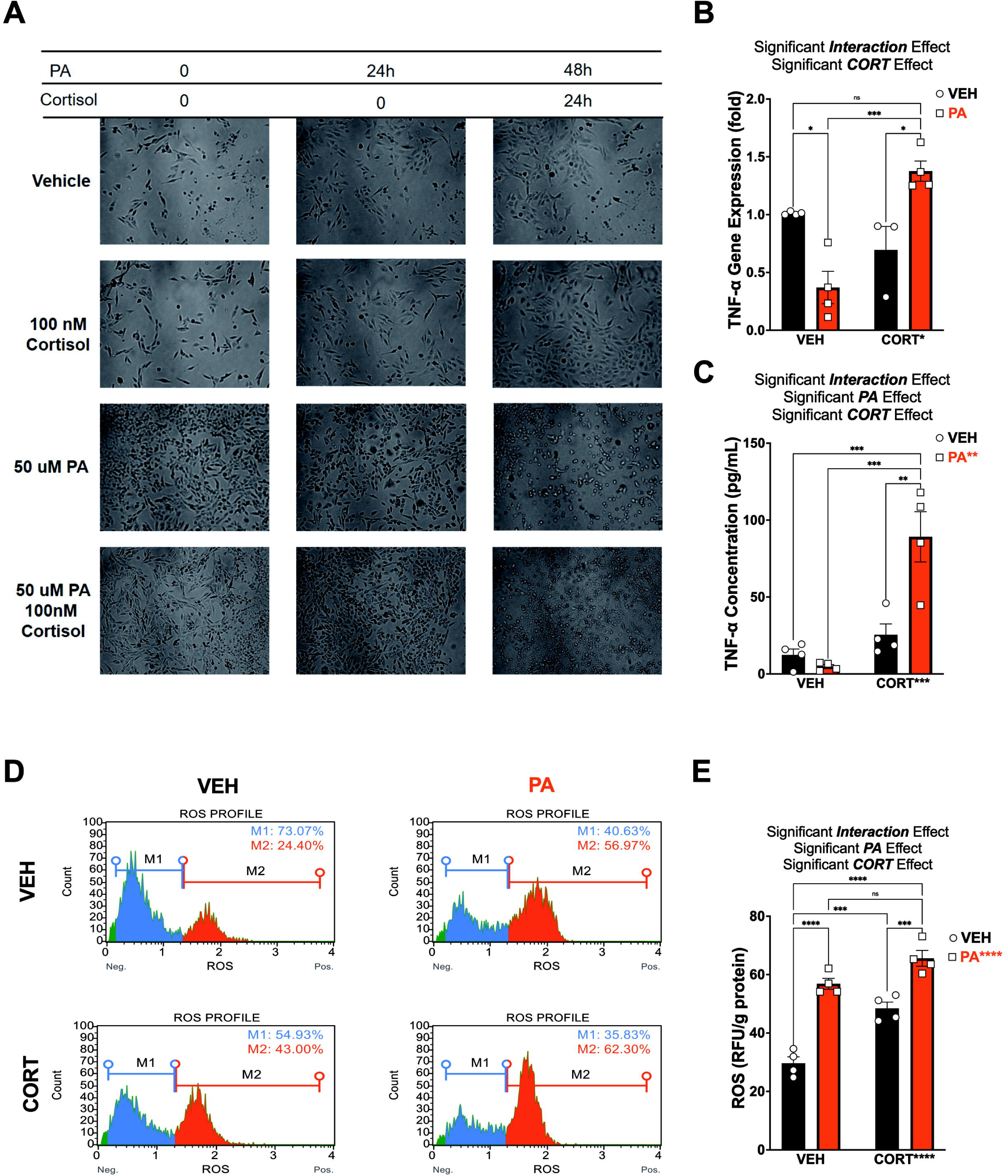
FKBP5 integrates microglial pro-inflammatory signals under obesogenic conditions. **(A)** Representative brightfield images of Human microglial cells (HCM3 cells) pretreated with Vehicle, Palmitic Acid (PA) at 50 µM and Cortisol (CORT) at 100 nm and in PA alone and combination. **(B)** TNF-α Gene expression was significantly influenced by PA+CORT interaction (*p* = 0.0002) and CORT alone (*p* = 0.0146). **(C)** TNF-α protein expression was also significantly influenced by PA+CORT (*p* = 0.0021) and PA (*p* = 0.0096), and CORT (*p* = 0.0002) alone. **(D)** Reactive Oxygen Species (ROS-Intracellular superoxide) were detected by dihydroethidium (DHE) measurements, showing an increase of M2 polarization **(E)** ROS expression normalized by grams of protein measurements was also significantly influenced by PA+CORT (*p* = 0.0459), and PA (*p* < 0.0001) and CORT (*p* < 0.0001) alone. RFU: relative fluorescence unit. Error bars represent standard errors (n = 3). \**p* < 0.05, \*\**p* < 0.01 and \*\*\**p* < 0.001 \*\*\*\**p* < 0.0001 by Two Way ANOVA. Sample size: 4 rats/group.

### FKBP5 plays a crucial role in neuroendocrine immunometabolism regulation

We evaluated the influence of PA and CORT regulation of the FKBP5 signaling in microglial function in our system. We found that FKBP5 gene expression was significantly decreased by PA (*p* < 0.0001), CORT (*p* = 0.0038), and PA+CORT (*p* = 0.0275). CORT treatment significantly increased FKBP5 expression (p = 0.0049), and PA pretreatment in CORT-treated cells showed a blunted expression of FKBP5 (p = 0.0002)—**Figure 8** and **S Table 21**. Interestingly, FKBP5 Gene expression in HCM3 cells treated with increasing concentrations of CORT (0-1000nm) showed an interesting rise and fall pattern, suggesting a dampening of the expression at high CORT concentrations (**S Fig 24A**).

**Figure 8.**
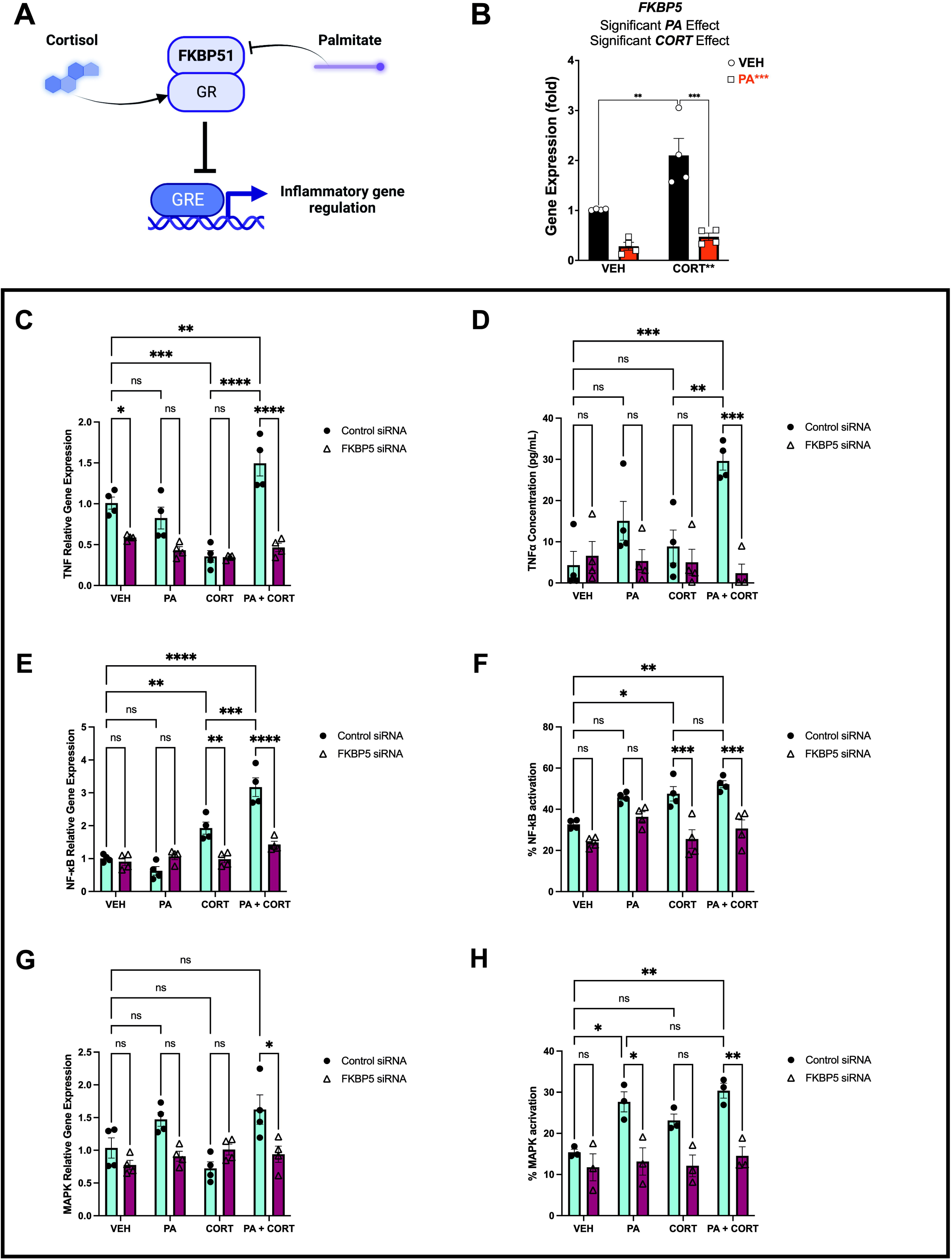
FKBP5 plays a key role on neuroendocrine immunometabolism regulation in HCM3. **(A)** Schematic representation of PA+CORT regulation of FKBP5 signaling **(B)** FKBP5 Gene expression was significantly decreased by PA (*p* < 0.0001), CORT (*p* = 0.0038) and PA+CORT (*p* = 0.0275). Sample size: 4 rats/group. **FKBP5 plays a key role on neuroendocrine immunometabolism regulation in HCM3. (C)** After silencing FKBP5, TNF-α Gene expression was significantly influenced by Treatment + SiRNA interaction, Treatment and SiRNA (*p* < 0.0001). **(D)** TNF-α protein release was significantly influenced by Treatment + SiRNA interaction (*p* = 0.0012), Treatment (*p* = 0.0206) and SiRNA (*p* = 0.0004). **(E)** NF-κB Gene expression was significantly influenced by Treatment + SiRNA interaction, Treatment and SiRNA (*p* < 0.0001) **(F)** NF-κB percent activation was significantly influenced by Treatment + SiRNA interaction (*p* = 0.0350), Treatment (*p* = 0.0003) and SiRNA (*p* < 0.0001). **(G)** MAPK Gene expression was significantly influenced by Treatment + SiRNA interaction (*p* = 0.0039), Treatment (*p* = 0.0064) and SiRNA (*p* = 0.0024). **(H)** MAPK percent activation was significantly influenced by Treatment (*p* = 0.011) and SiRNA (*p* < 0.0001). Sample size: 4 rats/group.

To further investigate the role of FKBP5 in regulating immunometabolism, we used small interfering RNA (siRNA) to silence FKBP5. We evaluated the response and activation of essential pathways involved in the immunoregulation of the stress response. We found TNF-α mRNA levels were significantly influenced by treatment + siRNA interaction, Treatment, and SiRNA (*p* < 0.0001). TNF-α protein release was significantly influenced by Treatment + siRNA interaction (*p* = 0.0012), Treatment (*p* = 0.0206), and SiRNA (*p* = 0.0004). NF-κB gene expression was significantly influenced by treatment + siRNA interaction, treatment, and siRNA (*p* < 0.0001). NF-κB activation was significantly influenced by treatment + siRNA interaction (*p* = 0.0350), treatment (*p* = 0.0003), and siRNA (*p* < 0.0001). Similarly, MAPK gene expression was significantly influenced by treatment + siRNA interaction (*p* = 0.0039), treatment (*p* = 0.0064), and siRNA (*p* = 0.0024). MAPK activation was significantly influenced by treatment (*p* = 0.011) and siRNA (*p* < 0.0001). **Figure 8C** and **Table 9** show the statistical details of these analyses. Altogether these data demonstrate that FKBP5 plays a crucial role in neuroendocrine immunometabolism regulation, and appropriate levels of expression seem to be required for a normal response.

## DISCUSSION

Despite the high comorbidity between obesity and stress-related mental disorders, the pathways contributing to this bidirectional association remain relatively unexplored. This study investigated the biological underpinnings of early exposure to an obesogenic environment characterized by access to an obesogenic diet and chronic psychosocial stress. The main findings are as follows: **1)** we demonstrate that early chronic psychosocial stress leads to increased food intake and weight gain, **2)** early exposure to an obesogenic diet and exposure to psychosocial stress impair social behaviors while increasing fear and anxiety-like behaviors, **3)** the obesogenic conditions altered the structural integrity within specific hippocampal formation regions and subfields, **4)** these conditions were associated with significant shifts in gut microbiome distribution, neuroinflammation, and immunometabolic alterations, **5)** mechanistically, we identified the corticosterone receptor chaperone gene FKBP5, as a potential molecular link connecting obesogenic conditions to altered inflammation and stress reactivity. This study identifies novel mechanisms and adaptations by which obesogenic environments shape the maturational trajectories of common neurobiological correlates of stress resilience.

Stress disorders, including post-traumatic stress disorder (PTSD), represent a public health challenge. In the United States, approximately 40% of high school students are subjects of violence, and 6% are diagnosed with PTSD.^102, 103^ Growing epidemiological evidence shows that PTSD is linked to obesity and cardiometabolic complications.^104–112^ Psychosocial stress is a prevalent environmental factor that contributes to the development of obesity. On the other hand, mounting evidence supports that obesity and metabolic disorders confer vulnerability to stress-related disorders.^63, 64, 113, 114^ The findings presented herein support that obesity predisposes individuals to stress-related complications. Consumption of foods rich in saturated fats and refined sugars has been strongly correlated with the modulation of emotional states. While high palatable foods can relieve negative emotions by dampening perceived stress and anxiety in animals and humans, these effects have been found temporary and pose a risk for increased consumption, obesity, and heightened vulnerability to depression and anxiety.^115^

We found that the obesogenic environment altered acoustic startle reflex increased of fear-potentiated startle, and reduced sociability. Sociability is a fundamental and adaptive aspect of adaptive biology. In rodents, social recognition of conspecifics is important to maintain social hierarchy and normal development of social cognition. Moreover, several neuropsychiatric disorders are characterized by disrupting social behavior, including anxiety, depression, and obsessive-compulsive disorders. Studies have tried identifying and quantifying molecular biomarkers or signatures that complement the behavioral test. Research has uncovered the critical role of hippocampal subregions vCA1 and dCA2 in regulating social recognition, supporting our neuroimaing and behavioral findings.^116^ Recent studies have implicated corticosteroid hormones as potential modulators of social memory. Stimulation early in life resulted in significant enhancement of adult rat social recognition associated with a significant decrease in basal corticosterone. Interestingly recognition was highly negatively correlated with basal corticosterone levels, which indicates that elevated basal corticosterone could impair an individual’s social recognition performance.^117^ Since oxytocin is a vital regulator of the HPA axis, these interactions might explain the decreased sociability in our model when animals are exposed to stress.

Microglia have been shown to participate in pruning mechanisms while the brain is developing primarily and in the mature brain, providing trophic support. Emerging research suggests that the maturation and function of selected neurocircuits may be regulated by the molecular identity that microglia adopt on different regions in the brain.^118^ Given the increased sensing ability of microglia to environmental cues, defective microglia may significantly contribute to the function and maturation of the neurocircuitries affected by obesogenic environments.

The gut microbiome plays a crucial role in neural development, modulating neurotransmission and inflammation and affecting behavior.^119^ Increased levels of pro-inflammatory cytokines and environmental stress activate the limbic system. We found that stress induced a reduction in the abundance of short-chain fatty acid (SCFA)- producing bacteria. Our results expand on the increasing number of reports of studies and animals and humans that link the ontogeny and pathology of stress-related disorders to the microbiome composition.

Epigenetic changes and environmental factors can influence the effects of these allele-specific variants, impacting the response to GCs of the FKBP5 gene. The decreased methylation level of FKBP5 was observed in adolescents with persistent depressive symptoms. Early life stress is thought to imprint long-standing patterns of demethylation resulting in chronically dysregulated feedback and risk for PTSD.^120, 121^

## LIMITATIONS AND FUTURE STUDIES

In this study, we sought to evaluate the brain, behavior, and immunometabolic responses to an obesogenic environment in adolescent rats. We initially postulated that the early consumption of an obesogenic WD would exacerbate the adverse effects of adolescent trauma on the brain and behavior. Future studies should include female rats to investigate whether the sex differences observed in humans are recapitulated. Moreover, including female rats can provide robust evidence for the mechanisms studied here. Additionally, performing the two-hit model in different rat strains can provide insight into strain-dependent differences and confounding variables. Testing different housing conditions can help describe the effects of social isolation. Future studies are needed to examine causal mechanisms and determine the involvement of other specific brain regions implicated with anxiety and stress-related disorders.

### Images and Figures

Illustrations were prepared using BioRender (www.biorender.com), OmniGraffle Pro (The Omni Group; Seattle, WA), and GraphPad Prism version 9.4.1.681 (GraphPad Software, San Diego, California USA, www.graphpad.com).

## Acknowledgments

The authors thank the Advanced Magnetic Resonance Imaging and Spectroscopy (AMRIS) facility and the National High Magnetic Field Laboratory (NHMFL) for their continued support (National Science Foundation Cooperative Agreement No. DMR- 1157490 and the State of Florida).

The authors would like to acknowledge the Animal Care Facility at LLU. Additionally, we would like to honor with a special mention the predators used for this study: Nahla, the female cat, and Chaos, the male cat; for their contribution to the validation of our model and the valuable findings of this study.

## Conflict of Interest statement

The authors declare no conflict of interest.

## Notes

### Competing Interest Statement

The authors have declared no competing interest.

