## Supplemental for "Early-life obesogenic environment integrates immunometabolic and epigenetic signatures governing neuroinflammation"

**SUPPLEMENTAL FIGURES**

***Supplemental Figure 1*. WD and PSS consumption increase food consumption and body weight. (A)** Summary of the daily caloric intake of rats from all experimental groups. Two-way ANOVA demonstrated that PPS was a significant factor contributing to increased caloric intake (*p* = 0.0033). Post hoc testing revealed that the PSS effect on caloric intake was particularly significantly in CD animals (*p* = 0.0230). **(B)** Body weight evaluation of percent change in body weight before (PND51) and after the PSS model (PND91) showed that both diet (*p* < 0.0001) and stress (*p* = 0.0004) are significant factors contributing to weight increase. Two-way ANOVA revealed a significant effect in body weight increase by diet alone (*p* = 0.0125), by stress alone in the WD-fed group (*p* = 0.0192), and among the exposed groups regardless of diet type (*p* = 0.0041). No significant increase in body weight was observed in the groups consuming CD when exposed to stress (*p* = 0.1028). Interestingly, no significant difference between animals that consumed WD and those consuming CD and exposed to the PSS model (*p* = 0.8836), suggesting that WD has a similar effect on weight gain than consumption of a CD under stressful environmental conditions. Notably, diet and stress synergize to significantly increase weight gain (*p <* 0.0001). Sample numbers: Caloric intake: CD UNEXP, *n =* 7; CD EXP, *n =* 8; WD UNEXP, *n =* 8, WD EXP; *n =* 8. Body weight CD UNEXP, *n =* 13; CD EXP, *n =* 14; WD UNEXP, *n =* 15, WD EXP; *n =* 14.

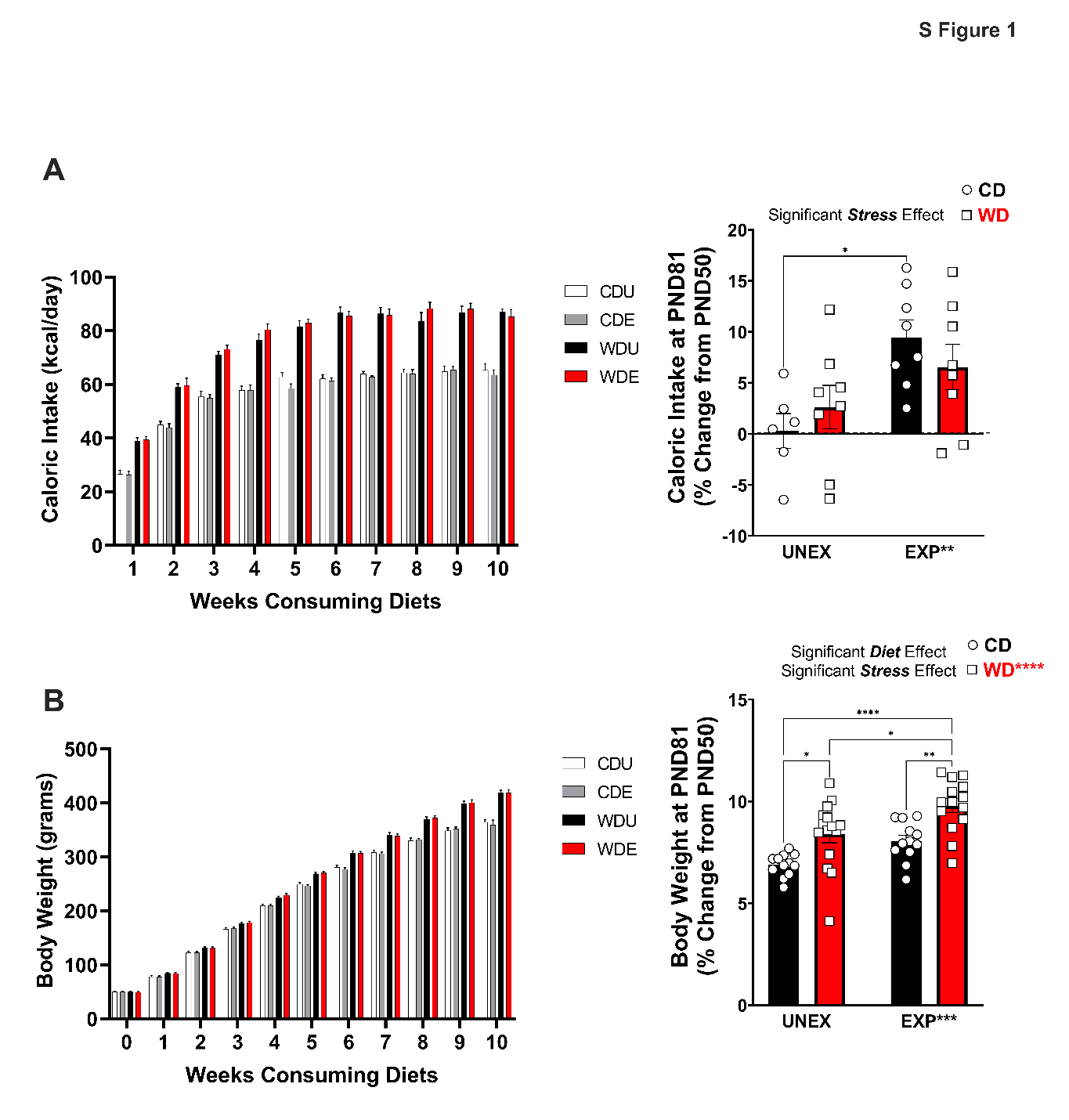

**Supplemental Figure 2. PSS induce decreases fasting glucose.** Fasting blood glucose levels (FBG) were measured at the end of the study. Stress had a significant effect on FBG (*p =* 0.0010); exposure to stress in CD-fed animals significantly decreased FBG (*p =* 0.0196) when compared to WD diet unexposed animals. Sample numbers: Fasting Glucose: CD UNEXP, *n =* 13; CD EXP, *n =* 14; WD UNEXP, *n =* 15, WD EXP; *n =* 14.

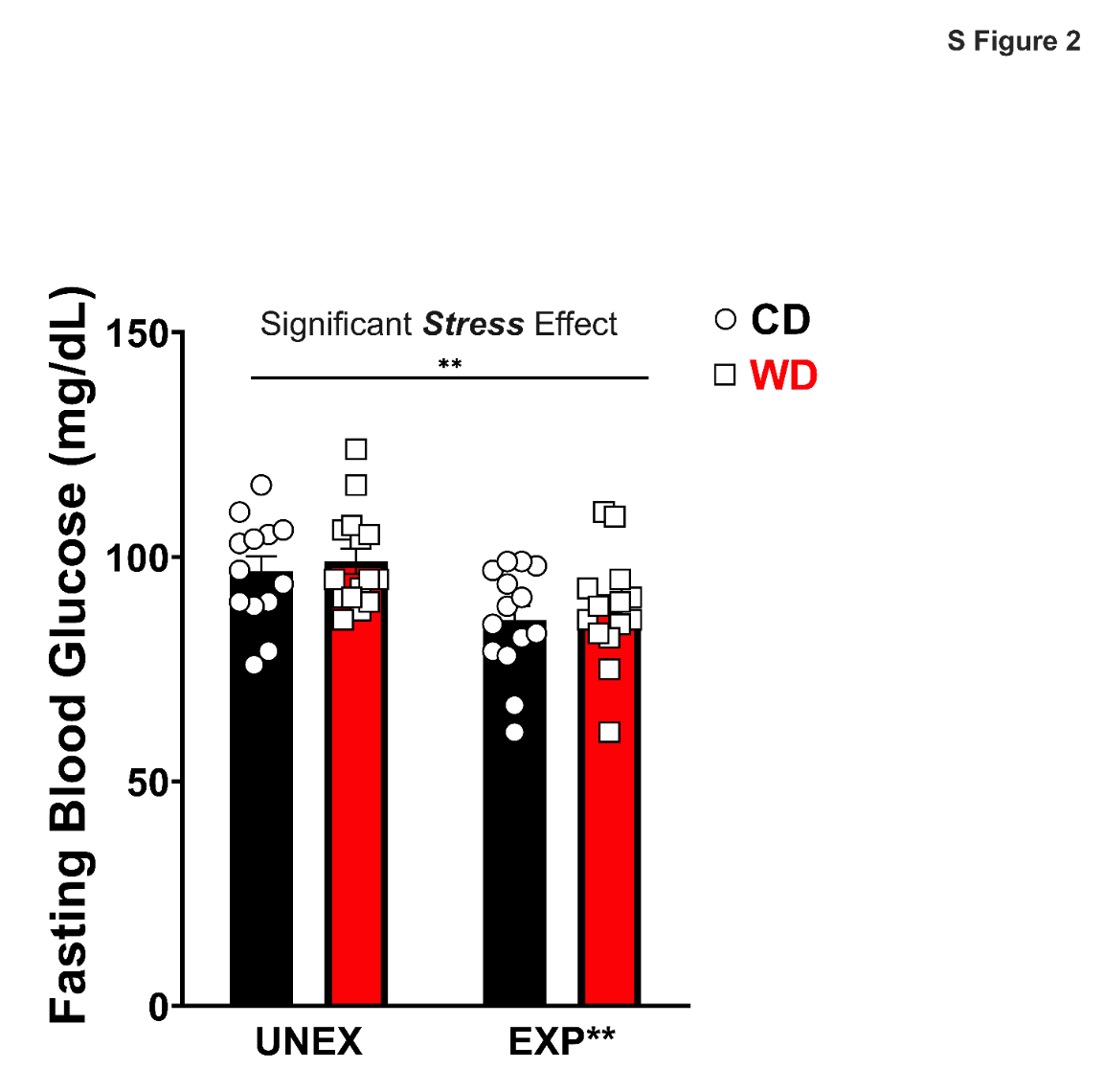

**Supplemental Figure 3. WD consumption decreased diastolic pressure and heart rate.** Measures of cardiovascular health showed no significant interactions or main effect of diet or stress in mean arterial blood pressure and systolic pressure. However, WD consumption alone had a significant contribution to the decrease of diastolic blood pressure (*p =* 0.0195) and heart rate (*p =* 0.0096). Sample numbers: CD UNEXP, *n =* 6; CD EXP, *n =* 7; WD UNEXP, *n =* 7, WD EXP; *n =* 7.

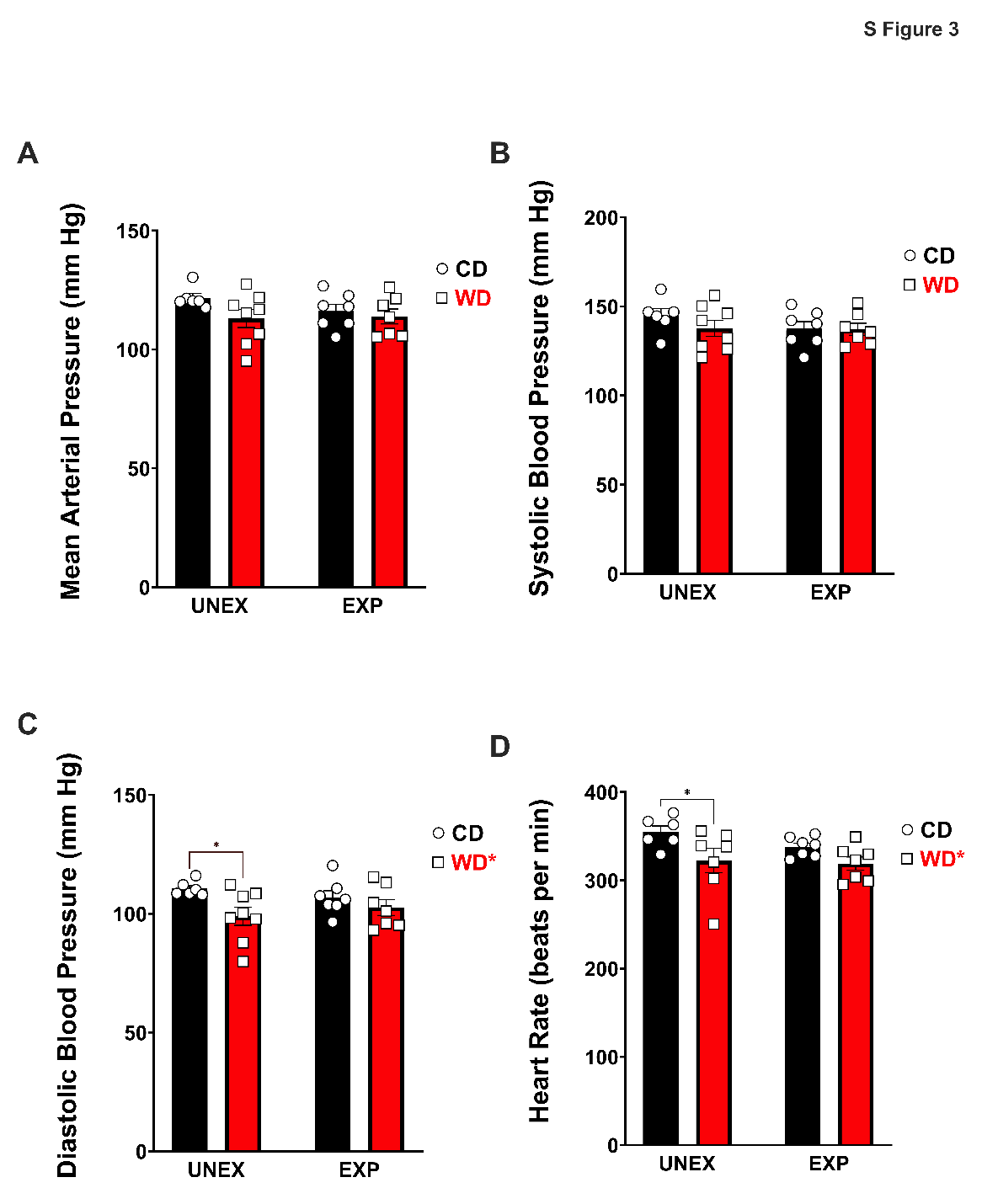

**Supplemental Figure 4. Behavioral phenotypes associated with early access to an obesogenic diet and psychosocial stress. (A)** There were no significant effects or interactions in ASR magnitude (*p* > 0.0050). **(B)** FPS Foot shock reactivity significantly decreased by WD consumption alone (*p =* 0.0302) and when combined with PSS (*p =* 0.0090). **(C)** Percent change from the FPS learning session had a significant Diet/Stress interaction (*p =* 0.0216). CD EXP rats showed a significant decrease (*p =* 0.0473) in FPS. **(D)** EPM Open arms entries were significantly decreased in WD EXP (*p =* 0.0120) and CD-exposed rats (*p =* 0.0392). Diet was a main contributor to the number of open arm entries (*p =* 0.0014) **(E)** EPM Nose in head pinning zone frequency showed a significant diet effect (*p =* 0.0002) and a significant decrease in the WD EXP animals (*p =* 0.0003). **(F)** EPM Stretch Attend frequency showed a significant diet effect (*p* < 0.0001). WD alone (*p =* 0.0031) and in combination with stress (*p =* 0.005) induced a significant reduction in the frequency of stretch-attend behaviors. **(G)** EPM Total Distance traveled was affected by stress (*p =* 0.0328). **(H)** SYM Sociability index corrected by distance traveled had a stress effect (*p =* 0.0060) **(I)** SYM Latency to first approach a conspecific showed no significant differences or main effects. For all behavioral samples: sample size = 14 rats/group (before outlier testing).

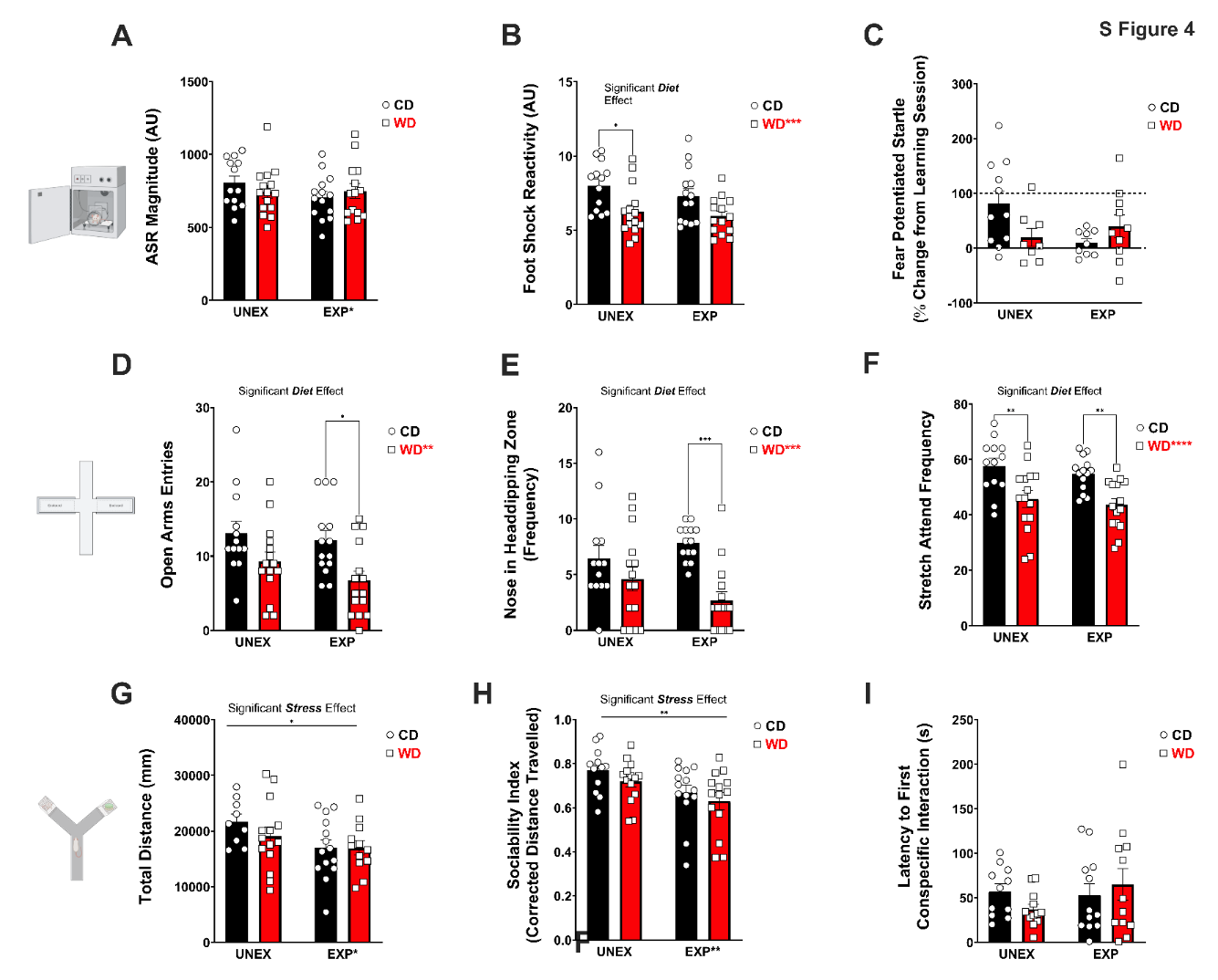

***Supplemental Figure 5*. Immunohistochemistry controls for Iba-1 immunodetection in hippocampal CA1.** Representative sections from the CA1 region of CD UNEXP rat were Iba-1 antibody was omitted. Nonspecific binding control validates the specificity of the commercial anti-Iba1 primary antibody. Brightfield image magnification: 10× and 40×. Scale bars: 100 microns

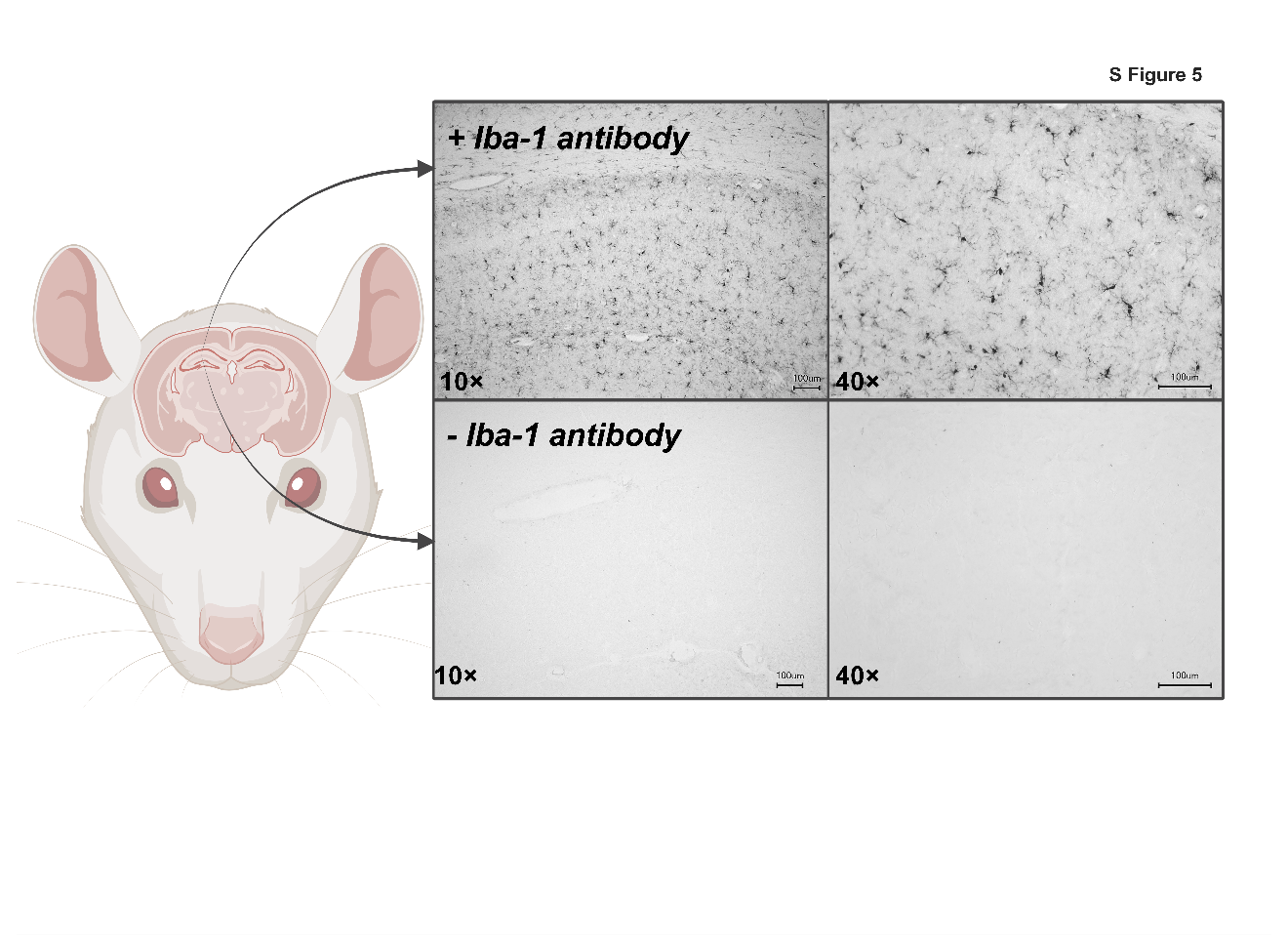

**6**

**Supplemental Figure 6. PC reveals distinctive microglial morphological descriptors.** Principal component (PC) analysis on seven (7) morphometric parameters in microglia. Circularity, Density, Span Ratio and Lacunarity were distinctly separated as morphological descriptors contributing to the morphological changes observed. Sample size:16,250 Iba-1+ cells, 4 rats/group.

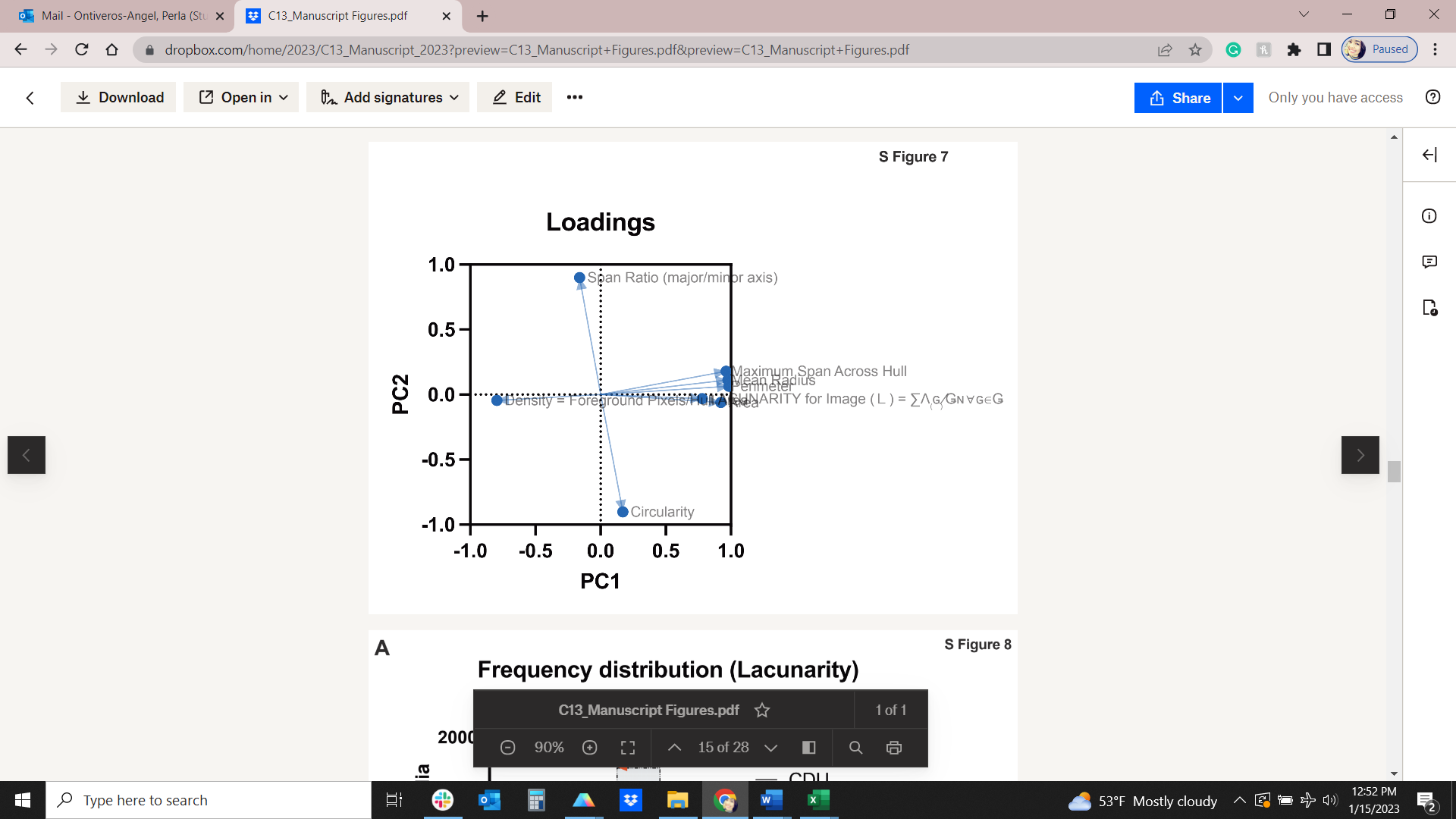

**Supplemental Figure 7. Frequency of microglial distribution. (A)** Frequency of cumulative distribution based on Lacunarity shows distinctly bimodal and upward trending populations (shifting towards each other) in WD and WD EXP animals **(B)** Span Ratio frequency distribution shows a unimodal right-skewed distribution with experimental group distinction in an upward forward trend when animals are exposed to an obesogenic diet and stress. Sample size:16,250 Iba-1+ cells, 4 rats/group.

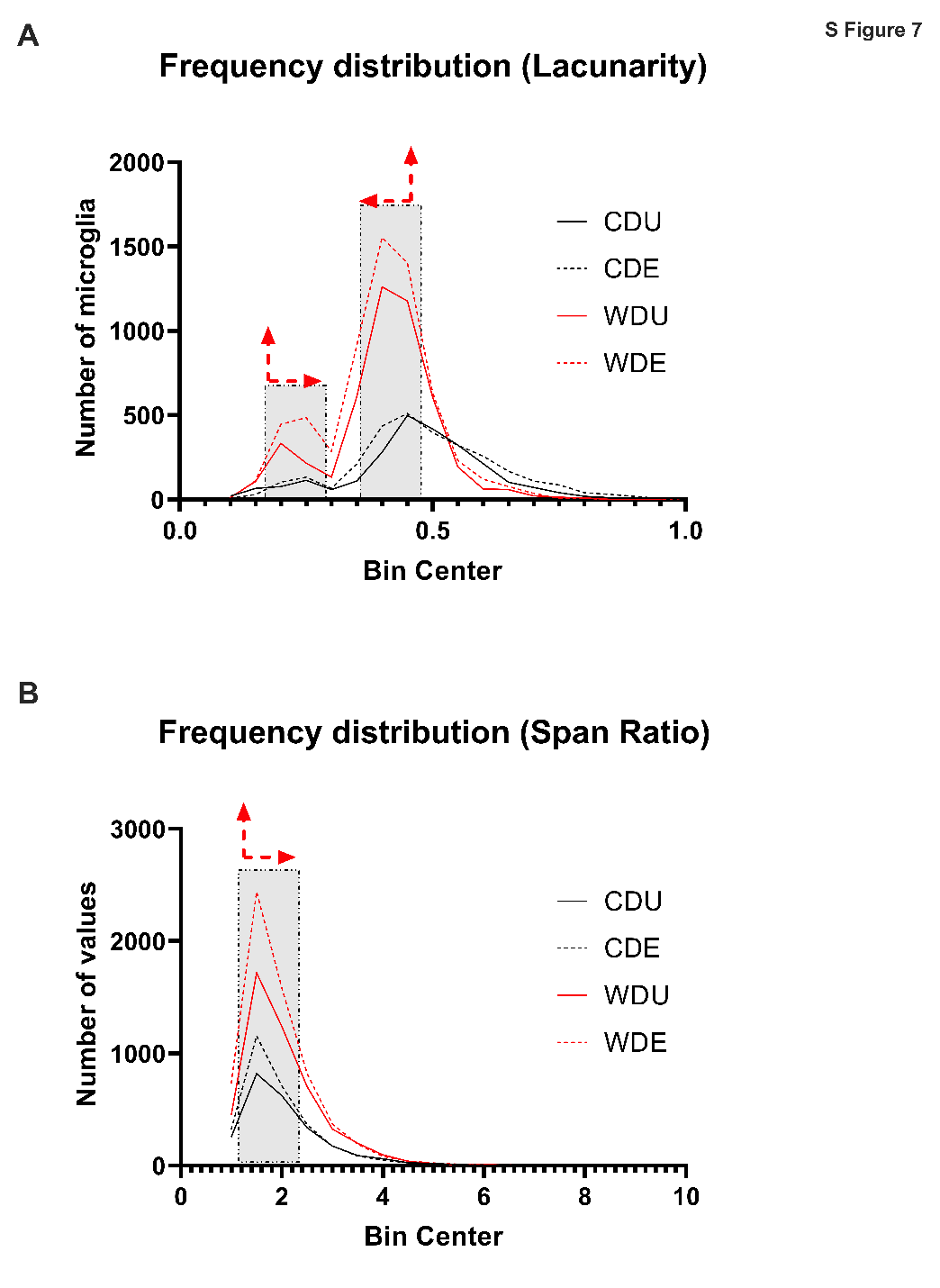

**Supplemental Figure 8. Clustering of microglia based on morphology and additional measures. (A)** *k*-means clustering based on solidity and form factor (Lacunarity) in each of the four experimental groups. Sample size:16,250 Iba-1+ cells, 4 rats/group. **(B)** Microglial density cell solidity) was significantly modified by diet (*p =* 0.0285) **(C)** Mean Radius (Roundness) was also modified by Diet (*p =* 0.0352). **(D)** Circularity was significantly modified by stress by stress (*p =* 0.0250) and increased (*p =* 0.045) in WD EXP rats. Sample size: 4 rats/group.

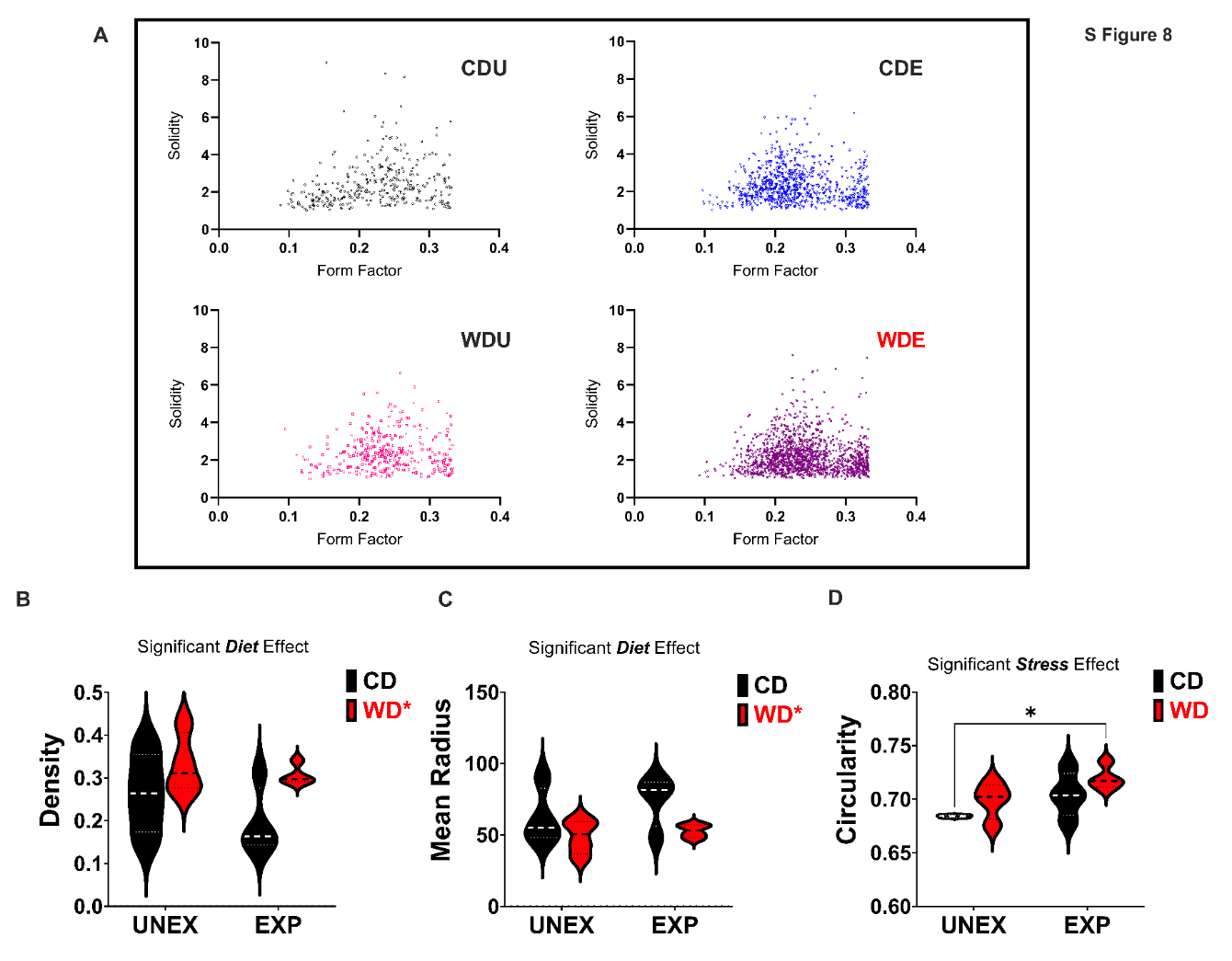

**Supplemental Figure 9. Representative images of brain parcellation in structural parameters using MRI.** Using Magnetic Resonance Imaging (MRI) derived Diffusion Tensor Imaging (DTI), B0 scan masks outlining rat brain boundaries were generated in MATLAB and adjusted manually using ITKSNAP. Independent images of corrected tensor element reconstruction for the Fractional Anisotropy (FA), Mean, Axial, and Radial Diffusivity (MD, AD, RD, respectively) parameters. MRI-derived neurite orientation dispersion and density imaging (NODDI) representative images of Orientation dispersion. Representative images of the rat brain template and the aligned B0 images, and the corresponding Log Normalized Jacobian Matrices.

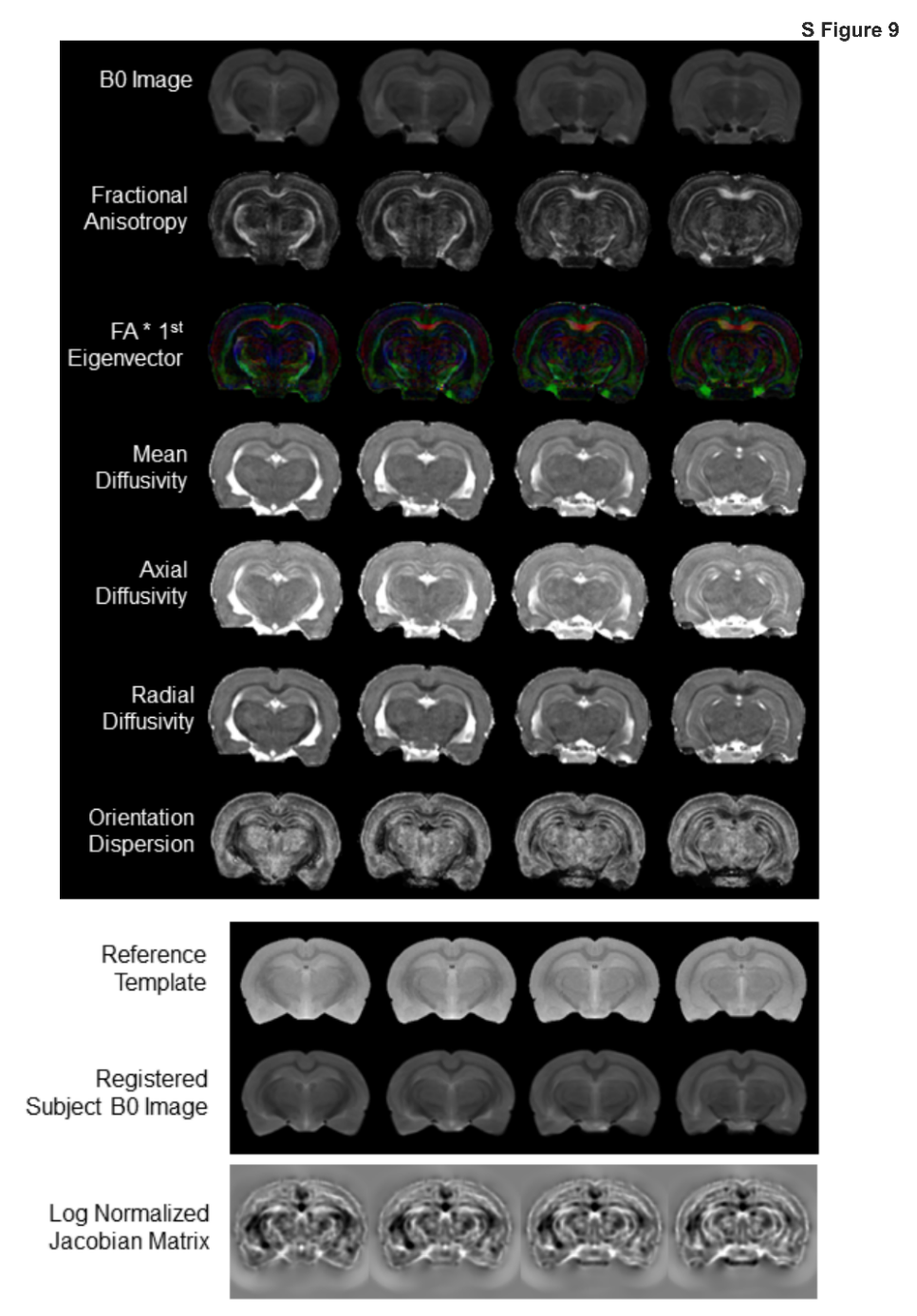

**Supplemental Figure 10. Heatmaps of hippocampal structural changes based on neuroimaging parameters.** Selected Parameters with contribution to microstructural integrity with region specific significance (*p* < 0.0001). **(A)** Fractional Anisotropy was significantly modified by diet (*p* < 0.0001), stress (*p =* 0.0022), and the interactions of diet and stress (*p =* 0.0219) **(B-D)** Medial, Axial and Radial Diffusivity were also significantly disturbed by diet (*p =* 0.0304) (*p =* 0.0357) (*p =* 0.0284) and stress (*p =* 0.0296) (*p =* 0.0333) (*p =* 0.0295) respectively. **(E-H)** Orientation Dispersion, Intracellular volume fraction, and Jacobian matrix indices derived from DTI and NODDI analysis showed to be significantly modified by diet (*p* < 0.0001) (*p* < 0.0001) (*p =* 0.0023), and diet x stress (*p =* 0.0131) *(p =* 0.0057) (*p =* 0.0270) interactions, respectively. Sample numbers: CD UNEXP, *n =* 6; CD EXP, *n =* 7; WD UNEXP, *n =* 8, WD EXP; *n =* 7.

**S**

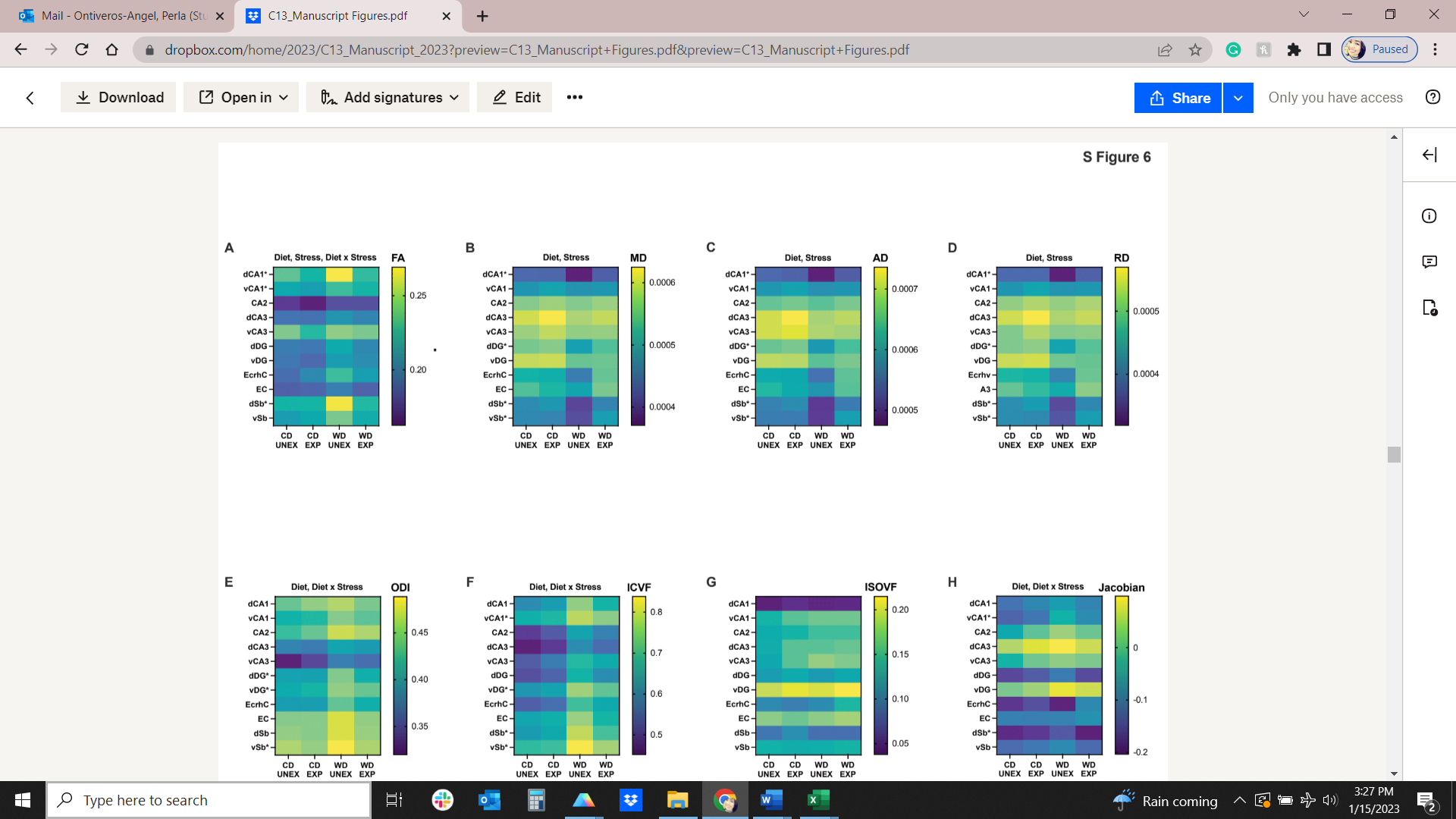

**Supplemental Figure 11. Microbiome analysis statistical details. (A)** PC Analysis showed a significant contribution of Diet (*p* < 0.001) to the explained variances. (B) Analysis of Chao1 (richness estimator) and Shannon (diversity index) did not show significant differences among the experimental groups. However, (D-E) Lower taxonomic ranks such as family and genus showed a distinct distribution among the groups. (F) Prevotellacea Relative Abundance was significantly reduced by stress (*p* = 0.004).

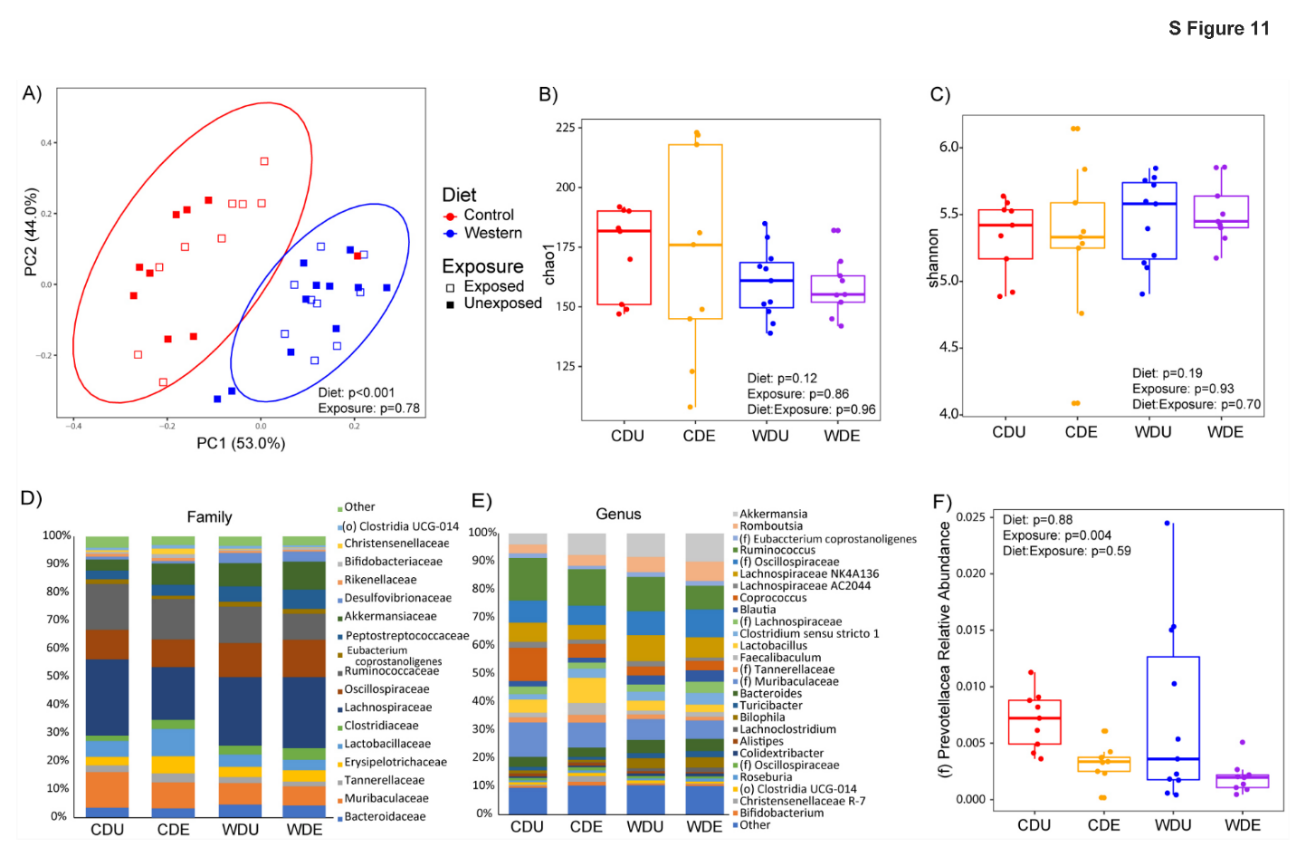

**Supplemental Figure 12A**. Statistical details of gut-brain correlation. Heatmap of *p*-value of the correlation between brain microstructure, microglia morphology, and selected bacteria groups.

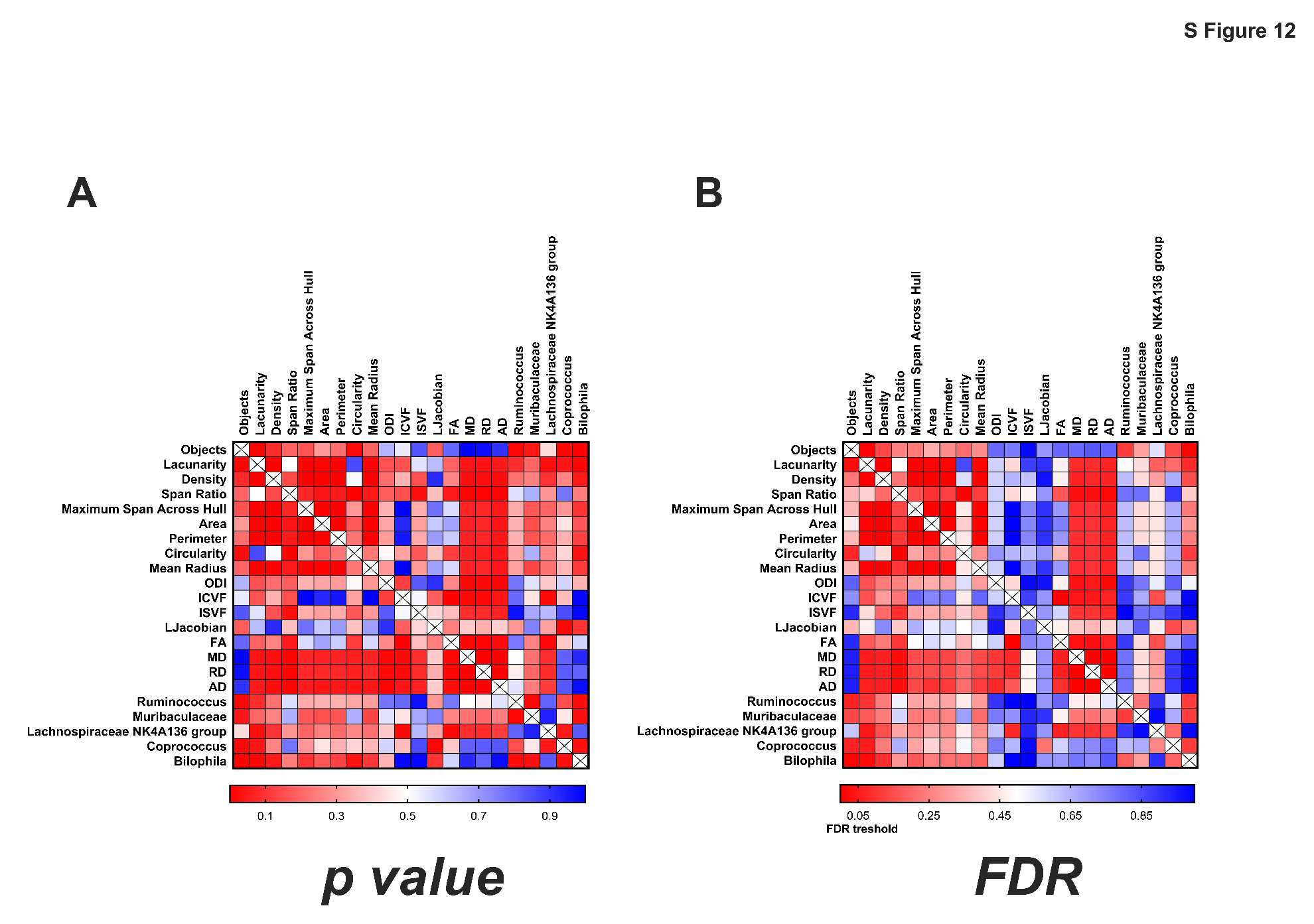

**Supplemental Figure 12B**. Statistical details of gut-brain correlation. Heatmap of False discovery rate (*FDR)* values of the correlation between brain microstructure, microglia morphology, and selected bacteria groups.

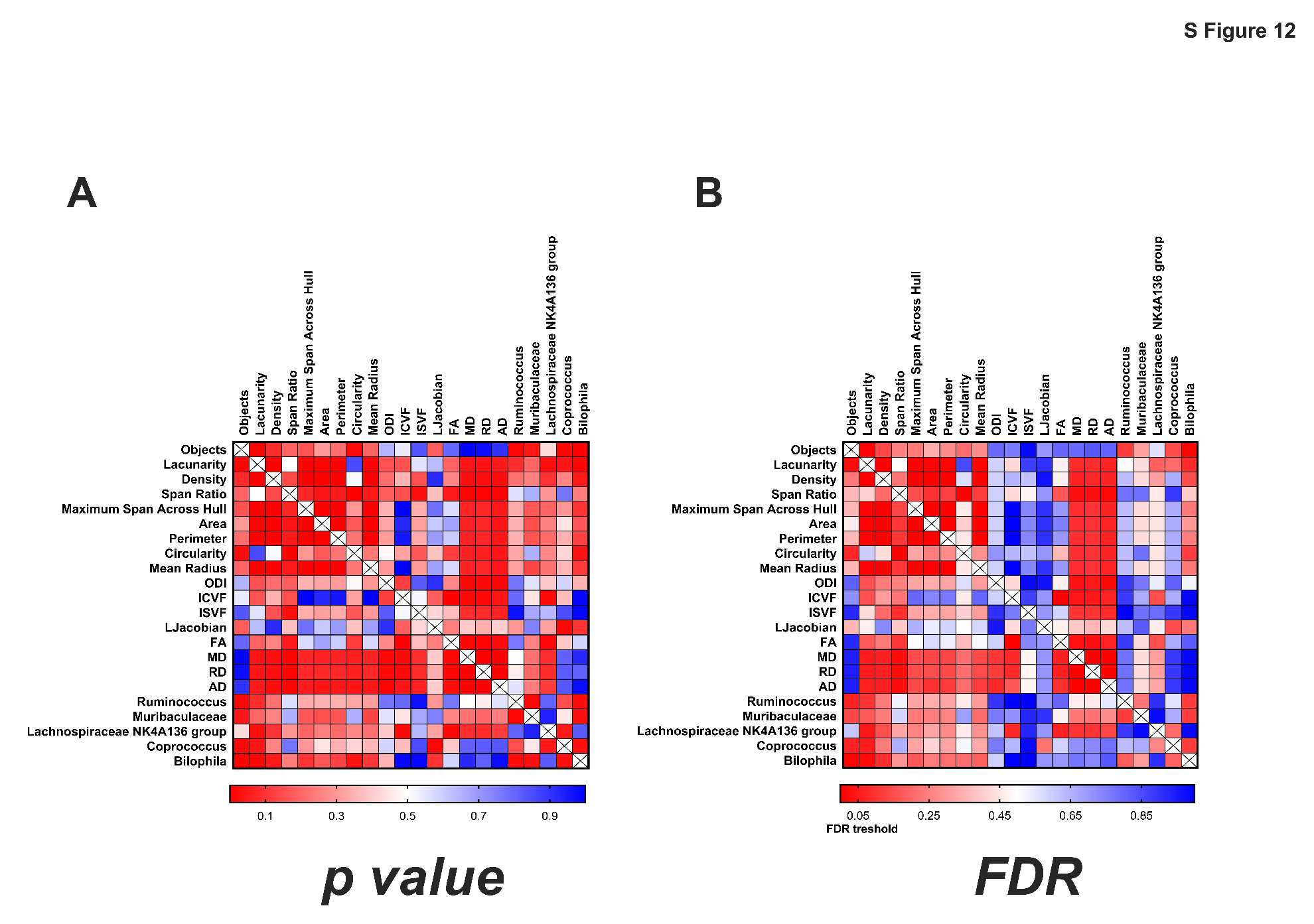

**Supplemental Figure 13. Evaluation of peripheral cytokines. (A) IL-6** circulating levels significant decreased by the diet (*p* = 0.010), **(B)** IL-1α had a significant interaction of diet and stress (*p* = 0.003) and a stress effect (*p* = 0.004), **(C)** IL-10 levels were significantly modified by the interaction of diet and stress (*p* = 0.017) **(D)** GM-CSF showed a significant increase by diet (*p* = 0.001) and diet and stress interaction(*p* = 0.009). **(E)** IL-12p70 had a significant interaction effect (*p* = 0.020)**,** and **(F)** IL-33 also had a significant interaction effect (*p* = 0.010). For a complete table of statistical analysis, see Supplemental Tables 13 and 14.

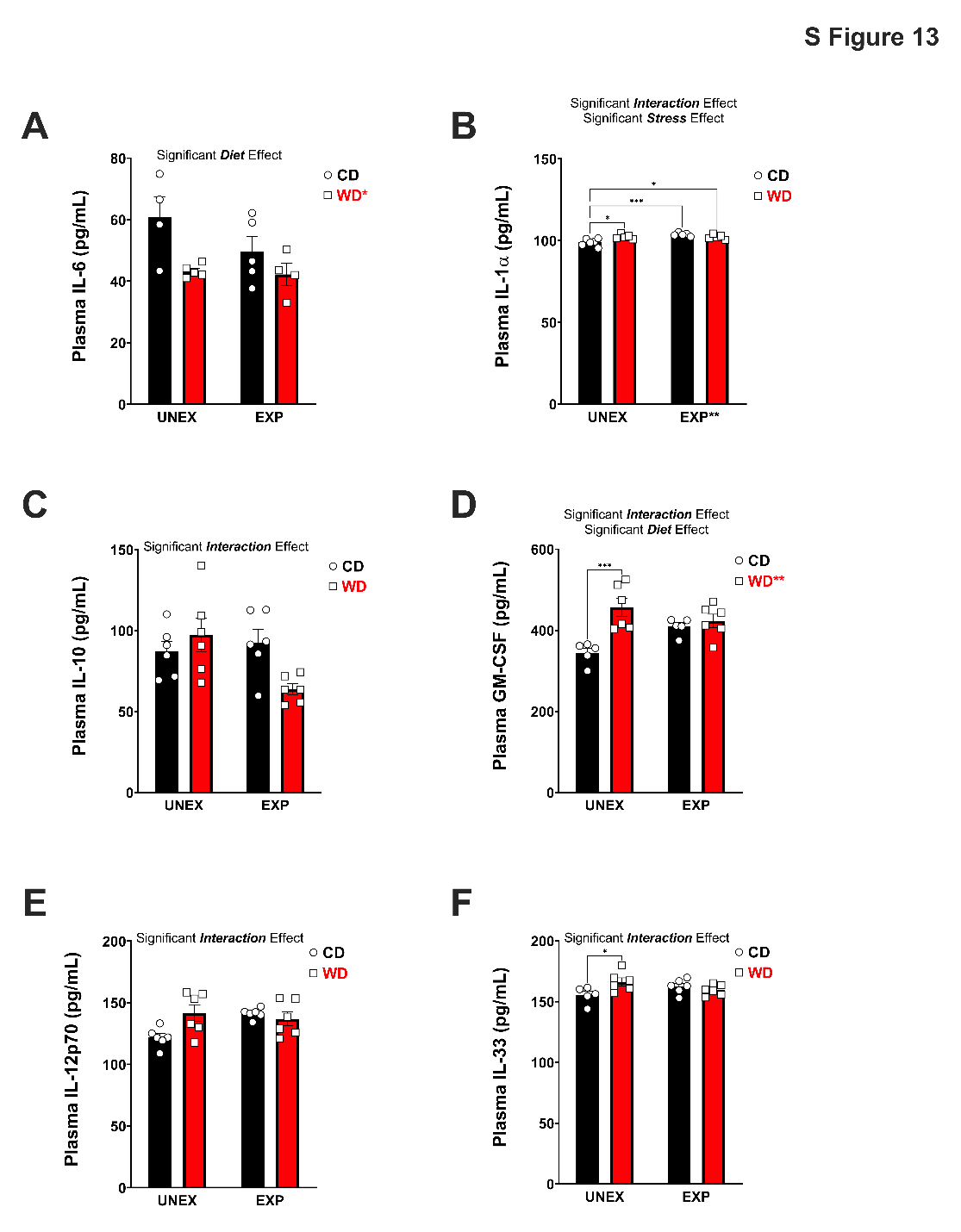

***Supplemental Figure 14*. Evaluation of supplemental peripheral cytokines. (A-E)** IFN-ϒ, CCL-2, IL-18, TNFα, and IL-17A circulating levels did not show any significant effect with Diet, Stress, or interactions of Diet and Stress. For a complete table of statistical analysis, see Supplemental Table 13.

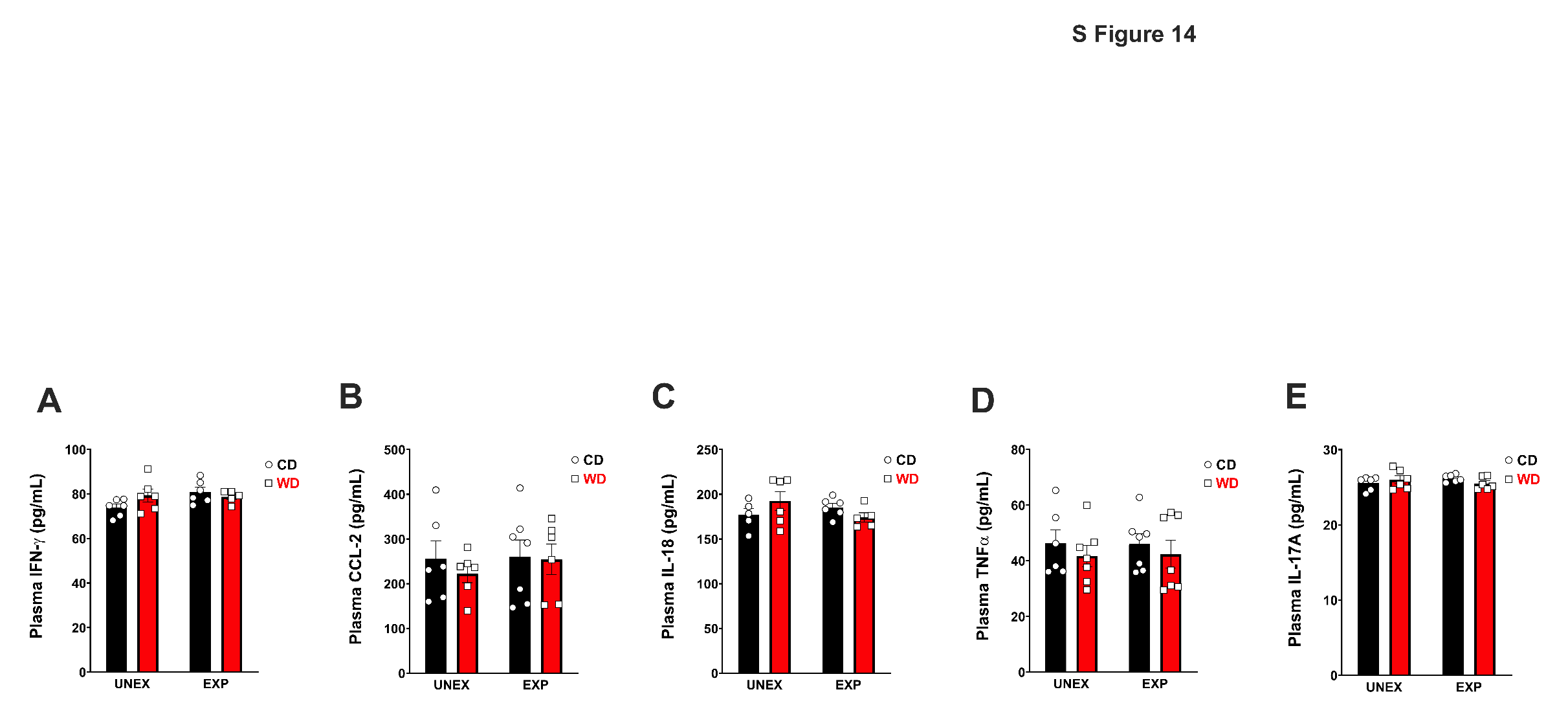

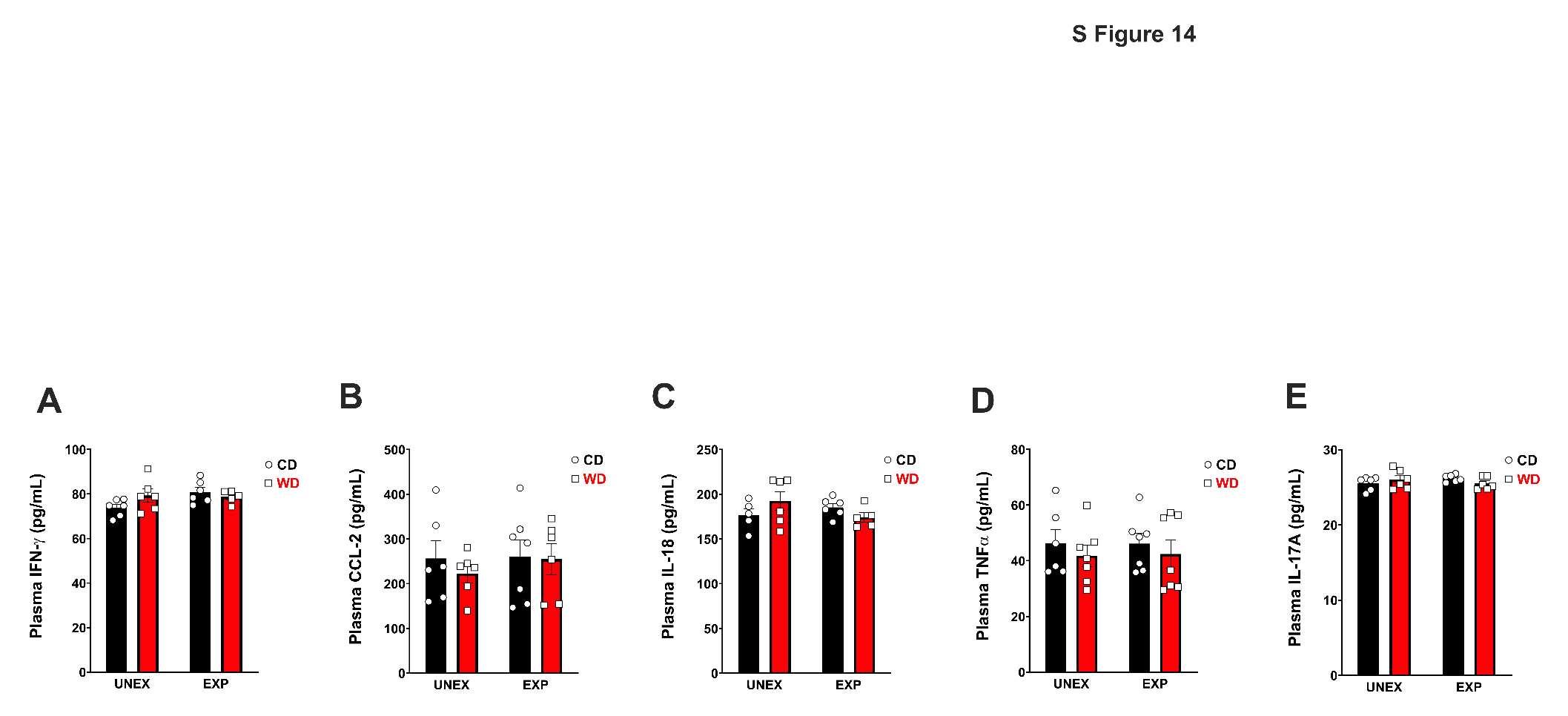

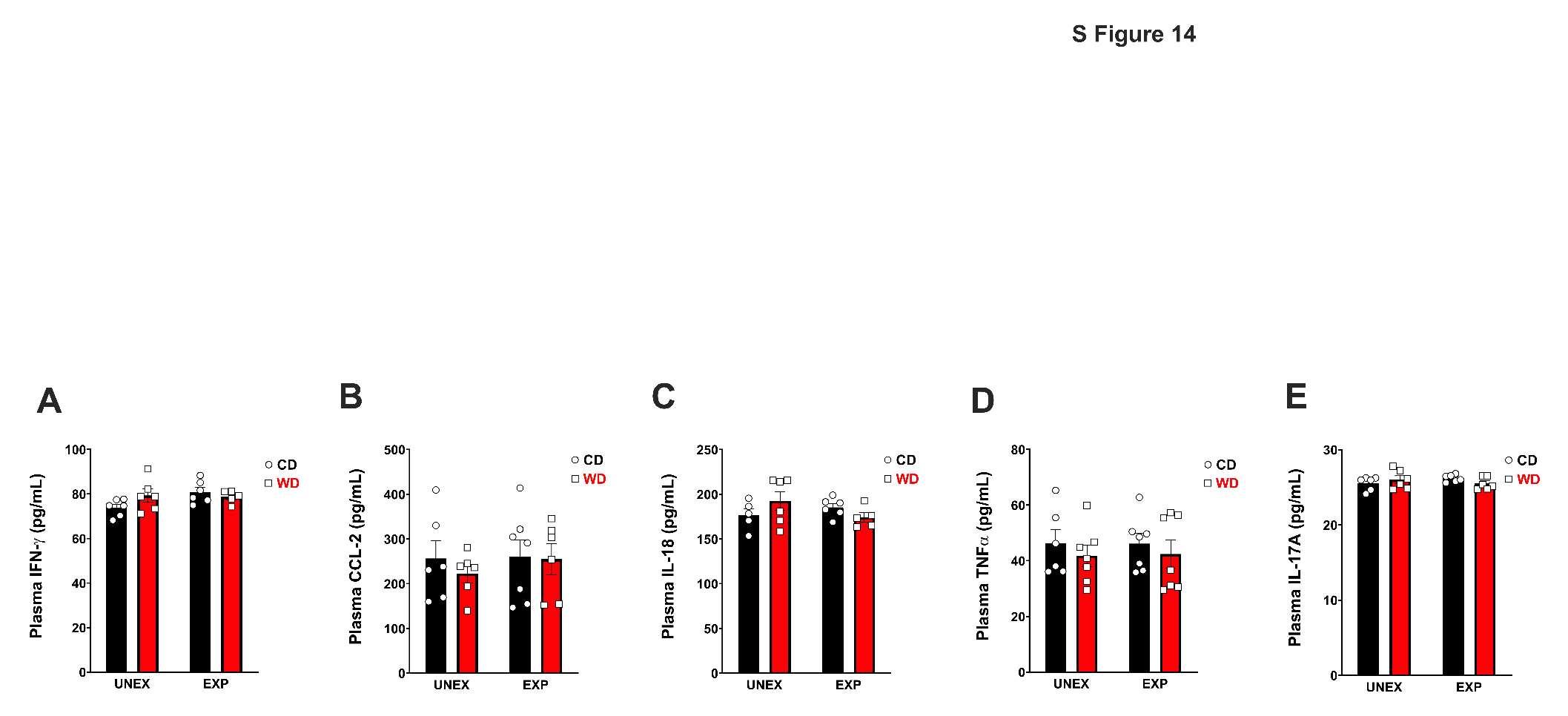

***Supplemental Figure 15.* Evaluation of genes relevant to metabolism and inflammation.** Gene expression of hippocampal levels of FKBP4, NR3C1, NR3C2, HSD11b1, IL6, NFκB, DUSP1, and SV2A were evaluated, and no significant effects were observed by the interventions. For a complete table of statistical analysis, see Supplemental Table 15.

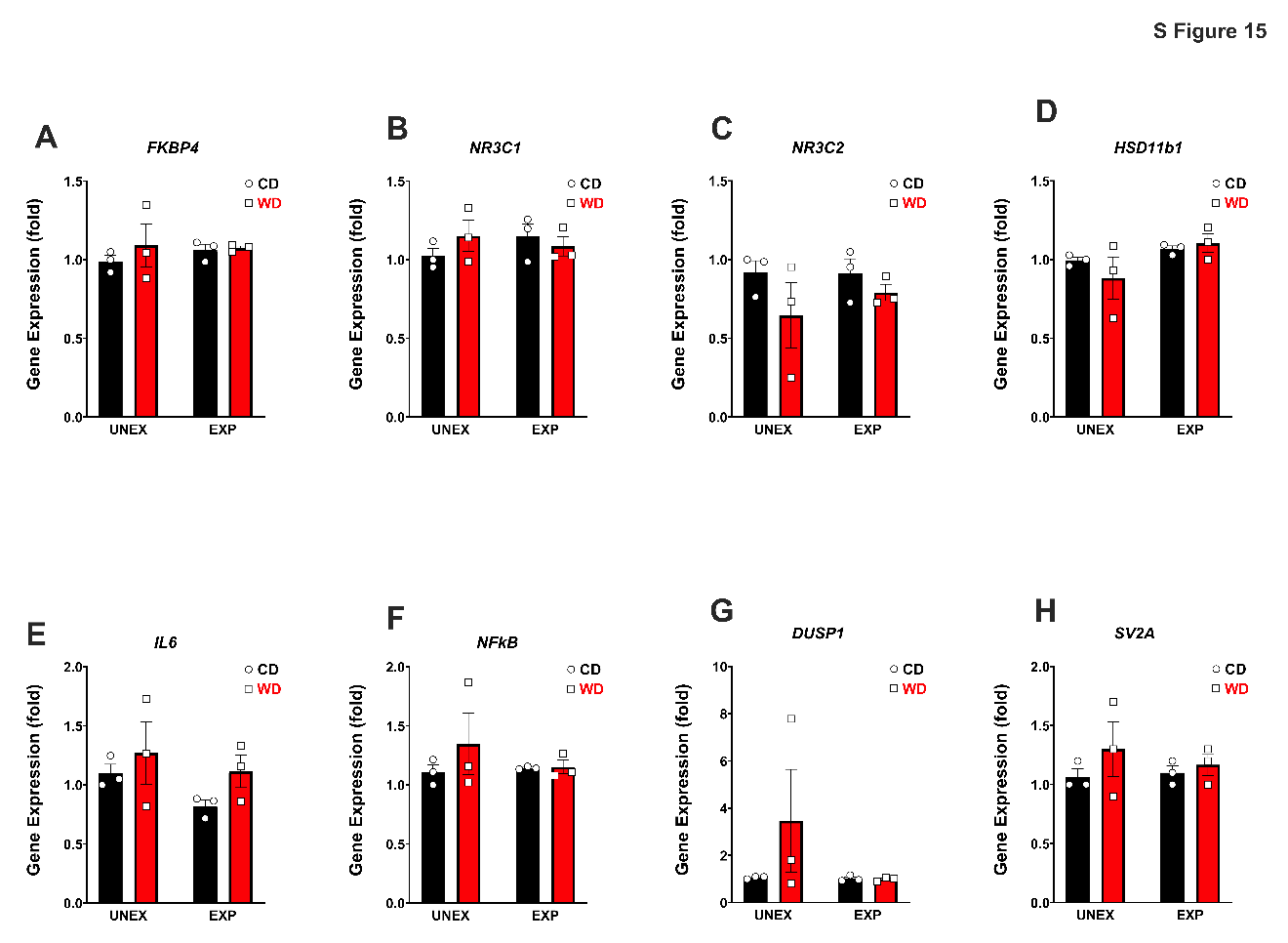

***Supplemental Figure 16,17*. Ensembl Genome of rat Fkbp5.** The complete sequence of rat Fkbp5 gene, including transcription length (3387 bp, 456 aa), chromosomal location [Chr20:7,976,713-8,019,020(-)], and mapping of twenty-three (23) regulatory regions.

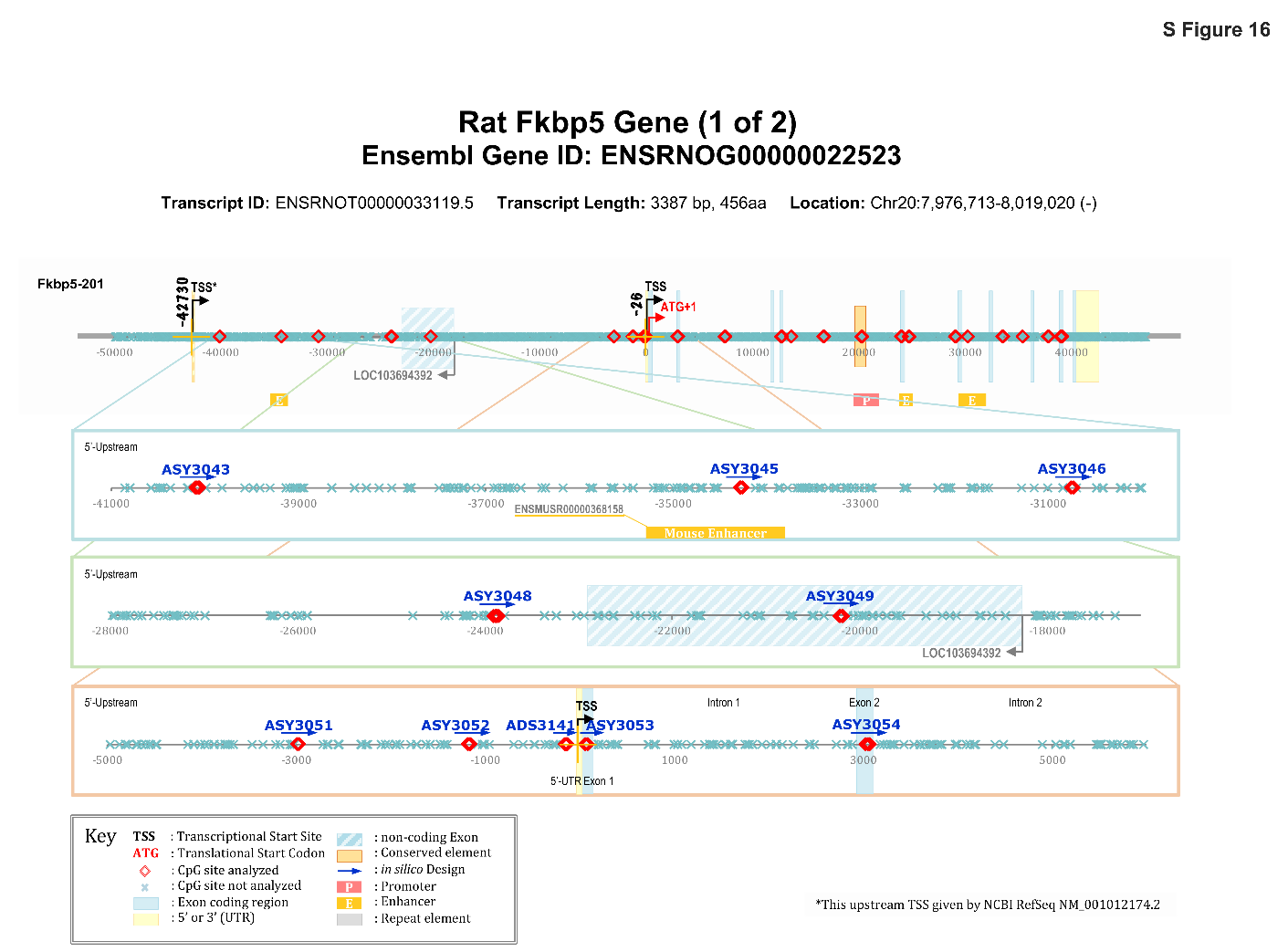

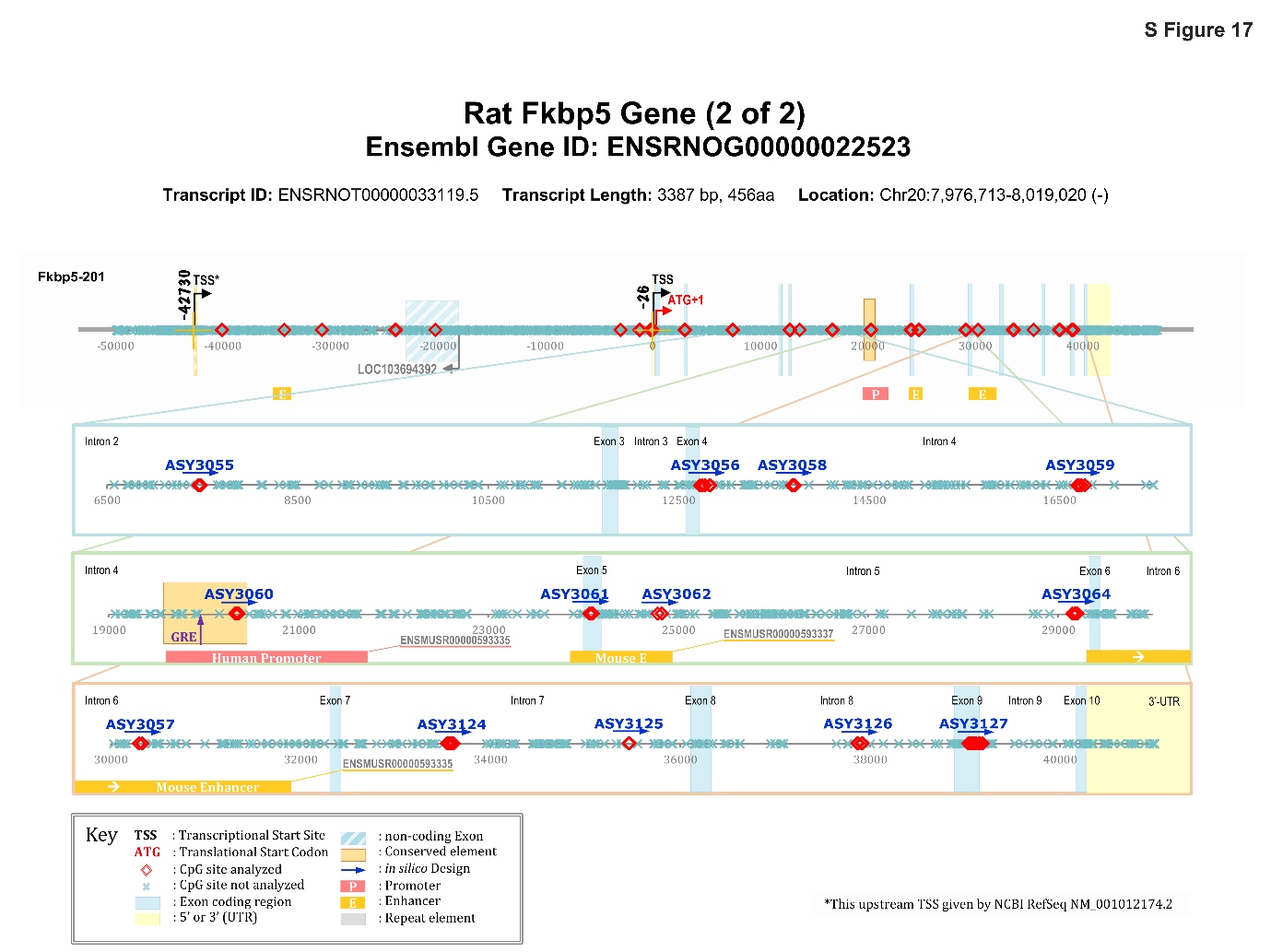

***Supplemental Figure 18*. Complete list of regulatory regions of Rat Fkbp5.** List of the twenty-three (23) optimized Rat Fkpb5 regulatory regions with assay location, coordinates from Analytical and Translational Genomics (ATG) and Transcriptional Start sites (TSS), Rnor_6.0 (-) Chromosomal regions, and number of CpG sites.

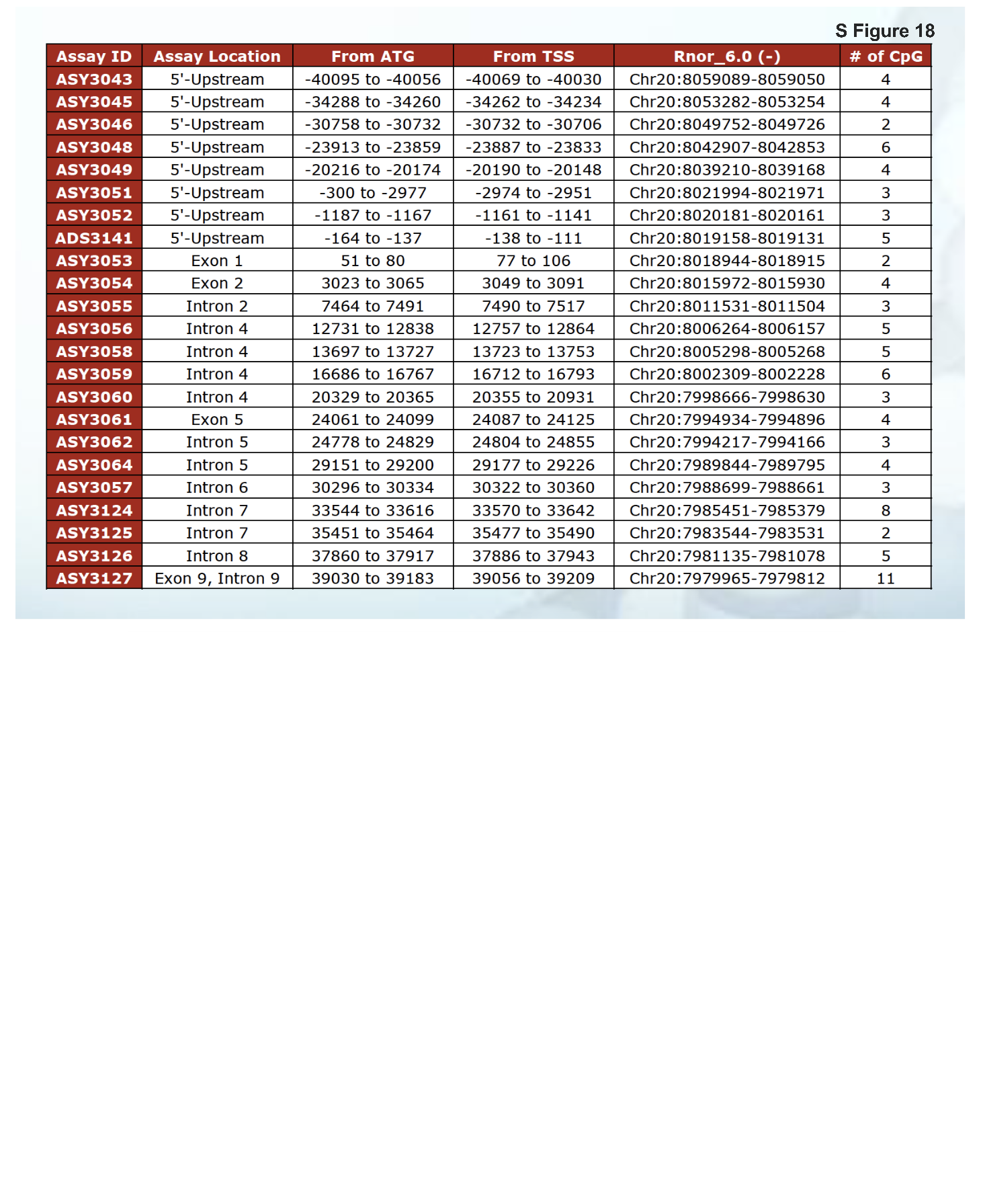

.

***Supplemental Figure 19.* Heatmap and analysis of Fkbp5 methylation signatures per gene region. (A)** Global percentages of Fkbp5 methylation per region **(B)** Breakdown of significance per region. Methylation in the 5-Upstream was significantly modified by PSS (*p* = 0.0001) and the interaction of WD x PPS (*p* = 0.009). Intron 4 was significantly modified by PSS (*p* = 0.003). Intron 6 was modified by WD x PSS (*p* = 0.032). Intron 7 was modified by the diet (*p* = 0.032); interestingly Intron 8 was modified by WD (*p* = 0.024), PSS (*p* = 0.017), and WD x PSS (0.024). The summary of statistical values of multiple comparisons in FKBP5 by regions can be found in Supplemental Table 16A-C and 17.

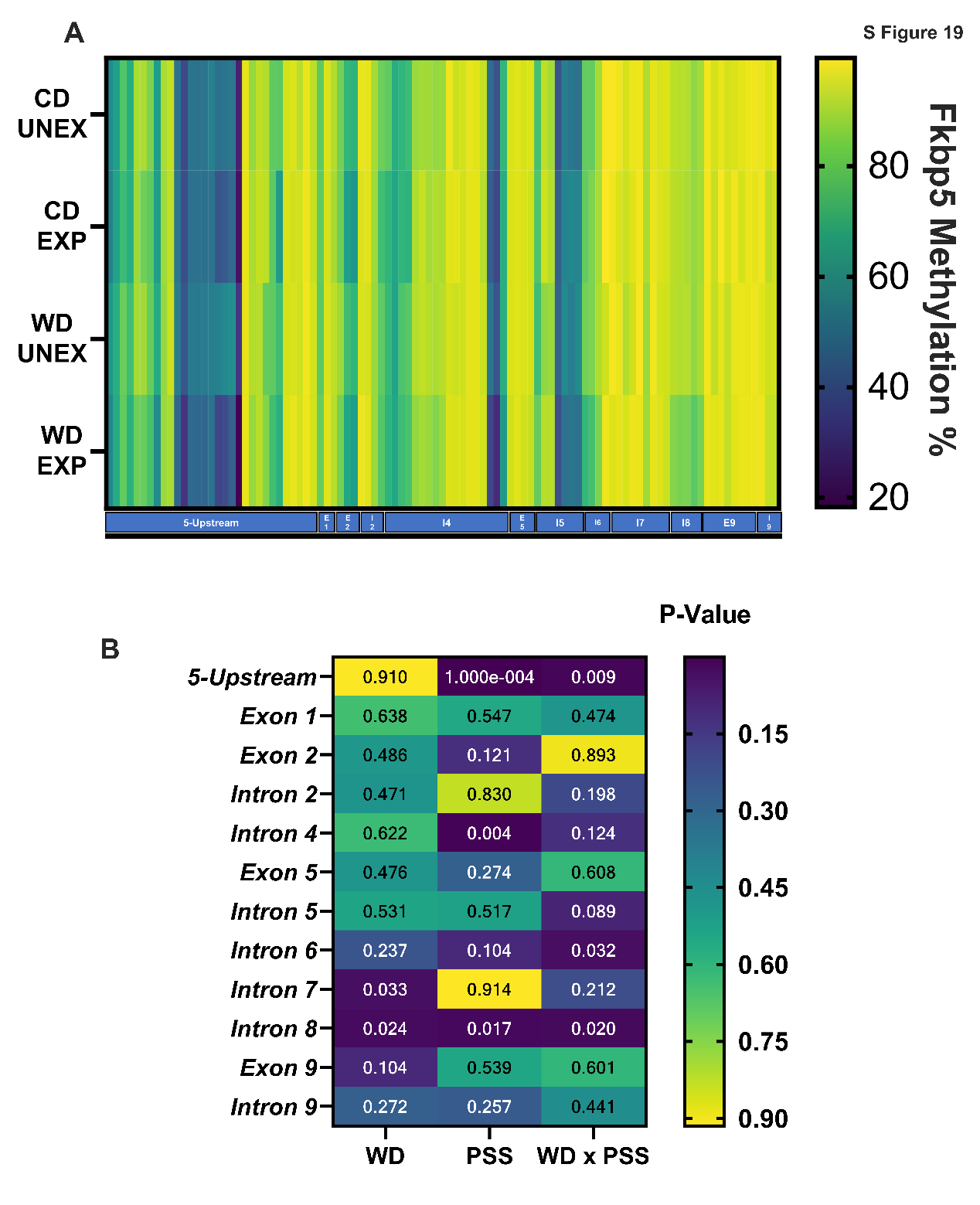

***Supplemental Figure 20.* Evaluation of Viability and the half maximal inhibitory concentration (IC_50_) of HMC3 cells treated with PA and Hydrocortisone. (A)** Evaluation viability and IC**_50_** of HCM3 cells treated with increasing concentrations of PA (0-800 µM). No significant difference in the percent of viability between cells treated with PA or PA + Hydrocortisone. IC_50_ for PA was determined to be 33.52 µM, and PA + Hydrocortisone was 50.84 µM. **(B)** HMC3 cells treated with increasing concentrations of Hydrocortisone (0-1000 nM) showed a 50% of growth inhibition (IC_50_) at 119.3 nM.

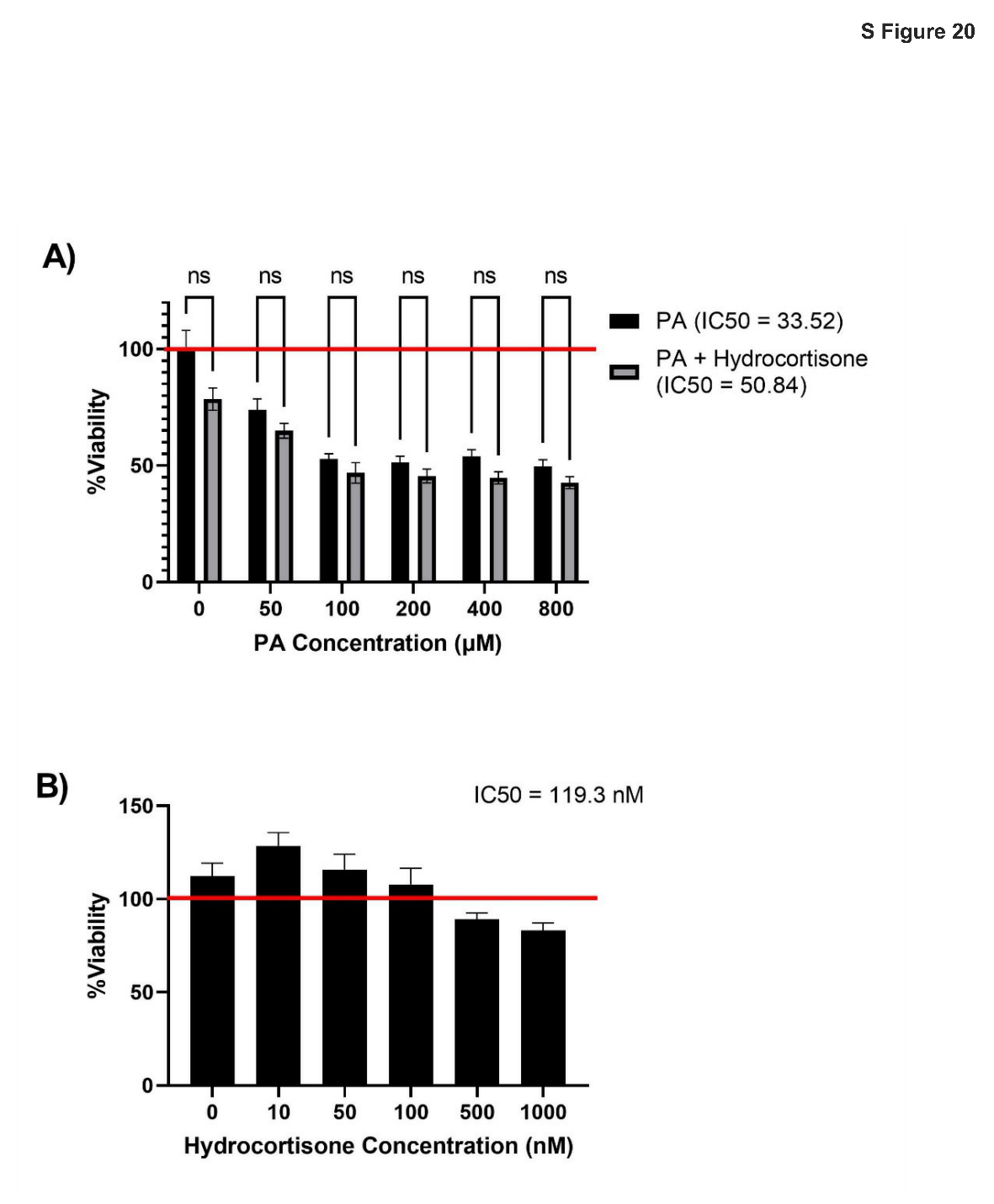

***Supplemental Figure 21.* Confocal microscopy images of HMC3 cells treated with PA and Hydrocortisone** Representative images of cells treated with either vehicle, Hydrocortisone at 100 nM, PA at 50 µM, alone or pretreated with PA (50 µM) for 24 h followed by Hydrocortisone (100 nM). The image shows individual markers CD68 (yellow) and nuclear DAPI staining (pink) and the merged images. Changes in microglia morphology are noticeable with the combination of PA + Hydrocortisone.

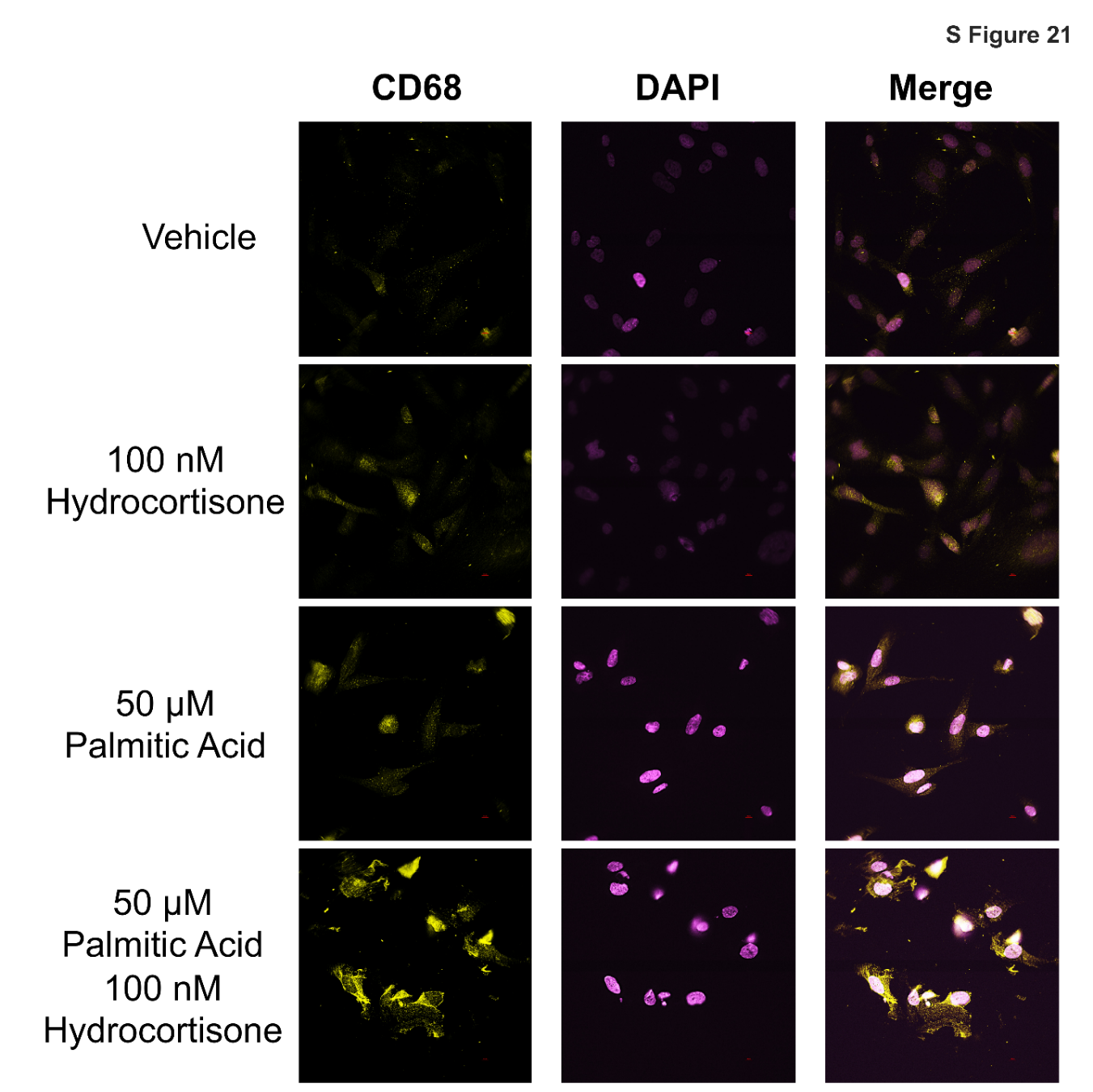

***Supplemental Figure 22.* Confocal microscopy images of HMC3 cells treated with PA and hydrocortisone.** Representative images of cells treated with either vehicle, Hydrocortisone at 100 nM, PA at 50 µM, alone or pretreated with PA (50 µM) for 24 h followed by Hydrocortisone (100 nM). The image shows individual markers CD68 (red) and nuclear DAPI staining (blue) and the merged images. Changes in microglia morphology are noticeable with the combination of PA + hydrocortisone.

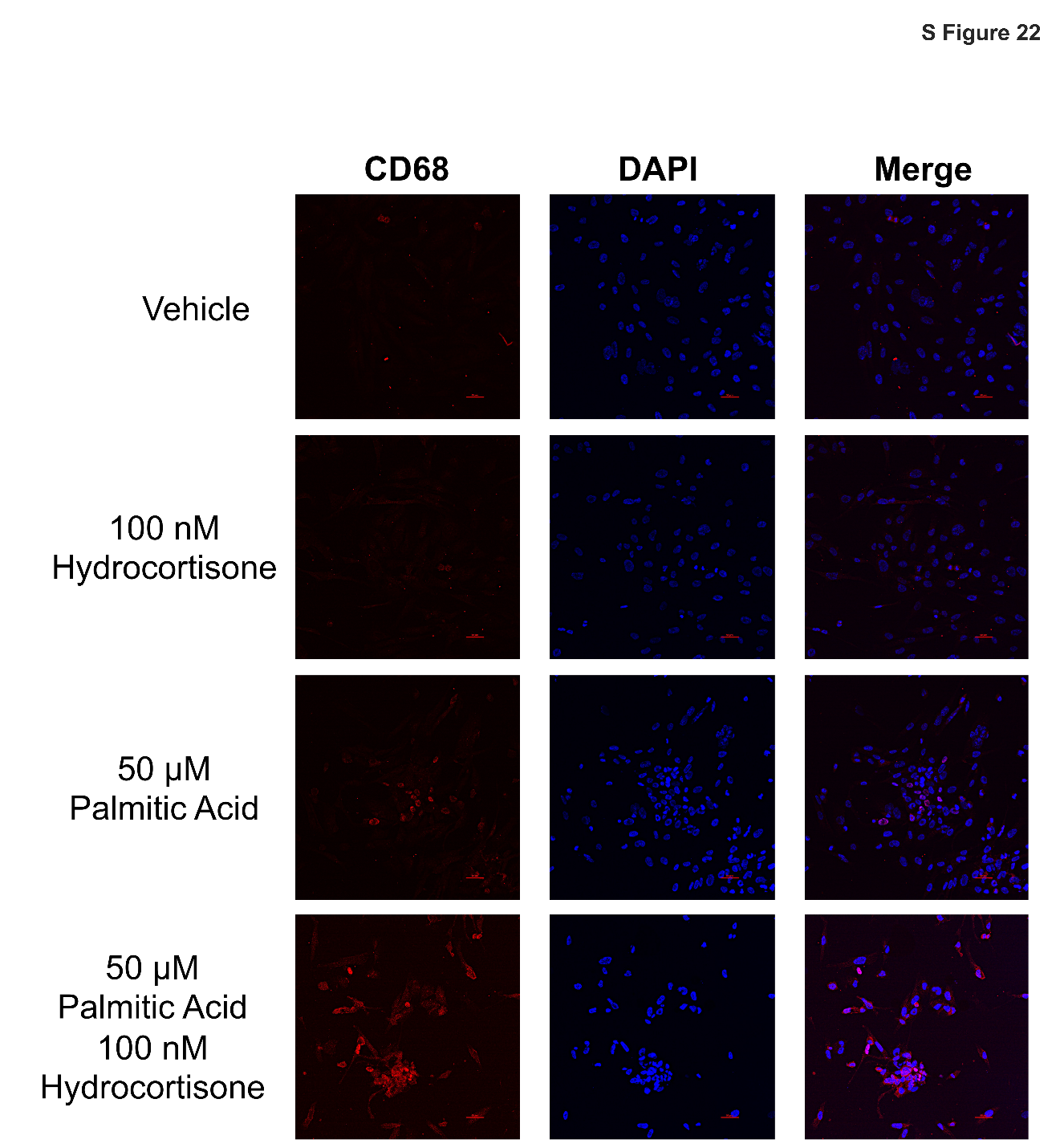

***Supplemental Figure 23.* Evaluation of IL-6 gene and protein expression in HMC3 cells treated with PA and Hydrocortisone, and HCM3 with Silenced FKBP5. (A)** IL-6 Gene expression was significantly affected by the interaction of PA + CORT (*p* < 0.0001) and by PA treatment alone (*p* = 0.0324). **(B)** IL-6 protein concentration was also significantly affected by the interaction of PA + CORT (*p* < 0.014) and by CORT treatment alone (*p* = 0.0022). **(C)** IL-6 Gene expression was significantly modified by SiRNA, treatment, and SiRNA x Treatment, all with (*p* < 0.0001). IL-6 protein concentration was also significantly modified by SiRNA (*p* < 0.0001), treatment (*p* < 0.0001), and SiRNA x Treatment (*p* = 0.0022). A summary of the statistical values of multiple comparisons is in Supplemental Tables 18-20.

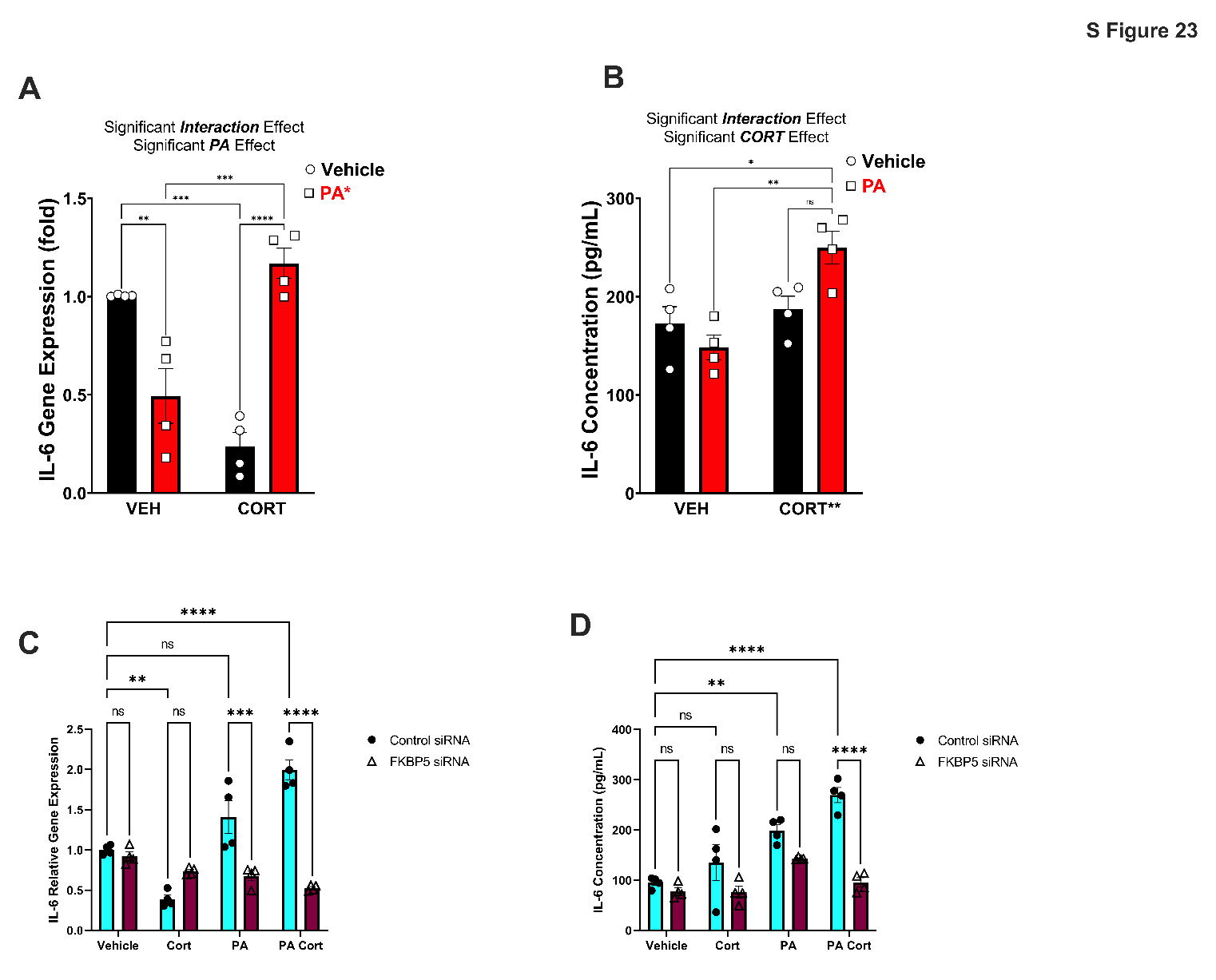

***Supplemental Figure 24.* FKBP5 Gene expression of HCM3 cells treated with CORT and FKBP5 SiRNA. (A)** FKBP5 Gene expression in HCM3 cells treated with increasing concentrations of CORT (0-1000nm) showed an interesting rise and fall pattern, suggesting a dampening of the expression at high CORT concentrations. **(B)** FKBP5 Gene expression with Control siRNA and FKBP5 SiRNA show a significant decrease in the silenced samples (*t* = 3.559, *p* = 0074). A summary of the statistical values of multiple comparisons is in Supplemental Tables 21, and 22.

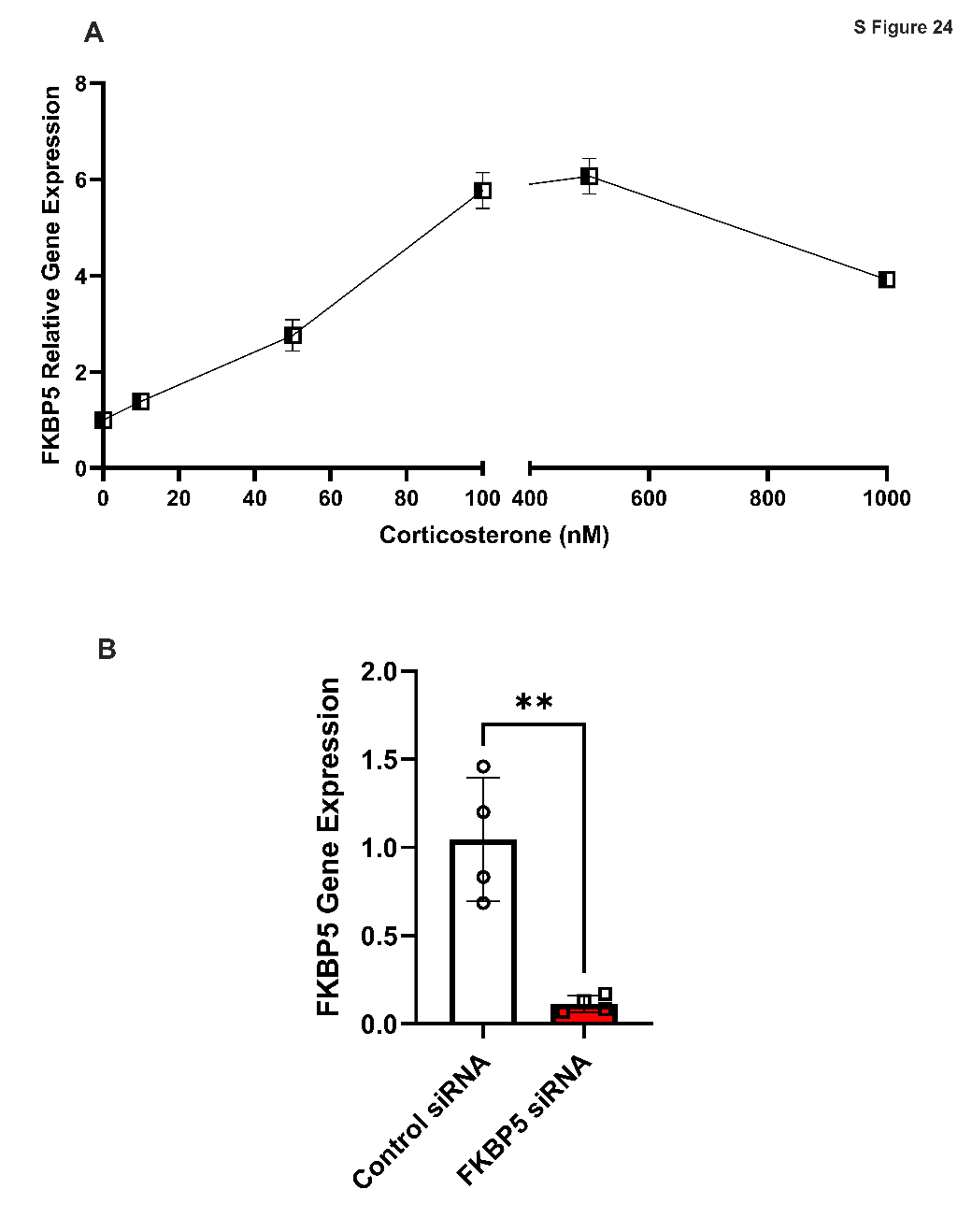

**ABBREVIATIONS**

| AD | Axial Diffusivity |
| --- | --- |
| ADAM17 | TNF-α-converting enzyme |
| ADOL | Adolescence |
| ADUL | Adulthood |
| AI | Anxiety index |
| ANOVA | Analysis of variance |
| ASR | Acoustic startle reflex |
| ASR | Acoustic Startle Reflex |
| BA | Background anxiety |
| BED | Binge eating disorders |
| BEH | Behavior |
| BGM | Brain-gut-microbiome |
| BL | Anterior basolateral amygda |
| BLA | Basolateral amygdala |
| BM | Posterior basomedial amygdala |
| BMI | Body mass index |
| BMI | Body mass index |
| BV | Brain volume |
| CA | Closed arm |
| CCL2 | Chemokine (C-C motif) ligand 2 |
| CD | Control diet |
| CD EXP | Control diet exposed |
| CD UNEX | Control diet unexposed |
| CD68 | CD68 (Cluster of Differentiation 68) |
| CDE | Control-diet exposed |
| cDNA | Complementary deoxyribonucleic acid |
| CDU | Control diet unexposed |
| CHOW | Standard chow diet |
| CORT | Corticosterone |
| CORT | Hydrocortisone |
| CRF | Corticotropin releasing factor |
| CRH | Corticotropin-releasing hormone |
| CRHR1 | Corticotropin-releasing hormone receptor 1 |
| CS | Conditioned stimulus |
| CS + US | Conditioned stimulus + unconditioned stimulus |
| CXCL-1 | Chemokine (C-X-C motif) ligand 1 |
| dB | Decibel |
| dCA1 | Dorsal CA1 |
| dCA3 | Dorsal CA3 |
| dDG | Dorsal Dentate Gyrus |
| DIO | Diet-induced obesity |
| dSb | dorsal subiculum |
| DTI | Diffusion tensor imaging |
| EC | entorhinal cortex |
| EcrhC | ectorhinal cortex |
| ED | Eating Disorders |
| ELISA | Enzyme-linked immunosorbent assay |
| ELS | Early life stress |
| ELS | Early Life Stress |
| ELT | Early life trauma |
| EPM | Elevated Plus Maze |
| EXP | Exposed |
| FA | Fractional anisotropy |
| FCM | Fecal corticosterone metabolites |
| FCM | Fecal corticosterone metabolites |
| FDR | False discovery rate |
| FKBP51 | FK506-binding protein 51 |
| FPS | Fear Potentiated Startle |
| FST | Forced swim test |
| GAPDH | Glyceraldehyde 3-phosphate dehydrogenase |
| GE | Gene expression |
| GFAP | Glial fibrillary acidic protein |
| GM-CSF | Granulocyte-macrophage colony-stimulating factor |
| GPx | Glutathione peroxidase |
| GR | Glucocorticoid receptor |
| GSH | Glutathione |
| GSR | Glutathione reductase |
| h | hour |
| H_2_O_2_ | Hydrogen peroxide |
| HFD | High-fat diet |
| HMC3 | Human Microglial cell line |
| HPA | Hypothalamic-pituitary adrenal |
| HPA | The hypothalamic–pituitary–adrenal axis |
| HPC | Hippocampus |
| HPC CA | Cornu Ammonis |
| HPG | Hypothalamic–pituitary–gonadal axis |
| HPO | Hipocampus-Pitutary-ovaries |
| IACUC | Institutional animal care and use committee |
| IC_50_ | the half maximal inhibitory concentration |
| ICVF | Intracellular Volume Fraction |
| IHC | Immunohistochemistry |
| IL | Interleukin |
| IL-10 | Interleukin-10 |
| IL-12p70 | Interleukin-12p70 |
| IL-17a | Interlukin-17a |
| IL-18 | Interleukin-18 |
| IL-1α | Interleukin-1 alpha |
| IL-1β | Interleukin-1 beta |
| IL-33 | Interleukin-33 |
| IL-6 | Interleukin 6 |
| INF-γ | Interferon gamma |
| IPGTT | Intraperitoneal glucose tolerance test |
| ISOVF | Isometric Volume Fraction |
| ITI | Inter trial interval |
| LJ | Jacobian Integration Index |
| *Ls NK4A136* | Lacnospiracae NK4A136 |
| MD | Medial Diffusivity |
| mPFC | Medial prefrontal cortex |
| MR | Mineralocorticoid receptors |
| MRI | Magnetic resonance imaging |
| mRNA | Messenger ribonucleic acid |
| na | not applicable |
| nd | not detected |
| ND | Not determined |
| NHP | Non-human primate |
| NIR | Near infrared-backlit |
| NODDI | Neurite Orientation dispersion and density imaging |
| ns | not significant |
| OA | Open arm |
| ODI | Orientation Dispersion Index |
| OFT | Open field test |
| PA | Palmitic Acid |
| PBG | Postprandial blood glucose |
| PBS | Phosphate buffered saline |
| PFA | Paraformaldehyde |
| PFC | Prefrontal cortex |
| PHc | Palatable Hypercaloric Foods |
| PND | Post-natal day |
| POS | Predator odor stress |
| PPI | Pre-pulse inhibition |
| PS | Psychogenic stress |
| PSS | Psychosocial Stress |
| PTSD | Post-traumatic stress disorder |
| RD | Radial Diffusivity |
| ROI | Region of interest |
| ROS | Reactive oxygen species |
| RT-qPCR | Real time quantitative polymerase chain reaction |
| SYM | Social Y Maze |
| T2WI | T2-weighted |
| TACE | TNF-α-converting enzyme |
| TE | Echo time |
| TG | Triglycerides |
| TNF | Tumor necrosis factor |
| TR | Repetition time |
| UNEXP | Unexposed |
| US | Unconditioned stimulus |
| vCA1 | Ventral CA1 |
| vCA3 | Ventral CA3 |
| vDG | Ventral Dentate Gyrus |
| VML | ventromedial lateral amygdaloid nucleus |
| vSb | ventral subiculum |
| VTA | Ventral tegmental area |
| WB | Western blot |
| WD | Western Diet |
| WD EXP | Western diet exposed |
| WD UNEX | Western diet unexposed |
| WDE | Western diet exposed |
| WDU | Western diet unexposed |
| 3D | Three dimensional |

**Supplemental TABLES**

***Supplemental Table 1*.** Macronutrient composition of the custom purified diets. Detailed composition of the matched low-fat purified control diet (CD, 5-gm% fat, product *Cat. No F7463*) and Western-like high-saturated fat diet (WD, 20-gm% fat, product *Cat. No. F7462*).

| **Macronutrient (main source)** | **CD** | **WD** |
| --- | --- | --- |
| % kcal Carbohydrates (corn starch) | 63.5 | 49.2 |
| % kcal Protein (casein) | 17.7 | 15.5 |
| % kcal Fat (milk fat) | 5.2 | 21 |
| % kcal Fiber | 4.8 | 4.8 |
| Total Caloric Profile (kcal/gram) | 3.72 | 4.57 |
| **Fatty Acid Class** | **g/kg Diet** | **g/kg Diet** |
| Butyric (C4:0) | 0.8 | 4.8 |
| Caproic (6:0) | 0.6 | 3.4 |
| Caprylic (C8:0) | 0.3 | 2.0 |
| Capric (C10:0) | 0.8 | 4.7 |
| Lauric (C12:0) | 0.9 | 5.6 |
| Myristic (C14:0) | 3.1 | 18.8 |
| Pentadecanoic (15:0) | 0.3 | 2.0 |
| Palmitic (C16:0) | 12.1 | 60 |
| Heptadecanoic (17:0) | 0.2 | 1.0 |
| Stearic (C18:0) | 3.7 | 20.8 |
| Arachidic (20:0) | 0.1 | 0.3 |
| Myristoleic (C14:1) | 0.3 | 1.6 |
| Palmitoleic (C16:1) | 0.5 | 2.9 |
| Oleic (C18:1) | 11.4 | 38.8 |
| Linoleic (C18:2) | 11.3 | 9.9 |
| Alpha Linolenic (C18:3) | 0.66 | 3.0 |
| Homogamma Linolenic (20:3) | <0.07 | 0.2 |
| Arachidonic (n6) 20:4 | <0.07 | 0.3 |
| EPA (20:5 n-3) | <0.07 | <0.14 |
| DHA (22:6 n-3) | <0.07 | <0.14 |
| **Total per Fatty Acid Class** | **g/kg Diet** | **g/kg Diet** |
| Saturated (SFA) | 21.1 | 125 |
| Monosaturated (MUFA) | 14.1 | 60.2 |
| Polyunsaturated (PUFA) | 12.0 | 12.8 |

***Supplemental Table 2.*** Summary of statistical analysis of growth chart, food consumption, fasting glucose, and cardiovascular function in Supplemental Figures 1-3.

|  | **Caloric intake** (% change before and after PSS) | **Body Weight**  (% change before and after PSS) | **Fasting Blood Glucose** |
| --- | --- | --- | --- |
| Source of Variation | F Statistic, post-hoc | F Statistic, post-hoc | F Statistic, post-hoc |
| Stress: Diet | F(1, 26) = 1.67, *p =* 0.207 | F(1, 49) = 0.0993, *p =* 0.754 | F(1, 52) = 0.0005, *p=*0.98 |
| **STRESS** | **F(1, 26) = 10.5, *p =* 0.003** | **F(1, 49) = 14.27, *p =* 0.0004** | **F(1, 52) = 12.06, *p =* 0.001** |
| DIET | F(1, 26) = 0.019, *p =* 0.88 | **F(1, 49) = 23.08, p < 0.0001** | F(1, 52) = 0.53, *p =* 0.47 |
| CDU vs WDU | *p =* 0.8597 | ***p =* 0.0125** | *p =* 0.959 |
| CDU vs CDE | ***p =* 0.023** | *p =* 0.1028 | *p =* 0.0843 |
| CDU vs WDE | *p =* 0.176 | **p < 0.0001** | *p =* 0.2377 |
| WDU vs CDE | *p =* 0.0858 | *p =* 0.8836 | ***p =* 0.0196** |
| WDU vs WDE | *p =* 0.4923 | ***p =* 0.0192** | *p =* 0.0741 |
| CDE vs WDE | *p =* 0.7188 | ***p =* 0.0041** | *p =* 0.9512 |

***Supplemental Table 3.*** Summary of statistical analysis of cardiovascular function

|  | **Blood Pressure**  Diastolic | **Heart Rate**  Beats per minute |
| --- | --- | --- |
| Source of Variation | F Statistic, post-hoc | F Statistic, post-hoc |
| Stress: Diet | F(1, 24) = 1.325, *p =* 0.2611 | F(1, 23) = 0.5281, *p =* 0.4747 |
| STRESS | F(1, 24) = 3.6e-7, *p =* 0.9995 | F(1, 23) = 1.240, *p =* 0.2771 |
| **DIET** | **F(1, 24) = 6.263, *p =* 0.0195** | **F(1, 23) = 7.978, *p =* 0.0096** |
| **UNEX** | ***p =* 0.0333** | ***p =* 0.043** |
| EXP | *p =* 0.5726 | *p =* 0.2667 |

***Supplementary Table 4*.** Summary of behavioral measurements in Supplementary Figure 4

|  | **FPS**  Foot shock reactivity | **FPS**  (% change from learning session) | **EPM**  (% change) |
| --- | --- | --- | --- |
| Source of Variation | F Statistic, post-hoc | F Statistic, post-hoc | F Statistic, post-hoc |
| Stress: Diet | F(1, 52) = 0.2, *p =* 0.65 | F(1, 49) = 0.09, *p =* 0.75 | F(1, 53) = 0.34, *p =* 0.56 |
| STRESS | F(1, 52) = 1.35, *p =* 0.25 | **F(1, 49) = 14.3, *p =* 0.0004** | F(1, 53) = 1.6, *p =* 0.21 |
| **DIET** | **F(1, 52) = 12.9, *p =* 0.0007** | **F(1, 49) = 23.1, p < 0.0001** | **F(1, 53) = 11.3, *p =* 0.0014** |
| CDU vs WDU | ***p =* 0.0302** | **F(1, 34) = 5.8, *p =* 0.0216** | *p =* 0.2934 |
| CDU vs CDE | *p =* 0.6808 | F(1, 34) = 1.8, *p =* 0.188 | *p =* 0.9978 |
| CDU vs WDE | ***p =* 0.009** | F(1, 34) = 0.67, *p =* 0.416 | ***p =* 0.012** |
| WDU vs CDE | *p =* 0.3054 | *p =* 0.125 | *p =* 0.5924 |
| WDU vs WDE | *p =* 0.9555 | ***p =* 0.0473** | *p =* 0.7043 |
| CDE vs WDE | *p =* 0.128 | *p =* 0.3822 | ***p =* 0.0392** |

***Supplementary Table 5***. Summary of behavioral measurements in Supplementary Figure 4

|  | **EPM**  Nose in head dipping zone | **EPM**  Stretch attend |
| --- | --- | --- |
| Source of Variation | F Statistic, post-hoc | F Statistic, post-hoc |
| Stress: Diet | F(1, 53) = 3.441, *p =* 0.0691 | F(1, 53) = 0.02393, *p =* 0.8776 |
| STRESS | F(1, 53) = 0.08980, *p =* 0.7656 | F(1, 53) = 1.017, *p =* 0.3179 |
| **DIET** | **F(1, 53) = 15.44, *p =* 0.0002** | **F(1, 53) = 21.19, *p* < 0.0001** |
| **UNEX** | ***p =* 0.2811** | ***p =* 0.0031** |
| **EXP** | ***p =* 0.0003** | ***p =* 0.005** |

***Supplementary Table 6*.** Summary of supplemental behavioral measurements in Supplementary Figure 4

|  | **SYM**  Total distance | **SYM**  (Sociability Index) | **SYM**  (Latency to first) |
| --- | --- | --- | --- |
| Source of Variation | F Statistic, post-hoc | F Statistic, post-hoc | F Statistic, post-hoc |
| Stress: Diet | F(1, 45) = 0.75, *p =* 0.39 | F(1, 50) = 0.04, *p =* 0.83 | F(1, 43) = 1.7, *p =* 0.19 |
| STRESS | **F(1, 45) = 4.8, *p =* 0.03** | **F(1, 50) = 8.2, *p =* 0.006** | F(1, 43) = 1.0, *p =* 0.32 |
| DIET | F(1, 45) = 0.72, *p =* 0.4 | F(1, 50) = 1.8, *p =* 0.181 | F(1, 43) = 0.12, *p =* 0.72 |
| CD | *p =* 0.0834 | *p =* 0.0739 | *p =* 0.9676 |
| WD | *p =* 0.5506 | *p =* 0.1181 | *p =* 0.1948 |

***Supplementary Table 7.*** Statistical Information of supplemental microglia measurements

|  | **Density** | **Mean Radius** | **Circularity** |
| --- | --- | --- | --- |
| Source of Variation | F Statistic, post-hoc | F Statistic, post-hoc | F Statistic, post-hoc |
| Stress: Diet | F(1, 12) = 0.40, *p =* 0.53 | F(1, 12) = 0.3, *p =* 0.5 | F(1, 11) = 0.002, *p =* 0.96 |
| STRESS | F(1, 12) = 1.7, *p =* 0.21 | F(1, 12) = 1.2, *p =* 0.27 | **F(1, 11) = 6.7, *p =* 0.025** |
| DIET | **F(1, 12) = 6.1, *p =* 0.028** | **F(1, 12) = 5.6, *p =* 0.03** | F(1, 11) = 3.6, *p =* 0.08 |
| CDU vs WDU |  |  | *p =*0.5967 |
| CDU vs CDE |  |  | *p =*0.3531 |
| CDU vs WDE |  |  | ***p =*0.045** |
| WDU vs CDE |  |  | *p =*0.9582 |
| WDU vs WDE |  |  | *p =*0.2662 |
| CDE vs WDE |  |  | *p =*0.4989 |

***Supplemental Table 8*.** Summary of Neuroimaging parameter descriptors

| **Parameter** | **Abbreviation** | **Description** |
| --- | --- | --- |
| **Fractional Anisotropy** | **FA** | Measures the degree of anisotropy of water molecules. Without obstacles, water molecules diffuse freely in any direction, a pattern that may be changed by the presence of macromolecules, cell membranes, etc. If water molecules are conducted within a tube (or an axon), diffusion only occurs along the axis of this tube. Therefore, diffusion is not isotropic, but anisotropic. The degree of anisotropy can be measured, and allows to infer alterations in the axonal diameter, fiber density or myelin structure. Reductions in FA have been linked to reduced neurite density, increased dispersion of orientation suggests decreased white matter integrity. |
| **Axial Diffusivity** | **AD** | Coefficient of diffusion across the long axis of the ellipsoid, typically lying along the axon. Decreased AD indicates decreased of white matter integrity. |
| **Medial Diffusivity** | **MD** | Describes the rotationally invariant magnitude of water diffusion within brain tissue. Differences in MD could reflect variations within the intra- and extracellular space, and/or index global increases in CSF. MD itself is a rather non-specific, albeit sensitive, measure that can be affected by any disease process that affects the barriers that restrict the motion of water, such as cell membranes. |
| **Radial Diffusivity** | **RD** | Coefficient of diffusion perpendicular to the long axis. Increased RD is an indicator of decreased white matter integrity. |
| **Jacobian Integration Index** | **LJ** | Longitudinal registration-based method and a type of tensor-based morphometry. The output of the Jacobian integration method is a percent change in volume. |
| **Intracellular Volume Fraction** | **ICVF** | Compartmentalizes non-Gaussian water diffusion into geometric space encompassing hindered anisotropic components. Corresponds to intra-neurite water (of axons and dendrites). |
| **Isometric Volume Fraction** | **ISOVF** | Compartmentalizes non-Gaussian water diffusion into a geometric space encompassing isotropic (or free) components. Corresponds to free water/CSF. |
| **Orientation Dispersion Index** | **ODI** | Quantifies the angular variation of neurite orientation (ranging from 0 for perfectly coherently oriented structures) to 1 (for isotropic structures). Typically high in GM, low in WM). |

***Supplemental Table 9A*.** Statistical Information of hippocampal structural changes by neuroimaging parameters detected via DTI.

|  | **FA** | **MD** | **AD** | **RD** |
| --- | --- | --- | --- | --- |
| Source of Variation | F Statistic, post-hoc | F Statistic, post-hoc | F Statistic, post-hoc | F Statistic, posthoc |
| **Region** | **F(10, 242) = 19.4, p < 0.0001** | **F(10, 242) = 12.8, p < 0.0001** | **F(10, 242) = 12.8, p < 0.0001** | **F(10, 242) = 12.8, p < 0.0001** |
| Diet | **F(1, 242) = 25.1, p < 0.0001** | **F(1, 242) = 4.7, *p =* 0.03** | **F(1, 242) = 4.4, *p =* 0.035** | **F(1, 242) = 4.8, *p =* 0.028** |
| Stress | **F(1, 242) = 9.6, *p =* 0.002** | **F(1, 242) = 4.7, *p =* 0.029** | **F(1, 242) = 4.5, *p =* 0.03** | **F(1, 242) = 4.7, *p =* 0.029** |
| Region x Diet | F(10, 242) = 0.41, *p =* 0.94 | F(10, 242) = 0.2, *p =* 0.99 | F(10, 242) = 0.2, *p =* 0.9 | F(10, 242) = 0.2, *p =* 0.98 |
| Region x Stress | F(10, 242) = 0.5, *p =* 0.88 | F(10, 242) = 0.1, *p =* 0.99 | F(10, 242) = 0.19, *p =* 0.9 | F(10, 242) = 0.12, *p =* 0.9 |
| Diet x Stress | **F(1, 242) = 5.3, *p =* 0.0219** | F(1, 242) = 2.4, *p =* 0.11 | F(1, 242) = 3.2, *p =* 0.07 | F(1, 242) = 2.0 *p =* 0.15 |
| Region x Diet x Stress | F(10, 242) = 0.75, *p =* 0.67 | F(10, 242) = 0.4, *p =* 0.94 | F(10, 242) = 0.3, *p =* 0.9 | F(10, 242) = 0.4, *p =* 0.9 |

***Supplemental Table 9B*.** Statistical Information of hippocampal structural changes by neuroimaging parameters detected via NODDI.

|  | **ODI** | **ICVF** | **ISOVF** | **LJ** |
| --- | --- | --- | --- | --- |
| Source of Variation | F Statistic, post-hoc | F Statistic, post-hoc | F Statistic, post-hoc | F Statistic, post-hoc |
| **Region** | **F(10, 242)=22.2, p < 0.0001** | **F(10, 231)=7.4, p < 0.0001** | **F(10, 244)=17.05, p < 0.0001** | **F(10, 242) = 32.9, p < 0.0001** |
| Diet | **F(1, 242) 28.5, p < 0.0001** | **F(1, 231) = 44.5, p < 0.0001** | F(1, 244) = 2.3, *p =* 0.12 | **F(1, 242) = 9.5, *p =* 0.0023** |
| Stress | F(1, 242) = 2.8, *p =* 0.09 | F(1, 231) = 1.3, *p =* 0.24 | F(1, 244) = 1.7, *p =*0.19 | F(1, 242) = 0.08, *p =* 0.7 |
| Region x Diet | F(10, 242) = 0.3, *p =* 0.9 | F(10, 231) =0.09, *p=*0.99 | F(10, 244) = 0.38, *p =* 0.9 | F(10, 242) = 0.3, *p =* 0.9 |
| Region x Stress | F(10, 242) = 0.2, *p =* 0.9 | F(10, 231)=0.05, p > 0.99 | F(10, 244) = 0.06, p > 0.9 | F(10, 242) = 0.6, *p =* 0.8 |
| Diet x Stress | **F(1, 242) = 6.2, p = 0.013** | **F(1, 231)= 7.8, p = 0.0057** | F(1, 244) = 0.05, *p =* 0.8 | **F(1, 242) = 4.9, *p =* 0.02** |
| Region x Diet x Stress | F(10, 242) = 0.08, p > 0.9 | F(10, 231)=0.03, p > 0.99 | F(10, 244) = 0.2, *p =* 0.9 | F(10, 242) = 0.6, *p =* 0.7 |

***Supplemental Table 10A*.** Statistical Information of DTI neuroimaging parameters in hippocampal subfields

|  |  |  |  |  |  | **FA** | | **MD** | | **AD** | | **RD** | |
| --- | --- | --- | --- | --- | --- | --- | --- | --- | --- | --- | --- | --- | --- |
| **R0I 1** | **Diet1** | **Exp1** | **R0I 2** | **Diet2** | **Exp 2** | **Mean Diff** | ***p*** | **Mean Diff** | ***p*** | **Mean Diff** | ***p*** | **Mean Diff** | ***p*** |
| dCA1 | CD | EXP | EcrhC | CD | EXP | 0.0242 | 0.9997 | -0.0001 | >0.999 | -0.0001 | >0.999 | -0.0001 | 0.9997 |
| dCA1 | CD | EXP | CA2 | CD | EXP | 0.0613 | **0.0064** | -0.0001 | 0.4618 | -0.0001 | 0.8049 | -0.0002 | 0.3068 |
| dCA1 | CD | EXP | dCA3 | CD | EXP | 0.0337 | 0.9099 | -0.0002 | **0.0210** | -0.0002 | **0.0250** | -0.0002 | **0.0216** |
| dCA1 | CD | EXP | dDG | CD | EXP | 0.0313 | 0.9653 | -0.0001 | 0.6839 | -0.0001 | 0.7896 | -0.0001 | 0.6387 |
| dCA1 | CD | EXP | dSb | CD | EXP | 0.0002 | >0.999 | 0.0000 | >0.999 | 0.0000 | >0.999 | -0.0001 | >0.999 |
| dCA1 | CD | EXP | EC | CD | EXP | 0.0384 | 0.6946 | -0.0001 | 0.9977 | -0.0001 | >0.999 | -0.0001 | 0.9894 |
| dCA1 | CD | EXP | vCA1 | CD | EXP | 0.0138 | >0.999 | -0.0001 | >0.999 | -0.0001 | >0.999 | -0.0001 | >0.999 |
| dCA1 | CD | EXP | vCA3 | CD | EXP | -0.0028 | >0.999 | -0.0002 | 0.1857 | -0.0002 | 0.0766 | -0.0002 | 0.2984 |
| dCA1 | CD | EXP | vDG | CD | EXP | 0.0368 | 0.7864 | -0.0002 | 0.1464 | -0.0002 | 0.2492 | -0.0002 | 0.1152 |
| dCA1 | CD | EXP | vSb | CD | EXP | 0.0084 | >0.999 | 0.0000 | >0.999 | 0.0000 | >0.999 | 0.0000 | >0.999 |
| dCA1 | CD | EXP | EcrhC | CD | UNEX | 0.0357 | 0.8903 | -0.0001 | 0.9998 | -0.0001 | >0.999 | -0.0001 | 0.9993 |
| dCA1 | CD | EXP | CA2 | CD | UNEX | 0.0526 | 0.1072 | -0.0001 | 0.7788 | -0.0001 | 0.9353 | -0.0001 | 0.6753 |
| dCA1 | CD | EXP | dCA3 | CD | UNEX | 0.0333 | 0.9532 | -0.0002 | 0.1838 | -0.0002 | 0.1936 | -0.0002 | 0.1924 |
| dCA1 | CD | EXP | dDG | CD | UNEX | 0.0288 | 0.9952 | -0.0001 | 0.8690 | -0.0001 | 0.9192 | -0.0001 | 0.8455 |
| dCA1 | CD | EXP | dSb | CD | UNEX | 0.0015 | >0.999 | 0.0000 | >0.999 | 0.0000 | >0.999 | 0.0000 | >0.999 |
| dCA1 | CD | EXP | EC | CD | UNEX | 0.0402 | 0.6826 | -0.0001 | 0.9769 | -0.0001 | 0.9979 | -0.0001 | 0.9467 |
| dCA1 | CD | EXP | vCA1 | CD | UNEX | 0.0083 | >0.999 | 0.0000 | >0.999 | -0.0001 | >0.999 | 0.0000 | >0.999 |
| dCA1 | CD | EXP | vCA3 | CD | UNEX | -0.0166 | >0.999 | -0.0001 | 0.5856 | -0.0002 | 0.1920 | -0.0001 | 0.8246 |
| dCA1 | CD | EXP | vDG | CD | UNEX | 0.0314 | 0.9802 | -0.0002 | 0.2459 | -0.0002 | 0.2987 | -0.0002 | 0.2236 |
| dCA1 | CD | EXP | vSb | CD | UNEX | 0.0146 | >0.999 | 0.0000 | >0.999 | 0.0000 | >0.999 | 0.0000 | >0.999 |
| dCA1 | CD | UNEX | CA2 | CD | EXP | 0.0718 | **0.0005** | -0.0001 | 0.5868 | -0.0001 | 0.9195 | -0.0002 | 0.3878 |
| dCA1 | CD | UNEX | dCA1 | CD | EXP | 0.0105 | >0.999 | 0.0000 | >0.999 | 0.0000 | >0.999 | 0.0000 | >0.999 |
| dCA1 | CD | UNEX | dCA3 | CD | EXP | 0.0442 | 0.4471 | -0.0002 | **0.0418** | -0.0002 | 0.0647 | -0.0002 | **0.0365** |
| dCA1 | CD | UNEX | dDG | CD | EXP | 0.0418 | 0.5910 | -0.0001 | 0.7879 | -0.0001 | 0.9107 | -0.0001 | 0.7155 |
| dCA1 | CD | UNEX | dSb | CD | EXP | 0.0107 | >0.999 | 0.0000 | >0.999 | 0.0000 | >0.999 | -0.0001 | >0.999 |
| dCA1 | CD | UNEX | EC | CD | EXP | 0.0490 | 0.2181 | -0.0001 | 0.9992 | -0.0001 | >0.999 | -0.0001 | 0.9938 |
| dCA1 | CD | UNEX | EcrhC | CD | EXP | 0.0347 | 0.9207 | -0.0001 | >0.999 | -0.0001 | >0.999 | -0.0001 | 0.9999 |
| dCA1 | CD | UNEX | vCA1 | CD | EXP | 0.0243 | 0.9999 | -0.0001 | >0.999 | -0.0001 | >0.999 | -0.0001 | >0.999 |
| dCA1 | CD | UNEX | vCA3 | CD | EXP | 0.0077 | >0.999 | -0.0002 | 0.2792 | -0.0002 | 0.1658 | -0.0002 | 0.3787 |
| dCA1 | CD | UNEX | vDG | CD | EXP | 0.0473 | 0.2893 | -0.0002 | 0.2283 | -0.0002 | 0.4227 | -0.0002 | 0.1647 |
| dCA1 | CD | UNEX | vSb | CD | EXP | 0.0189 | >0.999 | 0.0000 | >0.999 | 0.0000 | >0.999 | 0.0000 | >0.999 |
| dCA1 | CD | UNEX | CA2 | CD | UNEX | 0.0631 | **0.0148** | -0.0001 | 0.8576 | -0.0001 | 0.9801 | -0.0001 | 0.7425 |
| dCA1 | CD | UNEX | dCA3 | CD | UNEX | 0.0438 | 0.5663 | -0.0002 | 0.2698 | -0.0002 | 0.3384 | -0.0002 | 0.2531 |
| dCA1 | CD | UNEX | dDG | CD | UNEX | 0.0393 | 0.8062 | -0.0001 | 0.9231 | -0.0001 | 0.9734 | -0.0001 | 0.8868 |
| dCA1 | CD | UNEX | dSb | CD | UNEX | 0.0120 | >0.999 | 0.0000 | >0.999 | 0.0000 | >0.999 | 0.0000 | >0.999 |
| dCA1 | CD | UNEX | EC | CD | UNEX | 0.0507 | 0.2205 | -0.0001 | 0.9894 | -0.0001 | 0.9997 | -0.0001 | 0.9641 |
| dCA1 | CD | UNEX | EcrhC | CD | UNEX | 0.0462 | 0.4289 | -0.0001 | >0.999 | -0.0001 | >0.999 | -0.0001 | 0.9996 |
| dCA1 | CD | UNEX | vCA1 | CD | UNEX | 0.0188 | >0.999 | 0.0000 | >0.999 | 0.0000 | >0.999 | 0.0000 | >0.999 |
| dCA1 | CD | UNEX | vCA3 | CD | UNEX | -0.0061 | >0.999 | -0.0001 | 0.6960 | -0.0002 | 0.3362 | -0.0001 | 0.8699 |
| dCA1 | CD | UNEX | vDG | CD | UNEX | 0.0419 | 0.6767 | -0.0002 | 0.3452 | -0.0002 | 0.4726 | -0.0002 | 0.2890 |
| dCA1 | CD | UNEX | vSb | CD | UNEX | 0.0251 | 0.9999 | 0.0000 | >0.999 | 0.0000 | >0.999 | 0.0000 | >0.999 |
| dCA1 | CD | EXP | EcrhC | WD | EXP | 0.0138 | >0.999 | -0.0001 | 0.9004 | -0.0001 | 0.9327 | -0.0001 | 0.8888 |
| dCA1 | CD | EXP | CA2 | WD | EXP | 0.0457 | 0.2759 | -0.0001 | 0.5019 | -0.0001 | 0.7973 | -0.0001 | 0.3551 |
| dCA1 | CD | EXP | dCA1 | WD | EXP | -0.0051 | >0.999 | 0.0000 | >0.999 | 0.0000 | >0.999 | 0.0000 | >0.999 |
| dCA1 | CD | EXP | dCA3 | WD | EXP | 0.0264 | 0.9979 | -0.0002 | 0.1813 | -0.0002 | 0.2260 | -0.0002 | 0.1706 |
| dCA1 | CD | EXP | dDG | WD | EXP | 0.0249 | 0.9994 | -0.0001 | 0.9610 | -0.0001 | 0.9684 | -0.0001 | 0.9620 |
| dCA1 | CD | EXP | dSb | WD | EXP | -0.0020 | >0.999 | 0.0000 | >0.999 | 0.0000 | >0.999 | 0.0000 | >0.999 |
| dCA1 | CD | EXP | EC | WD | EXP | 0.0399 | 0.6091 | -0.0001 | 0.7862 | -0.0001 | 0.9503 | -0.0001 | 0.6643 |
| dCA1 | CD | EXP | vCA1 | WD | EXP | 0.0004 | >0.999 | 0.0000 | >0.999 | -0.0001 | >0.999 | 0.0000 | >0.999 |
| dCA1 | CD | EXP | vCA3 | WD | EXP | -0.0152 | >0.999 | -0.0001 | 0.5426 | -0.0002 | 0.4152 | -0.0001 | 0.6362 |
| dCA1 | CD | EXP | vDG | WD | EXP | 0.0215 | >0.999 | -0.0001 | 0.8563 | -0.0001 | 0.7817 | -0.0001 | 0.9030 |
| dCA1 | CD | EXP | vSb | WD | EXP | 0.0059 | >0.999 | -0.0001 | >0.999 | -0.0001 | >0.999 | -0.0001 | >0.999 |
| dCA1 | CD | EXP | EcrhC | WD | UNEX | -0.0062 | >0.999 | 0.0000 | >0.999 | 0.0000 | >0.999 | 0.0000 | >0.999 |
| dCA1 | CD | EXP | CA2 | WD | UNEX | 0.0462 | 0.3388 | -0.0001 | 0.7997 | -0.0001 | 0.9668 | -0.0001 | 0.6612 |
| dCA1 | CD | EXP | dCA1 | WD | UNEX | -0.0453 | 0.3860 | 0.0001 | >0.999 | 0.0001 | >0.999 | 0.0001 | >0.999 |
| dCA1 | CD | EXP | dCA3 | WD | UNEX | 0.0188 | >0.999 | -0.0002 | 0.4439 | -0.0002 | 0.4917 | -0.0002 | 0.4322 |
| dCA1 | CD | EXP | dDG | WD | UNEX | 0.0076 | >0.999 | -0.0001 | >0.999 | -0.0001 | >0.999 | -0.0001 | >0.999 |
| dCA1 | CD | EXP | dSb | WD | UNEX | -0.0435 | 0.4883 | 0.0000 | >0.999 | 0.0000 | >0.999 | 0.0000 | >0.999 |
| dCA1 | CD | EXP | EC | WD | UNEX | 0.0283 | 0.9964 | -0.0001 | >0.999 | -0.0001 | >0.999 | -0.0001 | >0.999 |
| dCA1 | CD | EXP | vCA1 | WD | UNEX | -0.0064 | >0.999 | 0.0000 | >0.999 | 0.0000 | >0.999 | 0.0000 | >0.999 |
| dCA1 | CD | EXP | vCA3 | WD | UNEX | -0.0203 | >0.999 | -0.0001 | 0.6619 | -0.0002 | 0.4263 | -0.0001 | 0.7928 |
| dCA1 | CD | EXP | vDG | WD | UNEX | 0.0142 | >0.999 | -0.0001 | 0.9252 | -0.0001 | 0.9687 | -0.0001 | 0.9009 |
| dCA1 | CD | EXP | vSb | WD | UNEX | -0.0155 | >0.999 | 0.0000 | >0.999 | 0.0000 | >0.999 | 0.0000 | >0.999 |
| dCA1 | CD | UNEX | CA2 | WD | EXP | 0.0562 | **0.0481** | -0.0001 | 0.6259 | -0.0001 | 0.9152 | -0.0002 | 0.4394 |
| dCA1 | CD | UNEX | dCA1 | WD | EXP | 0.0054 | >0.999 | 0.0000 | >0.999 | 0.0000 | >0.999 | 0.0000 | >0.999 |
| dCA1 | CD | UNEX | dCA3 | WD | EXP | 0.0369 | 0.8456 | -0.0002 | 0.2736 | -0.0002 | 0.3926 | -0.0002 | 0.2329 |
| dCA1 | CD | UNEX | dDG | WD | EXP | 0.0354 | 0.9023 | -0.0001 | 0.9819 | -0.0001 | 0.9927 | -0.0001 | 0.9759 |
| dCA1 | CD | UNEX | dSb | WD | EXP | 0.0085 | >0.999 | 0.0000 | >0.999 | 0.0000 | >0.999 | 0.0000 | >0.999 |
| dCA1 | CD | UNEX | EC | WD | EXP | 0.0504 | 0.1681 | -0.0001 | 0.8679 | -0.0001 | 0.9869 | -0.0001 | 0.7383 |
| dCA1 | CD | UNEX | EcrhC | WD | EXP | 0.0243 | 0.9999 | -0.0001 | 0.9464 | -0.0001 | 0.9806 | -0.0001 | 0.9233 |
| dCA1 | CD | UNEX | vCA1 | WD | EXP | 0.0109 | >0.999 | 0.0000 | >0.999 | 0.0000 | >0.999 | 0.0000 | >0.999 |
| dCA1 | CD | UNEX | vCA3 | WD | EXP | -0.0047 | >0.999 | -0.0001 | 0.6643 | -0.0002 | 0.6122 | -0.0001 | 0.7132 |
| dCA1 | CD | UNEX | vDG | WD | EXP | 0.0320 | 0.9737 | -0.0001 | 0.9177 | -0.0001 | 0.9061 | -0.0001 | 0.9339 |
| dCA1 | CD | UNEX | vSb | WD | EXP | 0.0164 | >0.999 | -0.0001 | >0.999 | -0.0001 | >0.999 | -0.0001 | >0.999 |
| dCA1 | CD | UNEX | CA2 | WD | UNEX | 0.0568 | 0.0683 | -0.0001 | 0.8734 | -0.0001 | 0.9915 | -0.0001 | 0.7299 |
| dCA1 | CD | UNEX | dCA1 | WD | UNEX | -0.0348 | 0.9480 | 0.0001 | >0.999 | 0.0001 | >0.999 | 0.0001 | >0.999 |
| dCA1 | CD | UNEX | dCA3 | WD | UNEX | 0.0293 | 0.9966 | -0.0002 | 0.5606 | -0.0002 | 0.6762 | -0.0002 | 0.5119 |
| dCA1 | CD | UNEX | dDG | WD | UNEX | 0.0181 | >0.999 | -0.0001 | >0.999 | -0.0001 | >0.999 | -0.0001 | >0.999 |
| dCA1 | CD | UNEX | dSb | WD | UNEX | -0.0330 | 0.9751 | 0.0000 | >0.999 | 0.0000 | >0.999 | 0.0000 | >0.999 |
| dCA1 | CD | UNEX | EC | WD | UNEX | 0.0388 | 0.8253 | -0.0001 | >0.999 | 0.0000 | >0.999 | -0.0001 | >0.999 |
| dCA1 | CD | UNEX | EcrhC | WD | UNEX | 0.0043 | >0.999 | 0.0000 | >0.999 | 0.0000 | >0.999 | 0.0000 | >0.999 |
| dCA1 | CD | UNEX | vCA1 | WD | UNEX | 0.0041 | >0.999 | 0.0000 | >0.999 | 0.0000 | >0.999 | 0.0000 | >0.999 |
| dCA1 | CD | UNEX | vCA3 | WD | UNEX | -0.0098 | >0.999 | -0.0001 | 0.7631 | -0.0002 | 0.6126 | -0.0001 | 0.8438 |
| dCA1 | CD | UNEX | vDG | WD | UNEX | 0.0247 | >0.999 | -0.0001 | 0.9598 | -0.0001 | 0.9921 | -0.0001 | 0.9301 |
| dCA1 | CD | UNEX | vSb | WD | UNEX | -0.0050 | >0.999 | 0.0000 | >0.999 | 0.0000 | >0.999 | 0.0000 | >0.999 |
| vCA1 | CD | UNEX | CA2 | CD | UNEX | 0.0444 | 0.5351 | -0.0001 | 0.9999 | -0.0001 | >0.999 | -0.0001 | 0.9989 |
| vCA1 | CD | UNEX | dCA3 | CD | UNEX | 0.0250 | 0.9999 | -0.0001 | 0.9087 | -0.0001 | 0.9448 | -0.0001 | 0.8932 |
| vCA1 | CD | UNEX | dDG | CD | UNEX | 0.0205 | >0.999 | -0.0001 | >0.999 | -0.0001 | >0.999 | -0.0001 | >0.999 |
| vCA1 | CD | UNEX | dSb | CD | UNEX | -0.0068 | >0.999 | 0.0000 | >0.999 | 0.0000 | >0.999 | 0.0000 | >0.999 |
| vCA1 | CD | UNEX | EC | CD | UNEX | 0.0320 | 0.9850 | -0.0001 | >0.999 | 0.0000 | >0.999 | -0.0001 | >0.999 |
| vCA1 | CD | UNEX | EcrhC | CD | UNEX | 0.0275 | 0.9991 | 0.0000 | >0.999 | 0.0000 | >0.999 | 0.0000 | >0.999 |
| vCA1 | CD | UNEX | vCA3 | CD | UNEX | -0.0248 | >0.999 | -0.0001 | 0.9980 | -0.0001 | 0.9439 | -0.0001 | >0.999 |
| vCA1 | CD | UNEX | vDG | CD | UNEX | 0.0231 | >0.999 | -0.0001 | 0.9478 | -0.0001 | 0.9809 | -0.0001 | 0.9176 |
| vCA1 | CD | UNEX | vSb | CD | UNEX | 0.0063 | >0.999 | 0.0000 | >0.999 | 0.0000 | >0.999 | 0.0000 | >0.999 |
| vCA1 | CD | UNEX | CA2 | CD | EXP | 0.0531 | 0.0977 | -0.0001 | 0.9950 | -0.0001 | >0.999 | -0.0001 | 0.9676 |
| vCA1 | CD | UNEX | dCA3 | CD | EXP | 0.0255 | 0.9996 | -0.0002 | 0.4744 | -0.0002 | 0.5837 | -0.0002 | 0.4358 |
| vCA1 | CD | UNEX | dDG | CD | EXP | 0.0230 | >0.999 | -0.0001 | 0.9997 | -0.0001 | >0.999 | -0.0001 | 0.9988 |
| vCA1 | CD | UNEX | dSb | CD | EXP | -0.0080 | >0.999 | 0.0000 | >0.999 | 0.0000 | >0.999 | 0.0000 | >0.999 |
| vCA1 | CD | UNEX | EC | CD | EXP | 0.0302 | 0.9892 | 0.0000 | >0.999 | 0.0000 | >0.999 | -0.0001 | >0.999 |
| vCA1 | CD | UNEX | EcrhC | CD | EXP | 0.0160 | >0.999 | 0.0000 | >0.999 | 0.0000 | >0.999 | 0.0000 | >0.999 |
| vCA1 | CD | UNEX | vCA3 | CD | EXP | -0.0110 | >0.999 | -0.0001 | 0.9275 | -0.0001 | 0.8245 | -0.0001 | 0.9652 |
| vCA1 | CD | UNEX | vDG | CD | EXP | 0.0285 | 0.9960 | -0.0001 | 0.8926 | -0.0001 | 0.9769 | -0.0001 | 0.8183 |
| vCA1 | CD | UNEX | vSb | CD | EXP | 0.0001 | >0.999 | 0.0000 | >0.999 | 0.0000 | >0.999 | 0.0000 | >0.999 |
| vCA1 | CD | UNEX | vCA1 | CD | EXP | 0.0055 | >0.999 | 0.0000 | >0.999 | 0.0000 | >0.999 | 0.0000 | >0.999 |
| vCA1 | CD | UNEX | CA2 | WD | UNEX | 0.0380 | 0.8595 | -0.0001 | >0.999 | -0.0001 | >0.999 | -0.0001 | 0.9986 |
| vCA1 | CD | UNEX | dCA3 | WD | UNEX | 0.0105 | >0.999 | -0.0001 | 0.9914 | -0.0001 | 0.9975 | -0.0001 | 0.9857 |
| vCA1 | CD | UNEX | dDG | WD | UNEX | -0.0007 | >0.999 | 0.0000 | >0.999 | 0.0000 | >0.999 | 0.0000 | >0.999 |
| vCA1 | CD | UNEX | dSb | WD | UNEX | -0.0518 | 0.1830 | 0.0001 | >0.999 | 0.0001 | >0.999 | 0.0001 | >0.999 |
| vCA1 | CD | UNEX | EC | WD | UNEX | 0.0201 | >0.999 | 0.0000 | >0.999 | 0.0000 | >0.999 | 0.0000 | >0.999 |
| vCA1 | CD | UNEX | EcrhC | WD | UNEX | -0.0145 | >0.999 | 0.0000 | >0.999 | 0.0000 | >0.999 | 0.0000 | >0.999 |
| vCA1 | CD | UNEX | vCA3 | WD | UNEX | -0.0285 | 0.9980 | -0.0001 | 0.9992 | -0.0001 | 0.9948 | -0.0001 | 0.9998 |
| vCA1 | CD | UNEX | vDG | WD | UNEX | 0.0059 | >0.999 | -0.0001 | >0.999 | -0.0001 | >0.999 | -0.0001 | >0.999 |
| vCA1 | CD | UNEX | vSb | WD | UNEX | -0.0238 | >0.999 | 0.0001 | >0.999 | 0.0001 | >0.999 | 0.0001 | >0.999 |
| vCA1 | CD | UNEX | vCA1 | WD | UNEX | -0.0147 | >0.999 | 0.0000 | >0.999 | 0.0000 | >0.999 | 0.0000 | >0.999 |
| vCA1 | CD | UNEX | CA2 | WD | EXP | 0.0374 | 0.8251 | -0.0001 | 0.9967 | -0.0001 | >0.999 | -0.0001 | 0.9789 |
| vCA1 | CD | UNEX | dCA3 | WD | EXP | 0.0182 | >0.999 | -0.0001 | 0.9243 | -0.0001 | 0.9703 | -0.0001 | 0.8925 |
| vCA1 | CD | UNEX | dDG | WD | EXP | 0.0166 | >0.999 | -0.0001 | >0.999 | -0.0001 | >0.999 | -0.0001 | >0.999 |
| vCA1 | CD | UNEX | dSb | WD | EXP | -0.0103 | >0.999 | 0.0000 | >0.999 | 0.0000 | >0.999 | 0.0000 | >0.999 |
| vCA1 | CD | UNEX | EC | WD | EXP | 0.0316 | 0.9777 | -0.0001 | >0.999 | -0.0001 | >0.999 | -0.0001 | 0.9992 |
| vCA1 | CD | UNEX | EcrhC | WD | EXP | 0.0055 | >0.999 | -0.0001 | >0.999 | -0.0001 | >0.999 | -0.0001 | >0.999 |
| vCA1 | CD | UNEX | vCA3 | WD | EXP | -0.0235 | >0.999 | -0.0001 | 0.9979 | -0.0001 | 0.9962 | -0.0001 | 0.9988 |
| vCA1 | CD | UNEX | vDG | WD | EXP | 0.0132 | >0.999 | -0.0001 | >0.999 | -0.0001 | >0.999 | -0.0001 | >0.999 |
| vCA1 | CD | UNEX | vSb | WD | EXP | -0.0024 | >0.999 | 0.0000 | >0.999 | 0.0000 | >0.999 | 0.0000 | >0.999 |
| vCA1 | CD | UNEX | vCA1 | WD | EXP | -0.0079 | >0.999 | 0.0000 | >0.999 | 0.0000 | >0.999 | 0.0000 | >0.999 |
| vCA1 | CD | EXP | EcrhC | CD | UNEX | 0.0220 | >0.999 | 0.0000 | >0.999 | 0.0000 | >0.999 | 0.0000 | >0.999 |
| vCA1 | CD | EXP | CA2 | CD | UNEX | 0.0389 | 0.7563 | -0.0001 | >0.999 | -0.0001 | >0.999 | -0.0001 | >0.999 |
| vCA1 | CD | EXP | dCA3 | CD | UNEX | 0.0196 | >0.999 | -0.0001 | 0.9790 | -0.0001 | 0.9828 | -0.0001 | 0.9785 |
| vCA1 | CD | EXP | dDG | CD | UNEX | 0.0150 | >0.999 | -0.0001 | >0.999 | -0.0001 | >0.999 | -0.0001 | >0.999 |
| vCA1 | CD | EXP | dSb | CD | UNEX | -0.0123 | >0.999 | 0.0000 | >0.999 | 0.0000 | >0.999 | 0.0000 | >0.999 |
| vCA1 | CD | EXP | EC | CD | UNEX | 0.0265 | 0.9991 | 0.0000 | >0.999 | 0.0000 | >0.999 | -0.0001 | >0.999 |
| vCA1 | CD | EXP | vCA3 | CD | UNEX | -0.0303 | 0.9884 | -0.0001 | >0.999 | -0.0001 | 0.9824 | -0.0001 | >0.999 |
| vCA1 | CD | EXP | vDG | CD | UNEX | 0.0176 | >0.999 | -0.0001 | 0.9911 | -0.0001 | 0.9960 | -0.0001 | 0.9859 |
| vCA1 | CD | EXP | vSb | CD | UNEX | 0.0009 | >0.999 | 0.0000 | >0.999 | 0.0000 | >0.999 | 0.0000 | >0.999 |
| vCA1 | CD | EXP | EcrhC | CD | EXP | 0.0105 | >0.999 | 0.0000 | >0.999 | 0.0000 | >0.999 | 0.0000 | >0.999 |
| vCA1 | CD | EXP | CA2 | CD | EXP | 0.0476 | 0.1970 | -0.0001 | 0.9998 | -0.0001 | >0.999 | -0.0001 | 0.9971 |
| vCA1 | CD | EXP | dCA3 | CD | EXP | 0.0200 | >0.999 | -0.0001 | 0.6766 | -0.0001 | 0.7208 | -0.0001 | 0.6643 |
| vCA1 | CD | EXP | dDG | CD | EXP | 0.0175 | >0.999 | -0.0001 | >0.999 | -0.0001 | >0.999 | -0.0001 | >0.999 |
| vCA1 | CD | EXP | dSb | CD | EXP | -0.0135 | >0.999 | 0.0000 | >0.999 | 0.0000 | >0.999 | 0.0000 | >0.999 |
| vCA1 | CD | EXP | EC | CD | EXP | 0.0247 | 0.9995 | 0.0000 | >0.999 | 0.0000 | >0.999 | 0.0000 | >0.999 |
| vCA1 | CD | EXP | vCA3 | CD | EXP | -0.0165 | >0.999 | -0.0001 | 0.9860 | -0.0001 | 0.9169 | -0.0001 | 0.9967 |
| vCA1 | CD | EXP | vDG | CD | EXP | 0.0230 | 0.9999 | -0.0001 | 0.9742 | -0.0001 | 0.9949 | -0.0001 | 0.9512 |
| vCA1 | CD | EXP | vSb | CD | EXP | -0.0054 | >0.999 | 0.0000 | >0.999 | 0.0000 | >0.999 | 0.0000 | >0.999 |
| vCA1 | CD | EXP | EcrhC | WD | UNEX | -0.0200 | >0.999 | 0.0000 | >0.999 | 0.0001 | >0.999 | 0.0000 | >0.999 |
| vCA1 | CD | EXP | CA2 | WD | UNEX | 0.0325 | 0.9666 | -0.0001 | >0.999 | -0.0001 | >0.999 | -0.0001 | >0.999 |
| vCA1 | CD | EXP | dCA3 | WD | UNEX | 0.0050 | >0.999 | -0.0001 | 0.9994 | -0.0001 | 0.9997 | -0.0001 | 0.9991 |
| vCA1 | CD | EXP | dDG | WD | UNEX | -0.0062 | >0.999 | 0.0000 | >0.999 | 0.0000 | >0.999 | 0.0000 | >0.999 |
| vCA1 | CD | EXP | dSb | WD | UNEX | -0.0573 | **0.0370** | 0.0001 | 0.9988 | 0.0001 | 0.9981 | 0.0001 | 0.9992 |
| vCA1 | CD | EXP | EC | WD | UNEX | 0.0146 | >0.999 | 0.0000 | >0.999 | 0.0000 | >0.999 | 0.0000 | >0.999 |
| vCA1 | CD | EXP | vCA1 | WD | UNEX | -0.0202 | >0.999 | 0.0000 | >0.999 | 0.0000 | >0.999 | 0.0000 | >0.999 |
| vCA1 | CD | EXP | vCA3 | WD | UNEX | -0.0340 | 0.9384 | -0.0001 | >0.999 | -0.0001 | 0.9993 | -0.0001 | >0.999 |
| vCA1 | CD | EXP | vDG | WD | UNEX | 0.0004 | >0.999 | -0.0001 | >0.999 | -0.0001 | >0.999 | -0.0001 | >0.999 |
| vCA1 | CD | EXP | vSb | WD | UNEX | -0.0293 | 0.9936 | 0.0001 | 0.9998 | 0.0001 | 0.9986 | 0.0001 | >0.999 |
| vCA1 | CD | EXP | EcrhC | WD | EXP | 0.0001 | >0.999 | -0.0001 | >0.999 | -0.0001 | >0.999 | -0.0001 | >0.999 |
| vCA1 | CD | EXP | CA2 | WD | EXP | 0.0319 | 0.9542 | -0.0001 | 0.9999 | -0.0001 | >0.999 | -0.0001 | 0.9985 |
| vCA1 | CD | EXP | dCA3 | WD | EXP | 0.0127 | >0.999 | -0.0001 | 0.9850 | -0.0001 | 0.9929 | -0.0001 | 0.9795 |
| vCA1 | CD | EXP | dDG | WD | EXP | 0.0111 | >0.999 | 0.0000 | >0.999 | 0.0000 | >0.999 | 0.0000 | >0.999 |
| vCA1 | CD | EXP | dSb | WD | EXP | -0.0158 | >0.999 | 0.0000 | >0.999 | 0.0000 | >0.999 | 0.0000 | >0.999 |
| vCA1 | CD | EXP | EC | WD | EXP | 0.0261 | 0.9984 | -0.0001 | >0.999 | 0.0000 | >0.999 | -0.0001 | >0.999 |
| vCA1 | CD | EXP | vCA1 | WD | EXP | -0.0134 | >0.999 | 0.0000 | >0.999 | 0.0000 | >0.999 | 0.0000 | >0.999 |
| vCA1 | CD | EXP | vCA3 | WD | EXP | -0.0290 | 0.9892 | -0.0001 | >0.999 | -0.0001 | 0.9996 | -0.0001 | >0.999 |
| vCA1 | CD | EXP | vDG | WD | EXP | 0.0077 | >0.999 | -0.0001 | >0.999 | -0.0001 | >0.999 | -0.0001 | >0.999 |
| vCA1 | CD | EXP | vSb | WD | EXP | -0.0079 | >0.999 | 0.0000 | >0.999 | 0.0000 | >0.999 | 0.0000 | >0.999 |
| CA2 | CD | UNEX | EcrhC | CD | UNEX | -0.0169 | >0.999 | 0.0000 | >0.999 | 0.0000 | >0.999 | 0.0000 | >0.999 |
| CA2 | CD | UNEX | dCA3 | CD | UNEX | -0.0193 | >0.999 | 0.0000 | >0.999 | -0.0001 | >0.999 | 0.0000 | >0.999 |
| CA2 | CD | UNEX | dDG | CD | UNEX | -0.0239 | >0.999 | 0.0000 | >0.999 | 0.0000 | >0.999 | 0.0000 | >0.999 |
| CA2 | CD | UNEX | dSb | CD | UNEX | -0.0512 | 0.2047 | 0.0001 | 0.9981 | 0.0001 | 0.9997 | 0.0001 | 0.9962 |
| CA2 | CD | UNEX | EC | CD | UNEX | -0.0124 | >0.999 | 0.0000 | >0.999 | 0.0000 | >0.999 | 0.0000 | >0.999 |
| CA2 | CD | UNEX | vCA3 | CD | UNEX | -0.0692 | **0.0028** | 0.0000 | >0.999 | -0.0001 | >0.999 | 0.0000 | >0.999 |
| CA2 | CD | UNEX | vDG | CD | UNEX | -0.0213 | >0.999 | 0.0000 | >0.999 | 0.0000 | >0.999 | 0.0000 | >0.999 |
| CA2 | CD | UNEX | vSb | CD | UNEX | -0.0380 | 0.8583 | 0.0001 | 0.9975 | 0.0001 | 0.9993 | 0.0001 | 0.9958 |
| CA2 | CD | UNEX | EcrhC | CD | EXP | -0.0284 | 0.9963 | 0.0001 | >0.999 | 0.0001 | >0.999 | 0.0001 | >0.999 |
| CA2 | CD | UNEX | CA2 | CD | EXP | 0.0087 | >0.999 | 0.0000 | >0.999 | 0.0000 | >0.999 | 0.0000 | >0.999 |
| CA2 | CD | UNEX | dCA3 | CD | EXP | -0.0189 | >0.999 | -0.0001 | >0.999 | -0.0001 | >0.999 | -0.0001 | >0.999 |
| CA2 | CD | UNEX | dDG | CD | EXP | -0.0213 | >0.999 | 0.0000 | >0.999 | 0.0000 | >0.999 | 0.0000 | >0.999 |
| CA2 | CD | UNEX | dSb | CD | EXP | -0.0524 | 0.1126 | 0.0001 | 0.9998 | 0.0001 | >0.999 | 0.0001 | 0.9997 |
| CA2 | CD | UNEX | EC | CD | EXP | -0.0142 | >0.999 | 0.0000 | >0.999 | 0.0000 | >0.999 | 0.0000 | >0.999 |
| CA2 | CD | UNEX | vCA3 | CD | EXP | -0.0554 | 0.0583 | 0.0000 | >0.999 | -0.0001 | >0.999 | 0.0000 | >0.999 |
| CA2 | CD | UNEX | vDG | CD | EXP | -0.0159 | >0.999 | 0.0000 | >0.999 | 0.0000 | >0.999 | 0.0000 | >0.999 |
| CA2 | CD | UNEX | vSb | CD | EXP | -0.0442 | 0.4482 | 0.0001 | 0.9880 | 0.0001 | 0.9971 | 0.0001 | 0.9795 |
| CA2 | CD | UNEX | EcrhC | WD | UNEX | -0.0589 | **0.0425** | 0.0001 | 0.9823 | 0.0001 | 0.9936 | 0.0001 | 0.9760 |
| CA2 | CD | UNEX | CA2 | WD | UNEX | -0.0064 | >0.999 | 0.0000 | >0.999 | 0.0000 | >0.999 | 0.0000 | >0.999 |
| CA2 | CD | UNEX | dCA3 | WD | UNEX | -0.0338 | 0.9651 | 0.0000 | >0.999 | 0.0000 | >0.999 | 0.0000 | >0.999 |
| CA2 | CD | UNEX | dDG | WD | UNEX | -0.0451 | 0.4943 | 0.0001 | >0.999 | 0.0001 | >0.999 | 0.0001 | >0.999 |
| CA2 | CD | UNEX | dSb | WD | UNEX | -0.0962 | <0.0001 | 0.0002 | 0.4018 | 0.0002 | 0.6126 | 0.0002 | 0.3185 |
| CA2 | CD | UNEX | EC | WD | UNEX | -0.0243 | >0.999 | 0.0001 | >0.999 | 0.0001 | >0.999 | 0.0001 | >0.999 |
| CA2 | CD | UNEX | vCA3 | WD | UNEX | -0.0729 | **0.0009** | 0.0000 | >0.999 | 0.0000 | >0.999 | 0.0000 | >0.999 |
| CA2 | CD | UNEX | vDG | WD | UNEX | -0.0385 | 0.8412 | 0.0000 | >0.999 | 0.0000 | >0.999 | 0.0000 | >0.999 |
| CA2 | CD | UNEX | vSb | WD | UNEX | -0.0681 | **0.0038** | 0.0002 | 0.5219 | 0.0002 | 0.6332 | 0.0002 | 0.4811 |
| CA2 | CD | UNEX | EcrhC | WD | EXP | -0.0388 | 0.7590 | 0.0000 | >0.999 | 0.0000 | >0.999 | 0.0000 | >0.999 |
| CA2 | CD | UNEX | CA2 | WD | EXP | -0.0069 | >0.999 | 0.0000 | >0.999 | 0.0000 | >0.999 | 0.0000 | >0.999 |
| CA2 | CD | UNEX | dCA3 | WD | EXP | -0.0262 | 0.9993 | 0.0000 | >0.999 | 0.0000 | >0.999 | 0.0000 | >0.999 |
| CA2 | CD | UNEX | dDG | WD | EXP | -0.0278 | 0.9976 | 0.0000 | >0.999 | 0.0000 | >0.999 | 0.0000 | >0.999 |
| CA2 | CD | UNEX | dSb | WD | EXP | -0.0546 | 0.0692 | 0.0001 | 0.9860 | 0.0001 | 0.9977 | 0.0001 | 0.9729 |
| CA2 | CD | UNEX | EC | WD | EXP | -0.0128 | >0.999 | 0.0000 | >0.999 | 0.0000 | >0.999 | 0.0000 | >0.999 |
| CA2 | CD | UNEX | vCA3 | WD | EXP | -0.0679 | **0.0019** | 0.0000 | >0.999 | 0.0000 | >0.999 | 0.0000 | >0.999 |
| CA2 | CD | UNEX | vDG | WD | EXP | -0.0312 | 0.9819 | 0.0000 | >0.999 | 0.0000 | >0.999 | 0.0000 | >0.999 |
| CA2 | CD | UNEX | vSb | WD | EXP | -0.0468 | 0.3136 | 0.0001 | >0.999 | 0.0001 | >0.999 | 0.0001 | 0.9999 |
| CA2 | CD | EXP | EcrhC | CD | UNEX | -0.0256 | 0.9995 | 0.0001 | >0.999 | 0.0001 | >0.999 | 0.0001 | >0.999 |
| CA2 | CD | EXP | dDG | CD | UNEX | -0.0326 | 0.9657 | 0.0000 | >0.999 | 0.0000 | >0.999 | 0.0000 | >0.999 |
| CA2 | CD | EXP | dSb | CD | UNEX | -0.0599 | **0.0192** | 0.0001 | 0.9695 | 0.0001 | 0.9964 | 0.0001 | 0.9344 |
| CA2 | CD | EXP | EC | CD | UNEX | -0.0211 | >0.999 | 0.0000 | >0.999 | 0.0000 | >0.999 | 0.0000 | >0.999 |
| CA2 | CD | EXP | vDG | CD | UNEX | -0.0300 | 0.9903 | 0.0000 | >0.999 | 0.0000 | >0.999 | 0.0000 | >0.999 |
| CA2 | CD | EXP | vSb | CD | UNEX | -0.0467 | 0.3154 | 0.0001 | 0.9632 | 0.0001 | 0.9933 | 0.0001 | 0.9308 |
| CA2 | CD | EXP | CA2 | CD | UNEX | -0.0280 | 0.9971 | 0.0000 | >0.999 | 0.0000 | >0.999 | 0.0000 | >0.999 |
| CA2 | CD | EXP | dCA3 | CD | UNEX | -0.0779 | **<0.0001** | 0.0000 | >0.999 | 0.0000 | >0.999 | 0.0000 | >0.999 |
| CA2 | CD | EXP | dCA3 | CD | EXP | -0.0276 | 0.9954 | -0.0001 | >0.999 | -0.0001 | >0.999 | 0.0000 | >0.999 |
| CA2 | CD | EXP | vCA3 | CD | EXP | -0.0641 | **0.0028** | 0.0000 | >0.999 | -0.0001 | >0.999 | 0.0000 | >0.999 |
| CA2 | CD | EXP | EcrhC | CD | EXP | -0.0371 | 0.7695 | 0.0001 | >0.999 | 0.0001 | >0.999 | 0.0001 | >0.999 |
| CA2 | CD | EXP | dDG | CD | EXP | -0.0300 | 0.9811 | 0.0000 | >0.999 | 0.0000 | >0.999 | 0.0000 | >0.999 |
| CA2 | CD | EXP | dSb | CD | EXP | -0.0611 | **0.0069** | 0.0001 | 0.9932 | 0.0001 | 0.9996 | 0.0001 | 0.9819 |
| CA2 | CD | EXP | EC | CD | EXP | -0.0229 | >0.999 | 0.0001 | >0.999 | 0.0001 | >0.999 | 0.0001 | >0.999 |
| CA2 | CD | EXP | vDG | CD | EXP | -0.0246 | 0.9996 | 0.0000 | >0.999 | 0.0000 | >0.999 | 0.0000 | >0.999 |
| CA2 | CD | EXP | vSb | CD | EXP | -0.0529 | 0.0623 | 0.0001 | 0.8949 | 0.0001 | 0.9786 | 0.0001 | 0.8177 |
| CA2 | CD | EXP | EcrhC | WD | UNEX | -0.0676 | **0.0021** | 0.0001 | 0.8768 | 0.0001 | 0.9650 | 0.0001 | 0.8151 |
| CA2 | CD | EXP | CA2 | WD | UNEX | -0.0151 | >0.999 | 0.0000 | >0.999 | 0.0000 | >0.999 | 0.0000 | >0.999 |
| CA2 | CD | EXP | dDG | WD | UNEX | -0.0538 | 0.0837 | 0.0001 | 0.9997 | 0.0001 | >0.999 | 0.0001 | 0.9984 |
| CA2 | CD | EXP | dSb | WD | UNEX | -0.1050 | **<0.0001** | 0.0002 | 0.1610 | 0.0002 | 0.3919 | 0.0002 | 0.0965 |
| CA2 | CD | EXP | EC | WD | UNEX | -0.0330 | 0.9587 | 0.0001 | >0.999 | 0.0001 | >0.999 | 0.0001 | >0.999 |
| CA2 | CD | EXP | vDG | WD | UNEX | -0.0472 | 0.2945 | 0.0000 | >0.999 | 0.0000 | >0.999 | 0.0000 | >0.999 |
| CA2 | CD | EXP | vSb | WD | UNEX | -0.0768 | **0.0001** | 0.0002 | 0.2388 | 0.0002 | 0.4118 | 0.0002 | 0.1800 |
| CA2 | CD | EXP | dCA3 | WD | UNEX | -0.0425 | 0.5477 | 0.0000 | >0.999 | 0.0000 | >0.999 | 0.0000 | >0.999 |
| CA2 | CD | EXP | vCA3 | WD | UNEX | -0.0816 | **<0.0001** | 0.0000 | >0.999 | 0.0000 | >0.999 | 0.0000 | >0.999 |
| CA2 | CD | EXP | EcrhC | WD | EXP | -0.0475 | 0.1989 | 0.0000 | >0.999 | 0.0000 | >0.999 | 0.0000 | >0.999 |
| CA2 | CD | EXP | CA2 | WD | EXP | -0.0156 | >0.999 | 0.0000 | >0.999 | 0.0000 | >0.999 | 0.0000 | >0.999 |
| CA2 | CD | EXP | dDG | WD | EXP | -0.0365 | 0.8015 | 0.0000 | >0.999 | 0.0000 | >0.999 | 0.0000 | >0.999 |
| CA2 | CD | EXP | dSb | WD | EXP | -0.0633 | **0.0035** | 0.0001 | 0.8849 | 0.0001 | 0.9825 | 0.0001 | 0.7890 |
| CA2 | CD | EXP | EC | WD | EXP | -0.0215 | >0.999 | 0.0000 | >0.999 | 0.0000 | >0.999 | 0.0000 | >0.999 |
| CA2 | CD | EXP | vDG | WD | EXP | -0.0399 | 0.6085 | 0.0000 | >0.999 | 0.0000 | >0.999 | 0.0000 | >0.999 |
| CA2 | CD | EXP | vSb | WD | EXP | -0.0555 | **0.0332** | 0.0001 | 0.9991 | 0.0001 | >0.999 | 0.0001 | 0.9899 |
| CA2 | CD | EXP | dCA3 | WD | EXP | -0.0349 | 0.8705 | 0.0000 | >0.999 | 0.0000 | >0.999 | 0.0000 | >0.999 |
| CA2 | CD | EXP | vCA3 | WD | EXP | -0.0766 | <0.0001 | 0.0000 | >0.999 | 0.0000 | >0.999 | 0.0000 | >0.999 |
| dCA3 | CD | UNEX | dDG | CD | UNEX | -0.0045 | >0.999 | 0.0000 | >0.999 | 0.0001 | >0.999 | 0.0000 | >0.999 |
| dCA3 | CD | UNEX | EC | CD | UNEX | 0.0069 | >0.999 | 0.0001 | >0.999 | 0.0001 | >0.999 | 0.0001 | >0.999 |
| dCA3 | CD | UNEX | vDG | CD | UNEX | -0.0020 | >0.999 | 0.0000 | >0.999 | 0.0000 | >0.999 | 0.0000 | >0.999 |
| dCA3 | CD | UNEX | dSb | CD | UNEX | -0.0319 | 0.9858 | 0.0001 | 0.7771 | 0.0002 | 0.6935 | 0.0001 | 0.8276 |
| dCA3 | CD | UNEX | EcrhC | CD | UNEX | 0.0024 | >0.999 | 0.0001 | 0.9999 | 0.0001 | 0.9986 | 0.0001 | >0.999 |
| dCA3 | CD | UNEX | vSb | CD | UNEX | -0.0187 | >0.999 | 0.0001 | 0.7560 | 0.0002 | 0.6383 | 0.0001 | 0.8213 |
| dCA3 | CD | UNEX | vCA3 | CD | UNEX | -0.0499 | 0.2548 | 0.0000 | >0.999 | 0.0000 | >0.999 | 0.0000 | >0.999 |
| dCA3 | CD | UNEX | dDG | CD | EXP | -0.0020 | >0.999 | 0.0000 | >0.999 | 0.0000 | >0.999 | 0.0000 | >0.999 |
| dCA3 | CD | UNEX | EC | CD | EXP | 0.0051 | >0.999 | 0.0001 | >0.999 | 0.0001 | 0.9976 | 0.0001 | >0.999 |
| dCA3 | CD | UNEX | vDG | CD | EXP | 0.0035 | >0.999 | 0.0000 | >0.999 | 0.0000 | >0.999 | 0.0000 | >0.999 |
| dCA3 | CD | UNEX | dSb | CD | EXP | -0.0331 | 0.9574 | 0.0001 | 0.8869 | 0.0001 | 0.8117 | 0.0001 | 0.9262 |
| dCA3 | CD | UNEX | EcrhC | CD | EXP | -0.0091 | >0.999 | 0.0001 | 0.9986 | 0.0001 | 0.9937 | 0.0001 | 0.9995 |
| dCA3 | CD | UNEX | vSb | CD | EXP | -0.0249 | 0.9997 | 0.0001 | 0.5889 | 0.0002 | 0.4906 | 0.0001 | 0.6551 |
| dCA3 | CD | UNEX | dCA3 | CD | EXP | 0.0004 | >0.999 | 0.0000 | >0.999 | 0.0000 | >0.999 | 0.0000 | >0.999 |
| dCA3 | CD | UNEX | vCA3 | CD | EXP | -0.0361 | 0.8793 | 0.0000 | >0.999 | 0.0000 | >0.999 | 0.0000 | >0.999 |
| dCA3 | CD | UNEX | dCA3 | WD | UNEX | -0.0145 | >0.999 | 0.0000 | >0.999 | 0.0000 | >0.999 | 0.0000 | >0.999 |
| dCA3 | CD | UNEX | vCA3 | WD | UNEX | -0.0536 | 0.1314 | 0.0000 | >0.999 | 0.0000 | >0.999 | 0.0000 | >0.999 |
| dCA3 | CD | UNEX | dDG | WD | UNEX | -0.0258 | 0.9998 | 0.0001 | 0.9783 | 0.0001 | 0.9589 | 0.0001 | 0.9864 |
| dCA3 | CD | UNEX | EC | WD | UNEX | -0.0050 | >0.999 | 0.0001 | 0.9907 | 0.0001 | 0.9254 | 0.0001 | 0.9984 |
| dCA3 | CD | UNEX | vDG | WD | UNEX | -0.0191 | >0.999 | 0.0001 | >0.999 | 0.0001 | >0.999 | 0.0000 | >0.999 |
| dCA3 | CD | UNEX | dSb | WD | UNEX | -0.0768 | **0.0003** | 0.0002 | **0.0492** | 0.0002 | **0.0466** | 0.0002 | 0.0553 |
| dCA3 | CD | UNEX | EcrhC | WD | UNEX | -0.0395 | 0.7941 | 0.0002 | 0.5689 | 0.0002 | 0.4524 | 0.0001 | 0.6554 |
| dCA3 | CD | UNEX | vSb | WD | UNEX | -0.0488 | 0.3000 | 0.0002 | 0.0794 | 0.0002 | 0.0507 | 0.0002 | 0.1075 |
| dCA3 | CD | UNEX | dDG | WD | EXP | -0.0084 | >0.999 | 0.0001 | >0.999 | 0.0001 | >0.999 | 0.0001 | >0.999 |
| dCA3 | CD | UNEX | EC | WD | EXP | 0.0066 | >0.999 | 0.0000 | >0.999 | 0.0001 | >0.999 | 0.0000 | >0.999 |
| dCA3 | CD | UNEX | vDG | WD | EXP | -0.0119 | >0.999 | 0.0001 | >0.999 | 0.0000 | >0.999 | 0.0001 | >0.999 |
| dCA3 | CD | UNEX | dSb | WD | EXP | -0.0353 | 0.9041 | 0.0001 | 0.5717 | 0.0002 | 0.5135 | 0.0001 | 0.6205 |
| dCA3 | CD | UNEX | EcrhC | WD | EXP | -0.0195 | >0.999 | 0.0001 | >0.999 | 0.0001 | >0.999 | 0.0001 | >0.999 |
| dCA3 | CD | UNEX | vSb | WD | EXP | -0.0274 | 0.9981 | 0.0001 | 0.9587 | 0.0001 | 0.9750 | 0.0001 | 0.9501 |
| dCA3 | CD | UNEX | dCA3 | WD | EXP | -0.0069 | >0.999 | 0.0000 | >0.999 | 0.0000 | >0.999 | 0.0000 | >0.999 |
| dCA3 | CD | UNEX | vCA3 | WD | EXP | -0.0485 | 0.2342 | 0.0000 | >0.999 | 0.0000 | >0.999 | 0.0000 | >0.999 |
| dCA3 | CD | EXP | vCA3 | CD | UNEX | -0.0503 | 0.1705 | 0.0001 | >0.999 | 0.0000 | >0.999 | 0.0001 | >0.999 |
| dCA3 | CD | EXP | dDG | CD | UNEX | -0.0050 | >0.999 | 0.0001 | >0.999 | 0.0001 | >0.999 | 0.0001 | >0.999 |
| dCA3 | CD | EXP | dSb | CD | UNEX | -0.0323 | 0.9694 | 0.0002 | 0.2928 | 0.0002 | 0.2371 | 0.0002 | 0.3401 |
| dCA3 | CD | EXP | EC | CD | UNEX | 0.0065 | >0.999 | 0.0001 | 0.9996 | 0.0001 | 0.9946 | 0.0001 | >0.999 |
| dCA3 | CD | EXP | EcrhC | CD | UNEX | 0.0020 | >0.999 | 0.0001 | 0.9658 | 0.0001 | 0.9143 | 0.0001 | 0.9837 |
| dCA3 | CD | EXP | vDG | CD | UNEX | -0.0024 | >0.999 | 0.0000 | >0.999 | 0.0000 | >0.999 | 0.0000 | >0.999 |
| dCA3 | CD | EXP | vSb | CD | UNEX | -0.0191 | >0.999 | 0.0002 | 0.2730 | 0.0002 | 0.1987 | 0.0002 | 0.3326 |
| dCA3 | CD | EXP | dDG | CD | EXP | -0.0025 | >0.999 | 0.0001 | >0.999 | 0.0001 | >0.999 | 0.0001 | >0.999 |
| dCA3 | CD | EXP | dSb | CD | EXP | -0.0335 | 0.9170 | 0.0001 | 0.4107 | 0.0002 | 0.3274 | 0.0001 | 0.4797 |
| dCA3 | CD | EXP | EC | CD | EXP | 0.0047 | >0.999 | 0.0001 | 0.9752 | 0.0001 | 0.8798 | 0.0001 | 0.9936 |
| dCA3 | CD | EXP | EcrhC | CD | EXP | -0.0095 | >0.999 | 0.0001 | 0.8921 | 0.0001 | 0.8161 | 0.0001 | 0.9277 |
| dCA3 | CD | EXP | vDG | CD | EXP | 0.0030 | >0.999 | 0.0000 | >0.999 | 0.0000 | >0.999 | 0.0000 | >0.999 |
| dCA3 | CD | EXP | vSb | CD | EXP | -0.0254 | 0.9991 | 0.0002 | 0.1452 | 0.0002 | 0.1105 | 0.0002 | 0.1764 |
| dCA3 | CD | EXP | vCA3 | CD | EXP | -0.0365 | 0.7995 | 0.0000 | >0.999 | 0.0000 | >0.999 | 0.0000 | >0.999 |
| dCA3 | CD | EXP | dCA3 | WD | UNEX | -0.0149 | >0.999 | 0.0000 | >0.999 | 0.0000 | >0.999 | 0.0000 | >0.999 |
| dCA3 | CD | EXP | vCA3 | WD | UNEX | -0.0540 | 0.0794 | 0.0001 | >0.999 | 0.0000 | >0.999 | 0.0001 | >0.999 |
| dCA3 | CD | EXP | dDG | WD | UNEX | -0.0262 | 0.9992 | 0.0001 | 0.6948 | 0.0002 | 0.6294 | 0.0001 | 0.7387 |
| dCA3 | CD | EXP | dSb | WD | UNEX | -0.0773 | **<0.0001** | 0.0002 | **0.0039** | 0.0002 | **0.0041** | 0.0002 | **0.0044** |
| dCA3 | CD | EXP | EC | WD | UNEX | -0.0054 | >0.999 | 0.0001 | 0.7836 | 0.0002 | 0.5323 | 0.0001 | 0.8935 |
| dCA3 | CD | EXP | EcrhC | WD | UNEX | -0.0400 | 0.6965 | 0.0002 | 0.1474 | 0.0002 | 0.1045 | 0.0002 | 0.1912 |
| dCA3 | CD | EXP | vDG | WD | UNEX | -0.0196 | >0.999 | 0.0001 | >0.999 | 0.0001 | 0.9999 | 0.0001 | >0.999 |
| dCA3 | CD | EXP | vSb | WD | UNEX | -0.0493 | 0.2066 | 0.0002 | **0.0074** | 0.0002 | **0.0046** | 0.0002 | **0.0107** |
| dCA3 | CD | EXP | dDG | WD | EXP | -0.0089 | >0.999 | 0.0001 | 0.9989 | 0.0001 | 0.9991 | 0.0001 | 0.9989 |
| dCA3 | CD | EXP | dSb | WD | EXP | -0.0358 | 0.8346 | 0.0002 | 0.1368 | 0.0002 | 0.1202 | 0.0002 | 0.1568 |
| dCA3 | CD | EXP | EC | WD | EXP | 0.0061 | >0.999 | 0.0001 | >0.999 | 0.0001 | 0.9996 | 0.0001 | >0.999 |
| dCA3 | CD | EXP | EcrhC | WD | EXP | -0.0199 | >0.999 | 0.0001 | 0.9999 | 0.0001 | 0.9998 | 0.0001 | >0.999 |
| dCA3 | CD | EXP | vDG | WD | EXP | -0.0123 | >0.999 | 0.0001 | >0.999 | 0.0001 | >0.999 | 0.0001 | 0.9999 |
| dCA3 | CD | EXP | vSb | WD | EXP | -0.0279 | 0.9945 | 0.0001 | 0.5836 | 0.0001 | 0.6757 | 0.0001 | 0.5448 |
| dCA3 | CD | EXP | dCA3 | WD | EXP | -0.0073 | >0.999 | 0.0000 | >0.999 | 0.0000 | >0.999 | 0.0000 | >0.999 |
| dCA3 | CD | EXP | vCA3 | WD | EXP | -0.0490 | 0.1496 | 0.0001 | >0.999 | 0.0000 | >0.999 | 0.0001 | >0.999 |
| vCA3 | CD | UNEX | dDG | CD | UNEX | 0.0453 | 0.4798 | 0.0000 | >0.999 | 0.0001 | >0.999 | 0.0000 | >0.999 |
| vCA3 | CD | UNEX | dSb | CD | UNEX | 0.0180 | >0.999 | 0.0001 | 0.9854 | 0.0002 | 0.6910 | 0.0001 | 0.9995 |
| vCA3 | CD | UNEX | EC | CD | UNEX | 0.0568 | 0.0679 | 0.0000 | >0.999 | 0.0001 | >0.999 | 0.0000 | >0.999 |
| vCA3 | CD | UNEX | vDG | CD | UNEX | 0.0479 | 0.3432 | 0.0000 | >0.999 | 0.0000 | >0.999 | 0.0000 | >0.999 |
| vCA3 | CD | UNEX | vSb | CD | UNEX | 0.0312 | 0.9900 | 0.0001 | 0.9819 | 0.0002 | 0.6357 | 0.0001 | 0.9995 |
| vCA3 | CD | UNEX | EcrhC | CD | UNEX | 0.0523 | 0.1673 | 0.0001 | >0.999 | 0.0001 | 0.9985 | 0.0000 | >0.999 |
| vCA3 | CD | UNEX | dDG | CD | EXP | 0.0479 | 0.2632 | 0.0000 | >0.999 | 0.0000 | >0.999 | 0.0000 | >0.999 |
| vCA3 | CD | UNEX | dSb | CD | EXP | 0.0168 | >0.999 | 0.0001 | 0.9974 | 0.0001 | 0.8097 | 0.0001 | >0.999 |
| vCA3 | CD | UNEX | EC | CD | EXP | 0.0550 | 0.0637 | 0.0001 | >0.999 | 0.0001 | 0.9976 | 0.0000 | >0.999 |
| vCA3 | CD | UNEX | vDG | CD | EXP | 0.0533 | 0.0925 | 0.0000 | >0.999 | 0.0000 | >0.999 | 0.0000 | >0.999 |
| vCA3 | CD | UNEX | vSb | CD | EXP | 0.0250 | 0.9997 | 0.0001 | 0.9425 | 0.0002 | 0.4879 | 0.0001 | 0.9960 |
| vCA3 | CD | UNEX | EcrhC | CD | EXP | 0.0408 | 0.6507 | 0.0001 | >0.999 | 0.0001 | 0.9935 | 0.0000 | >0.999 |
| vCA3 | CD | UNEX | dDG | WD | UNEX | 0.0241 | >0.999 | 0.0001 | >0.999 | 0.0001 | 0.9581 | 0.0001 | >0.999 |
| vCA3 | CD | UNEX | dSb | WD | UNEX | -0.0270 | 0.9994 | 0.0002 | 0.2400 | 0.0002 | **0.0461** | 0.0002 | 0.4700 |
| vCA3 | CD | UNEX | EC | WD | UNEX | 0.0449 | 0.5047 | 0.0001 | >0.999 | 0.0001 | 0.9242 | 0.0001 | >0.999 |
| vCA3 | CD | UNEX | vCA3 | WD | UNEX | -0.0037 | >0.999 | 0.0000 | >0.999 | 0.0000 | >0.999 | 0.0000 | >0.999 |
| vCA3 | CD | UNEX | vDG | WD | UNEX | 0.0307 | 0.9922 | 0.0000 | >0.999 | 0.0001 | >0.999 | 0.0000 | >0.999 |
| vCA3 | CD | UNEX | vSb | WD | UNEX | 0.0010 | >0.999 | 0.0002 | 0.3358 | 0.0002 | 0.0501 | 0.0001 | 0.6471 |
| vCA3 | CD | UNEX | EcrhC | WD | UNEX | 0.0103 | >0.999 | 0.0001 | 0.9279 | 0.0002 | 0.4499 | 0.0001 | 0.9947 |
| vCA3 | CD | UNEX | vCA3 | CD | EXP | 0.0138 | >0.999 | 0.0000 | >0.999 | 0.0000 | >0.999 | 0.0000 | >0.999 |
| vCA3 | CD | UNEX | dDG | WD | EXP | 0.0414 | 0.6136 | 0.0000 | >0.999 | 0.0001 | >0.999 | 0.0000 | >0.999 |
| vCA3 | CD | UNEX | dSb | WD | EXP | 0.0145 | >0.999 | 0.0001 | 0.9361 | 0.0002 | 0.5108 | 0.0001 | 0.9942 |
| vCA3 | CD | UNEX | EC | WD | EXP | 0.0564 | **0.0456** | 0.0000 | >0.999 | 0.0001 | >0.999 | 0.0000 | >0.999 |
| vCA3 | CD | UNEX | vCA3 | WD | EXP | 0.0013 | >0.999 | 0.0000 | >0.999 | 0.0000 | >0.999 | 0.0000 | >0.999 |
| vCA3 | CD | UNEX | vDG | WD | EXP | 0.0380 | 0.7991 | 0.0000 | >0.999 | 0.0000 | >0.999 | 0.0000 | >0.999 |
| vCA3 | CD | UNEX | vSb | WD | EXP | 0.0224 | >0.999 | 0.0001 | 0.9997 | 0.0001 | 0.9744 | 0.0001 | >0.999 |
| vCA3 | CD | UNEX | EcrhC | WD | EXP | 0.0304 | 0.9881 | 0.0000 | >0.999 | 0.0001 | >0.999 | 0.0000 | >0.999 |
| vCA3 | CD | EXP | dDG | CD | UNEX | 0.0315 | 0.9784 | 0.0000 | >0.999 | 0.0001 | >0.999 | 0.0000 | >0.999 |
| vCA3 | CD | EXP | dSb | CD | UNEX | 0.0042 | >0.999 | 0.0001 | 0.8039 | 0.0002 | 0.4618 | 0.0001 | 0.9303 |
| vCA3 | CD | EXP | EC | CD | UNEX | 0.0430 | 0.5204 | 0.0001 | >0.999 | 0.0001 | 0.9998 | 0.0000 | >0.999 |
| vCA3 | CD | EXP | EcrhC | CD | UNEX | 0.0385 | 0.7748 | 0.0001 | >0.999 | 0.0001 | 0.9872 | 0.0001 | >0.999 |
| vCA3 | CD | EXP | vDG | CD | UNEX | 0.0341 | 0.9364 | 0.0000 | >0.999 | 0.0000 | >0.999 | 0.0000 | >0.999 |
| vCA3 | CD | EXP | vSb | CD | UNEX | 0.0174 | >0.999 | 0.0001 | 0.7832 | 0.0002 | 0.4054 | 0.0001 | 0.9266 |
| vCA3 | CD | EXP | EC | CD | UNEX | 0.0311 | 0.9825 | 0.0001 | 0.9944 | 0.0001 | 0.7845 | 0.0001 | >0.999 |
| vCA3 | CD | EXP | dDG | CD | EXP | 0.0341 | 0.8999 | 0.0000 | >0.999 | 0.0001 | >0.999 | 0.0000 | >0.999 |
| vCA3 | CD | EXP | dSb | CD | EXP | 0.0030 | >0.999 | 0.0001 | 0.9070 | 0.0001 | 0.5904 | 0.0001 | 0.9803 |
| vCA3 | CD | EXP | EC | CD | EXP | 0.0412 | 0.5263 | 0.0001 | >0.999 | 0.0001 | 0.9798 | 0.0001 | >0.999 |
| vCA3 | CD | EXP | EcrhC | CD | EXP | 0.0270 | 0.9969 | 0.0001 | 0.9993 | 0.0001 | 0.9591 | 0.0001 | >0.999 |
| vCA3 | CD | EXP | vDG | CD | EXP | 0.0395 | 0.6301 | 0.0000 | >0.999 | 0.0000 | >0.999 | 0.0000 | >0.999 |
| vCA3 | CD | EXP | vSb | CD | EXP | 0.0112 | >0.999 | 0.0001 | 0.6116 | 0.0002 | 0.2648 | 0.0001 | 0.8097 |
| vCA3 | CD | EXP | dDG | WD | UNEX | 0.0103 | >0.999 | 0.0001 | 0.9856 | 0.0001 | 0.8567 | 0.0001 | 0.9982 |
| vCA3 | CD | EXP | dSb | WD | UNEX | -0.0408 | 0.6513 | 0.0002 | **0.0479** | 0.0002 | **0.0146** | 0.0002 | 0.0931 |
| vCA3 | CD | EXP | EcrhC | WD | UNEX | -0.0035 | >0.999 | 0.0001 | 0.5938 | 0.0002 | 0.2456 | 0.0001 | 0.8074 |
| vCA3 | CD | EXP | vCA3 | WD | UNEX | -0.0175 | >0.999 | 0.0000 | >0.999 | 0.0000 | >0.999 | 0.0000 | >0.999 |
| vCA3 | CD | EXP | vDG | WD | UNEX | 0.0169 | >0.999 | 0.0000 | >0.999 | 0.0001 | >0.999 | 0.0000 | >0.999 |
| vCA3 | CD | EXP | vSb | WD | UNEX | -0.0128 | >0.999 | 0.0002 | 0.0789 | 0.0002 | **0.0161** | 0.0002 | 0.1744 |
| vCA3 | CD | EXP | EC | WD | EXP | 0.0426 | 0.4401 | 0.0000 | >0.999 | 0.0001 | >0.999 | 0.0000 | >0.999 |
| vCA3 | CD | EXP | dDG | WD | EXP | 0.0276 | 0.9953 | 0.0001 | >0.999 | 0.0001 | >0.999 | 0.0000 | >0.999 |
| vCA3 | CD | EXP | dSb | WD | EXP | 0.0008 | >0.999 | 0.0001 | 0.5938 | 0.0002 | 0.2830 | 0.0001 | 0.7804 |
| vCA3 | CD | EXP | EcrhC | WD | EXP | 0.0166 | >0.999 | 0.0000 | >0.999 | 0.0001 | >0.999 | 0.0000 | >0.999 |
| vCA3 | CD | EXP | vCA3 | WD | EXP | -0.0125 | >0.999 | 0.0000 | >0.999 | 0.0000 | >0.999 | 0.0000 | >0.999 |
| vCA3 | CD | EXP | vDG | WD | EXP | 0.0242 | 0.9997 | 0.0000 | >0.999 | 0.0001 | >0.999 | 0.0000 | >0.999 |
| vCA3 | CD | EXP | vSb | WD | EXP | 0.0086 | >0.999 | 0.0001 | 0.9699 | 0.0001 | 0.8922 | 0.0001 | 0.9888 |
| dDG | CD | UNEX | dSb | CD | UNEX | -0.0273 | 0.9992 | 0.0001 | 0.9996 | 0.0001 | 0.9995 | 0.0001 | 0.9997 |
| dDG | CD | UNEX | EcrhC | CD | UNEX | 0.0070 | >0.999 | 0.0000 | >0.999 | 0.0000 | >0.999 | 0.0000 | >0.999 |
| dDG | CD | UNEX | vDG | CD | UNEX | 0.0026 | >0.999 | 0.0000 | >0.999 | 0.0000 | >0.999 | 0.0000 | >0.999 |
| dDG | CD | UNEX | vSb | CD | UNEX | -0.0142 | >0.999 | 0.0001 | 0.9994 | 0.0001 | 0.9989 | 0.0001 | 0.9996 |
| dDG | CD | UNEX | EC | CD | UNEX | 0.0115 | >0.999 | 0.0000 | >0.999 | 0.0000 | >0.999 | 0.0000 | >0.999 |
| dDG | CD | UNEX | EC | CD | UNEX | -0.0004 | >0.999 | 0.0001 | >0.999 | 0.0001 | >0.999 | 0.0001 | >0.999 |
| dDG | CD | UNEX | EC | CD | EXP | 0.0097 | >0.999 | 0.0000 | >0.999 | 0.0000 | >0.999 | 0.0000 | >0.999 |
| dDG | CD | UNEX | dDG | CD | EXP | 0.0025 | >0.999 | 0.0000 | >0.999 | 0.0000 | >0.999 | 0.0000 | >0.999 |
| dDG | CD | UNEX | dSb | CD | EXP | -0.0285 | 0.9959 | 0.0001 | >0.999 | 0.0001 | >0.999 | 0.0001 | >0.999 |
| dDG | CD | UNEX | EcrhC | CD | EXP | -0.0045 | >0.999 | 0.0000 | >0.999 | 0.0001 | >0.999 | 0.0000 | >0.999 |
| dDG | CD | UNEX | vDG | CD | EXP | 0.0080 | >0.999 | 0.0000 | >0.999 | 0.0000 | >0.999 | 0.0000 | >0.999 |
| dDG | CD | UNEX | vSb | CD | EXP | -0.0204 | >0.999 | 0.0001 | 0.9965 | 0.0001 | 0.9955 | 0.0001 | 0.9971 |
| dDG | CD | UNEX | dDG | WD | UNEX | -0.0212 | >0.999 | 0.0001 | >0.999 | 0.0001 | >0.999 | 0.0001 | >0.999 |
| dDG | CD | UNEX | dSb | WD | UNEX | -0.0723 | **0.0011** | 0.0002 | 0.5164 | 0.0002 | 0.5762 | 0.0002 | 0.4978 |
| dDG | CD | UNEX | EcrhC | WD | UNEX | -0.0350 | 0.9446 | 0.0001 | 0.9941 | 0.0001 | 0.9909 | 0.0001 | 0.9961 |
| dDG | CD | UNEX | vDG | WD | UNEX | -0.0146 | >0.999 | 0.0000 | >0.999 | 0.0000 | >0.999 | 0.0000 | >0.999 |
| dDG | CD | UNEX | vSb | WD | UNEX | -0.0443 | 0.5391 | 0.0001 | 0.6402 | 0.0002 | 0.5970 | 0.0001 | 0.6745 |
| dDG | CD | UNEX | EC | WD | EXP | 0.0111 | >0.999 | 0.0000 | >0.999 | 0.0000 | >0.999 | 0.0000 | >0.999 |
| dDG | CD | UNEX | dDG | WD | EXP | -0.0039 | >0.999 | 0.0000 | >0.999 | 0.0000 | >0.999 | 0.0000 | >0.999 |
| dDG | CD | UNEX | dSb | WD | EXP | -0.0308 | 0.9851 | 0.0001 | 0.9958 | 0.0001 | 0.9965 | 0.0001 | 0.9957 |
| dDG | CD | UNEX | EcrhC | WD | EXP | -0.0150 | >0.999 | 0.0000 | >0.999 | 0.0000 | >0.999 | 0.0000 | >0.999 |
| dDG | CD | UNEX | vDG | WD | EXP | -0.0073 | >0.999 | 0.0000 | >0.999 | 0.0000 | >0.999 | 0.0000 | >0.999 |
| dDG | CD | UNEX | vSb | WD | EXP | -0.0229 | >0.999 | 0.0001 | >0.999 | 0.0001 | >0.999 | 0.0001 | >0.999 |
| dDG | CD | EXP | dSb | CD | UNEX | -0.0298 | 0.9910 | 0.0001 | 0.9956 | 0.0001 | 0.9956 | 0.0001 | 0.9960 |
| dDG | CD | EXP | dSb | CD | UNEX | -0.0748 | **0.0002** | 0.0002 | 0.3038 | 0.0002 | 0.3751 | 0.0002 | 0.2813 |
| dDG | CD | EXP | EC | CD | UNEX | 0.0089 | >0.999 | 0.0000 | >0.999 | 0.0000 | >0.999 | 0.0000 | >0.999 |
| dDG | CD | EXP | EcrhC | CD | UNEX | 0.0045 | >0.999 | 0.0000 | >0.999 | 0.0001 | >0.999 | 0.0000 | >0.999 |
| dDG | CD | EXP | vDG | CD | UNEX | 0.0001 | >0.999 | 0.0000 | >0.999 | 0.0000 | >0.999 | 0.0000 | >0.999 |
| dDG | CD | EXP | vSb | CD | UNEX | -0.0167 | >0.999 | 0.0001 | 0.9943 | 0.0001 | 0.9920 | 0.0001 | 0.9956 |
| dDG | CD | EXP | dSb | CD | EXP | -0.0311 | 0.9689 | 0.0001 | 0.9995 | 0.0001 | 0.9994 | 0.0001 | 0.9997 |
| dDG | CD | EXP | EC | CD | EXP | 0.0072 | >0.999 | 0.0000 | >0.999 | 0.0001 | >0.999 | 0.0000 | >0.999 |
| dDG | CD | EXP | EcrhC | CD | EXP | -0.0071 | >0.999 | 0.0001 | >0.999 | 0.0001 | >0.999 | 0.0001 | >0.999 |
| dDG | CD | EXP | vDG | CD | EXP | 0.0055 | >0.999 | 0.0000 | >0.999 | 0.0000 | >0.999 | 0.0000 | >0.999 |
| dDG | CD | EXP | vSb | CD | EXP | -0.0229 | >0.999 | 0.0001 | 0.9752 | 0.0001 | 0.9751 | 0.0001 | 0.9767 |
| dDG | CD | EXP | vSb | WD | UNEX | -0.0468 | 0.3111 | 0.0002 | 0.4163 | 0.0002 | 0.3947 | 0.0002 | 0.4422 |
| dDG | CD | EXP | EC | WD | UNEX | -0.0030 | >0.999 | 0.0001 | >0.999 | 0.0001 | >0.999 | 0.0001 | >0.999 |
| dDG | CD | EXP | EcrhC | WD | UNEX | -0.0375 | 0.8204 | 0.0001 | 0.9664 | 0.0001 | 0.9601 | 0.0001 | 0.9734 |
| dDG | CD | EXP | vDG | WD | UNEX | -0.0171 | >0.999 | 0.0000 | >0.999 | 0.0000 | >0.999 | 0.0000 | >0.999 |
| dDG | CD | EXP | dDG | WD | UNEX | -0.0237 | >0.999 | 0.0001 | >0.999 | 0.0001 | >0.999 | 0.0001 | >0.999 |
| dDG | CD | EXP | dDG | WD | EXP | -0.0064 | >0.999 | 0.0000 | >0.999 | 0.0000 | >0.999 | 0.0000 | >0.999 |
| dDG | CD | EXP | dSb | WD | EXP | -0.0333 | 0.9226 | 0.0001 | 0.9716 | 0.0001 | 0.9794 | 0.0001 | 0.9691 |
| dDG | CD | EXP | EC | WD | EXP | 0.0086 | >0.999 | 0.0000 | >0.999 | 0.0000 | >0.999 | 0.0000 | >0.999 |
| dDG | CD | EXP | EcrhC | WD | EXP | -0.0175 | >0.999 | 0.0000 | >0.999 | 0.0000 | >0.999 | 0.0000 | >0.999 |
| dDG | CD | EXP | vDG | WD | EXP | -0.0099 | >0.999 | 0.0000 | >0.999 | 0.0000 | >0.999 | 0.0000 | >0.999 |
| dDG | CD | EXP | vSb | WD | EXP | -0.0254 | 0.9991 | 0.0001 | >0.999 | 0.0001 | >0.999 | 0.0001 | 0.9999 |
| vDG | CD | UNEX | dSb | CD | UNEX | -0.0299 | 0.9952 | 0.0001 | 0.8484 | 0.0001 | 0.8183 | 0.0001 | 0.8609 |
| vDG | CD | UNEX | EC | CD | UNEX | 0.0089 | >0.999 | 0.0001 | >0.999 | 0.0001 | >0.999 | 0.0001 | >0.999 |
| vDG | CD | UNEX | EcrhC | CD | UNEX | 0.0044 | >0.999 | 0.0001 | >0.999 | 0.0001 | 0.9998 | 0.0001 | >0.999 |
| vDG | CD | UNEX | vSb | CD | UNEX | -0.0167 | >0.999 | 0.0001 | 0.8309 | 0.0001 | 0.7727 | 0.0001 | 0.8553 |
| vDG | CD | UNEX | dSb | CD | EXP | -0.0311 | 0.9824 | 0.0001 | 0.9341 | 0.0001 | 0.9080 | 0.0001 | 0.9457 |
| vDG | CD | UNEX | EC | CD | EXP | 0.0071 | >0.999 | 0.0001 | >0.999 | 0.0001 | 0.9997 | 0.0001 | >0.999 |
| vDG | CD | UNEX | EcrhC | CD | EXP | -0.0071 | >0.999 | 0.0001 | 0.9996 | 0.0001 | 0.9989 | 0.0001 | 0.9998 |
| vDG | CD | UNEX | vSb | CD | EXP | -0.0230 | >0.999 | 0.0001 | 0.6843 | 0.0002 | 0.6412 | 0.0001 | 0.7027 |
| vDG | CD | UNEX | vDG | CD | EXP | 0.0054 | >0.999 | 0.0000 | >0.999 | 0.0000 | >0.999 | 0.0000 | >0.999 |
| vDG | CD | UNEX | dSb | WD | UNEX | -0.0749 | **0.0005** | 0.0002 | 0.0708 | 0.0002 | 0.0818 | 0.0002 | 0.0666 |
| vDG | CD | UNEX | EC | WD | UNEX | -0.0030 | >0.999 | 0.0001 | 0.9964 | 0.0001 | 0.9718 | 0.0001 | 0.9991 |
| vDG | CD | UNEX | EcrhC | WD | UNEX | -0.0376 | 0.8736 | 0.0001 | 0.6618 | 0.0002 | 0.5970 | 0.0001 | 0.7013 |
| vDG | CD | UNEX | vDG | WD | UNEX | -0.0172 | >0.999 | 0.0000 | >0.999 | 0.0001 | >0.999 | 0.0000 | >0.999 |
| vDG | CD | UNEX | vSb | WD | UNEX | -0.0469 | 0.3962 | 0.0002 | 0.1114 | 0.0002 | 0.0884 | 0.0002 | 0.1270 |
| vDG | CD | UNEX | dSb | WD | EXP | -0.0334 | 0.9522 | 0.0001 | 0.6679 | 0.0001 | 0.6637 | 0.0001 | 0.6693 |
| vDG | CD | UNEX | EC | WD | EXP | 0.0085 | >0.999 | 0.0000 | >0.999 | 0.0001 | >0.999 | 0.0000 | >0.999 |
| vDG | CD | UNEX | EcrhC | WD | EXP | -0.0175 | >0.999 | 0.0000 | >0.999 | 0.0001 | >0.999 | 0.0000 | >0.999 |
| vDG | CD | UNEX | vDG | WD | EXP | -0.0099 | >0.999 | 0.0000 | >0.999 | 0.0000 | >0.999 | 0.0001 | >0.999 |
| vDG | CD | UNEX | vSb | WD | EXP | -0.0255 | 0.9996 | 0.0001 | 0.9801 | 0.0001 | 0.9935 | 0.0001 | 0.9645 |
| vDG | CD | EXP | dSb | CD | UNEX | -0.0353 | 0.9042 | 0.0001 | 0.7445 | 0.0001 | 0.7886 | 0.0001 | 0.7304 |
| vDG | CD | EXP | EC | CD | UNEX | 0.0035 | >0.999 | 0.0001 | >0.999 | 0.0001 | >0.999 | 0.0001 | >0.999 |
| vDG | CD | EXP | EcrhC | CD | UNEX | -0.0010 | >0.999 | 0.0001 | 0.9998 | 0.0001 | 0.9998 | 0.0001 | 0.9999 |
| vDG | CD | EXP | vSb | CD | UNEX | -0.0222 | >0.999 | 0.0001 | 0.7213 | 0.0001 | 0.7379 | 0.0001 | 0.7224 |
| vDG | CD | EXP | dSb | WD | UNEX | -0.0803 | **<0.0001** | 0.0002 | **0.0359** | 0.0002 | 0.0617 | 0.0002 | **0.0288** |
| vDG | CD | EXP | EC | WD | UNEX | -0.0084 | >0.999 | 0.0001 | 0.9889 | 0.0001 | 0.9659 | 0.0001 | 0.9952 |
| vDG | CD | EXP | EcrhC | WD | UNEX | -0.0430 | 0.5195 | 0.0001 | 0.5225 | 0.0002 | 0.5497 | 0.0001 | 0.5304 |
| vDG | CD | EXP | vDG | WD | UNEX | -0.0226 | >0.999 | 0.0001 | >0.999 | 0.0001 | >0.999 | 0.0001 | >0.999 |
| vDG | CD | EXP | vSb | WD | UNEX | -0.0523 | 0.1152 | 0.0002 | 0.0602 | 0.0002 | 0.0671 | 0.0002 | 0.0612 |
| vDG | CD | EXP | dSb | CD | EXP | -0.0365 | 0.7983 | 0.0001 | 0.8646 | 0.0001 | 0.8882 | 0.0001 | 0.8618 |
| vDG | CD | EXP | EC | CD | EXP | 0.0017 | >0.999 | 0.0001 | >0.999 | 0.0001 | 0.9996 | 0.0001 | >0.999 |
| vDG | CD | EXP | EcrhC | CD | EXP | -0.0125 | >0.999 | 0.0001 | 0.9983 | 0.0001 | 0.9986 | 0.0001 | 0.9982 |
| vDG | CD | EXP | vSb | CD | EXP | -0.0284 | 0.9925 | 0.0001 | 0.5375 | 0.0001 | 0.5904 | 0.0001 | 0.5212 |
| vDG | CD | EXP | dSb | WD | EXP | -0.0388 | 0.6753 | 0.0001 | 0.5197 | 0.0001 | 0.6145 | 0.0001 | 0.4849 |
| vDG | CD | EXP | EC | WD | EXP | 0.0031 | >0.999 | 0.0000 | >0.999 | 0.0001 | >0.999 | 0.0000 | >0.999 |
| vDG | CD | EXP | EcrhC | WD | EXP | -0.0230 | >0.999 | 0.0001 | >0.999 | 0.0000 | >0.999 | 0.0001 | >0.999 |
| vDG | CD | EXP | vDG | WD | EXP | -0.0153 | >0.999 | 0.0000 | >0.999 | 0.0000 | >0.999 | 0.0001 | >0.999 |
| vDG | CD | EXP | vSb | WD | EXP | -0.0309 | 0.9711 | 0.0001 | 0.9495 | 0.0001 | 0.9918 | 0.0001 | 0.9003 |
| EcrhC | CD | UNEX | dSb | CD | UNEX | -0.0343 | 0.9579 | 0.0001 | >0.999 | 0.0000 | >0.999 | 0.0001 | >0.999 |
| EcrhC | CD | UNEX | EC | CD | UNEX | 0.0045 | >0.999 | 0.0000 | >0.999 | 0.0000 | >0.999 | 0.0000 | >0.999 |
| EcrhC | CD | UNEX | vSb | CD | UNEX | -0.0211 | >0.999 | 0.0001 | >0.999 | 0.0001 | >0.999 | 0.0001 | >0.999 |
| EcrhC | CD | UNEX | dSb | CD | EXP | -0.0355 | 0.8981 | 0.0000 | >0.999 | 0.0000 | >0.999 | 0.0000 | >0.999 |
| EcrhC | CD | UNEX | EC | CD | EXP | 0.0027 | >0.999 | 0.0000 | >0.999 | 0.0000 | >0.999 | 0.0000 | >0.999 |
| EcrhC | CD | UNEX | EcrhC | CD | EXP | -0.0115 | >0.999 | 0.0000 | >0.999 | 0.0000 | >0.999 | 0.0000 | >0.999 |
| EcrhC | CD | UNEX | vSb | CD | EXP | -0.0274 | 0.9982 | 0.0001 | >0.999 | 0.0001 | >0.999 | 0.0001 | >0.999 |
| EcrhC | CD | UNEX | dSb | WD | UNEX | -0.0793 | **0.0001** | 0.0001 | 0.9758 | 0.0001 | 0.9934 | 0.0001 | 0.9595 |
| EcrhC | CD | UNEX | EC | WD | UNEX | -0.0074 | >0.999 | 0.0000 | >0.999 | 0.0000 | >0.999 | 0.0000 | >0.999 |
| EcrhC | CD | UNEX | EcrhC | WD | UNEX | -0.0420 | 0.6697 | 0.0001 | >0.999 | 0.0001 | >0.999 | 0.0001 | >0.999 |
| EcrhC | CD | UNEX | vSb | WD | UNEX | -0.0513 | 0.2016 | 0.0001 | 0.9918 | 0.0001 | 0.9947 | 0.0001 | 0.9905 |
| EcrhC | CD | UNEX | dSb | WD | EXP | -0.0378 | 0.8106 | 0.0001 | >0.999 | 0.0001 | >0.999 | 0.0001 | >0.999 |
| EcrhC | CD | UNEX | EC | WD | EXP | 0.0041 | >0.999 | 0.0000 | >0.999 | 0.0000 | >0.999 | 0.0000 | >0.999 |
| EcrhC | CD | UNEX | EcrhC | WD | EXP | -0.0219 | >0.999 | 0.0000 | >0.999 | 0.0000 | >0.999 | 0.0000 | >0.999 |
| EcrhC | CD | UNEX | vSb | WD | EXP | -0.0299 | 0.9908 | 0.0000 | >0.999 | 0.0000 | >0.999 | 0.0000 | >0.999 |
| EcrhC | CD | EXP | dSb | CD | UNEX | -0.0228 | >0.999 | 0.0000 | >0.999 | 0.0000 | >0.999 | 0.0000 | >0.999 |
| EcrhC | CD | EXP | EC | CD | UNEX | 0.0160 | >0.999 | 0.0000 | >0.999 | 0.0000 | >0.999 | 0.0000 | >0.999 |
| EcrhC | CD | EXP | vSb | CD | UNEX | -0.0096 | >0.999 | 0.0000 | >0.999 | 0.0000 | >0.999 | 0.0000 | >0.999 |
| EcrhC | CD | EXP | dSb | CD | EXP | -0.0240 | 0.9997 | 0.0000 | >0.999 | 0.0000 | >0.999 | 0.0000 | >0.999 |
| EcrhC | CD | EXP | EC | CD | EXP | 0.0142 | >0.999 | 0.0000 | >0.999 | 0.0000 | >0.999 | 0.0000 | >0.999 |
| EcrhC | CD | EXP | vSb | CD | EXP | -0.0158 | >0.999 | 0.0001 | >0.999 | 0.0001 | >0.999 | 0.0001 | >0.999 |
| EcrhC | CD | EXP | Ecrh | WD | UNEX | -0.0305 | 0.9873 | 0.0001 | >0.999 | 0.0001 | >0.999 | 0.0001 | >0.999 |
| EcrhC | CD | EXP | dSb | WD | UNEX | -0.0678 | **0.0020** | 0.0001 | 0.9814 | 0.0001 | 0.9937 | 0.0001 | 0.9728 |
| EcrhC | CD | EXP | EC | WD | UNEX | 0.0041 | >0.999 | 0.0000 | >0.999 | 0.0000 | >0.999 | 0.0000 | >0.999 |
| EcrhC | CD | EXP | vSb | WD | UNEX | -0.0397 | 0.7098 | 0.0001 | 0.9944 | 0.0001 | 0.9950 | 0.0001 | 0.9948 |
| EcrhC | CD | EXP | Ecrh | WD | EXP | -0.0104 | >0.999 | 0.0000 | >0.999 | 0.0000 | >0.999 | 0.0000 | >0.999 |
| EcrhC | CD | EXP | dSb | WD | EXP | -0.0262 | 0.9982 | 0.0001 | >0.999 | 0.0001 | >0.999 | 0.0001 | >0.999 |
| EcrhC | CD | EXP | EC | WD | EXP | 0.0156 | >0.999 | 0.0000 | >0.999 | 0.0000 | >0.999 | -0.0001 | >0.999 |
| EcrhC | CD | EXP | vSb | WD | EXP | -0.0184 | >0.999 | 0.0000 | >0.999 | 0.0000 | >0.999 | 0.0000 | >0.999 |
| EC | CD | UNEX | dSb | CD | UNEX | -0.0388 | 0.8281 | 0.0001 | >0.999 | 0.0001 | >0.999 | 0.0001 | >0.999 |
| EC | CD | UNEX | vSb | CD | UNEX | -0.0256 | 0.9998 | 0.0001 | >0.999 | 0.0001 | >0.999 | 0.0001 | >0.999 |
| EC | CD | UNEX | dSb | CD | EXP | -0.0400 | 0.6961 | 0.0001 | >0.999 | 0.0001 | >0.999 | 0.0001 | >0.999 |
| EC | CD | UNEX | EC | CD | EXP | -0.0018 | >0.999 | 0.0000 | >0.999 | 0.0000 | >0.999 | 0.0000 | >0.999 |
| EC | CD | UNEX | vSb | CD | EXP | -0.0318 | 0.9751 | 0.0001 | >0.999 | 0.0001 | >0.999 | 0.0001 | 0.9998 |
| EC | CD | UNEX | dSb | WD | UNEX | -0.0838 | **<0.0001** | 0.0001 | 0.7725 | 0.0001 | 0.9109 | 0.0001 | 0.6853 |
| EC | CD | UNEX | EC | WD | UNEX | -0.0119 | >0.999 | 0.0000 | >0.999 | 0.0001 | >0.999 | 0.0000 | >0.999 |
| EC | CD | UNEX | vSb | WD | UNEX | -0.0557 | 0.0849 | 0.0001 | 0.8648 | 0.0001 | 0.9207 | 0.0001 | 0.8358 |
| EC | CD | UNEX | dSb | WD | EXP | -0.0422 | 0.5651 | 0.0001 | >0.999 | 0.0001 | >0.999 | 0.0001 | 0.9996 |
| EC | CD | UNEX | EC | WD | EXP | -0.0004 | >0.999 | 0.0000 | >0.999 | 0.0000 | >0.999 | 0.0000 | >0.999 |
| EC | CD | UNEX | vSb | WD | EXP | -0.0344 | 0.9304 | 0.0001 | >0.999 | 0.0000 | >0.999 | 0.0001 | >0.999 |
| EC | CD | EXP | dSb | CD | UNEX | -0.0370 | 0.8438 | 0.0001 | >0.999 | 0.0001 | >0.999 | 0.0001 | >0.999 |
| EC | CD | EXP | vSb | CD | UNEX | -0.0238 | >0.999 | 0.0001 | >0.999 | 0.0001 | >0.999 | 0.0001 | >0.999 |
| EC | CD | EXP | dSb | CD | EXP | -0.0382 | 0.7084 | 0.0000 | >0.999 | 0.0000 | >0.999 | 0.0000 | >0.999 |
| EC | CD | EXP | vSb | CD | EXP | -0.0301 | 0.9809 | 0.0001 | >0.999 | 0.0001 | >0.999 | 0.0001 | >0.999 |
| EC | CD | EXP | dSb | WD | UNEX | -0.0820 | **<0.0001** | 0.0001 | 0.9152 | 0.0001 | 0.9854 | 0.0001 | 0.8423 |
| EC | CD | EXP | EC | WD | UNEX | -0.0101 | >0.999 | 0.0000 | >0.999 | 0.0000 | >0.999 | 0.0000 | >0.999 |
| EC | CD | EXP | vSb | WD | UNEX | -0.0540 | 0.0805 | 0.0001 | 0.9624 | 0.0001 | 0.9879 | 0.0001 | 0.9409 |
| EC | CD | EXP | dSb | WD | EXP | -0.0405 | 0.5728 | 0.0001 | >0.999 | 0.0001 | >0.999 | 0.0001 | >0.999 |
| EC | CD | EXP | EC | WD | EXP | 0.0014 | >0.999 | 0.0000 | >0.999 | 0.0000 | >0.999 | 0.0000 | >0.999 |
| EC | CD | EXP | vSb | WD | EXP | -0.0326 | 0.9409 | 0.0000 | >0.999 | 0.0000 | >0.999 | 0.0000 | >0.999 |
| dSb | CD | UNEX | dSb | CD | EXP | -0.0012 | >0.999 | 0.0000 | >0.999 | 0.0000 | >0.999 | 0.0000 | >0.999 |
| dSb | CD | UNEX | vSb | CD | EXP | 0.0069 | >0.999 | 0.0000 | >0.999 | 0.0000 | >0.999 | 0.0000 | >0.999 |
| dSb | CD | UNEX | vSb | CD | UNEX | 0.0132 | >0.999 | 0.0000 | >0.999 | 0.0000 | >0.999 | 0.0000 | >0.999 |
| dSb | CD | UNEX | dSb | WD | EXP | -0.0035 | >0.999 | 0.0000 | >0.999 | 0.0000 | >0.999 | 0.0000 | >0.999 |
| dSb | CD | UNEX | dSb | WD | UNEX | -0.0450 | 0.4988 | 0.0001 | >0.999 | 0.0001 | >0.999 | 0.0001 | >0.999 |
| dSb | CD | UNEX | vSb | WD | EXP | 0.0044 | >0.999 | 0.0000 | >0.999 | 0.0000 | >0.999 | 0.0000 | >0.999 |
| dSb | CD | UNEX | vSb | WD | UNEX | -0.0170 | >0.999 | 0.0001 | >0.999 | 0.0001 | >0.999 | 0.0001 | >0.999 |
| dSb | CD | EXP | vSb | CD | EXP | 0.0082 | >0.999 | 0.0000 | >0.999 | 0.0000 | >0.999 | 0.0000 | >0.999 |
| dSb | CD | EXP | vSb | CD | UNEX | 0.0144 | >0.999 | 0.0000 | >0.999 | 0.0000 | >0.999 | 0.0000 | >0.999 |
| dSb | CD | EXP | dSb | WD | EXP | -0.0023 | >0.999 | 0.0000 | >0.999 | 0.0000 | >0.999 | 0.0000 | >0.999 |
| dSb | CD | EXP | dSb | WD | UNEX | -0.0438 | 0.4743 | 0.0001 | >0.999 | 0.0001 | >0.999 | 0.0001 | >0.999 |
| dSb | CD | EXP | vSb | WD | EXP | 0.0056 | >0.999 | 0.0000 | >0.999 | 0.0000 | >0.999 | 0.0000 | >0.999 |
| dSb | CD | EXP | vSb | WD | UNEX | -0.0158 | >0.999 | 0.0001 | >0.999 | 0.0001 | >0.999 | 0.0001 | >0.999 |
| vSb | CD | UNEX | vSb | CD | EXP | -0.0062 | >0.999 | 0.0000 | >0.999 | 0.0000 | >0.999 | 0.0000 | >0.999 |
| vSb | CD | UNEX | vSb | WD | UNEX | -0.0301 | 0.9945 | 0.0001 | >0.999 | 0.0001 | >0.999 | 0.0001 | >0.999 |
| vSb | CD | UNEX | vSb | WD | EXP | -0.0087 | >0.999 | 0.0000 | >0.999 | 0.0000 | >0.999 | 0.0000 | >0.999 |
| vSb | CD | EXP | vSb | WD | EXP | -0.0025 | >0.999 | 0.0000 | >0.999 | 0.0000 | >0.999 | 0.0000 | >0.999 |
| vSb | CD | EXP | vSb | WD | UNEX | -0.0239 | >0.999 | 0.0000 | >0.999 | 0.0001 | >0.999 | 0.0000 | >0.999 |

***Supplemental Table 10B*.** Statistical Information of NODDI neuroimaging parameters in hippocampal subfields

|  |  |  |  |  |  | **ODI** | | **ICVF** | | **ISOVF** | | **LJ** | |
| --- | --- | --- | --- | --- | --- | --- | --- | --- | --- | --- | --- | --- | --- |
| **R0I 1** | **Diet1** | **Exp1** | **R0I 2** | **Diet2** | **Exp 2** | **Mean Diff** | ***p*** | **Mean Diff** | ***p*** | **Mean Diff** | ***p*** | **Mean Diff** | ***p*** |
| dCA1 | CD | EXP | EcrhC | CD | EXP | 0.0552 | 0.7933 | na | na | -0.0461 | >0.999 | 0.0583 | >0.999 |
| dCA1 | CD | EXP | CA2 | CD | EXP | 0.0132 | >0.999 | na | na | -0.0913 | 0.4447 | -0.0988 | 0.9536 |
| dCA1 | CD | EXP | dCA3 | CD | EXP | 0.0801 | 0.0594 | na | na | -0.1090 | 0.1055 | -0.1780 | **0.0179** |
| dCA1 | CD | EXP | dDG | CD | EXP | 0.0520 | 0.8844 | na | na | -0.0815 | 0.7221 | 0.0476 | >0.999 |
| dCA1 | CD | EXP | dSb | CD | EXP | 0.0037 | >0.999 | na | na | -0.0583 | 0.9965 | 0.0800 | 0.9986 |
| dCA1 | CD | EXP | EC | CD | EXP | 0.0035 | >0.999 | na | na | -0.1160 | **0.0495** | 0.0104 | >0.999 |
| dCA1 | CD | EXP | vCA1 | CD | EXP | 0.0300 | >0.999 | na | na | -0.1080 | 0.1104 | 0.0206 | >0.999 |
| dCA1 | CD | EXP | vCA3 | CD | EXP | 0.1100 | **0.0001** | na | na | -0.1080 | 0.1103 | -0.1090 | 0.8537 |
| dCA1 | CD | EXP | vDG | CD | EXP | 0.0335 | >0.999 | na | na | -0.1600 | **<0.0001** | -0.1390 | 0.3176 |
| dCA1 | CD | EXP | vSb | CD | EXP | -0.0012 | >0.999 | na | na | -0.0850 | 0.6251 | 0.0183 | >0.999 |
| dCA1 | CD | EXP | EcrhC | CD | UNEX | 0.0566 | 0.8196 | na | na | -0.0546 | 0.9996 | 0.0830 | 0.9987 |
| dCA1 | CD | EXP | CA2 | CD | UNEX | 0.0234 | >0.999 | na | na | -0.0826 | 0.7764 | -0.0444 | >0.999 |
| dCA1 | CD | EXP | dCA3 | CD | UNEX | 0.0743 | 0.2059 | na | na | -0.0841 | 0.7406 | -0.1520 | 0.2070 |
| dCA1 | CD | EXP | dDG | CD | UNEX | 0.0507 | 0.9466 | na | na | -0.0699 | 0.9656 | 0.0678 | >0.999 |
| dCA1 | CD | EXP | dSb | CD | UNEX | -0.0003 | >0.999 | na | na | -0.0396 | >0.999 | 0.0927 | 0.9901 |
| dCA1 | CD | EXP | EC | CD | UNEX | 0.0041 | >0.999 | na | na | -0.1250 | **0.0285** | 0.0075 | >0.999 |
| dCA1 | CD | EXP | vCA1 | CD | UNEX | 0.0369 | 0.9998 | na | na | -0.0921 | 0.5243 | 0.0393 | >0.999 |
| dCA1 | CD | EXP | vCA3 | CD | UNEX | 0.1300 | **<0.0001** | na | na | -0.0776 | 0.8757 | -0.0666 | >0.999 |
| dCA1 | CD | EXP | vDG | CD | UNEX | 0.0428 | 0.9963 | na | na | -0.1490 | **0.0012** | -0.1160 | 0.8220 |
| dCA1 | CD | EXP | vSb | CD | UNEX | -0.0107 | >0.999 | na | na | -0.0887 | 0.6190 | 0.0310 | >0.999 |
| dCA1 | CD | UNEX | CA2 | CD | EXP | -0.0001 | >0.999 | 0.0701 | 0.9985 | -0.0998 | 0.3262 | -0.1230 | 0.7032 |
| dCA1 | CD | UNEX | dCA1 | CD | EXP | -0.0133 | >0.999 | -0.0306 | 0.9994 | -0.0085 | >0.999 | -0.0244 | >0.999 |
| dCA1 | CD | UNEX | dCA3 | CD | EXP | 0.0668 | 0.4431 | 0.1090 | 0.9246 | -0.1170 | 0.0700 | -0.2030 | **0.0038** |
| dCA1 | CD | UNEX | dDG | CD | EXP | 0.0387 | 0.9995 | 0.0590 | 0.9988 | -0.0899 | 0.5842 | 0.0233 | >0.999 |
| dCA1 | CD | UNEX | dSb | CD | EXP | -0.0096 | >0.999 | -0.0528 | 0.9990 | -0.0668 | 0.9822 | 0.0556 | >0.999 |
| dCA1 | CD | UNEX | EC | CD | EXP | -0.0098 | >0.999 | -0.0657 | 0.9986 | -0.1240 | **0.0323** | -0.0139 | >0.999 |
| dCA1 | CD | UNEX | EcrhC | CD | EXP | 0.0419 | 0.9975 | 0.0441 | 0.9992 | -0.0546 | 0.9996 | 0.0339 | >0.999 |
| dCA1 | CD | UNEX | vCA1 | CD | EXP | 0.0167 | >0.999 | -0.0863 | 0.9861 | -0.1170 | 0.0734 | -0.0038 | >0.999 |
| dCA1 | CD | UNEX | vCA3 | CD | EXP | 0.0967 | **0.0058** | 0.0225 | 0.9996 | -0.1170 | 0.0733 | -0.1330 | 0.5087 |
| dCA1 | CD | UNEX | vDG | CD | EXP | 0.0202 | >0.999 | -0.0425 | 0.9992 | -0.1690 | **<0.0001** | -0.1630 | 0.1049 |
| dCA1 | CD | UNEX | vSb | CD | EXP | -0.0145 | >0.999 | -0.1040 | 0.9503 | -0.0935 | 0.4870 | -0.0060 | >0.999 |
| dCA1 | CD | UNEX | CA2 | CD | UNEX | 0.0101 | >0.999 | 0.1080 | 0.9525 | -0.0910 | 0.6454 | -0.0688 | >0.999 |
| dCA1 | CD | UNEX | dCA3 | CD | UNEX | 0.0610 | 0.7504 | 0.1400 | 0.6875 | -0.0925 | 0.6058 | -0.1760 | 0.0630 |
| dCA1 | CD | UNEX | dDG | CD | UNEX | 0.0374 | >0.999 | 0.0828 | 0.9982 | -0.0783 | 0.9081 | 0.0435 | >0.999 |
| dCA1 | CD | UNEX | dSb | CD | UNEX | -0.0135 | >0.999 | -0.0268 | 0.9996 | -0.0481 | >0.999 | 0.0684 | >0.999 |
| dCA1 | CD | UNEX | EC | CD | UNEX | -0.0092 | >0.999 | -0.0452 | 0.9992 | -0.1340 | **0.0184** | -0.0168 | >0.999 |
| dCA1 | CD | UNEX | EcrhC | CD | UNEX | 0.0433 | 0.9977 | 0.0644 | 0.9987 | -0.0630 | 0.9965 | 0.0587 | >0.999 |
| dCA1 | CD | UNEX | vCA1 | CD | UNEX | 0.0236 | >0.999 | -0.0703 | 0.9986 | -0.1010 | 0.3956 | 0.0150 | >0.999 |
| dCA1 | CD | UNEX | vCA3 | CD | UNEX | 0.1160 | **0.0002** | 0.0696 | 0.9986 | -0.0861 | 0.7672 | -0.0910 | 0.9963 |
| dCA1 | CD | UNEX | vDG | CD | UNEX | 0.0295 | >0.999 | -0.0183 | 0.9997 | -0.1570 | **0.0008** | -0.1400 | 0.4767 |
| dCA1 | CD | UNEX | vSb | CD | UNEX | -0.0240 | >0.999 | -0.0969 | 0.9828 | -0.0972 | 0.4826 | 0.0067 | >0.999 |
| dCA1 | CD | EXP | EcrhC | WD | EXP | 0.0394 | 0.9984 | na | na | -0.0943 | 0.3661 | -0.0022 | >0.999 |
| dCA1 | CD | EXP | CA2 | WD | EXP | -0.0106 | >0.999 | na | na | -0.1040 | 0.1621 | -0.1070 | 0.8804 |
| dCA1 | CD | EXP | dCA1 | WD | EXP | 0.0080 | >0.999 | na | na | 0.0032 | >0.999 | 0.0040 | >0.999 |
| dCA1 | CD | EXP | dCA3 | WD | EXP | 0.0574 | 0.7159 | na | na | -0.1140 | 0.0591 | -0.1670 | **0.0488** |
| dCA1 | CD | EXP | dDG | WD | EXP | 0.0393 | 0.9984 | na | na | -0.0784 | 0.7969 | 0.0646 | >0.999 |
| dCA1 | CD | EXP | dSb | WD | EXP | -0.0023 | >0.999 | na | na | -0.0352 | >0.999 | 0.1030 | 0.9201 |
| dCA1 | CD | EXP | EC | WD | EXP | -0.0009 | >0.999 | na | na | -0.1350 | **0.0042** | -0.0153 | >0.999 |
| dCA1 | CD | EXP | vCA1 | WD | EXP | 0.0172 | >0.999 | na | na | -0.1160 | **0.0465** | -0.0061 | >0.999 |
| dCA1 | CD | EXP | vCA3 | WD | EXP | 0.0885 | **0.0136** | na | na | -0.1210 | **0.0262** | -0.1160 | 0.7394 |
| dCA1 | CD | EXP | vDG | WD | EXP | 0.0133 | >0.999 | na | na | -0.1680 | **<0.0001** | -0.1680 | **0.0454** |
| dCA1 | CD | EXP | vSb | WD | EXP | -0.0101 | >0.999 | na | na | -0.0912 | 0.4476 | 0.0321 | >0.999 |
| dCA1 | CD | EXP | EcrhC | WD | UNEX | 0.0156 | >0.999 | na | na | -0.0560 | 0.9993 | 0.1020 | 0.9601 |
| dCA1 | CD | EXP | CA2 | WD | UNEX | -0.0210 | >0.999 | na | na | -0.1040 | 0.2339 | -0.1370 | 0.4491 |
| dCA1 | CD | EXP | dCA1 | WD | UNEX | -0.0121 | >0.999 | na | na | 0.0039 | >0.999 | -0.0305 | >0.999 |
| dCA1 | CD | EXP | dCA3 | WD | UNEX | 0.0496 | 0.9604 | na | na | -0.1080 | 0.1662 | -0.1990 | **0.0054** |
| dCA1 | CD | EXP | dDG | WD | UNEX | 0.0058 | >0.999 | na | na | -0.0665 | 0.9836 | 0.0148 | >0.999 |
| dCA1 | CD | EXP | dSb | WD | UNEX | -0.0253 | >0.999 | na | na | -0.0355 | >0.999 | 0.0501 | >0.999 |
| dCA1 | CD | EXP | EC | WD | UNEX | -0.0250 | >0.999 | na | na | -0.1260 | **0.0262** | 0.0069 | >0.999 |
| dCA1 | CD | EXP | vCA1 | WD | UNEX | 0.0044 | >0.999 | na | na | -0.1160 | 0.0763 | -0.0630 | >0.999 |
| dCA1 | CD | EXP | vCA3 | WD | UNEX | 0.0780 | 0.1288 | na | na | -0.1290 | **0.0175** | -0.1290 | 0.5986 |
| dCA1 | CD | EXP | vDG | WD | UNEX | 0.0000 | >0.999 | na | na | -0.1550 | **0.0005** | -0.1860 | **0.0177** |
| dCA1 | CD | EXP | vSb | WD | UNEX | -0.0389 | 0.9995 | na | na | -0.0854 | 0.7068 | -0.0036 | >0.999 |
| dCA1 | CD | UNEX | CA2 | WD | EXP | -0.0239 | >0.999 | 0.0157 | 0.9997 | -0.1130 | 0.1093 | -0.1310 | 0.5493 |
| dCA1 | CD | UNEX | dCA1 | WD | EXP | -0.0053 | >0.999 | -0.0740 | 0.9985 | -0.0053 | >0.999 | -0.0204 | >0.999 |
| dCA1 | CD | UNEX | dCA3 | WD | EXP | 0.0441 | 0.9936 | 0.0460 | 0.9992 | -0.1230 | **0.0387** | -0.1910 | **0.0113** |
| dCA1 | CD | UNEX | dDG | WD | EXP | 0.0260 | >0.999 | -0.0037 | >0.999 | -0.0869 | 0.6669 | 0.0403 | >0.999 |
| dCA1 | CD | UNEX | dSb | WD | EXP | -0.0156 | >0.999 | -0.1080 | 0.9500 | -0.0437 | >0.999 | 0.0788 | 0.9996 |
| dCA1 | CD | UNEX | EC | WD | EXP | -0.0142 | >0.999 | -0.0826 | 0.9982 | -0.1430 | **0.0027** | -0.0397 | >0.999 |
| dCA1 | CD | UNEX | EcrhC | WD | EXP | 0.0261 | >0.999 | -0.0512 | 0.9991 | -0.1030 | 0.2621 | -0.0265 | >0.999 |
| dCA1 | CD | UNEX | vCA1 | WD | EXP | 0.0039 | >0.999 | -0.1520 | 0.5517 | -0.1250 | **0.0304** | -0.0304 | >0.999 |
| dCA1 | CD | UNEX | vCA3 | WD | EXP | 0.0753 | 0.1826 | -0.0548 | 0.9990 | -0.1290 | **0.0170** | -0.1410 | 0.3756 |
| dCA1 | CD | UNEX | vDG | WD | EXP | 0.0000 | >0.999 | -0.1210 | 0.8652 | -0.1770 | **<0.0001** | -0.1920 | **0.0105** |
| dCA1 | CD | UNEX | vSb | WD | EXP | -0.0234 | >0.999 | -0.1720 | 0.3587 | -0.0997 | 0.3286 | 0.0078 | >0.999 |
| dCA1 | CD | UNEX | CA2 | WD | UNEX | -0.0343 | >0.999 | -0.0434 | 0.9993 | -0.1130 | 0.1613 | -0.1610 | 0.1736 |
| dCA1 | CD | UNEX | dCA1 | WD | UNEX | -0.0254 | >0.999 | -0.1590 | 0.4827 | -0.0046 | >0.999 | -0.0548 | >0.999 |
| dCA1 | CD | UNEX | dCA3 | WD | UNEX | 0.0363 | >0.999 | -0.0039 | >0.999 | -0.1170 | 0.1122 | -0.2230 | **0.0011** |
| dCA1 | CD | UNEX | dDG | WD | UNEX | -0.0075 | >0.999 | -0.0764 | 0.9984 | -0.0749 | 0.9466 | -0.0096 | >0.999 |
| dCA1 | CD | UNEX | dSb | WD | UNEX | -0.0386 | 0.9998 | -0.1990 | 0.1773 | -0.0440 | >0.999 | 0.0258 | >0.999 |
| dCA1 | CD | UNEX | EC | WD | UNEX | -0.0383 | 0.9998 | -0.1650 | 0.4241 | -0.1340 | **0.0169** | -0.0174 | >0.999 |
| dCA1 | CD | UNEX | EcrhC | WD | UNEX | 0.0023 | >0.999 | -0.1070 | 0.9578 | -0.0645 | 0.9947 | 0.0773 | 0.9999 |
| dCA1 | CD | UNEX | vCA1 | WD | UNEX | -0.0089 | >0.999 | -0.1930 | 0.2093 | -0.1250 | 0.0501 | -0.0874 | 0.9983 |
| dCA1 | CD | UNEX | vCA3 | WD | UNEX | 0.0647 | 0.6176 | -0.0916 | 0.9852 | -0.1380 | **0.0113** | -0.1530 | 0.2675 |
| dCA1 | CD | UNEX | vDG | WD | UNEX | -0.0133 | >0.999 | -0.1700 | 0.3794 | -0.1630 | **0.0003** | -0.2100 | **0.0039** |
| dCA1 | CD | UNEX | vSb | WD | UNEX | -0.0522 | 0.9521 | -0.2460 | **0.0365** | -0.0939 | 0.5700 | -0.0279 | >0.999 |
| vCA1 | CD | UNEX | CA2 | CD | UNEX | -0.0135 | >0.999 | na | na | 0.0096 | >0.999 | -0.0838 | 0.9993 |
| vCA1 | CD | UNEX | dCA3 | CD | UNEX | 0.0374 | >0.999 | na | na | 0.0081 | >0.999 | -0.1910 | **0.0200** |
| vCA1 | CD | UNEX | dDG | CD | UNEX | 0.0138 | >0.999 | na | na | 0.0222 | >0.999 | 0.0285 | >0.999 |
| vCA1 | CD | UNEX | dSb | CD | UNEX | -0.0372 | >0.999 | na | na | 0.0525 | >0.999 | 0.0534 | >0.999 |
| vCA1 | CD | UNEX | EC | CD | UNEX | -0.0328 | >0.999 | na | na | -0.0331 | >0.999 | -0.0318 | >0.999 |
| vCA1 | CD | UNEX | EcrhC | CD | UNEX | 0.0197 | >0.999 | na | na | 0.0375 | >0.999 | 0.0437 | >0.999 |
| vCA1 | CD | UNEX | vCA3 | CD | UNEX | 0.0927 | **0.0225** | na | na | 0.0145 | >0.999 | -0.1060 | 0.9576 |
| vCA1 | CD | UNEX | vDG | CD | UNEX | 0.0059 | >0.999 | na | na | -0.0565 | 0.9996 | -0.1550 | 0.2398 |
| vCA1 | CD | UNEX | vSb | CD | UNEX | -0.0476 | 0.9875 | na | na | 0.0034 | >0.999 | -0.0083 | >0.999 |
| vCA1 | CD | UNEX | CA2 | CD | EXP | -0.0237 | >0.999 | na | na | 0.0008 | >0.999 | -0.1380 | 0.4209 |
| vCA1 | CD | UNEX | dCA3 | CD | EXP | 0.0432 | 0.9956 | na | na | -0.0165 | >0.999 | -0.2180 | **0.0008** |
| vCA1 | CD | UNEX | dDG | CD | EXP | 0.0151 | >0.999 | na | na | 0.0106 | >0.999 | 0.0083 | >0.999 |
| vCA1 | CD | UNEX | dSb | CD | EXP | -0.0332 | >0.999 | na | na | 0.0338 | >0.999 | 0.0406 | >0.999 |
| vCA1 | CD | UNEX | EC | CD | EXP | -0.0334 | >0.999 | na | na | -0.0235 | >0.999 | -0.0289 | >0.999 |
| vCA1 | CD | UNEX | EcrhC | CD | EXP | 0.0183 | >0.999 | na | na | 0.0460 | >0.999 | 0.0190 | >0.999 |
| vCA1 | CD | UNEX | vCA3 | CD | EXP | 0.0731 | 0.2350 | na | na | -0.0161 | >0.999 | -0.1480 | 0.2552 |
| vCA1 | CD | UNEX | vDG | CD | EXP | -0.0034 | >0.999 | na | na | -0.0679 | 0.9770 | -0.1780 | **0.0345** |
| vCA1 | CD | UNEX | vSb | CD | EXP | -0.0381 | 0.9997 | na | na | 0.0071 | >0.999 | -0.0210 | >0.999 |
| vCA1 | CD | UNEX | vCA1 | CD | EXP | -0.0069 | >0.999 | na | na | -0.0161 | >0.999 | -0.0187 | >0.999 |
| vCA1 | CD | UNEX | CA2 | WD | UNEX | -0.0579 | 0.8434 | na | na | -0.0121 | >0.999 | -0.1760 | 0.0655 |
| vCA1 | CD | UNEX | dCA3 | WD | UNEX | 0.0127 | >0.999 | na | na | -0.0161 | >0.999 | -0.2380 | **0.0002** |
| vCA1 | CD | UNEX | dDG | WD | UNEX | -0.0311 | >0.999 | na | na | 0.0257 | >0.999 | -0.0246 | >0.999 |
| vCA1 | CD | UNEX | dSb | WD | UNEX | -0.0622 | 0.7069 | na | na | 0.0566 | 0.9996 | 0.0108 | >0.999 |
| vCA1 | CD | UNEX | EC | WD | UNEX | -0.0619 | 0.7195 | na | na | -0.0338 | >0.999 | -0.0324 | >0.999 |
| vCA1 | CD | UNEX | EcrhC | WD | UNEX | -0.0213 | >0.999 | na | na | 0.0361 | >0.999 | 0.0623 | >0.999 |
| vCA1 | CD | UNEX | vCA3 | WD | UNEX | 0.0411 | 0.9992 | na | na | -0.0371 | >0.999 | -0.1680 | 0.1117 |
| vCA1 | CD | UNEX | vDG | WD | UNEX | -0.0369 | >0.999 | na | na | -0.0624 | 0.9971 | -0.2250 | **0.0009** |
| vCA1 | CD | UNEX | vSb | WD | UNEX | -0.0758 | 0.2376 | na | na | 0.0067 | >0.999 | -0.0429 | >0.999 |
| vCA1 | CD | UNEX | vCA1 | WD | UNEX | -0.0325 | >0.999 | na | na | -0.0242 | >0.999 | -0.1020 | 0.9739 |
| vCA1 | CD | UNEX | CA2 | WD | EXP | -0.0475 | 0.9783 | na | na | -0.0121 | >0.999 | -0.1460 | 0.2855 |
| vCA1 | CD | UNEX | dCA3 | WD | EXP | 0.0205 | >0.999 | na | na | -0.0219 | >0.999 | -0.2060 | **0.0028** |
| vCA1 | CD | UNEX | dDG | WD | EXP | 0.0024 | >0.999 | na | na | 0.0137 | >0.999 | 0.0253 | >0.999 |
| vCA1 | CD | UNEX | dSb | WD | EXP | -0.0392 | 0.9993 | na | na | 0.0569 | 0.9990 | 0.0638 | >0.999 |
| vCA1 | CD | UNEX | EC | WD | EXP | -0.0378 | 0.9997 | na | na | -0.0426 | >0.999 | -0.0547 | >0.999 |
| vCA1 | CD | UNEX | EcrhC | WD | EXP | 0.0025 | >0.999 | na | na | -0.0022 | >0.999 | -0.0415 | >0.999 |
| vCA1 | CD | UNEX | vCA3 | WD | EXP | 0.0516 | 0.9328 | na | na | -0.0288 | >0.999 | -0.1560 | 0.1676 |
| vCA1 | CD | UNEX | vDG | WD | EXP | -0.0236 | >0.999 | na | na | -0.0763 | 0.8963 | -0.2070 | **0.0026** |
| vCA1 | CD | UNEX | vSb | WD | EXP | -0.0470 | 0.9815 | na | na | 0.0009 | >0.999 | -0.0072 | >0.999 |
| vCA1 | CD | UNEX | vCA1 | WD | EXP | -0.0197 | >0.999 | na | na | -0.0240 | >0.999 | -0.0454 | >0.999 |
| vCA1 | CD | EXP | EcrhC | CD | UNEX | 0.0266 | >0.999 | na | na | 0.0536 | 0.9997 | 0.0625 | >0.999 |
| vCA1 | CD | EXP | CA2 | CD | UNEX | -0.0067 | >0.999 | na | na | 0.0256 | >0.999 | -0.0650 | >0.999 |
| vCA1 | CD | EXP | dCA3 | CD | UNEX | 0.0443 | 0.9932 | na | na | 0.0241 | >0.999 | -0.1730 | 0.0516 |
| vCA1 | CD | EXP | dDG | CD | UNEX | 0.0207 | >0.999 | na | na | 0.0383 | >0.999 | 0.0472 | >0.999 |
| vCA1 | CD | EXP | dSb | CD | UNEX | -0.0303 | >0.999 | na | na | 0.0685 | 0.9738 | 0.0721 | >0.999 |
| vCA1 | CD | EXP | EC | CD | UNEX | -0.0259 | >0.999 | na | na | -0.0170 | >0.999 | -0.0131 | >0.999 |
| vCA1 | CD | EXP | vCA3 | CD | UNEX | 0.0996 | **0.0034** | na | na | 0.0306 | >0.999 | -0.0872 | 0.9966 |
| vCA1 | CD | EXP | vDG | CD | UNEX | 0.0128 | >0.999 | na | na | -0.0404 | >0.999 | -0.1360 | 0.4519 |
| vCA1 | CD | EXP | vSb | CD | UNEX | -0.0407 | 0.9986 | na | na | 0.0195 | >0.999 | 0.0105 | >0.999 |
| vCA1 | CD | EXP | EcrhC | CD | EXP | 0.0252 | >0.999 | na | na | 0.0621 | 0.9895 | 0.0377 | >0.999 |
| vCA1 | CD | EXP | CA2 | CD | EXP | -0.0168 | >0.999 | na | na | 0.0169 | >0.999 | -0.1190 | 0.6831 |
| vCA1 | CD | EXP | dCA3 | CD | EXP | 0.0501 | 0.9250 | na | na | -0.0004 | >0.999 | -0.1990 | **0.0025** |
| vCA1 | CD | EXP | dDG | CD | EXP | 0.0220 | >0.999 | na | na | 0.0267 | >0.999 | 0.0271 | >0.999 |
| vCA1 | CD | EXP | dSb | CD | EXP | -0.0263 | >0.999 | na | na | 0.0498 | 0.9999 | 0.0594 | >0.999 |
| vCA1 | CD | EXP | EC | CD | EXP | -0.0266 | >0.999 | na | na | -0.0074 | >0.999 | -0.0101 | >0.999 |
| vCA1 | CD | EXP | vCA3 | CD | EXP | 0.0800 | 0.0600 | na | na | 0.0000 | >0.999 | -0.1300 | 0.4798 |
| vCA1 | CD | EXP | vDG | CD | EXP | 0.0035 | >0.999 | na | na | -0.0519 | 0.9997 | -0.1590 | 0.0868 |
| vCA1 | CD | EXP | vSb | CD | EXP | -0.0312 | >0.999 | na | na | 0.0232 | >0.999 | -0.0023 | >0.999 |
| vCA1 | CD | EXP | EcrhC | WD | UNEX | -0.0144 | >0.999 | na | na | 0.0522 | 0.9999 | 0.0811 | 0.9992 |
| vCA1 | CD | EXP | CA2 | WD | UNEX | -0.0510 | 0.9427 | na | na | 0.0040 | >0.999 | -0.1570 | 0.1526 |
| vCA1 | CD | EXP | dCA3 | WD | UNEX | 0.0196 | >0.999 | na | na | 0.0000 | >0.999 | -0.2200 | **0.0007** |
| vCA1 | CD | EXP | dDG | WD | UNEX | -0.0242 | >0.999 | na | na | 0.0417 | >0.999 | -0.0058 | >0.999 |
| vCA1 | CD | EXP | dSb | WD | UNEX | -0.0554 | 0.8545 | na | na | 0.0727 | 0.9421 | 0.0296 | >0.999 |
| vCA1 | CD | EXP | EC | WD | UNEX | -0.0550 | 0.8637 | na | na | -0.0177 | >0.999 | -0.0137 | >0.999 |
| vCA1 | CD | EXP | vCA1 | WD | UNEX | -0.0256 | >0.999 | na | na | -0.0081 | >0.999 | -0.0836 | 0.9985 |
| vCA1 | CD | EXP | vCA3 | WD | UNEX | 0.0480 | 0.9753 | na | na | -0.0210 | >0.999 | -0.1490 | 0.2426 |
| vCA1 | CD | EXP | vDG | WD | UNEX | -0.0300 | >0.999 | na | na | -0.0463 | >0.999 | -0.2060 | **0.0027** |
| vCA1 | CD | EXP | vSb | WD | UNEX | -0.0689 | 0.3662 | na | na | 0.0228 | >0.999 | -0.0242 | >0.999 |
| vCA1 | CD | EXP | EcrhC | WD | EXP | 0.0093 | >0.999 | na | na | 0.0139 | >0.999 | -0.0228 | >0.999 |
| vCA1 | CD | EXP | CA2 | WD | EXP | -0.0406 | 0.9970 | na | na | 0.0040 | >0.999 | -0.1280 | 0.5216 |
| vCA1 | CD | EXP | dCA3 | WD | EXP | 0.0274 | >0.999 | na | na | -0.0059 | >0.999 | -0.1870 | **0.0080** |
| vCA1 | CD | EXP | dDG | WD | EXP | 0.0093 | >0.999 | na | na | 0.0297 | >0.999 | 0.0440 | >0.999 |
| vCA1 | CD | EXP | dSb | WD | EXP | -0.0323 | >0.999 | na | na | 0.0730 | 0.9020 | 0.0826 | 0.9975 |
| vCA1 | CD | EXP | EC | WD | EXP | -0.0309 | >0.999 | na | na | -0.0265 | >0.999 | -0.0359 | >0.999 |
| vCA1 | CD | EXP | vCA1 | WD | EXP | -0.0128 | >0.999 | na | na | -0.0080 | >0.999 | -0.0267 | >0.999 |
| vCA1 | CD | EXP | vCA3 | WD | EXP | 0.0585 | 0.6718 | na | na | -0.0128 | >0.999 | -0.1370 | 0.3449 |
| vCA1 | CD | EXP | vDG | WD | EXP | -0.0168 | >0.999 | na | na | -0.0602 | 0.9938 | -0.1880 | **0.0073** |
| vCA1 | CD | EXP | vSb | WD | EXP | -0.0402 | 0.9976 | na | na | 0.0170 | >0.999 | 0.0115 | >0.999 |
| CA2 | CD | UNEX | EcrhC | CD | UNEX | 0.0332 | >0.999 | na | na | 0.0280 | >0.999 | 0.1280 | 0.7081 |
| CA2 | CD | UNEX | dCA3 | CD | UNEX | 0.0509 | 0.9658 | na | na | -0.0015 | >0.999 | -0.1080 | 0.9477 |
| CA2 | CD | UNEX | dDG | CD | UNEX | 0.0273 | >0.999 | na | na | 0.0127 | >0.999 | 0.1120 | 0.9128 |
| CA2 | CD | UNEX | dSb | CD | UNEX | -0.0236 | >0.999 | na | na | 0.0429 | >0.999 | 0.1370 | 0.5323 |
| CA2 | CD | UNEX | EC | CD | UNEX | -0.0193 | >0.999 | na | na | -0.0426 | >0.999 | 0.0520 | >0.999 |
| CA2 | CD | UNEX | vCA3 | CD | UNEX | 0.1060 | **0.0019** | na | na | 0.0050 | >0.999 | -0.0222 | >0.999 |
| CA2 | CD | UNEX | vDG | CD | UNEX | 0.0195 | >0.999 | na | na | -0.0661 | 0.9919 | -0.0714 | >0.999 |
| CA2 | CD | UNEX | vSb | CD | UNEX | -0.0341 | >0.999 | na | na | -0.0061 | >0.999 | 0.0755 | >0.999 |
| CA2 | CD | UNEX | EcrhC | CD | EXP | 0.0318 | >0.999 | na | na | 0.0365 | >0.999 | 0.1030 | 0.9541 |
| CA2 | CD | UNEX | CA2 | CD | EXP | -0.0101 | >0.999 | na | na | -0.0088 | >0.999 | -0.0543 | >0.999 |
| CA2 | CD | UNEX | dCA3 | CD | EXP | 0.0567 | 0.8162 | na | na | -0.0261 | >0.999 | -0.1340 | 0.4966 |
| CA2 | CD | UNEX | dDG | CD | EXP | 0.0287 | >0.999 | na | na | 0.0011 | >0.999 | 0.0921 | 0.9912 |
| CA2 | CD | UNEX | dSb | CD | EXP | -0.0197 | >0.999 | na | na | 0.0242 | >0.999 | 0.1240 | 0.6801 |
| CA2 | CD | UNEX | EC | CD | EXP | -0.0199 | >0.999 | na | na | -0.0331 | >0.999 | 0.0549 | >0.999 |
| CA2 | CD | UNEX | vCA3 | CD | EXP | 0.0867 | **0.0351** | na | na | -0.0256 | >0.999 | -0.0646 | >0.999 |
| CA2 | CD | UNEX | vDG | CD | EXP | 0.0102 | >0.999 | na | na | -0.0775 | 0.8777 | -0.0941 | 0.9874 |
| CA2 | CD | UNEX | vSb | CD | EXP | -0.0245 | >0.999 | na | na | -0.0024 | >0.999 | 0.0627 | >0.999 |
| CA2 | CD | UNEX | EcrhC | WD | UNEX | -0.0078 | >0.999 | na | na | 0.0266 | >0.999 | 0.1460 | 0.3735 |
| CA2 | CD | UNEX | CA2 | WD | UNEX | -0.0443 | 0.9964 | na | na | -0.0216 | >0.999 | -0.0921 | 0.9954 |
| CA2 | CD | UNEX | dCA3 | WD | UNEX | 0.0262 | >0.999 | na | na | -0.0257 | >0.999 | -0.1550 | 0.2460 |
| CA2 | CD | UNEX | dDG | WD | UNEX | -0.0176 | >0.999 | na | na | 0.0161 | >0.999 | 0.0592 | >0.999 |
| CA2 | CD | UNEX | dSb | WD | UNEX | -0.0487 | 0.9821 | na | na | 0.0470 | >0.999 | 0.0946 | 0.9926 |
| CA2 | CD | UNEX | EC | WD | UNEX | -0.0483 | 0.9841 | na | na | -0.0433 | >0.999 | 0.0514 | >0.999 |
| CA2 | CD | UNEX | vCA3 | WD | UNEX | 0.0546 | 0.9160 | na | na | -0.0467 | >0.999 | -0.0843 | 0.9992 |
| CA2 | CD | UNEX | vDG | WD | UNEX | -0.0233 | >0.999 | na | na | -0.0719 | 0.9692 | -0.1410 | 0.4548 |
| CA2 | CD | UNEX | vSb | WD | UNEX | -0.0623 | 0.7065 | na | na | -0.0028 | >0.999 | 0.0409 | >0.999 |
| CA2 | CD | UNEX | EcrhC | WD | EXP | 0.0160 | >0.999 | na | na | -0.0117 | >0.999 | 0.0422 | >0.999 |
| CA2 | CD | UNEX | CA2 | WD | EXP | -0.0340 | >0.999 | na | na | -0.0217 | >0.999 | -0.0625 | >0.999 |
| CA2 | CD | UNEX | dCA3 | WD | EXP | 0.0340 | >0.999 | na | na | -0.0315 | >0.999 | -0.1220 | 0.7202 |
| CA2 | CD | UNEX | dDG | WD | EXP | 0.0160 | >0.999 | na | na | 0.0041 | >0.999 | 0.1090 | 0.9045 |
| CA2 | CD | UNEX | dSb | WD | EXP | -0.0257 | >0.999 | na | na | 0.0473 | >0.999 | 0.1480 | 0.2661 |
| CA2 | CD | UNEX | EC | WD | EXP | -0.0243 | >0.999 | na | na | -0.0521 | 0.9999 | 0.0291 | >0.999 |
| CA2 | CD | UNEX | vCA3 | WD | EXP | 0.0652 | 0.5052 | na | na | -0.0384 | >0.999 | -0.0719 | >0.999 |
| CA2 | CD | UNEX | vDG | WD | EXP | -0.0101 | >0.999 | na | na | -0.0859 | 0.6947 | -0.1230 | 0.7041 |
| CA2 | CD | UNEX | vSb | WD | EXP | -0.0335 | >0.999 | na | na | -0.0087 | >0.999 | 0.0766 | 0.9998 |
| CA2 | CD | EXP | EcrhC | CD | UNEX | 0.0434 | 0.9953 | na | na | 0.0367 | >0.999 | 0.1820 | **0.0249** |
| CA2 | CD | EXP | dDG | CD | UNEX | 0.0375 | 0.9998 | na | na | 0.0214 | >0.999 | 0.1670 | 0.0808 |
| CA2 | CD | EXP | dSb | CD | UNEX | -0.0135 | >0.999 | na | na | 0.0517 | 0.9999 | 0.1920 | **0.0108** |
| CA2 | CD | EXP | EC | CD | UNEX | -0.0092 | >0.999 | na | na | -0.0339 | >0.999 | 0.1060 | 0.9291 |
| CA2 | CD | EXP | vDG | CD | UNEX | 0.0296 | >0.999 | na | na | -0.0573 | 0.9989 | -0.0171 | >0.999 |
| CA2 | CD | EXP | vSb | CD | UNEX | -0.0240 | >0.999 | na | na | 0.0026 | >0.999 | 0.1300 | 0.5776 |
| CA2 | CD | EXP | CA2 | CD | UNEX | 0.0610 | 0.6675 | na | na | 0.0073 | >0.999 | -0.0533 | >0.999 |
| CA2 | CD | EXP | dCA3 | CD | UNEX | 0.1160 | **<0.0001** | na | na | 0.0137 | >0.999 | 0.0321 | >0.999 |
| CA2 | CD | EXP | dCA3 | CD | EXP | 0.0669 | 0.3430 | na | na | -0.0173 | >0.999 | -0.0797 | 0.9987 |
| CA2 | CD | EXP | vCA3 | CD | EXP | 0.0968 | **0.0026** | na | na | -0.0169 | >0.999 | -0.0103 | >0.999 |
| CA2 | CD | EXP | EcrhC | CD | EXP | 0.0420 | 0.9946 | na | na | 0.0452 | >0.999 | 0.1570 | 0.1005 |
| CA2 | CD | EXP | dDG | CD | EXP | 0.0388 | 0.9988 | na | na | 0.0098 | >0.999 | 0.1460 | 0.2035 |
| CA2 | CD | EXP | dSb | CD | EXP | -0.0096 | >0.999 | na | na | 0.0330 | >0.999 | 0.1790 | **0.0174** |
| CA2 | CD | EXP | EC | CD | EXP | -0.0098 | >0.999 | na | na | -0.0243 | >0.999 | 0.1090 | 0.8514 |
| CA2 | CD | EXP | vDG | CD | EXP | 0.0203 | >0.999 | na | na | -0.0687 | 0.9525 | -0.0397 | >0.999 |
| CA2 | CD | EXP | vSb | CD | EXP | -0.0144 | >0.999 | na | na | 0.0063 | >0.999 | 0.1170 | 0.7254 |
| CA2 | CD | EXP | EcrhC | WD | UNEX | 0.0024 | >0.999 | na | na | 0.0353 | >0.999 | 0.2000 | **0.0047** |
| CA2 | CD | EXP | CA2 | WD | UNEX | -0.0342 | >0.999 | na | na | -0.0129 | >0.999 | -0.0378 | >0.999 |
| CA2 | CD | EXP | dDG | WD | UNEX | -0.0074 | >0.999 | na | na | 0.0249 | >0.999 | 0.1140 | 0.8532 |
| CA2 | CD | EXP | dSb | WD | UNEX | -0.0386 | 0.9995 | na | na | 0.0558 | 0.9994 | 0.1490 | 0.2478 |
| CA2 | CD | EXP | EC | WD | UNEX | -0.0382 | 0.9996 | na | na | -0.0346 | >0.999 | 0.1060 | 0.9338 |
| CA2 | CD | EXP | vDG | WD | UNEX | -0.0132 | >0.999 | na | na | -0.0632 | 0.9929 | -0.0871 | 0.9968 |
| CA2 | CD | EXP | vSb | WD | UNEX | -0.0521 | 0.9248 | na | na | 0.0059 | >0.999 | 0.0952 | 0.9848 |
| CA2 | CD | EXP | dCA3 | WD | UNEX | 0.0363 | 0.9999 | na | na | -0.0169 | >0.999 | -0.1000 | 0.9667 |
| CA2 | CD | EXP | vCA3 | WD | UNEX | 0.0647 | 0.5227 | na | na | -0.0379 | >0.999 | -0.0299 | >0.999 |
| CA2 | CD | EXP | EcrhC | WD | EXP | 0.0261 | >0.999 | na | na | -0.0030 | >0.999 | 0.0966 | 0.9658 |
| CA2 | CD | EXP | CA2 | WD | EXP | -0.0239 | >0.999 | na | na | -0.0129 | >0.999 | -0.0082 | >0.999 |
| CA2 | CD | EXP | dDG | WD | EXP | 0.0261 | >0.999 | na | na | 0.0129 | >0.999 | 0.1630 | 0.0628 |
| CA2 | CD | EXP | dSb | WD | EXP | -0.0155 | >0.999 | na | na | 0.0561 | 0.9984 | 0.2020 | **0.0018** |
| CA2 | CD | EXP | EC | WD | EXP | -0.0141 | >0.999 | na | na | -0.0434 | >0.999 | 0.0834 | 0.9969 |
| CA2 | CD | EXP | vDG | WD | EXP | 0.0000 | >0.999 | na | na | -0.0771 | 0.8263 | -0.0687 | >0.999 |
| CA2 | CD | EXP | vSb | WD | EXP | -0.0234 | >0.999 | na | na | 0.0001 | >0.999 | 0.1310 | 0.4549 |
| CA2 | CD | EXP | dCA3 | WD | EXP | 0.0442 | 0.9872 | na | na | -0.0227 | >0.999 | -0.0678 | >0.999 |
| CA2 | CD | EXP | vCA3 | WD | EXP | 0.0753 | 0.1217 | na | na | -0.0296 | >0.999 | -0.0175 | >0.999 |
| dCA3 | CD | UNEX | dDG | CD | UNEX | -0.0236 | >0.999 | na | na | 0.0142 | >0.999 | 0.2200 | **0.0015** |
| dCA3 | CD | UNEX | EC | CD | UNEX | -0.0702 | 0.4119 | na | na | -0.0411 | >0.999 | 0.1600 | 0.1871 |
| dCA3 | CD | UNEX | vDG | CD | UNEX | -0.0315 | >0.999 | na | na | -0.0646 | 0.9946 | 0.0363 | >0.999 |
| dCA3 | CD | UNEX | dSb | CD | UNEX | -0.0745 | 0.2723 | na | na | 0.0444 | >0.999 | 0.2450 | **0.0001** |
| dCA3 | CD | UNEX | EcrhC | CD | UNEX | -0.0177 | >0.999 | na | na | 0.0295 | >0.999 | 0.2350 | **0.0003** |
| dCA3 | CD | UNEX | vSb | CD | UNEX | -0.0850 | 0.0738 | na | na | -0.0046 | >0.999 | 0.1830 | **0.0384** |
| dCA3 | CD | UNEX | vCA3 | CD | UNEX | 0.0553 | 0.9025 | na | na | 0.0065 | >0.999 | 0.0854 | 0.9989 |
| dCA3 | CD | UNEX | dDG | CD | EXP | -0.0223 | >0.999 | na | na | 0.0026 | >0.999 | 0.2000 | **0.0051** |
| dCA3 | CD | UNEX | EC | CD | EXP | -0.0708 | 0.3030 | na | na | -0.0316 | >0.999 | 0.1630 | 0.1072 |
| dCA3 | CD | UNEX | vDG | CD | EXP | -0.0408 | 0.9986 | na | na | -0.0760 | 0.9011 | 0.0136 | >0.999 |
| dCA3 | CD | UNEX | dSb | CD | EXP | -0.0706 | 0.3101 | na | na | 0.0257 | >0.999 | 0.2320 | **0.0002** |
| dCA3 | CD | UNEX | EcrhC | CD | EXP | -0.0191 | >0.999 | na | na | 0.0380 | >0.999 | 0.2100 | **0.0018** |
| dCA3 | CD | UNEX | vSb | CD | EXP | -0.0754 | 0.1785 | na | na | -0.0009 | >0.999 | 0.1700 | 0.0612 |
| dCA3 | CD | UNEX | dCA3 | CD | EXP | 0.0058 | >0.999 | na | na | -0.0246 | >0.999 | -0.0264 | >0.999 |
| dCA3 | CD | UNEX | vCA3 | CD | EXP | 0.0358 | >0.999 | na | na | -0.0241 | >0.999 | 0.0431 | >0.999 |
| dCA3 | CD | UNEX | dCA3 | WD | UNEX | -0.0247 | >0.999 | na | na | -0.0242 | >0.999 | -0.0470 | >0.999 |
| dCA3 | CD | UNEX | vCA3 | WD | UNEX | 0.0037 | >0.999 | na | na | -0.0452 | >0.999 | 0.0234 | >0.999 |
| dCA3 | CD | UNEX | dDG | WD | UNEX | -0.0685 | 0.4741 | na | na | 0.0176 | >0.999 | 0.1670 | 0.1205 |
| dCA3 | CD | UNEX | EC | WD | UNEX | -0.0992 | **0.0072** | na | na | -0.0418 | >0.999 | 0.1590 | 0.1936 |
| dCA3 | CD | UNEX | vDG | WD | UNEX | -0.0742 | 0.2809 | na | na | -0.0704 | 0.9774 | -0.0337 | >0.999 |
| dCA3 | CD | UNEX | dSb | WD | UNEX | -0.0996 | **0.0067** | na | na | 0.0485 | >0.999 | 0.2020 | **0.0080** |
| dCA3 | CD | UNEX | EcrhC | WD | UNEX | -0.0587 | 0.8217 | na | na | 0.0281 | >0.999 | 0.2540 | **<0.0001** |
| dCA3 | CD | UNEX | vSb | WD | UNEX | -0.1130 | **0.0005** | na | na | -0.0013 | >0.999 | 0.1490 | 0.3345 |
| dCA3 | CD | UNEX | dDG | WD | EXP | -0.0349 | >0.999 | na | na | 0.0056 | >0.999 | 0.2170 | **0.0009** |
| dCA3 | CD | UNEX | EC | WD | EXP | -0.0752 | 0.1847 | na | na | -0.0506 | >0.999 | 0.1370 | 0.4454 |
| dCA3 | CD | UNEX | vDG | WD | EXP | -0.0610 | 0.6687 | na | na | -0.0844 | 0.7330 | -0.0154 | >0.999 |
| dCA3 | CD | UNEX | dSb | WD | EXP | -0.0766 | 0.1548 | na | na | 0.0488 | >0.999 | 0.2550 | **<0.0001** |
| dCA3 | CD | UNEX | EcrhC | WD | EXP | -0.0349 | >0.999 | na | na | -0.0102 | >0.999 | 0.1500 | 0.2345 |
| dCA3 | CD | UNEX | vSb | WD | EXP | -0.0844 | 0.0504 | na | na | -0.0071 | >0.999 | 0.1840 | **0.0204** |
| dCA3 | CD | UNEX | dCA3 | WD | EXP | -0.0169 | >0.999 | na | na | -0.0300 | >0.999 | -0.0145 | >0.999 |
| dCA3 | CD | UNEX | vCA3 | WD | EXP | 0.0143 | >0.999 | na | na | -0.0369 | >0.999 | 0.0358 | >0.999 |
| dCA3 | CD | EXP | vCA3 | CD | UNEX | 0.0495 | 0.9613 | na | na | 0.0310 | >0.999 | 0.1120 | 0.8744 |
| dCA3 | CD | EXP | dDG | CD | UNEX | -0.0294 | >0.999 | na | na | 0.0388 | >0.999 | 0.2460 | **<0.0001** |
| dCA3 | CD | EXP | dSb | CD | UNEX | -0.0803 | 0.0928 | na | na | 0.0690 | 0.9713 | 0.2710 | **<0.0001** |
| dCA3 | CD | EXP | EC | CD | UNEX | -0.0760 | 0.1664 | na | na | -0.0165 | >0.999 | 0.1860 | **0.0175** |
| dCA3 | CD | EXP | EcrhC | CD | UNEX | -0.0235 | >0.999 | na | na | 0.0541 | 0.9997 | 0.2620 | **<0.0001** |
| dCA3 | CD | EXP | vDG | CD | UNEX | -0.0373 | 0.9998 | na | na | -0.0400 | >0.999 | 0.0626 | >0.999 |
| dCA3 | CD | EXP | vSb | CD | UNEX | -0.0908 | **0.0173** | na | na | 0.0200 | >0.999 | 0.2100 | **0.0020** |
| dCA3 | CD | EXP | dDG | CD | EXP | -0.0281 | >0.999 | na | na | 0.0272 | >0.999 | 0.2260 | **0.0001** |
| dCA3 | CD | EXP | dSb | CD | EXP | -0.0764 | 0.1041 | na | na | 0.0503 | 0.9998 | 0.2580 | **<0.0001** |
| dCA3 | CD | EXP | EC | CD | EXP | -0.0766 | 0.1008 | na | na | -0.0070 | >0.999 | 0.1890 | **0.0068** |
| dCA3 | CD | EXP | EcrhC | CD | EXP | -0.0249 | >0.999 | na | na | 0.0626 | 0.9882 | 0.2370 | **<0.0001** |
| dCA3 | CD | EXP | vDG | CD | EXP | -0.0466 | 0.9711 | na | na | -0.0514 | 0.9997 | 0.0400 | >0.999 |
| dCA3 | CD | EXP | vSb | CD | EXP | -0.0813 | **0.0491** | na | na | 0.0237 | >0.999 | 0.1970 | **0.0031** |
| dCA3 | CD | EXP | vCA3 | CD | EXP | 0.0299 | >0.999 | na | na | 0.0004 | >0.999 | 0.0694 | >0.999 |
| dCA3 | CD | EXP | dCA3 | WD | UNEX | -0.0305 | >0.999 | na | na | 0.0004 | >0.999 | -0.0206 | >0.999 |
| dCA3 | CD | EXP | vCA3 | WD | UNEX | -0.0021 | >0.999 | na | na | -0.0206 | >0.999 | 0.0497 | >0.999 |
| dCA3 | CD | EXP | dDG | WD | UNEX | -0.0743 | 0.2052 | na | na | 0.0422 | >0.999 | 0.1930 | **0.0092** |
| dCA3 | CD | EXP | dSb | WD | UNEX | -0.1050 | **0.0010** | na | na | 0.0731 | 0.9374 | 0.2290 | **0.0003** |
| dCA3 | CD | EXP | EC | WD | UNEX | -0.1050 | **0.0011** | na | na | -0.0172 | >0.999 | 0.1850 | **0.0184** |
| dCA3 | CD | EXP | EcrhC | WD | UNEX | -0.0645 | 0.5326 | na | na | 0.0526 | 0.9998 | 0.2800 | **<0.0001** |
| dCA3 | CD | EXP | vDG | WD | UNEX | -0.0800 | 0.0968 | na | na | -0.0459 | >0.999 | -0.0074 | >0.999 |
| dCA3 | CD | EXP | vSb | WD | UNEX | -0.1190 | **<0.0001** | na | na | 0.0232 | >0.999 | 0.1750 | **0.0435** |
| dCA3 | CD | EXP | dDG | WD | EXP | -0.0408 | 0.9968 | na | na | 0.0302 | >0.999 | 0.2430 | **<0.0001** |
| dCA3 | CD | EXP | dSb | WD | EXP | -0.0824 | **0.0407** | na | na | 0.0734 | 0.8950 | 0.2820 | **<0.0001** |
| dCA3 | CD | EXP | EC | WD | EXP | -0.0810 | 0.0514 | na | na | -0.0260 | >0.999 | 0.1630 | 0.0641 |
| dCA3 | CD | EXP | EcrhC | WD | EXP | -0.0407 | 0.9969 | na | na | 0.0144 | >0.999 | 0.1760 | **0.0217** |
| dCA3 | CD | EXP | vDG | WD | EXP | -0.0668 | 0.3441 | na | na | -0.0598 | 0.9946 | 0.0110 | >0.999 |
| dCA3 | CD | EXP | vSb | WD | EXP | -0.0902 | **0.0099** | na | na | 0.0174 | >0.999 | 0.2110 | **0.0007** |
| dCA3 | CD | EXP | dCA3 | WD | EXP | -0.0227 | >0.999 | na | na | -0.0054 | >0.999 | 0.0119 | >0.999 |
| dCA3 | CD | EXP | vCA3 | WD | EXP | 0.0085 | >0.999 | na | na | -0.0123 | >0.999 | 0.0622 | >0.999 |
| vCA3 | CD | UNEX | dDG | CD | UNEX | -0.0789 | 0.1663 | na | na | 0.0077 | >0.999 | 0.1350 | 0.5821 |
| vCA3 | CD | UNEX | dSb | CD | UNEX | -0.1300 | **<0.0001** | na | na | 0.0380 | >0.999 | 0.1590 | 0.1899 |
| vCA3 | CD | UNEX | EC | CD | UNEX | -0.1260 | **<0.0001** | na | na | -0.0476 | >0.999 | 0.0742 | >0.999 |
| vCA3 | CD | UNEX | vDG | CD | UNEX | -0.0868 | 0.0570 | na | na | -0.0710 | 0.9745 | -0.0492 | >0.999 |
| vCA3 | CD | UNEX | vSb | CD | UNEX | -0.1400 | **<0.0001** | na | na | -0.0111 | >0.999 | 0.0977 | 0.9873 |
| vCA3 | CD | UNEX | EcrhC | CD | UNEX | -0.0730 | 0.3184 | na | na | 0.0230 | >0.999 | 0.1500 | 0.3163 |
| vCA3 | CD | UNEX | dDG | CD | EXP | -0.0776 | 0.1359 | na | na | -0.0039 | >0.999 | 0.1140 | 0.8434 |
| vCA3 | CD | UNEX | dSb | CD | EXP | -0.1260 | **<0.0001** | na | na | 0.0193 | >0.999 | 0.1470 | 0.2804 |
| vCA3 | CD | UNEX | EC | CD | EXP | -0.1260 | **<0.0001** | na | na | -0.0380 | >0.999 | 0.0771 | 0.9997 |
| vCA3 | CD | UNEX | vDG | CD | EXP | -0.0961 | **0.0066** | na | na | -0.0824 | 0.7791 | -0.0719 | >0.999 |
| vCA3 | CD | UNEX | vSb | CD | EXP | -0.1310 | **<0.0001** | na | na | -0.0074 | >0.999 | 0.0850 | 0.9980 |
| vCA3 | CD | UNEX | EcrhC | CD | EXP | -0.0744 | 0.2032 | na | na | 0.0315 | >0.999 | 0.1250 | 0.6702 |
| vCA3 | CD | UNEX | dDG | WD | UNEX | -0.1240 | **<0.0001** | na | na | 0.0112 | >0.999 | 0.0814 | 0.9996 |
| vCA3 | CD | UNEX | dSb | WD | UNEX | -0.1550 | **<0.0001** | na | na | 0.0421 | >0.999 | 0.1170 | 0.8658 |
| vCA3 | CD | UNEX | EC | WD | UNEX | -0.1550 | **<0.0001** | na | na | -0.0483 | >0.999 | 0.0736 | >0.999 |
| vCA3 | CD | UNEX | vCA3 | WD | UNEX | -0.0516 | 0.9586 | na | na | -0.0516 | >0.999 | -0.0621 | >0.999 |
| vCA3 | CD | UNEX | vDG | WD | UNEX | -0.1300 | **<0.0001** | na | na | -0.0769 | 0.9263 | -0.1190 | 0.8358 |
| vCA3 | CD | UNEX | vSb | WD | UNEX | -0.1690 | **<0.0001** | na | na | -0.0078 | >0.999 | 0.0631 | >0.999 |
| vCA3 | CD | UNEX | EcrhC | WD | UNEX | -0.1140 | **0.0004** | na | na | 0.0216 | >0.999 | 0.1680 | 0.1098 |
| vCA3 | CD | UNEX | vCA3 | CD | EXP | -0.0196 | >0.999 | na | na | -0.0306 | >0.999 | -0.0424 | >0.999 |
| vCA3 | CD | UNEX | dDG | WD | EXP | -0.0903 | **0.0191** | na | na | -0.0008 | >0.999 | 0.1310 | 0.5499 |
| vCA3 | CD | UNEX | dSb | WD | EXP | -0.1320 | **<0.0001** | na | na | 0.0424 | >0.999 | 0.1700 | 0.0640 |
| vCA3 | CD | UNEX | EC | WD | EXP | -0.1310 | **<0.0001** | na | na | -0.0571 | 0.9990 | 0.0513 | >0.999 |
| vCA3 | CD | UNEX | vCA3 | WD | EXP | -0.0410 | 0.9983 | na | na | -0.0433 | >0.999 | -0.0497 | >0.999 |
| vCA3 | CD | UNEX | vDG | WD | EXP | -0.1160 | **<0.0001** | na | na | -0.0908 | 0.5602 | -0.1010 | 0.9642 |
| vCA3 | CD | UNEX | vSb | WD | EXP | -0.1400 | **<0.0001** | na | na | -0.0136 | >0.999 | 0.0988 | 0.9734 |
| vCA3 | CD | UNEX | EcrhC | WD | EXP | -0.0902 | **0.0192** | na | na | -0.0166 | >0.999 | 0.0645 | >0.999 |
| vCA3 | CD | EXP | dDG | CD | UNEX | -0.0593 | 0.7306 | na | na | 0.0383 | >0.999 | 0.1770 | **0.0372** |
| vCA3 | CD | EXP | dSb | CD | UNEX | -0.1100 | **0.0004** | na | na | 0.0686 | 0.9738 | 0.2020 | **0.0042** |
| vCA3 | CD | EXP | EC | CD | UNEX | -0.1060 | **0.0009** | na | na | -0.0170 | >0.999 | 0.1170 | 0.8112 |
| vCA3 | CD | EXP | EcrhC | CD | UNEX | -0.0534 | 0.8999 | na | na | 0.0536 | 0.9997 | 0.1920 | **0.0102** |
| vCA3 | CD | EXP | vDG | CD | UNEX | -0.0672 | 0.4271 | na | na | -0.0404 | >0.999 | -0.0068 | >0.999 |
| vCA3 | CD | EXP | vSb | CD | UNEX | -0.1210 | **<0.0001** | na | na | 0.0195 | >0.999 | 0.1400 | 0.3856 |
| vCA3 | CD | EXP | EC | CD | UNEX | -0.1350 | **<0.0001** | na | na | -0.0177 | >0.999 | 0.1160 | 0.8200 |
| vCA3 | CD | EXP | dDG | CD | EXP | -0.0580 | 0.6919 | na | na | 0.0267 | >0.999 | 0.1570 | 0.1033 |
| vCA3 | CD | EXP | dSb | CD | EXP | -0.1060 | **0.0003** | na | na | 0.0499 | 0.9999 | 0.1890 | **0.0067** |
| vCA3 | CD | EXP | EC | CD | EXP | -0.1070 | **0.0003** | na | na | -0.0074 | >0.999 | 0.1200 | 0.6806 |
| vCA3 | CD | EXP | EcrhC | CD | EXP | -0.0548 | 0.8057 | na | na | 0.0621 | 0.9895 | 0.1670 | **0.0460** |
| vCA3 | CD | EXP | vDG | CD | EXP | -0.0765 | 0.1024 | na | na | -0.0519 | 0.9997 | -0.0295 | >0.999 |
| vCA3 | CD | EXP | vSb | CD | EXP | -0.1110 | **0.0001** | na | na | 0.0232 | >0.999 | 0.1270 | 0.5247 |
| vCA3 | CD | EXP | dDG | WD | UNEX | -0.1040 | **0.0013** | na | na | 0.0418 | >0.999 | 0.1240 | 0.6908 |
| vCA3 | CD | EXP | dSb | WD | UNEX | -0.1350 | **<0.0001** | na | na | 0.0727 | 0.9419 | 0.1590 | 0.1339 |
| vCA3 | CD | EXP | EcrhC | WD | UNEX | -0.0944 | **0.0090** | na | na | 0.0522 | 0.9999 | 0.2110 | **0.0017** |
| vCA3 | CD | EXP | vCA3 | WD | UNEX | -0.0321 | >0.999 | na | na | -0.0210 | >0.999 | -0.0197 | >0.999 |
| vCA3 | CD | EXP | vDG | WD | UNEX | -0.1100 | **0.0004** | na | na | -0.0463 | >0.999 | -0.0768 | 0.9998 |
| vCA3 | CD | EXP | vSb | WD | UNEX | -0.1490 | **<0.0001** | na | na | 0.0228 | >0.999 | 0.1060 | 0.9356 |
| vCA3 | CD | EXP | EC | WD | EXP | -0.1110 | **0.0001** | na | na | -0.0265 | >0.999 | 0.0937 | 0.9779 |
| vCA3 | CD | EXP | dDG | WD | EXP | -0.0707 | 0.2227 | na | na | 0.0298 | >0.999 | 0.1740 | **0.0272** |
| vCA3 | CD | EXP | dSb | WD | EXP | -0.1120 | **<0.0001** | na | na | 0.0730 | 0.9017 | 0.2120 | **0.0006** |
| vCA3 | CD | EXP | EcrhC | WD | EXP | -0.0707 | 0.2236 | na | na | 0.0140 | >0.999 | 0.1070 | 0.8813 |
| vCA3 | CD | EXP | vCA3 | WD | EXP | -0.0215 | >0.999 | na | na | -0.0127 | >0.999 | -0.0073 | >0.999 |
| vCA3 | CD | EXP | vDG | WD | EXP | -0.0968 | **0.0027** | na | na | -0.0602 | 0.9938 | -0.0585 | >0.999 |
| vCA3 | CD | EXP | vSb | WD | EXP | -0.1200 | **<0.0001** | na | na | 0.0170 | >0.999 | 0.1410 | 0.2756 |
| dDG | CD | UNEX | dSb | CD | UNEX | -0.0510 | 0.9652 | na | na | 0.0302 | >0.999 | 0.0249 | >0.999 |
| dDG | CD | UNEX | EcrhC | CD | UNEX | 0.0059 | >0.999 | na | na | 0.0153 | >0.999 | 0.0152 | >0.999 |
| dDG | CD | UNEX | vDG | CD | UNEX | -0.0079 | >0.999 | na | na | -0.0788 | 0.9026 | -0.1840 | **0.0369** |
| dDG | CD | UNEX | vSb | CD | UNEX | -0.0614 | 0.7347 | na | na | -0.0188 | >0.999 | -0.0368 | >0.999 |
| dDG | CD | UNEX | EC | CD | UNEX | -0.0466 | 0.9912 | na | na | -0.0553 | 0.9998 | -0.0603 | >0.999 |
| dDG | CD | UNEX | EC | CD | UNEX | -0.0757 | 0.2402 | na | na | -0.0560 | 0.9997 | -0.0609 | >0.999 |
| dDG | CD | UNEX | EC | CD | EXP | -0.0472 | 0.9802 | na | na | -0.0457 | >0.999 | -0.0574 | >0.999 |
| dDG | CD | UNEX | dDG | CD | EXP | 0.0013 | >0.999 | na | na | -0.0116 | >0.999 | -0.0202 | >0.999 |
| dDG | CD | UNEX | dSb | CD | EXP | -0.0470 | 0.9816 | na | na | 0.0115 | >0.999 | 0.0121 | >0.999 |
| dDG | CD | UNEX | EcrhC | CD | EXP | 0.0045 | >0.999 | na | na | 0.0238 | >0.999 | -0.0095 | >0.999 |
| dDG | CD | UNEX | vDG | CD | EXP | -0.0172 | >0.999 | na | na | -0.0902 | 0.5778 | -0.2060 | **0.0027** |
| dDG | CD | UNEX | vSb | CD | EXP | -0.0519 | 0.9289 | na | na | -0.0151 | >0.999 | -0.0495 | >0.999 |
| dDG | CD | UNEX | dDG | WD | UNEX | -0.0449 | 0.9954 | na | na | 0.0034 | >0.999 | -0.0531 | >0.999 |
| dDG | CD | UNEX | dSb | WD | UNEX | -0.0761 | 0.2309 | na | na | 0.0343 | >0.999 | -0.0177 | >0.999 |
| dDG | CD | UNEX | EcrhC | WD | UNEX | -0.0351 | >0.999 | na | na | 0.0139 | >0.999 | 0.0338 | >0.999 |
| dDG | CD | UNEX | vDG | WD | UNEX | -0.0507 | 0.9680 | na | na | -0.0846 | 0.7992 | -0.2540 | <0.0001 |
| dDG | CD | UNEX | vSb | WD | UNEX | -0.0896 | **0.0370** | na | na | -0.0155 | >0.999 | -0.0714 | >0.999 |
| dDG | CD | UNEX | EC | WD | EXP | -0.0516 | 0.9334 | na | na | -0.0648 | 0.9891 | -0.0832 | 0.9987 |
| dDG | CD | UNEX | dDG | WD | EXP | -0.0114 | >0.999 | na | na | -0.0086 | >0.999 | -0.0032 | >0.999 |
| dDG | CD | UNEX | dSb | WD | EXP | -0.0530 | 0.9084 | na | na | 0.0346 | >0.999 | 0.0353 | >0.999 |
| dDG | CD | UNEX | EcrhC | WD | EXP | -0.0114 | >0.999 | na | na | -0.0244 | >0.999 | -0.0700 | >0.999 |
| dDG | CD | UNEX | vDG | WD | EXP | -0.0374 | 0.9998 | na | na | -0.0986 | 0.3552 | -0.2350 | **0.0001** |
| dDG | CD | UNEX | vSb | WD | EXP | -0.0609 | 0.6744 | na | na | -0.0213 | >0.999 | -0.0357 | >0.999 |
| dDG | CD | EXP | dSb | CD | UNEX | -0.0523 | 0.9223 | na | na | 0.0418 | >0.999 | 0.0451 | >0.999 |
| dDG | CD | EXP | dSb | CD | UNEX | -0.0774 | 0.1397 | na | na | 0.0459 | >0.999 | 0.0025 | >0.999 |
| dDG | CD | EXP | EC | CD | UNEX | -0.0479 | 0.9755 | na | na | -0.0437 | >0.999 | -0.0401 | >0.999 |
| dDG | CD | EXP | EcrhC | CD | UNEX | 0.0046 | >0.999 | na | na | 0.0269 | >0.999 | 0.0354 | >0.999 |
| dDG | CD | EXP | vDG | CD | UNEX | -0.0092 | >0.999 | na | na | -0.0672 | 0.9807 | -0.1640 | 0.1006 |
| dDG | CD | EXP | vSb | CD | UNEX | -0.0627 | 0.6011 | na | na | -0.0072 | >0.999 | -0.0166 | >0.999 |
| dDG | CD | EXP | dSb | CD | EXP | -0.0483 | 0.9518 | na | na | 0.0231 | >0.999 | 0.0323 | >0.999 |
| dDG | CD | EXP | EC | CD | EXP | -0.0486 | 0.9489 | na | na | -0.0341 | >0.999 | -0.0372 | >0.999 |
| dDG | CD | EXP | EcrhC | CD | EXP | 0.0032 | >0.999 | na | na | 0.0354 | >0.999 | 0.0107 | >0.999 |
| dDG | CD | EXP | vDG | CD | EXP | -0.0185 | >0.999 | na | na | -0.0786 | 0.7939 | -0.1860 | **0.0088** |
| dDG | CD | EXP | vSb | CD | EXP | -0.0532 | 0.8544 | na | na | -0.0035 | >0.999 | -0.0293 | >0.999 |
| dDG | CD | EXP | vSb | WD | UNEX | -0.0909 | **0.0170** | na | na | -0.0039 | >0.999 | -0.0512 | >0.999 |
| dDG | CD | EXP | EC | WD | UNEX | -0.0770 | 0.1464 | na | na | -0.0444 | >0.999 | -0.0407 | >0.999 |
| dDG | CD | EXP | EcrhC | WD | UNEX | -0.0364 | 0.9999 | na | na | 0.0255 | >0.999 | 0.0540 | >0.999 |
| dDG | CD | EXP | vDG | WD | UNEX | -0.0520 | 0.9274 | na | na | -0.0730 | 0.9383 | -0.2340 | **0.0002** |
| dDG | CD | EXP | dDG | WD | UNEX | -0.0462 | 0.9860 | na | na | 0.0150 | >0.999 | -0.0329 | >0.999 |
| dDG | CD | EXP | dDG | WD | EXP | -0.0127 | >0.999 | na | na | 0.0030 | >0.999 | 0.0170 | >0.999 |
| dDG | CD | EXP | dSb | WD | EXP | -0.0543 | 0.8216 | na | na | 0.0462 | >0.999 | 0.0555 | >0.999 |
| dDG | CD | EXP | EC | WD | EXP | -0.0529 | 0.8619 | na | na | -0.0532 | 0.9995 | -0.0630 | >0.999 |
| dDG | CD | EXP | EcrhC | WD | EXP | -0.0127 | >0.999 | na | na | -0.0128 | >0.999 | -0.0498 | >0.999 |
| dDG | CD | EXP | vDG | WD | EXP | -0.0388 | 0.9988 | na | na | -0.0870 | 0.5689 | -0.2150 | **0.0004** |
| dDG | CD | EXP | vSb | WD | EXP | -0.0622 | 0.5243 | na | na | -0.0097 | >0.999 | -0.0155 | >0.999 |
| vDG | CD | UNEX | dSb | CD | UNEX | -0.0431 | 0.9979 | na | na | 0.1090 | 0.2183 | 0.2090 | **0.0045** |
| vDG | CD | UNEX | EC | CD | UNEX | -0.0387 | 0.9998 | na | na | 0.0235 | >0.999 | 0.1230 | 0.7758 |
| vDG | CD | UNEX | EcrhC | CD | UNEX | 0.0138 | >0.999 | na | na | 0.0941 | 0.5651 | 0.1990 | **0.0107** |
| vDG | CD | UNEX | vSb | CD | UNEX | -0.0535 | 0.9335 | na | na | 0.0599 | 0.9987 | 0.1470 | 0.3609 |
| vDG | CD | UNEX | dSb | CD | EXP | -0.0391 | 0.9994 | na | na | 0.0903 | 0.5748 | 0.1960 | **0.0073** |
| vDG | CD | UNEX | EC | CD | EXP | -0.0394 | 0.9993 | na | na | 0.0330 | >0.999 | 0.1260 | 0.6452 |
| vDG | CD | UNEX | EcrhC | CD | EXP | 0.0124 | >0.999 | na | na | 0.1030 | 0.2657 | 0.1740 | **0.0462** |
| vDG | CD | UNEX | vSb | CD | EXP | -0.0440 | 0.9939 | na | na | 0.0636 | 0.9919 | 0.1340 | 0.4944 |
| vDG | CD | UNEX | vDG | CD | EXP | -0.0093 | >0.999 | na | na | -0.0114 | >0.999 | -0.0227 | >0.999 |
| vDG | CD | UNEX | dSb | WD | UNEX | -0.0682 | 0.4860 | na | na | 0.1130 | 0.1550 | 0.1660 | 0.1276 |
| vDG | CD | UNEX | EC | WD | UNEX | -0.0678 | 0.4995 | na | na | 0.0228 | >0.999 | 0.1230 | 0.7850 |
| vDG | CD | UNEX | EcrhC | WD | UNEX | -0.0272 | >0.999 | na | na | 0.0926 | 0.6032 | 0.2180 | **0.0019** |
| vDG | CD | UNEX | vDG | WD | UNEX | -0.0428 | 0.9982 | na | na | -0.0059 | >0.999 | -0.0700 | >0.999 |
| vDG | CD | UNEX | vSb | WD | UNEX | -0.0817 | 0.1162 | na | na | 0.0632 | 0.9963 | 0.1120 | 0.9129 |
| vDG | CD | UNEX | dSb | WD | EXP | -0.0451 | 0.9906 | na | na | 0.1130 | 0.1022 | 0.2190 | **0.0007** |
| vDG | CD | UNEX | EC | WD | EXP | -0.0437 | 0.9945 | na | na | 0.0140 | >0.999 | 0.1010 | 0.9659 |
| vDG | CD | UNEX | EcrhC | WD | EXP | -0.0035 | >0.999 | na | na | 0.0544 | 0.9996 | 0.1140 | 0.8519 |
| vDG | CD | UNEX | vDG | WD | EXP | -0.0296 | >0.999 | na | na | -0.0198 | >0.999 | -0.0517 | >0.999 |
| vDG | CD | UNEX | vSb | WD | EXP | -0.0530 | 0.9095 | na | na | 0.0574 | 0.9988 | 0.1480 | 0.2610 |
| vDG | CD | EXP | dSb | CD | UNEX | -0.0338 | >0.999 | na | na | 0.1200 | **0.0491** | 0.2310 | **0.0002** |
| vDG | CD | EXP | EC | CD | UNEX | -0.0294 | >0.999 | na | na | 0.0349 | >0.999 | 0.1460 | 0.2888 |
| vDG | CD | EXP | EcrhC | CD | UNEX | 0.0231 | >0.999 | na | na | 0.1060 | 0.2102 | 0.2220 | **0.0006** |
| vDG | CD | EXP | vSb | CD | UNEX | -0.0442 | 0.9932 | na | na | 0.0714 | 0.9541 | 0.1700 | 0.0652 |
| vDG | CD | EXP | dSb | WD | UNEX | -0.0589 | 0.7469 | na | na | 0.1250 | **0.0307** | 0.1890 | **0.0139** |
| vDG | CD | EXP | EC | WD | UNEX | -0.0585 | 0.7593 | na | na | 0.0342 | >0.999 | 0.1450 | 0.2978 |
| vDG | CD | EXP | EcrhC | WD | UNEX | -0.0179 | >0.999 | na | na | 0.1040 | 0.2362 | 0.2400 | **<0.0001** |
| vDG | CD | EXP | vDG | WD | UNEX | -0.0335 | >0.999 | na | na | 0.0056 | >0.999 | -0.0473 | >0.999 |
| vDG | CD | EXP | vSb | WD | UNEX | -0.0724 | 0.2550 | na | na | 0.0747 | 0.9194 | 0.1350 | 0.4795 |
| vDG | CD | EXP | dSb | CD | EXP | -0.0298 | >0.999 | na | na | 0.1020 | 0.2036 | 0.2190 | **0.0003** |
| vDG | CD | EXP | EC | CD | EXP | -0.0301 | >0.999 | na | na | 0.0444 | >0.999 | 0.1490 | 0.1738 |
| vDG | CD | EXP | EcrhC | CD | EXP | 0.0217 | >0.999 | na | na | 0.1140 | 0.0596 | 0.1970 | **0.0031** |
| vDG | CD | EXP | vSb | CD | EXP | -0.0347 | 0.9999 | na | na | 0.0751 | 0.8669 | 0.1570 | 0.1022 |
| vDG | CD | EXP | dSb | WD | EXP | -0.0358 | 0.9998 | na | na | 0.1250 | **0.0160** | 0.2420 | **<0.0001** |
| vDG | CD | EXP | EC | WD | EXP | -0.0344 | >0.999 | na | na | 0.0254 | >0.999 | 0.1230 | 0.6084 |
| vDG | CD | EXP | EcrhC | WD | EXP | 0.0058 | >0.999 | na | na | 0.0658 | 0.9742 | 0.1360 | 0.3547 |
| vDG | CD | EXP | vDG | WD | EXP | -0.0203 | >0.999 | na | na | -0.0084 | >0.999 | -0.0290 | >0.999 |
| vDG | CD | EXP | vSb | WD | EXP | -0.0437 | 0.9894 | na | na | 0.0689 | 0.9516 | 0.1710 | **0.0351** |
| EcrhC | CD | UNEX | dSb | CD | UNEX | -0.0569 | 0.8689 | na | na | 0.0149 | >0.999 | 0.0097 | >0.999 |
| EcrhC | CD | UNEX | EC | CD | UNEX | -0.0525 | 0.9479 | na | na | -0.0706 | 0.9766 | -0.0755 | >0.999 |
| EcrhC | CD | UNEX | vSb | CD | UNEX | -0.0673 | 0.5168 | na | na | -0.0341 | >0.999 | -0.0520 | >0.999 |
| EcrhC | CD | UNEX | dSb | CD | EXP | -0.0529 | 0.9102 | na | na | -0.0038 | >0.999 | -0.0031 | >0.999 |
| EcrhC | CD | UNEX | EC | CD | EXP | -0.0531 | 0.9058 | na | na | -0.0610 | 0.9962 | -0.0726 | >0.999 |
| EcrhC | CD | UNEX | EcrhC | CD | EXP | -0.0014 | >0.999 | na | na | 0.0085 | >0.999 | -0.0248 | >0.999 |
| EcrhC | CD | UNEX | vSb | CD | EXP | -0.0578 | 0.7833 | na | na | -0.0304 | >0.999 | -0.0647 | >0.999 |
| EcrhC | CD | UNEX | dSb | WD | UNEX | -0.0819 | 0.1125 | na | na | 0.0191 | >0.999 | -0.0329 | >0.999 |
| EcrhC | CD | UNEX | EC | WD | UNEX | -0.0816 | 0.1180 | na | na | -0.0713 | 0.9730 | -0.0761 | >0.999 |
| EcrhC | CD | UNEX | EcrhC | WD | UNEX | -0.0410 | 0.9992 | na | na | -0.0014 | >0.999 | 0.0186 | >0.999 |
| EcrhC | CD | UNEX | vSb | WD | UNEX | -0.0955 | **0.0140** | na | na | -0.0308 | >0.999 | -0.0866 | 0.9986 |
| EcrhC | CD | UNEX | dSb | WD | EXP | -0.0589 | 0.7453 | na | na | 0.0193 | >0.999 | 0.0201 | >0.999 |
| EcrhC | CD | UNEX | EC | WD | EXP | -0.0575 | 0.7922 | na | na | -0.0801 | 0.8295 | -0.0984 | 0.9749 |
| EcrhC | CD | UNEX | EcrhC | WD | EXP | -0.0173 | >0.999 | na | na | -0.0397 | >0.999 | -0.0852 | 0.9978 |
| EcrhC | CD | UNEX | vSb | WD | EXP | -0.0667 | 0.4449 | na | na | -0.0366 | >0.999 | -0.0509 | >0.999 |
| EcrhC | CD | EXP | dSb | CD | UNEX | -0.0555 | 0.8517 | na | na | 0.0064 | >0.999 | 0.0344 | >0.999 |
| EcrhC | CD | EXP | EC | CD | UNEX | -0.0511 | 0.9407 | na | na | -0.0791 | 0.8492 | -0.0508 | >0.999 |
| EcrhC | CD | EXP | vSb | CD | UNEX | -0.0659 | 0.4758 | na | na | -0.0426 | >0.999 | -0.0273 | >0.999 |
| EcrhC | CD | EXP | dSb | CD | EXP | -0.0515 | 0.8957 | na | na | -0.0123 | >0.999 | 0.0217 | >0.999 |
| EcrhC | CD | EXP | EC | CD | EXP | -0.0517 | 0.8907 | na | na | -0.0695 | 0.9449 | -0.0478 | >0.999 |
| EcrhC | CD | EXP | vSb | CD | EXP | -0.0564 | 0.7530 | na | na | -0.0389 | >0.999 | -0.0400 | >0.999 |
| EcrhC | CD | EXP | Ecrh | WD | UNEX | -0.0396 | 0.9992 | na | na | -0.0099 | >0.999 | 0.0434 | >0.999 |
| EcrhC | CD | EXP | dSb | WD | UNEX | -0.0805 | 0.0900 | na | na | 0.0106 | >0.999 | -0.0082 | >0.999 |
| EcrhC | CD | EXP | EC | WD | UNEX | -0.0802 | 0.0948 | na | na | -0.0798 | 0.8358 | -0.0514 | >0.999 |
| EcrhC | CD | EXP | vSb | WD | UNEX | -0.0941 | **0.0096** | na | na | -0.0393 | >0.999 | -0.0619 | >0.999 |
| EcrhC | CD | EXP | Ecrh | WD | EXP | -0.0159 | >0.999 | na | na | -0.0482 | >0.999 | -0.0605 | >0.999 |
| EcrhC | CD | EXP | dSb | WD | EXP | -0.0575 | 0.7114 | na | na | 0.0108 | >0.999 | 0.0449 | >0.999 |
| EcrhC | CD | EXP | EC | WD | EXP | -0.0561 | 0.7628 | na | na | -0.0886 | 0.5212 | -0.0736 | 0.9998 |
| EcrhC | CD | EXP | vSb | WD | EXP | -0.0653 | 0.3978 | na | na | -0.0451 | >0.999 | -0.0262 | >0.999 |
| EC | CD | UNEX | dSb | CD | UNEX | -0.0043 | >0.999 | na | na | 0.0855 | 0.7796 | 0.0852 | 0.9990 |
| EC | CD | UNEX | vSb | CD | UNEX | -0.0148 | >0.999 | na | na | 0.0365 | >0.999 | 0.0235 | >0.999 |
| EC | CD | UNEX | dSb | CD | EXP | -0.0004 | >0.999 | na | na | 0.0668 | 0.9821 | 0.0724 | >0.999 |
| EC | CD | UNEX | EC | CD | EXP | -0.0006 | >0.999 | na | na | 0.0096 | >0.999 | 0.0029 | >0.999 |
| EC | CD | UNEX | vSb | CD | EXP | -0.0053 | >0.999 | na | na | 0.0402 | >0.999 | 0.0108 | >0.999 |
| EC | CD | UNEX | dSb | WD | UNEX | -0.0294 | >0.999 | na | na | 0.0896 | 0.6811 | 0.0426 | >0.999 |
| EC | CD | UNEX | EC | WD | UNEX | -0.0291 | >0.999 | na | na | -0.0007 | >0.999 | -0.0006 | >0.999 |
| EC | CD | UNEX | vSb | WD | UNEX | -0.0430 | 0.9980 | na | na | 0.0398 | >0.999 | -0.0111 | >0.999 |
| EC | CD | UNEX | dSb | WD | EXP | -0.0064 | >0.999 | na | na | 0.0899 | 0.5844 | 0.0956 | 0.9837 |
| EC | CD | UNEX | EC | WD | EXP | -0.0050 | >0.999 | na | na | -0.0095 | >0.999 | -0.0229 | >0.999 |
| EC | CD | UNEX | vSb | WD | EXP | -0.0142 | >0.999 | na | na | 0.0340 | >0.999 | 0.0246 | >0.999 |
| EC | CD | EXP | dSb | CD | UNEX | -0.0037 | >0.999 | na | na | 0.0760 | 0.9015 | 0.0823 | 0.9989 |
| EC | CD | EXP | vSb | CD | UNEX | -0.0142 | >0.999 | na | na | 0.0269 | >0.999 | 0.0206 | >0.999 |
| EC | CD | EXP | dSb | CD | EXP | 0.0002 | >0.999 | na | na | 0.0573 | 0.9975 | 0.0695 | >0.999 |
| EC | CD | EXP | vSb | CD | EXP | -0.0046 | >0.999 | na | na | 0.0306 | >0.999 | 0.0079 | >0.999 |
| EC | CD | EXP | dSb | WD | UNEX | -0.0288 | >0.999 | na | na | 0.0801 | 0.8300 | 0.0397 | >0.999 |
| EC | CD | EXP | EC | WD | UNEX | -0.0284 | >0.999 | na | na | -0.0103 | >0.999 | -0.0035 | >0.999 |
| EC | CD | EXP | vSb | WD | UNEX | -0.0424 | 0.9969 | na | na | 0.0302 | >0.999 | -0.0140 | >0.999 |
| EC | CD | EXP | dSb | WD | EXP | -0.0058 | >0.999 | na | na | 0.0804 | 0.7504 | 0.0927 | 0.9812 |
| EC | CD | EXP | EC | WD | EXP | -0.0044 | >0.999 | na | na | -0.0191 | >0.999 | -0.0258 | >0.999 |
| EC | CD | EXP | vSb | WD | EXP | -0.0136 | >0.999 | na | na | 0.0244 | >0.999 | 0.0217 | >0.999 |
| dSb | CD | UNEX | dSb | CD | EXP | 0.0039 | >0.999 | na | na | -0.0187 | >0.999 | -0.0128 | >0.999 |
| dSb | CD | UNEX | vSb | CD | EXP | -0.0009 | >0.999 | na | na | -0.0454 | >0.999 | -0.0744 | 0.9999 |
| dSb | CD | UNEX | vSb | CD | UNEX | -0.0105 | >0.999 | na | na | -0.0490 | >0.999 | -0.0617 | >0.999 |
| dSb | CD | UNEX | dSb | WD | EXP | -0.0021 | >0.999 | na | na | 0.0044 | >0.999 | 0.0104 | >0.999 |
| dSb | CD | UNEX | dSb | WD | UNEX | -0.0251 | >0.999 | na | na | 0.0041 | >0.999 | -0.0426 | >0.999 |
| dSb | CD | UNEX | vSb | WD | EXP | -0.0099 | >0.999 | na | na | -0.0516 | 0.9999 | -0.0606 | >0.999 |
| dSb | CD | UNEX | vSb | WD | UNEX | -0.0386 | 0.9998 | na | na | -0.0458 | >0.999 | -0.0963 | 0.9900 |
| dSb | CD | EXP | vSb | CD | EXP | -0.0049 | >0.999 | na | na | -0.0267 | >0.999 | -0.0616 | >0.999 |
| dSb | CD | EXP | vSb | CD | UNEX | -0.0144 | >0.999 | na | na | -0.0303 | >0.999 | -0.0489 | >0.999 |
| dSb | CD | EXP | dSb | WD | EXP | -0.0060 | >0.999 | na | na | 0.0231 | >0.999 | 0.0232 | >0.999 |
| dSb | CD | EXP | dSb | WD | UNEX | -0.0290 | >0.999 | na | na | 0.0228 | >0.999 | -0.0298 | >0.999 |
| dSb | CD | EXP | vSb | WD | EXP | -0.0138 | >0.999 | na | na | -0.0329 | >0.999 | -0.0478 | >0.999 |
| dSb | CD | EXP | vSb | WD | UNEX | -0.0426 | 0.9966 | na | na | -0.0271 | >0.999 | -0.0835 | 0.9985 |
| vSb | CD | UNEX | vSb | CD | EXP | 0.0096 | >0.999 | na | na | 0.0037 | >0.999 | -0.0127 | >0.999 |
| vSb | CD | UNEX | vSb | WD | UNEX | -0.0282 | >0.999 | na | na | 0.0033 | >0.999 | -0.0346 | >0.999 |
| vSb | CD | UNEX | vSb | WD | EXP | 0.0006 | >0.999 | na | na | -0.0025 | >0.999 | 0.0011 | >0.999 |
| vSb | CD | EXP | vSb | WD | EXP | -0.0090 | >0.999 | na | na | -0.0062 | >0.999 | 0.0138 | >0.999 |
| vSb | CD | EXP | vSb | WD | UNEX | -0.0377 | 0.9997 | na | na | -0.0004 | >0.999 | -0.0219 | >0.999 |

***Supplemental Table 11*.** Statistical details of microbiome analysis.

|  | ***Lachnospiraceae* NK4A136** Relative abundance | **PCA** | **Richness** Chao1 | **Diversity** Shannon | ***Prevotellacea*** Relative abundance |
| --- | --- | --- | --- | --- | --- |
| Source of Variation | F Statistic, post hoc | t test | F Statistic, post hoc | F Statistic, post hoc | F Statistic, post hoc |
| Stress:Diet | **F(1, 34) = 8.9, *p =* 0.005** | -- | F(1, 34) = 0.003, *p =*0.9 | F(1, 34) = 0.15, *p =* 0.07 | F(1, 34) = 0.29, *p =* 0.59 |
| STRESS | **F(1, 34) = 5.8, *p =* 0.021** | **t(34), *p* < 0.001** | F(1, 34) = 0.03, *p =* 0.8 | F(1, 34) = 1.7, *p =* 0.19 | F(1, 34) = 0.02, *p =* 0.87 |
| DIET | **F(1, 34) = 9.9, *p =* 0.003** | t(34), *p* = 0.78 | F(1, 34) = 2.5, *p =* 0.12 | F(1, 34) = 0.007, *p =* 0.93 | **F(1, 34) = 9.6, *p =* 0.003** |
| CDU vs WDU | ***p =* 0.0005** |  |  |  |  |
| CDU vs CDE | ***p =* 0.0048** |  |  |  |  |
| CDU vs WDE | ***p =* 0.0019** |  |  |  |  |
| WDU vs CDE | *p =* 0.954 |  |  |  |  |
| WDU vs WDE | *p =* 0.9743 |  |  |  |  |
| CDE vs WDE | *p =* 0.9994 |  |  |  |  |

***Supplementary Table 12*.** Statistical Information of peripheral cytokines

|  | **IL-1α** | **IL-10** | **GM-CSF** | **IL-33** |
| --- | --- | --- | --- | --- |
| Factor | F Statistic, post-hoc | F Statistic, post-hoc | F Statistic, post-hoc | F Statistic, post-hoc |
| Stress:Diet | **F(1, 17) = 11.6, p = 0.003** | **F(1, 20) = 6.7, p = 0.017** | **F(1, 18) = 8.5, p = 0.009** | **F(1, 19) = 6.8, p = 0.01** |
| Stress | **F(1, 17) = 10.5, p = 0.004** | F(1, 20) = 3.4, p = 0.07 | F(1, 18) = 0.8, p = 0.3 | F(1, 19) = 0.003, p = 0.9 |
| Diet | F(1, 17) = 1.9, p = 0.183 | F(1, 20) = 1.5, p = 0.2 | **F(1, 18) = 13.9, p = 0.001** | F(1, 19) = 2.1, p = 0.15 |
| CDU vs WDU | **p = 0.014** | p = 0.7784 | **p = 0.0009** | **p = 0.0488** |
| CDU vs CDE | **p = 0.0008** | p = 0.9542 | p = 0.0755 | p = 0.2835 |
| CDU vs WDE | **p = 0.018** | p = 0.1588 | **p = 0.0188** | p = 0.7205 |
| WDU vs CDE | p = 0.583 | p = 0.9719 | p = 0.2346 | p = 0.7368 |
| WDU vs WDE | p = 0.9994 | **p = 0.0242** | p = 0.4764 | p = 0.2791 |
| CDE vs WDE | p = 0.5156 | p = 0.0597 | p = 0.9386 | p = 0.8392 |

***Supplementary Table 13*.** Statistical Information of supplemental peripheral cytokines

|  | **IL-6** | **IL-12p70** | **IFN-ϒ** | **CCL-2** | **IL-18** | **TNF-α** | **IL-17A** |
| --- | --- | --- | --- | --- | --- | --- | --- |
| Factor | F Statistic, post-hoc | F Statistic, post-hoc | F Statistic, post-hoc | F Statistic, post-hoc | F Statistic, post-hoc | F Statistic, post-hoc | F Statistic, post-hoc |
| Stress:Diet | F(1, 14) = 1.3, p = 0.2 | **F(1, 20) = 6.2, p = 0.02** | F(1, 19) = 3.2, p = 0.08 | F(1, 21) = 0.17, p = 0.68 | F(1, 18) = 3.2, p = 0.08 | F(1, 23) = 0.007, p = 0.9 | F(1, 20) = 2.4, p = 0.13 |
| Stress | F(1, 14) = 1.9, p = 0.18 | F(1, 20) = 2.3,p = 0.13 | F(1, 19) = 2.3, p = 0.14 | F(1, 21) = 0.27, p = 0.6 | F(1, 18) = 0.4, p = 0.51 | F(1, 23) = 0.004, p = 0.9 | F(1, 20) = 0.05, p = 0.81 |
| Diet | **F(1, 14) = 8.4, p = 0.01** | F(1, 20) = 2.5, p = 0.12 | F(1, 19) = 0.6, p = 0.44 | F(1, 21) = 0.33, p = 0.57 | F(1, 18) = 0.09, p = 0.76 | F(1, 23) = 0.8, p = 0.3 | F(1, 20) = 0.07, p = 0.78 |

***Supplementary Table 14A*.** Statistical Information of gut brain correlations with microglia morphology Figures 5 and Supplemental Figure 11

|  | **Objects** | **Lacunarity** | **Density** | **Span Ratio** | **Max Span Across Hull** | **Area** | **Perimeter** | **Circularity** | **Mean Radius** |
| --- | --- | --- | --- | --- | --- | --- | --- | --- | --- |
| **Objects** |  | *r =* -0.72, p = 0.002, FDR =5E-3 | *r =* 0.45, *p =* 0.082, FDR = 0.13 | *r =* -0.34, p = 0.196, FDR = 0.25 | *r =* -0.36, *p =* 0.161, FDR = 0.24 | *r =* -0.27, *p =* 0.298, FDR = 0.34 | *r =* -0.32, *p =* 0.216, FDR = 0.27 | *r =* 0.582, *p =* 0.019, FDR = 0.21 | *r =* -0.34, *p =* 0.196, FDR = 0.27 |
| **Lacunarity** | *r =* -0.72 p = 0.002, FDR = 0.02 |  | *r =* -0.89, p = 1.E-05, FDR = 5E-5 | *r =* -0.18, p = 0.497, FDR = 0.5 | *r =* 0.791, *p =* 4E-4, FDR = 1E-3 | *r =* 0.723, *p =* 0.002, FDR = 7E-3 | *r =* 0.764, *p =* 8E-4, FDR = 3E-3 | *r =* -0.05, *p =* 0.856, FDR = 0.85 | *r =* 0.767, *p =* 8E-4 FDR = 3E-3 |
| **Density** | *r =* 0.45, p = 0.082, FDR = 0.23 | *r =* -0.89, p = 1.E-05, FDR = 2E-4 |  | *r =* 0.382, p = 0.144, FDR = 0.23 | *r =* -0.89, *p =* 1E-5,  FDR = 4E5 | *r =* -0.84, *p =* 8.E-05, FDR = 3E-4 | *r =* -0.87, *p =* 2E-05, FDR = 9E-5 | *r =* -0.17, *p =* 0.512, FDR = 0.56 | *r =* -0.89, *p =* 1E-05, FDR = 4E-5 |
| **Span Ratio** | *r =* -0.34, p = 0.196, FDR = 0.35 | *r =* -0.18, p = 0.497, FDR = 0.407 | *r =* 0.382, p = 0.144, FDR = 0.20 |  | *r =* -0.46, *p =* 0.073, FDR = 0.15 | *r =* -0.52, *p =* 0.04, FDR = 0.1 | *r =* -0.49, *p =* 0.052,  FDR = 0.13 | *r =* -0.91, *p =* 3E-06, FDR = 0 | *r =* -0.5,p = 0.051, FDR = 0.12 |
| **Max Span Across Hull** | *r =* -0.36, p = 0.161, FDR = 0.35 | *r =* 0.791, p = 0.0004, FDR = 0.003 | *r =* -0.89, *p =* 1.E-05, FDR = 5E-5 | *r =* -0.46, p = 0.073, FDR = 0.13 |  | *r =* 0.985, *p =* 3E-10, FDR = 1E-9 | *r =* 0.997, *p =* 1E-12, FDR = 1E-11 | *r =* 0.279, *p =* 0.293, FDR = 0.5 | *r =* 0.994, *p =* 1E-11, FDR = 8E-11 |
| **Area** | *r =* -0.27, p = 0.298, FDR = 0.46 | *r =* 0.723, p = 0.002, FDR = 0.005 | *r =* -0.84, p = 8E-05, FDR = 3E-4 | *r =* -0.52, p = 0.04, FDR = 0.11 | *r =* 0.985, *p =* 3E-10, FDR = 1E-9 |  | *r =* 0.994, *p =* 1E-11, FDR = 5E-11 | *r =* 0.358, *p =* 0.172, FDR = 0.40 | *r =* 0.988, *p =* 1E-10, FDR = 6E-10 |
| **Perimeter** | *r =* -0.32, p = 0.216, FDR = 0.36 | *r =* 0.764, p = 1E-05, FDR = 0.003 | *r =* -0.87, *p =* 2.E-05, FDR = 9E-5 | *r =* -0.49, p = 0.052, FDR = 0.1 | *r =* 0.997, *p =* 1E-12, FDR = 2E-11 | *r =* 0.994, *p =* 1E-11, FDR = 1E-10 |  | *r =* 0.32, *p =* 0.225, FDR = 0.45 | *r =* 0.997, *p =* 1E-12, FDR = 2E-11 |
| **Circularity** | *r =*0.582, p = 0.019, FDR = 0.08 | *r =* -0.05, p = 0.856, FDR = 0.6 | *r =* -0.17, *p =* 0.512, FDR = 0.43 | *r =* -0.91, p = 3E-06, FDR = 0 | *r =* 0.279, *p =* 0.293, FDR = 0.32 | *r =* 0.358, *p =* 0.172, FDR = 0.24 | *r =* 0.32,p = 0.225, FDR = 0.27 |  | *r =* 0.314, *p =* 0.234, FDR = 0.3 |
| **Mean Radius** | *r =* -0.34, p = 0.196, FDR =0.35 | *r =* 0.767, p = 0.0008, FDR = 0.003 | *r =* -0.89, *p =* 1E-05,  FDR = 5E-5 | *r =* -0.5, p = 0.05, FDR = 0.1 | *r =* 0.994, *p =* 1E-11, FDR = 8E-11 | *r =* 0.988, *p =* 1E-10, FDR = 1E-9 | *r =* 0.997, *p =* 1E-12, FDR = 1E-9 | *r =* 0.314, *p =* 0.234, FDR = 0.45 |  |
| **ODI** | *r =* 0.123, p = 0.648, FDR = 0.81 | *r =* -0.37 p = 0.158, FDR = 0.157 | *r =* 0.329, *p =* 0.212, FDR = 0.24 | *r =* 0.352, p = 0.18, FDR = 0.25 | *r =* -0.25, *p =* 0.343, FDR = 0.32 | *r =* -0.26, *p =* 0.32, FDR = 0.34 | *r =* -0.27, *p =* 0.304, FDR = 0.32 | *r =* -0.18, *p =* 0.483, FDR = 0.56 | *r =* -0.27, *p =* 0.293, FDR = 0.3 |
| **ICVF** | *r =* 0.171, p = 0.54, FDR = 0.72 | *r =* -0.38, *p =* 0.16, FDR = 0.157 | *r =* 0.26, *p =* 0.346, FDR = 0.31 | *r =* 0.389, p = 0.152, FDR = 0.23 | *r =* 0.003, *p =* 0.994, FDR = 0.8 | *r =* 0.028, *p =* 0.923, FDR = 0.74 | *r =* 0.017, *p =* 0.953, FDR = 0.763 | *r =* -0.27, *p =* 0.313, FDR = 0.45 | *r =* 0, p = 1, FDR = 0.8 |
| **ISVF** | *r =* 0.058, p = 0.83, FDR = 0.92 | *r =* 0.158, p = 0.556, FDR = 0.43 | *r =* -0.37, *p =* 0.158, FDR = 0.20 | *r =* -0.56, p = 0.025, FDR = 0.092 | *r =* 0.255, *p =* 0.337, FDR = 0.32 | *r =* 0.27, p = 0.309, FDR = 0.34 | *r =* 0.258, *p =* 0.331, FDR = 0.32 | *r =* 0.361, *p =* 0.169, FDR = 0.4 | *r =* 0.276, *p =* 0.298, FDR = 0.3 |
| **LJ** | *r =* 0.352, p = 0.18, FDR = 0.35 | *r =* -0.12, *p =* 0.632, FDR = 0.46 | *r =* -0.02, *p =* 0.917, FDR = 0.74 | *r =* -0.23, p = 0.372, FDR = 0.392 | *r =* 0.076, *p =* 0.779, FDR = 0.65 | *r =* 0.138, *p =* 0.609, FDR = 0.54 | *r =* 0.105, *p =* 0.696, FDR = 0.585 | *r =* 0.235, *p =* 0.379, FDR = 0.5 | *r =* 0.108, *p =* 0.688, FDR = 0.6 |
| **FA** | *r =* 0.07, p = 0.796, FDR = 0.92 | *r =* -0.34, *p =* 0.191, FDR = 0.17 | *r =* 0.3, *p =* 0.258, FDR = 0.24 | *r =* 0.505, p = 0.047, FDR = 0.104 | *r =* -0.15, *p =* 0.571, FDR = 0.5 | *r =* -0.11, p = 0.68, FDR = 0.57 | *r =* -0.14, p = 0.601, FDR = 0.532 | *r =* -0.4,p = 0.12, FDR = 0.36 | *r =* -0.15, *p =* 0.578, FDR = 0.5 |
| **MD** | *r =* -0.004, p = 0.989, FDR = 0.94 | *r =* 0.525, *p =* 0.038, FDR = 0.05 | *r =* -0.55, *p =* 0.028, FDR = 0.06 | *r =* -0.67, p = 0.005, FDR = 0.023 | *r =* 0.465, *p =* 0.071, FDR = 0.15 | *r =* 0.473, *p =* 0.065, FDR = 0.14 | *r =* 0.473, *p =* 0.065, FDR = 0.137 | *r =* 0.481, *p =* 0.06, FDR = 0.2 | *r =* 0.467, *p =* 0.069, FDR = 0.1 |
| **RD** | *r =* -0.009, p = 0.975, FDR = 0.94 | *r =* 0.522, p = 0.039, FDR = 0.05 | *r =* -0.54, *p =* 0.03, FDR = 0.06 | *r =* -0.67, p = 0.004, FDR = 0.02 | *r =* 0.453, *p =* 0.079, FDR = 0.15 | *r =* 0.453, *p =* 0.079, FDR = 0.15 | *r =* 0.459, *p =* 0.075, FDR = 0.14 | *r =* 0.479, *p =* 0.061, FDR = 0.21 | *r =* 0.454, *p =* 0.078, FDR = 0.1 |
| **AD** | *r =* 0.033, p = 0.901, FDR = 0.94 | *r =* 0.5, p = 0.05, FDR = 0.06 | *r =* -0.56, *p =* 0.024, FDR = 0.06 | *r =* -0.71, p = 0.002, FDR = 0.02 | *r =* 0.498, *p =* 0.051, FDR = 0.14 | *r =* 0.52, p = 0.04, FDR = 0.1 | *r =* 0.516, *p =* 0.042, FDR = 0.119 | *r =* 0.52, *p =* 0.04, FDR = 0.21 | *r =* 0.512, *p =* 0.044, FDR = 0.1 |
| ***Rumino coccus*** | *r =* -0.63, p = 0.01, FDR = 0.06 | *r =* 0.441, p = 0.088, FDR = 0.10 | *r =* -0.32, *p =* 0.221, FDR = 0.24 | *r =* 0.155, p = 0.563, FDR = 0.52 | *r =* 0.258, *p =* 0.331, FDR = 0.32 | *r =* 0.252, *p =* 0.343, FDR = 0.34 | *r =* 0.252, *p =* 0.343, FDR = 0.32 | *r =* -0.26, *p =* 0.326, FDR = 0.46 | *r =* 0.27, p = 0.309, FDR = 0.3 |
| ***Muri baculaceae*** | *r =* -0.51, p = 0.042, FDR = 0.14 | *r =* 0.338, *p =* 0.2, FDR = 0.17 | *r =* -0.3, *p =* 0.253, FDR 0.241 | *r =* 0.12, p = 0.656, FDR = 0.59 | *r =* 0.361, *p =* 0.169, FDR = 0.23 | *r =* 0.376, *p =* 0.151, FDR = 0.24 | *r =* 0.364, *p =* 0.165, FDR = 0.253 | *r =* -0.12, *p =* 0.656, FDR = 0.69 | *r =* 0.391, *p =* 0.135, FDR = 0.2 |
| ***Ls NK4A136*** | *r =* 0.211, p = 0.429, FDR = 0.61 | *r =* -0.56, p = 0.025, FDR = 0.05 | *r =* 0.429, *p =* 0.098, FDR = 0.15 | *r =* 0.282, p = 0.288, FDR = 0.32 | *r =* -0.34, *p =* 0.196, FDR = 0.25 | *r =* -0.3, p = 0.248, FDR = 0.32 | *r =* -0.33, *p =* 0.204, FDR = 0.271 | *r =* -0.29, *p =* 0.263, FDR = 0.45 | *r =* -0.32, *p =* 0.221, FDR = 0.3 |
| ***Copro coccus*** | *r =* -0.61, *p =* 0.013, FDR = 0.07 | *r =* 0.52, p = 0.04, FDR = 0.05 | *r =* -0.3, *p =* 0.258, FDR = 0.24 | *r =* 0.079, p = 0.771, FDR = 0.7 | *r =* 0.279, *p =* 0.293, FDR = 0.32 | *r =* 0.205, *p =* 0.442, FDR = 0.41 | *r =* 0.252, *p =* 0.343, FDR = 0.32 | *r =* -0.21, *p =* 0.416, FDR = 0.51 | *r =* 0.247, *p =* 0.355, FDR = 0.3 |
| **Bilophila** | *r =* 0.838,p = 1E-4,  FDR = 2E-3 | *r =* -0.65, p = 0.007, FDR = 0.01 | *r =* 0.502, *p =* 0.049, FDR = 0.09 | *r =* -0.3, p = 0.243, FDR = 0.29 | *r =* -0.42, *p =* 0.101, FDR = 0.17 | *r =* -0.36, *p =* 0.169, FDR = 0.24 | *r =* -0.39, *p =* 0.128, FDR = 0.217 | *r =* 0.535, *p =* 0.034, FDR = 0.21 | *r =* -0.41, *p =* 0.111, FDR = 0.2 |

***Supplementary Table 14B*.** Statistical Information of gut brain correlations with MRI measures in Figures 5 and Supplemental Figure 11.

|  | **ODI** | **ICVF** | **ISVF** | **LJ** | **FA** | **MD** | **RD** | **AD** |
| --- | --- | --- | --- | --- | --- | --- | --- | --- |
| **Objects** | *r =* 0.123, *p =* 0.648, FDR = 0.8 | *r =* 0.171, p = 0.54, FDR = 0.75 | *r =* 0.058, *p =* 0.83, FDR = 0.98 | *r =* 0.352, *p =* 0.18, FDR = 0.7 | *r =* 0.07, p = 0.796, FDR = 0.78 | *r =* -0.004, *p =* 0.989, FDR = 0.8 | *r =* -0.009, *p =* 0.975, FDR = 0.8 | *r =* 0.033, *p =* 0.901, FDR = 0.80 |
| **Lacunarity** | *r =* -0.37, *p =* 0.158, FDR = 0.6 | *r =* -0.38, *p =* 0.16, FDR = 0.42 | *r =* 0.158, *p =* 0.556, FDR = 0.87 | *r =* -0.12, *p =* 0.632, FDR = 0.9 | *r =* -0.34, *p =* 0.191, FDR = 0.47 | *r =* 0.525, *p =* 0.038, FDR = 0.08 | *r =* 0.522, *p =* 0.039, FDR = 0.1 | *r =* 0.5, p = 0.05, FDR = 0.07 |
| **Density** | *r =* 0.329, *p =* 0.212, FDR = 0.6 | *r =* 0.26, p = 0.346, FDR = 0.64 | *r =* -0.37, *p =* 0.158, FDR = 0.62 | *r =* -0.02, *p =* 0.917, FDR = 0.9 | *r =* 0.3, p = 0.258, FDR = 0.47 | *r =* -0.55, *p =* 0.028, FDR = 0.07 | *r =* -0.54, *p =* 0.03, FDR = 0.07 | *r =* -0.56, *p =* 0.024, FDR = 0.07 |
| **Span Ratio** | *r =* 0.352, *p =* 0.18, FDR = 0.6 | *r =* 0.389, *p =* 0.152, FDR = 0.42 | *r =* -0.56, *p =* 0.025, FDR = 0.48 | *r =* -0.23, *p =* 0.372, FDR = 0.7 | *r =* 0.505, *p =* 0.047, FDR = 0.16 | *r =* -0.67, *p =* 0.005, FDR = 0.03 | *r =* -0.67, *p =* 0.004, FDR = 0.03 | *r =* -0.71, *p =* 0.002, FDR = 0.015 |
| **Max Span Across Hull** | *r =* -0.25, *p =* 0.343, FDR = 0.6 | *r =* 0.003, *p =* 0.994, FDR = 1 | *r =* 0.255, *p =* 0.337, FDR = 0.73 | *r =* 0.076, *p =* 0.779, FDR = 0.9 | *r =* -0.15, *p =* 0.571, FDR = 0.7 | *r =* 0.465, *p =* 0.071, FDR = 0.09 | *r =* 0.453, *p =* 0.079, FDR = 0.1 | *r =* 0.498, *p =* 0.051, FDR = 0.07 |
| **Area** | *r =* -0.26, *p =* 0.32, FDR = 0.6 | *r =* 0.028, *p =* 0.923, FDR = 1 | *r =* 0.27, *p =* 0.309, FDR = 0.73 | *r =* 0.138, *p =* 0.609, FDR = 0.9 | *r =* -0.11, *p =* 0.68, FDR = 0.75 | *r =* 0.473, *p =* 0.065, FDR = 0.09 | *r =* 0.453, *p =* 0.079, FDR = 0.1 | *r =* 0.52, *p =* 0.04, FDR = 0.07 |
| **Perimeter** | *r =* -0.27, *p =* 0.304, FDR = 0.6 | *r =* 0.017, *p =* 0.953, FDR = 1 | *r =* 0.258, *p =* 0.331, FDR = 0.73 | *r =* 0.105, *p =* 0.696, FDR = 0.9 | *r =* -0.14, *p =* 0.601, FDR = 0.7 | *r =* 0.473, *p =* 0.065, FDR = 0.09 | *r =* 0.459, *p =* 0.075, FDR = 0.1 | *r =* 0.516, *p =* 0.042, FDR = 0.07 |
| **Circularity** | *r =* -0.18, *p =* 0.483, FDR = 0.7 | *r =* -0.27, *p =* 0.313, FDR = 0.64 | *r =* 0.361, *p =* 0.169, FDR = 0.62 | *r =* 0.235, *p =* 0.379, FDR = 0.7 | *r =* -0.4, p = 0.12, FDR = 0.36 | *r =* 0.481, *p =* 0.06, FDR = 0.09 | *r =* 0.479, *p =* 0.061, FDR = 0.1 | *r =* 0.52, *p =* 0.04, FDR = 0.07 |
| **Mean Radius** | *r =* -0.27, *p =* 0.293, FDR = 0.6 | *r =* 0, p = 1, FDR = 1 | *r =* 0.276, *p =* 0.298, FDR = 0.73 | *r =* 0.108, *p =* 0.688, FDR = 0.9 | *r =* -0.15, *p =* 0.578, FDR = 0.7 | *r =* 0.467, *p =* 0.069, FDR = 0.09 | *r =* 0.454, *p =* 0.078, FDR = 0.1 | *r =* 0.512, *p =* 0.044, FDR = 0.07 |
| **ODI** |  | *r =* 0.417, *p =* 0.122, FDR = 0.42 | *r =* -0.06, *p =* 0.822, FDR = 0.98 | *r =* 0.032, *p =* 0.908, FDR = 0.9 | *r =* 0.294, *p =* 0.268, FDR = 0.47 | *r =* -0.63, *p =* 0.009, FDR = 0.03 | *r =* -0.59, *p =* 0.015, FDR = 0.05 | *r =* -0.69, *p =* 0.003, FDR = 0.016 |
| **ICVF** | *r =* 0.417, *p =* 0.122, FDR = 0.6 |  | *r =* -0.18, *p =* 0.515, FDR = 0.87 | *r =* 0.364, *p =* 0.182, FDR = 0.7 | *r =* 0.942, *p =* 8E-7, FDR = 1E-5 | *r =* -0.62, p = 0.015, FDR = 0.04 | *r =* -0.63, *p =* 0.013, FDR = 0.05 | *r =* -0.53, *p =* 0.04, FDR = 0.07 |
| **ISVF** | *r =* -0.06, *p =* 0.822, FDR = 0.9 | *r =* -0.18, *p =* 0.515, FDR = 0.75 |  | *r =* 0.217, *p =* 0.416, FDR = 0.7 | *r =* -0.24, *p =* 0.366, FDR = 0.58 | *r =* 0.442, *p =* 0.086, FDR = 0.1 | *r =* 0.444, *p =* 0.085, FDR = 0.1 | *r =* 0.473, *p =* 0.065, FDR = 0.08 |
| **LJ** | *r =* 0.032, *p =* 0.908, FDR = 0.9 | *r =* 0.364, *p =* 0.182, FDR = 0.42 | *r =* 0.217, *p =* 0.416, FDR = 0.7 |  | *r =* 0.305, *p =* 0.248, FDR = 0.47 | *r =* 0.228, *p =* 0.392, FDR = 0.38 | *r =* 0.254, *p =* 0.338, FDR = 0.35 | *r =* 0.223, *p =* 0.401, FDR = 0.42 |
| **FA** | *r =* 0.294, *p =* 0.268, FDR = 0.6 | *r =* 0.942, *p =* 8E-7, FDR = 1E-5 | *r =* -0.24, *p =* 0.366, FDR = 0.73 | *r =* 0.305, *p =* 0.248, FDR = 0.7 |  | *r =* -0.64, *p =* 0.008, FDR = 0.03 | *r =* -0.66, *p =* 0.006, FDR = 0.03 | *r =* -0.57, *p =* 0.022, FDR = 0.07 |
| **MD** | *r =* -0.63, *p =* 0.009, FDR = 0.1 | *r =* -0.62, *p =* 0.015, FDR = 0.08 | *r =* 0.442, *p =* 0.086, FDR = 0.48 | *r =* 0.228, *p =* 0.392, FDR = 0.7 | *r =* -0.64, *p =* 0.008, FDR = 0.06 |  | *r =* 0.992, *p =* 1E-11, FDR = 2E-10 | *r =* 0.977, *p =* 1E-09, FDR = 2E-8 |
| **RD** | *r =* -0.59, *p =* 0.015, FDR = 0.1 | *r =* -0.63, *p =* 0.013, FDR = 0.08 | *r =* 0.444, *p =* 0.085, FDR = 0.48 | *r =* 0.254, *p =* 0.338, FDR = 0.7 | *r =* -0.66, *p =* 0.006, FDR = 0.06 | *r =* 0.992, *p =* 1E-11, FDR = 1E-10 |  | *r =* 0.956, *p =* 5.E-08, FDR = 4E-7 |
| **AD** | *r =* -0.69, *p =* 0.003, FDR = 0.08 | *r =* -0.53, *p =* 0.04, FDR = 0.17 | *r =* 0.473, *p =* 0.065, FDR = 0.48 | *r =* 0.223, *p =* 0.401, FDR = 0.7 | *r =* -0.57, *p =* 0.022, FDR = 0.09 | *r =* 0.977, *p =* 1E-09, FDR = 1E-8 | *r =* 0.956, *p =* 5E-08, FDR = 5E-7 |  |
| ***Rumino coccus*** | *r =* -0.07, *p =* 0.771, FDR = 0.8 | *r =* -0.07, *p =* 0.792, FDR = 1 | *r =* -0.01, *p =* 0.969, FDR = 1 | *r =* -0.27, *p =* 0.293, FDR = 0.7 | *r =* 0.082, *p =* 0.762, FDR = 0.8 | *r =* -0.18, *p =* 0.495, FDR = 0.46 | *r =* -0.19, *p =* 0.461, FDR = 0.45 | *r =* -0.15, *p =* 0.553, FDR = 0.55 |
| ***Muri baculaceae*** | *r =* 0.158, *p =* 0.556, FDR = 0.7 | *r =* 0.214, *p =* 0.442, FDR = 0.71 | *r =* -0.11,p = 0.672, FDR = 0.92 | *r =* -0.08, *p =* 0.746, FDR = 0.9 | *r =* 0.302, *p =* 0.253, FDR = 0.47 | *r =* -0.32, *p =* 0.222, FDR = 0.23 | *r =* -0.33, *p =* 0.198, FDR = 0.22 | *r =* -0.3, p = 0.244, FDR = 0.27 |
| ***Ls NK4A136*** | *r =* 0.229, *p =* 0.391, FDR = 0.6 | *r =* 0.628, *p =* 0.014, FDR = 0.08 | *r =* 0.126, *p =* 0.64, FDR = 0.92 | *r =* 0.267, *p =* 0.315, FDR = 0.7 | *r =* 0.617, *p =* 0.012, FDR = 0.06 | *r =* -0.43, *p =* 0.09, FDR = 0.1 | *r =* -0.44, *p =* 0.085, FDR = 0.1 | *r =* -0.4, p = 0.118, FDR = 0.14 |
| ***Coprococcus*** | *r =* -0.14, *p =* 0.586, FDR = 0.7 | *r =* -0.25, *p =* 0.367, FDR = 0.64 | *r =* -0.05, *p =* 0.847, FDR = 0.98 | *r =* -0.63, *p =* 0.01, FDR = 0.2 | *r =* -0.22, *p =* 0.391, FDR = 0.58 | *r =* -0.06, *p =* 0.811, FDR = 0.71 | *r =* -0.06, *p =* 0.819, FDR = 0.73 | *r =* -0.05, *p =* 0.828, FDR = 0.77 |
| ***Bilophila*** | *r =* 0.25, *p =* 0.349, FDR = 0.6 | *r =* -0.007, *p =* 0.984, FDR = 1 | *r =* -0.006, *p =* 0.986, FDR = 1 | *r =* 0.426, *p =* 0.101, FDR = 0.7 | *r =* -0.14, *p =* 0.586, FDR = 0.7 | *r =* 0.027, *p =* 0.919, FDR = 0.77 | *r =* 0.064, *p =* 0.811, FDR = 0.73 | *r =* 0.008, *p =* 0.975, FDR = 0.82 |

***Supplementary Table 14C*.** Statistical Information of gut brain correlations with microbiome measures in Figures 5 and Supplemental Figure 11.

|  | ***Rumino***  ***coccus*** | ***Muri***  ***baculaceae*** | ***Lachno***  ***spiraceae NK4A136*** | ***Copro***  ***coccus*** | **Bilophila** |
| --- | --- | --- | --- | --- | --- |
| **Objects** | *r =* -0.63, *p =* 0.01, FDR = 0.11 | *r =* -0.51,p = 0.042, FDR = 0.31 | *r =* 0.211, p = 0.429, FDR = 0.55 | *r =* -0.61, p = 0.013, FDR = 0.15 | *r =* 0.838, *p =* 1E-4, FDR = 0.002 |
| **Lacunarity** | *r =* 0.441,p = 0.088, FDR = 0.49 | *r =* 0.338,p = 0.2, FDR = 0.43 | *r =* -0.56,p = 0.025, FDR = 0.19 | *r =* 0.52,p = 0.04, FDR = 0.21 | *r =* -0.65,p = 0.007, FDR = 0.08 |
| **Density** | *r =* -0.32,p = 0.221, FDR = 0.63 | *r =* -0.3,p = 0.253, FDR = 0.43 | *r =* 0.429,p = 0.098, FDR = 0.31 | *r =* -0.3,p = 0.258, FDR = 0.66 | *r =* 0.502,p = 0.049, FDR = 0.14 |
| **Span Ratio** | *r =* 0.155,p = 0.563, FDR = 0.77 | *r =* 0.12, p = 0.656, FDR = 0.78 | *r =* 0.282,p = 0.288, FDR = 0.45 | *r =* 0.079,p = 0.771, FDR = 0.89 | *r =* -0.3,p = 0.243, FDR = 0.4 |
| **Max Span**  **Across Hull** | *r =* 0.258,p = 0.331, FDR = 0.63 | *r =* 0.361,p = 0.169, FDR = 0.43 | *r =* -0.34,p = 0.196, FDR = 0.44 | *r =* 0.279,p = 0.293, FDR = 0.66 | *r =* -0.42,p = 0.101, FDR = 0.23 |
| **Area** | *r =* 0.252,p = 0.343, FDR = 0.63 | *r =* 0.376,p = 0.151, FDR = 0.43 | *r =* -0.3,p = 0.248, FDR = 0.44 | *r =* 0.205,p = 0.442, FDR = 0.661 | *r =* -0.36,p = 0.169, FDR = 0.29 |
| **Perimeter** | *r =* 0.252,p = 0.343, FDR = 0.63 | *r =* 0.364,p = 0.165, FDR = 0.43 | *r =* -0.33,p = 0.204, FDR = 0.44 | *r =* 0.252,p = 0.343, FDR = 0.661 | *r =* -0.39,p = 0.128, FDR = 0.24 |
| **Circularity** | *r =* -0.26,p = 0.326, FDR = 0.63 | *r =* -0.12,p = 0.656, FDR = 0.78 | *r =* -0.29,p = 0.263, FDR = 0.44 | *r =* -0.21,p = 0.416, FDR = 0.66 | *r =* 0.535, p = 0.034, FDR = 0.12 |
| **Mean Radius** | *r =* 0.27,p = 0.309, FDR = 0.63 | *r =* 0.391,p = 0.135, FDR = 0.43 | *r =* -0.32,p = 0.221, FDR = 0.44 | *r =* 0.247,p = 0.355, FDR = 0.661 | *r =* -0.41, p = 0.111, FDR = 0.23 |
| **ODI** | *r =* -0.07,p = 0.771, FDR = 0.88 | *r =* 0.158,p = 0.556, FDR = 0.76 | *r =* 0.229,p = 0.391, FDR = 0.5 | *r =* -0.14,p = 0.586, FDR = 0.8 | *r =* 0.25,p = 0.349, FDR = 0.52 |
| **ICVF** | *r =* -0.07,p = 0.792, FDR = 0.88 | *r =* 0.214,p = 0.442, FDR = 0.66 | *r =* 0.628,p = 0.014, FDR = 0.15 | *r =* -0.25,p = 0.367, FDR = 0.66 | *r =* -0.007, p = 0.984, FDR = 0.98 |
| **ISVF** | *r =* -0.01,p = 0.969, FDR = 1 | *r =* -0.11,p = 0.672, FDR = 0.78 | *r =* 0.126,p = 0.64, FDR = 0.78 | *r =* -0.05,p = 0.847, FDR = 0.89 | *r =* -0.006,p = 0.986, FDR = 0.98 |
| **LJ** | *r =* -0.27, p = 0.293, FDR = 0.63 | *r =* -0.08,p = 0.746, FDR = 0.82 | *r =* 0.267,p = 0.315, FDR = 0.46 | *r =* -0.63,p = 0.01, FDR = 0.15 | *r =* 0.426, p = 0.101, FDR = 0.23 |
| **FA** | *r =* 0.082, *p =* 0.762, FDR = 0.88 | *r =* 0.302, *p =* 0.253, FDR = 0.43 | *r =* 0.617, *p =* 0.012, FDR = 0.15 | *r =* -0.22, *p =* 0.391, FDR = 0.66 | *r =* -0.14, *p =* 0.586, FDR = 0.82 |
| **MD** | *r =* -0.18, *p =* 0.495, FDR = 0.77 | *r =* -0.32, *p =* 0.222, FDR = 0.43 | *r =* -0.43, *p =* 0.09, FDR = 0.31 | *r =* -0.06, *p =* 0.811, FDR = 0.89 | *r =* 0.027, *p =* 0.919, FDR = 0.98 |
| **RD** | *r =* -0.19, *p =* 0.461, FDR = 0.7 | *r =* -0.33, *p =* 0.198, FDR = 0.43 | *r =* -0.44, *p =* 0.085, FDR = 0.31 | *r =* -0.06, *p =* 0.819, FDR = 0.89 | *r =* 0.064, *p =* 0.811, FDR = 0.98 |
| **AD** | *r =* -0.15, *p =* 0.553, FDR = 0.77 | *r =* -0.3, p = 0.244, FDR = 0.43 | *r =* -0.4, p = 0.118, FDR = 0.32 | *r =* -0.05, *p =* 0.828, FDR = 0.89 | *r =* 0.008, *p =* 0.975, FDR = 0.98 |
| ***Ruminococcus*** |  | *r =* 0.629, *p =* 0.01, FDR = 0.23 | *r =* 0.067, *p =* 0.805, FDR = 0.9 | *r =* 0.376, *p =* 0.151, FDR = 0.55 | *r =* -0.56, *p =* 0.024, FDR = 0.11 |
| ***Muribaculaceae*** | *r =* 0.629, *p =* 0.01, FDR = 0.11 |  | *r =* -0.02, *p =* 0.934, FDR = 0.98 | *r =* 0.202, *p =* 0.449, FDR = 0.66 | *r =* -0.55, *p =* 0.027, FDR = 0.11 |
| ***Ls NK4A136*** | *r =* 0.067, *p =* 0.805, FDR = 0.8 | *r =* -0.02, *p =* 0.934, FDR = 0.98 |  | *r =* -0.5,p = 0.047, FDR = 0.21 | *r =* 0.061, *p =* 0.822, FDR = 0.98 |
| ***Coprococcus*** | *r =* 0.376, *p =* 0.151, FDR = 0.631 | *r =* 0.202, *p =* 0.449, FDR = 0.6 | *r =* -0.5,p = 0.047, FDR = 0.2 |  | *r =* -0.55, *p =* 0.026, FDR = 0.11 |
| **Bilophila** | *r =* -0.56, *p =* 0.024, FDR = 0.18 | *r =* -0.55, *p =* 0.027, FDR = 0.3 | *r =* 0.061, *p =* 0.822, FDR = 0.9 | *r =* -0.55, *p =* 0.026, FDR = 0.19 |  |

***Supplementary Table 15*.** Statistical Information of Hippocampal proteins.

|  | **FKBP4** | **NRC31** | **NRC32** | **HSD11b1** | **IL-6** | **NF-κB** | **DUSP1** | **SV2A** |
| --- | --- | --- | --- | --- | --- | --- | --- | --- |
|  | Gene expression | Gene expression | Gene expression | Gene expression | Gene expression | Gene expression | Gene expression | Gene expression |
| Source of Variation | F Statistic, post-hoc | F Statistic, post-hoc | F Statistic, post-hoc | F Statistic, post-hoc | F Statistic, post-hoc | F Statistic, post-hoc | F Statistic, post-hoc | F Statistic, post-hoc |
| **Diet: Stress** | F(1, 8) = 0.35, p = 0.5690 | F(1, 8) = 1.6, p = 0.2407 | F(1, 8) = 0.38, p = 0.5514 | F(1, 8) = 1.01, p = 0.3434 | F(1, 8) = 0.15, p = 0.7056 | F(1, 8) = 0.7 p = 0.4242 | F(1, 8) = 1.24, p = 0.2981 | F(1, 8) = 0.4, p = 0.5432 |
| **Stress** | F(1, 8) = 0.13, p = 0.7204 | F(1, 8) = 0.14, p = 0.7160 | F(1, 8) = 0.32, p = 0.5858 | F(1, 8) = 3.84, p = 0.0856 | F(1, 8) = 1.9, p = 0.1999 | F(1, 8) = 0.34, p = 0.5753 | F(1, 8) = 1.3, p = 0.2813 | F(1, 8) = 0.14, p = 0.7131 |
| **Diet** | F(1, 8) = 0.6, p = 0.4587 | F(1, 8) = 0.19, p = 0.6692 | F(1, 8) = 2.49, p = 0.1526 | F(1, 8) = 0.25, p = 0.6297 | F(1, 8) = 2.2, p = 0.1731 | F(1, 8) = 0.81, p = 0.3941 | F(1, 8) = 1.1, p = 0.3076 | F(1, 8) = 1.3, p = 0.2861 |

***Supplementary Table 16A*.** Statistical Information of multiple comparisons in FKBP5 regions 5-Upstream CpG.

| Region | CpG# | CD UNEXP vs CD EXP | CD UNEXP vs WD UNEXP | CD UNEXP vs WD EXP | CD EXP vs WD UNEXP | CD EXP vs WD EXP | WD UNEXP vs WD EXP |
| --- | --- | --- | --- | --- | --- | --- | --- |
| 5-Upstream | 545 | p = 0.8378 | p = 0.6831 | p = 0.98 | p = 0.21 | p = 0.9704 | p = 0.4381 |
| 5-Upstream | 544 | p = 0.9906 | p = 0.5495 | p = 0.7417 | p = 0.7369 | p = 0.8918 | p = 0.9899 |
| 5-Upstream | 543 | p = 0.7786 | p = 0.8258 | p = 0.9321 | p = 0.2704 | p = 0.4091 | p = 0.9939 |
| 5-Upstream | 542 | p = 0.9996 | p = 0.4779 | p = 0.9864 | p = 0.5443 | p = 0.9954 | p = 0.6931 |
| 5-Upstream | 468 | p = 0.9672 | p = 0.9971 | p = 0.2067 | p = 0.9939 | p = 0.443 | p = 0.2981 |
| 5-Upstream | 467 | p = 0.2516 | p = 0.9913 | p = 0.1414 | p = 0.4043 | p = 0.9913 | p = 0.2516 |
| 5-Upstream | 466 | p = 0.7321 | p = 0.895 | p = 0.2233 | p = 0.9882 | p = 0.8134 | p = 0.622 |
| 5-Upstream | 465 | p > 0.9999 | p = 0.673 | p = 0.448 | p = 0.6831 | p = 0.4381 | p = 0.0455 |
| 5-Upstream | 416 | p = 0.9927 | p = 0.9526 | p = 0.9812 | p = 0.9944 | p = 0.9996 | p = 0.9989 |
| 5-Upstream | 415 | p = 0.9985 | p = 0.9882 | p = 0.8982 | p = 0.9621 | p = 0.9505 | p = 0.7369 |
| 5-Upstream | 336 | p = 0.9899 | p = 0.8607 | p = 0.7128 | p = 0.9639 | p = 0.5186 | p = 0.2516 |
| 5-Upstream | 335 | p = 0.9913 | p = 0.7963 | p = 0.5289 | p = 0.9243 | p = 0.3536 | p = 0.106 |
| 5-Upstream | 334 | p = 0.9977 | p = 0.9565 | p = 0.7786 | p = 0.8982 | p = 0.8715 | p = 0.4628 |
| 5-Upstream | 333 | p = 0.9995 | p = 0.8918 | p = 0.8784 | p = 0.9321 | p = 0.8258 | p = 0.4628 |
| 5-Upstream | 332 | p = 0.9812 | p = 0.9505 | p = 0.698 | p = 0.7963 | p = 0.895 | p = 0.3671 |
| 5-Upstream | 331 | p = 0.7369 | p = 0.9996 | p = 0.6578 | p = 0.7963 | p = 0.9992 | p = 0.7225 |
| 5-Upstream | **301** | p = 0.1594 | p = 0.5443 | **p = 0.0084** | **p = 0.0038** | p = 0.6931 | **p < 0.0001** |
| 5-Upstream | **300** | p = 0.2199 | p = 0.29 | **p = 0.0148** | **p = 0.0014** | p = 0.703 | **p < 0.0001** |
| 5-Upstream | **299** | p = 0.6881 | p = 0.2199 | p = 0.2821 | **p = 0.0137** | p = 0.9043 | **p = 0.0013** |
| 5-Upstream | **298** | p = 0.8134 | p = 0.5858 | p = 0.3063 | p = 0.139 | p = 0.8298 | **p = 0.0148** |
| 5-Upstream | 49 | p > 0.9999 | p > 0.9999 | p = 0.9998 | p > 0.9999 | p > 0.9999 | p > 0.9999 |
| 5-Upstream | 48 | p = 0.9295 | p = 0.8338 | p > 0.9999 | p = 0.9954 | p = 0.9346 | p = 0.8417 |
| 5-Upstream | 47 | p = 0.9954 | p = 0.9971 | p = 0.9994 | p = 0.9719 | p = 0.9844 | p = 0.9998 |
| 5-Upstream | 22 | p = 0.9947 | p = 0.127 | p = 0.5784 | p = 0.1177 | p = 0.506 | p = 0.8383 |
| 5-Upstream | 21 | p > 0.9999 | p > 0.9999 | p = 0.7512 | p > 0.9999 | p = 0.8134 | p = 0.787 |
| 5-Upstream | **20** | **p = 0.0004** | p = 0.73 | p = 0.2008 | p = 0.0546 | p = 0.3758 | p = 0.8285 |
| 5-Upstream | 5 | p = 0.9927 | p > 0.9999 | p = 0.9882 | p = 0.9963 | p > 0.9999 | p = 0.9933 |
| 5-Upstream | 4 | p = 0.9834 | p > 0.9999 | p = 0.9844 | p = 0.9899 | p = 0.8885 | p = 0.9762 |
| 5-Upstream | 3 | p = 0.8679 | p = 0.9216 | p = 0.8049 | p = 0.9991 | p = 0.9992 | p = 0.9933 |
| 5-Upstream | 2 | p = 0.9985 | p = 0.9882 | p = 0.9987 | p = 0.9985 | p > 0.9999 | p = 0.9982 |
| 5-Upstream | 1 | p = 0.9873 | p = 0.9189 | p = 0.9295 | p = 0.9899 | p = 0.9927 | p > 0.9999 |

***Supplementary Table 16B.*** Statistical Information of multiple comparisons in FKBP5 Exons by CpG.

| **Region** | **CpG#** | **CD UNEXP vs CD EXP** | **CD UNEXP vs WD UNEXP** | **CD UNEXP vs WD EXP** | **CD EXP vs WD UNEXP** | **CD EXP vs WD EXP** | **WD UNEXP vs WD EXP** |
| --- | --- | --- | --- | --- | --- | --- | --- |
| Exon 1 | 1 | p > 0.9999 | p = 0.9989 | p = 0.9899 | p = 0.9971 | p = 0.9944 | p = 0.9689 |
| Exon 1 | 2 | p = 0.9995 | p > 0.9999 | p = 0.9967 | p = 0.9991 | p = 0.9882 | p = 0.9977 |
| Exon 2 | 47 | p = 0.9873 | p = 0.9844 | p = 0.9995 | p = 0.9013 | p = 0.9704 | p = 0.9949 |
| Exon 2 | 48 | p = 0.9989 | p = 0.6013 | p > 0.9999 | p = 0.5083 | p = 0.9982 | p = 0.6168 |
| Exon 2 | 49 | p = 0.2166 | p = 0.9189 | p = 0.5806 | p = 0.0507 | p = 0.9132 | p = 0.2233 |
| Exon 2 | 50 | p = 0.1414 | p = 0.9949 | p = 0.3104 | p = 0.0811 | p = 0.9762 | p = 0.2003 |
| Exon 5 | 410 | p > 0.9999 | p > 0.9999 | p = 0.9913 | p > 0.9999 | p = 0.9933 | p = 0.9882 |
| Exon 5 | 411 | p > 0.9999 | p > 0.9999 | p > 0.9999 | p > 0.9999 | p > 0.9999 | p > 0.9999 |
| Exon 5 | 412 | p > 0.9999 | p = 0.9991 | p = 0.9967 | p = 0.9998 | p = 0.9933 | p = 0.9854 |
| Exon 5 | 413 | p = 0.9762 | p = 0.9891 | p = 0.8819 | p = 0.9997 | p = 0.9882 | p = 0.9748 |
| Exon 9 | 708 | p > 0.9999 | p > 0.9999 | p = 0.9915 | p > 0.9999 | p = 0.9946 | p = 0.9963 |
| Exon 9 | 709 | p > 0.9999 | p > 0.9999 | p = 0.9915 | p > 0.9999 | p = 0.9946 | p = 0.9963 |
| Exon 9 | 710 | p = 0.9822 | p = 0.9501 | p = 0.9832 | p = 0.9988 | p = 0.9065 | p = 0.8431 |
| Exon 9 | 711 | p = 0.9668 | p = 0.9987 | p = 0.9993 | p = 0.9421 | p = 0.9502 | p > 0.9999 |
| Exon 9 | 712 | p > 0.9999 | p = 0.9529 | p = 0.9472 | p = 0.9552 | p = 0.9502 | p > 0.9999 |
| Exon 9 | 713 | p = 0.9221 | p = 0.9779 | p = 0.9981 | p = 0.9968 | p = 0.9756 | p = 0.9968 |
| Exon 9 | 714 | p = 0.9827 | p = 0.9981 | p = 0.9973 | p = 0.9599 | p = 0.9552 | p > 0.9999 |
| Exon 9 | 715 | p > 0.9999 | p = 0.9994 | p > 0.9999 | p = 0.9998 | p > 0.9999 | p > 0.9999 |

**Supplementary Table 16C.** Statistical Information of multiple comparisons in FKBP5 Introns by CpG.

| **Region** | **CpG#** | **CD UNEXP vs CD EXP** | **CD UNEXP vs WD UNEXP** | **CD UNEXP vs WD EXP** | **CD EXP vs WD UNEXP** | **CD EXP vs WD EXP** | **WD UNEXP vs WD EXP** |
| --- | --- | --- | --- | --- | --- | --- | --- |
| Intron 2 | 123 | p = 0.9775 | p = 0.9711 | p = 0.9987 | p = 0.9998 | p = 0.944 | p = 0.9381 |
| Intron 2 | 124 | p = 0.9989 | p = 0.9676 | p = 0.998 | p = 0.9876 | p > 0.9999 | p = 0.9905 |
| Intron 2 | 125 | p = 0.9982 | p = 0.993 | p = 0.9906 | p = 0.9994 | p = 0.9992 | p > 0.9999 |
| Intron 4 | 218 | p = 0.3949 | p = 0.9639 | p = 0.8338 | p = 0.6931 | p = 0.8852 | p = 0.9834 |
| Intron 4 | 219 | p = 0.1649 | p = 0.3447 | p = 0.1439 | p = 0.9775 | p > 0.9999 | p = 0.9656 |
| Intron 4 | 220 | p = 0.9656 | p = 0.8852 | p = 0.9762 | p = 0.9939 | p > 0.9999 | p = 0.9891 |
| Intron 4 | 221 | p > 0.9999 | p = 0.565 | p = 0.9321 | p = 0.5806 | p = 0.9394 | p = 0.8982 |
| Intron 4 | 222 | p = 0.8679 | p = 0.6065 | p = 0.9216 | p = 0.188 | p = 0.4981 | p = 0.9321 |
| Intron 4 | 242 | p = 0.9269 | p = 0.8532 | p = 0.998 | p = 0.9977 | p = 0.857 | p = 0.7604 |
| Intron 4 | 243 | p = 0.7225 | p = 0.8258 | p > 0.9999 | p = 0.9977 | p = 0.7369 | p = 0.8378 |
| Intron 4 | 244 | p = 0.998 | p = 0.9321 | p = 0.9967 | p = 0.9748 | p = 0.98 | p = 0.8494 |
| Intron 4 | 245 | p = 0.8338 | p = 0.9013 | p = 0.9505 | p = 0.9987 | p = 0.5135 | p = 0.6117 |
| Intron 4 | 246 | p = 0.9394 | p = 0.8049 | p = 0.9994 | p = 0.9882 | p = 0.9689 | p = 0.8643 |
| Intron 4 | 296 | p = 0.9775 | p = 0.8494 | p = 0.9526 | p = 0.9775 | p = 0.9994 | p = 0.992 |
| Intron 4 | 297 | p = 0.857 | p > 0.9999 | p = 0.9800 | p = 0.8417 | p = 0.9775 | p = 0.9748 |
| Intron 4 | 298 | p > 0.9999 | p > 0.9999 | p = 0.9997 | p > 0.9999 | p = 0.9992 | p = 0.9985 |
| Intron 4 | 299 | p = 0.9992 | p > 0.9999 | p > 0.9999 | p > 0.9999 | p = 0.9999 | p > 0.9999 |
| Intron 4 | 300 | p = 0.9997 | p = 0.9992 | p > 0.9999 | p > 0.9999 | p > 0.9999 | p > 0.9999 |
| Intron 4 | 301 | p = 0.9762 | p = 0.9992 | p = 0.9788 | p = 0.9484 | p = 0.8494 | p = 0.9933 |
| **Intron 4** | **345** | p = 0.6629 | p = 0.8679 | **p = 0.0235** | p = 0.2233 | p = 0.3231 | **p = 0.0018** |
| **Intron 4** | **346** | p = 0.8643 | p = 0.9991 | p = 0.0529 | p = 0.7963 | p = 0.286 | **p = 0.0365** |
| Intron 4 | 347 | p = 0.9462 | p = 0.937 | p = 0.1101 | p > 0.9999 | p = 0.3231 | p = 0.3404 |
| Intron 5 | 433 | p = 0.9864 | p = 0.9844 | p = 0.6425 | p = 0.8982 | p = 0.8378 | p = 0.4187 |
| Intron 5 | 434 | p = 0.9565 | p = 0.565 | p = 0.9967 | p = 0.2704 | p = 0.8885 | p = 0.698 |
| Intron 5 | 435 | p = 0.9672 | p = 0.9873 | p = 0.9462 | p = 0.857 | p = 0.9997 | p = 0.8134 |
| Intron 5 | 524 | p = 0.8258 | p = 0.7369 | p = 0.9844 | p = 0.9985 | p = 0.6168 | p = 0.5135 |
| Intron 5 | 525 | p = 0.9043 | p = 0.765 | p = 0.7079 | p = 0.9913 | p = 0.2981 | p = 0.1733 |
| Intron 5 | 526 | p = 0.3902 | p = 0.9526 | p > 0.9999 | p = 0.7177 | p = 0.3902 | p = 0.9526 |
| Intron 5 | 527 | p = 0.3536 | p = 0.106 | p > 0.9999 | p = 0.9243 | p = 0.336 | p = 0.0984 |
| Intron 6 | 552 | p > 0.9999 | p = 0.9639 | p = 0.2199 | p = 0.9565 | p = 0.2337 | p = 0.078 |
| Intron 6 | 553 | p = 0.8134 | p = 0.8217 | p = 0.6781 | p > 0.9999 | p = 0.188 | p = 0.1941 |
| Intron 6 | 554 | p > 0.9999 | p = 0.9959 | p = 0.5961 | p = 0.9967 | p = 0.5858 | p = 0.4529 |
| Intron 7 | 612 | p = 0.9959 | p = 0.6972 | p = 0.992 | p = 0.5692 | p = 0.9565 | p = 0.8384 |
| Intron 7 | 613 | p = 0.9944 | p = 0.9993 | p = 0.8175 | p = 0.9996 | p = 0.9243 | p = 0.9121 |
| Intron 7 | 614 | p = 0.9132 | p = 0.999 | p = 0.9882 | p = 0.9704 | p = 0.9873 | p = 0.9988 |
| Intron 7 | 615 | p = 0.9854 | p = 0.7956 | p = 0.9997 | p = 0.9303 | p = 0.9704 | p = 0.747 |
| Intron 7 | 616 | p = 0.9864 | p = 0.8841 | p = 0.9992 | p = 0.9737 | p = 0.9656 | p = 0.8312 |
| Intron 7 | 617 | p = 0.9982 | p = 0.9915 | p = 0.9992 | p = 0.9991 | p = 0.9906 | p = 0.9774 |
| Intron 7 | 618 | p > 0.9999 | p = 0.9381 | p = 0.9584 | p = 0.952 | p = 0.9704 | p = 0.9992 |
| Intron 7 | 619 | p = 0.9944 | p = 0.9832 | p = 0.9999 | p = 0.999 | p = 0.998 | p = 0.9905 |
| Intron 7 | 655 | p > 0.9999 | p = 0.9982 | p = 0.9997 | p = 0.9959 | p > 0.9999 | p = 0.9939 |
| Intron 7 | 656 | p = 0.9834 | p = 0.9639 | p = 0.9216 | p = 0.9995 | p = 0.9933 | p = 0.9987 |
| Intron 8 | 687 | p > 0.9999 | p > 0.9999 | p = 0.336 | p > 0.9999 | p = 0.3671 | p = 0.3671 |
| Intron 8 | 688 | p > 0.9999 | p = 0.9974 | p = 0.2166 | p = 0.9992 | p = 0.2443 | p = 0.3063 |
| Intron 8 | 689 | p = 0.9959 | p > 0.9999 | p = 0.336 | p = 0.9933 | p = 0.4678 | p = 0.3146 |
| Intron 8 | 690 | p = 0.9963 | p = 0.9994 | p = 0.2781 | p = 0.9997 | p = 0.185 | p = 0.2233 |
| Intron 8 | 691 | p > 0.9999 | p = 0.9963 | p = 0.5135 | p = 0.9949 | p = 0.5289 | p = 0.3809 |
| Intron 9 | 716 | p = 0.9902 | p = 0.9997 | p = 0.9978 | p = 0.9976 | p = 0.9995 | p = 0.9998 |
| Intron 9 | 717 | p = 0.7875 | p = 0.9837 | p = 0.9055 | p = 0.9654 | p = 0.9987 | p = 0.9917 |
| Intron 9 | 718 | p = 0.7963 | p = 0.8522 | p = 0.6499 | p > 0.9999 | p = 0.9859 | p = 0.9879 |

***Supplementary Table 17*.** Statistical Information of methylation in FKBP5 by region.

| **Factor** | **WD** | **PSS** | **WD x PSS** |
| --- | --- | --- | --- |
| **5-Upstream** | p = 0.9100 | **p = 0.0001** | **p = 0.0090** |
| **Exon 1** | p = 0.63 | p = 0.54 | p = 0.47 |
| **Exon 2** | p = 0.48 | p = 0.12 | p = 0.89 |
| **Intron 4** | p = 0.621 | **p = 0.003** | p = 0.123 |
| **Exon 5** | p = 0.47 | p = 0.27 | p = 0.60 |
| **Exon 9** | p = 0.10 | p = 0.53 | p = 0.60 |
| **Intron 2** | p = 0.47 | p = 0.83 | p = 0.19 |
| **Intron 4** | p = 0.62 | **p = 0.003** | p = 0.12 |
| **Intron 5** | p = 0.08 | p = 0.51 | p = 0.53 |
| **Intron 6** | **p = 0.031** | p = 0.103 | p = 0.236 |
| **Intron 7** | p = 0.211 | p = 0.914 | **p = 0.032** |
| **Intron 8** | **p = 0.020** | **p = 0.016** | **p = 0.024** |
| **Intron 9** | p = 0.44 | p = 0.25 | p = 0.27 |

***Supplementary Table 18.*** Statistical Information of HCM3 treated gene and concentration of IL-6.

|  | **IL-6**  Gene expression | **IL-6**  concentration |
| --- | --- | --- |
| Factor | F Statistic, post-hoc | F Statistic, post-hoc |
| **CORT:PA** | **F(1, 12) = 68.10, p < 0.0001** | **F(1, 12) = 8.258, *p =* 0.0140** |
| CORT | F(1, 12) = 0.286, *p =* 0.6022 | **F(1, 12) = 14.96, *p =* 0.0022** |
| PA | **F(1, 12) = 5.852, *p =* 0.0324** | F(1, 12) = 1.630, *p =* 0.2259 |
| VEH:VEH vs VEH:PA | ***p =* 0.0067** | *p =* 0.6794 |
| VEH:VEH vs CORT:VEH | ***p =* 0.0002** | *p =* 0.8938 |
| VEH:VEH vs CORT:PA | *p =* 0.5615 | ***p =* 0.0155** |
| VEH:PA vs CORT:VEH | *p =* 0.2113 | *p =* 0.3059 |
| **VEH:PA vs CORT:PA** | ***p =* 0.0007** | ***p =* 0.0022** |
| CORT:VEH vs CORT:PA | ***p =* < 0.0001** | *p =* 0.053 |

***Supplementary Table 19*.** Statistical Information of HMC3 FKBP5 SiRNA and GE of IL-6.

|  | **FKBP5 siRNA** | **IL-6**  Gene expression | **IL-6**  concentration |
| --- | --- | --- | --- |
| Factor | t test | F Statistic, post-hoc | F Statistic, post-hoc |
| FKBP5 siRNA vs VEH | **t(58)= 3.559, *p =* 0.0074** | -- | -- |
| Treatment:SiRNA |  | **F(3, 24) = 36.64, p < 0.0001** | **F(3, 24) = 9.580, *p =* 0.0002** |
| Treatment |  | **F(3, 24) = 19.95, p < 0.0001** | **F(3, 24) = 18.78, p < 0.0001** |
| SiRNA |  | **F(1, 24) = 54.49, p < 0.0001** | **F(1, 24) = 48.34, p < 0.0001** |
| VEH:CTRL siRNA vs VEH:FKBP5 siRNA |  | *p =* 0.9987 | *p =* > 0.9999 |
| VEH:CTRL siRNA vs Cort:CTRL siRNA |  | ***p =* 0.002** | *p =* 0.8974 |
| VEH:CTRL siRNA vs Cort:FKBP5 siRNA |  | *p =* 0.4771 | *p =* > 0.9999 |
| VEH:CTRL siRNA vs PA:CTRL siRNA |  | *p =* 0.0755 | ***p =* 0.0022** |
| VEH:CTRL siRNA vs PA:FKBP5 siRNA |  | *p =* 0.2451 | *p =* 0.6464 |
| VEH:CTRL siRNA vs PA Cort:CTRL siRNA |  | ***p =* < 0.0001** | ***p =* < 0.0001** |
| VEH:CTRL siRNA vs PA Cort:FKBP5 siRNA |  | ***p =* 0.0252** | *p =* > 0.9999 |
| VEH:FKBP5 siRNA vs Cort:CTRL siRNA |  | ***p =* 0.0084** | *p =* 0.3565 |
| VEH:FKBP5 siRNA vs Cort:FKBP5 siRNA |  | *p =* 0.8249 | *p =* > 0.9999 |
| VEH:FKBP5 siRNA vs PA:CTRL siRNA |  | ***p =* 0.0207** | ***p =* 0.0003** |
| VEH:FKBP5 siRNA vs PA:FKBP5 siRNA |  | *p =* 0.5624 | *p =* 0.168 |
| VEH:FKBP5 siRNA vs PA Cort:CTRL siRNA |  | ***p =* < 0.0001** | ***p =* < 0.0001** |
| VEH:FKBP5 siRNA vs PA Cort:FKBP5 siRNA |  | *p =* 0.0904 | *p =* > 0.9999 |
| Cort:CTRL siRNA vs Cort:FKBP5 siRNA |  | *p =* 0.1888 | *p =* 0.3058 |
| Cort:CTRL siRNA vs PA:CTRL siRNA |  | ***p =* < 0.0001** | *p =* 0.1971 |
| Cort:CTRL siRNA vs PA:FKBP5 siRNA |  | *p =* 0.3902 | *p =* > 0.9999 |
| Cort:CTRL siRNA vs PA Cort:CTRL siRNA |  | ***p =* < 0.0001** | ***p =* < 0.0001** |
| Cort:CTRL siRNA vs PA Cort:FKBP5 siRNA |  | *p =* 0.9581 | *p =* 0.9144 |
| Cort:FKBP5 siRNA vs PA:CTRL siRNA |  | ***p =* 0.0006** | ***p =* 0.0003** |
| Cort:FKBP5 siRNA vs PA:FKBP5 siRNA |  | *p =* 0.9998 | *p =* 0.1402 |
| Cort:FKBP5 siRNA vs PA Cort:CTRL siRNA |  | ***p =* < 0.0001** | ***p =* < 0.0001** |
| Cort:FKBP5 siRNA vs PA Cort:FKBP5 siRNA |  | *p =* 0.7576 | *p =* > 0.9999 |
| PA:CTRL siRNA vs PA:FKBP5 siRNA |  | ***p =* 0.0002** | *p =* 0.4069 |
| PA:CTRL siRNA vs PA Cort:CTRL siRNA |  | ***p =* 0.0034** | *p =* 0.0962 |
| PA:CTRL siRNA vs PA Cort:FKBP5 siRNA |  | ***p =* < 0.0001** | ***p =* 0.0025** |
| PA:FKBP5 siRNA vs PA Cort:CTRL siRNA |  | ***p =* < 0.0001** | ***p =* 0.0002** |
| PA:FKBP5 siRNA vs PA Cort:FKBP5 siRNA |  | *p =* 0.9442 | *p =* 0.6764 |
| PA Cort:CTRL siRNA vs PA Cort:FKBP5 siRNA |  | ***p =* < 0.0001** | ***p =* < 0.0001** |

***Supplementary Table 20*.** Statistical Information of FKBP5 SiRNA.

|  | **FKBP5** | **FKBP5** |
| --- | --- | --- |
|  | SiRNA | SiRNA Gene Expression |
| Source of Variation | t test | F Statistic, post-hoc |
| **FKBP5 siRNA vs Control siRNA** | **t=5.259, df=6, p = 0.0019** | **---** |
| **Interaction** |  | **F(1, 20) = 6.703, p =0.0175** |
| **Stress** |  | F(1, 20) = 3.456, p =0.0778 |
| **Diet** |  | F(1, 20) = 1.550, p =0.2275 |
